# An anatomy-constrained transcriptomic screen identifies a 19-gene panel with an ordered regional similarity pattern in the human somatosensory system

**DOI:** 10.64898/2026.10.13.758993

**Authors:** Gerardo A. Boni, Alejandro Luque Hernández, Vivien Iran Bautista Ruedas

## Abstract

Neuroanatomical pathways link specialized brain regions, but whether regional transcriptomic profiles preserve relationships aligned with pathway order remains unclear. We defined candidate pathways and their available tissue representations from neuroanatomical literature, then screened 10,262 genes in the Body Index of Gene Expression (BIGE), a human microarray resource covering cells and tissues. The screen required expression across every mapped region of a route and limited single-region dominance. It identified 23 candidates in three routes with sufficient coverage; 19 met the criteria for the dorsal column-medial lemniscus (DCML) representation and formed the fixed panel. We evaluated this panel in six Allen Human Brain Atlas (AHBA) donors across three central regional representations: an aggregate of samples assigned to the cuneate and gracile nuclei (Cu/Gr), the lateral group of nuclei, ventral division, within the dorsal thalamus (DTLv), and postcentral gyrus. The regional-order contrast compared the mean similarity of the pairs adjacent in the anatomical sequence, Cu/Gr-DTLv and DTLv-postcentral gyrus, with Cu/Gr-postcentral gyrus similarity. Both adjacent-in-order correlations were positive in all six donors; the contrast was positive in five and reached a mean of T = 0.345175 (exact sign test, P = 0.109375). The panel exceeded two distributions of 50,000 gene sets generated under comparable selection constraints (P = 0.027919 and P = 0.026579). In descriptive anatomical comparisons, it ranked third among 240 combinations and first among the target and all 16 single-region substitutions. All 50,000 one-probe-per-gene analyses retained T > 0, and broad neuronal-expression contrasts were positive in medulla, thalamus, and cortex. Spatial support depended on the null model, with stronger evidence under whole-donor surrogates (P = 0.003860) than under the primary within-compartment model (P = 0.051179). These findings identify a fixed gene panel whose regional similarity is aligned with the represented central order of the DCML system and define a specific hypothesis for prospective evaluation in an independent, anatomically resolved dataset.

## Introduction

Classical neuroanatomy established specialized brain regions and the projections that relate them. Network neuroscience expanded this view by treating those regions as components of distributed, multiscale systems whose organization depends on local properties and interregional relationships [1]. In the human cortex, coordinated gene-expression patterns are associated with functional networks and macroscale anatomical hierarchies [2,3]. This evidence raises a complementary question for sensory systems that span subcortical and cortical structures: whether the anatomical order of their components is also reflected in regional transcriptomic relationships.

Regional transcriptomic atlases quantify ribonucleic acid (RNA) abundance in tissue samples and provide a molecular description complementary to projections, synapses, cell types, and signal direction. Similarity between regional expression profiles may reflect shared cellular composition, developmental programs, or spatial gradients. We therefore interpret transcriptomic similarity as regional molecular organization associated with an anatomical system, while treating cellular connectivity and causal direction as separate anatomical properties.

The dorsal column–medial lemniscus (DCML) system provides a well-described anatomical sequence. Somatosensory afferents reach the cuneate and gracile nuclei; second-order neurons project toward the somatosensory thalamus, and thalamocortical projections reach the somatosensory cortex of the postcentral gyrus [4,5]. In the Allen Human Brain Atlas (AHBA), this study represented the available central segment with Cu/Gr, an analysis-defined aggregate of samples assigned to the cuneate nucleus (Cu) and gracile nucleus (Gr); DTLv, the lateral group of nuclei, ventral division, within the dorsal thalamus, used as a regional approximation of the somatosensory thalamic stage; and postcentral gyrus, a gyral-level cortical representation. Cu/Gr–DTLv and DTLv–postcentral gyrus are adjacent in the anatomical sequence because they occupy positions 1–2 and 2–3. Cu/Gr–postcentral gyrus compares positions 1 and 3. These terms describe predefined order rather than compass direction, physical proximity, or connectivity between individual sampled cells.

An anatomy-guided discovery strategy can favor panels with particular expression properties, and spatial autocorrelation can produce similarity among nearby regions or regions embedded in the same gradient [6–10]. The study therefore compared the target with gene sets generated under similar selection constraints, alternative regional combinations, and surrogate maps that retained relevant spatial properties.

We evaluated 10,262 BIGE genes without a function- or disease-defined starting list. The screen identified 23 candidates in the three routes with sufficient coverage; 19 belonged to the DCML representation and were fixed before analysis in AHBA. We hypothesized that this fixed panel would show greater profile similarity between the pairs adjacent in the anatomical sequence than between the first and third regions. Donor-resolved AHBA data provided the primary regional evaluation, followed by selection-matched gene panels, anatomical comparisons, spatial surrogates, broad cell-class data, probe resampling, and reconstruction of Gene Expression Omnibus series GSE3526. Because the regional pattern was recognized in AHBA before the final robustness analyses, the study provides post-discovery characterization and a prospectively testable hypothesis.

## Methods

### Study design and inferential status

This study linked an anatomy-constrained discovery stage to a donor-resolved regional evaluation. A regional proxy was an available BIGE tissue label used to represent an anatomical component of a pathway. A nonexhaustive catalog of 16 candidate circuits was compiled from neuroanatomical literature. Five pathways were fixed before the gene-expression screen: the DCML pathway, descending pain modulation, a basal-ganglia motor loop, a cerebro-cerebellar loop, and a mesocorticolimbic reward loop. The first three had at least seven source samples for every regional proxy and entered the gene screen. The cerebro-cerebellar and reward loops were excluded because the pons and globus-pallidus proxies, respectively, each had one source sample. Supplementary Table S2 records the complete circuit catalog and screening decisions.

In BIGE, the DCML system was represented by dorsal root ganglion (n=8), medulla oblongata (n=9), thalamus (n=8), and parietal lobe (n=9). These broad tissue categories defined the screen. AHBA then evaluated the covered central segment using Cu/Gr, DTLv, and postcentral gyrus. Because AHBA does not include dorsal root ganglion samples, this was not a complete-pathway test.

The candidate pathways, regional mappings, and 23 candidate genes were defined before AHBA evaluation. The 19-gene DCML panel was therefore fixed with respect to AHBA, but the Cu/Gr–DTLv–postcentral pattern was recognized in AHBA before the final matched-panel and spatial analyses. Analysis outcomes and decision rules were then frozen in a post-discovery protocol. This sequence supports post-discovery characterization and a prospectively testable hypothesis rather than independent confirmation.

### BIGE source and anatomy-constrained gene screen

BIGE is a microarray resource described as containing expression information for more than 105 human cells and tissues [11–13]. Its documented source lineage includes studies of postmortem RNA quality and molecular relationships among 20 regions of the human central nervous system, including Gene Expression Omnibus series GSE3526 [11,12]. The processed neural matrix analyzed here contained aggregate regional means for 23 adult neural regions with adequate coverage and 10,262 reannotated genes. Individual-level observations, spatial coordinates, and donor covariates were unavailable in this matrix.

All 10,262 genes were evaluated without disease annotations or a preselected functional gene list. For each gene, expression was standardized across the 23 neural regions. Entry required a detection fraction of at least 0.50 in every regional proxy and a dominant-region share no greater than 0.50 of the summed positive regional contrast.

Candidate classes were assigned under frozen precedence rules. The route-wide class (B_ROUTEWIDE_90) required the lowest standardized expression among a pathway’s regions to exceed the 90th percentile of regions outside that pathway.

The balanced class (C_BALANCED) required the pathway minimum to exceed the off-route median by at least 0.50 standardized units and the pathway mean to exceed the off-route 90th percentile. A stricter route-exclusive class (A_EXCLUSIVE) required the pathway minimum to exceed the maximum off-route value; no gene met that criterion.

The screen yielded 23 unique candidates: 19 for DCML, comprising three route-wide and 16 balanced candidates; three balanced candidates for descending pain modulation; and one balanced candidate for the basal-ganglia pathway. The fixed DCML panel comprised *ATP2C1*, *RAI2*, *RAB3GAP2*, *CCT6A*, *MAP1B*, *SH3BGRL2*, *PDHX*, *C7orf25*, *TSHZ2*, *NEFM*, *ATXN3*, *ADAM23*, *HMGCS1*, *SLC16A6*, *NANS*, *IFT57*, *MEF2A*, *BBS7*, and *RAP1GDS1*. The screen selected genes with distributed expression across the mapped BIGE regions; the AHBA analysis tested relationships among regional profiles for the fixed panel.

Among the three eligible routes, DCML supplied the only candidate set large enough for a multigene profile-correlation analysis: 19 genes, compared with three candidates for descending pain modulation and one for the basal-ganglia pathway. The 19 DCML genes therefore became the fixed panel for the donor-resolved AHBA evaluation. The smaller candidate sets remained exploratory and were not subjected to the same panel-level contrast. This transition from screening to AHBA analysis was adaptive and is represented only partially by the subsequent matched-panel models.

### AHBA expression profiles and regional residuals

Microarray data from six adult AHBA donors were analyzed [6]. The source contained 3,702 anatomically annotated samples. Forty-four AHBA microarray probes with exact gene-symbol matches represented the 19 genes. Allen-provided probe z-score vectors were aligned to a frozen sample order. For genes represented by more than one probe, the primary profile used the median across probes.

The target mapping comprised 19 samples in the Cu/Gr analytical aggregate, 20 pooled DTLv samples, and 62 postcentral-gyrus samples across the six donors. Cu/Gr combines samples assigned to the cuneate and gracile nuclei. DTLv is the AHBA label for the lateral group of nuclei, ventral division, and should not be interpreted as an isolated measurement of the ventral posterolateral nucleus. Postcentral gyrus is a gyral-level cortical representation rather than an isolated cytoarchitectonic subdivision of primary somatosensory cortex. For each donor, region, and gene, a regional residual was calculated as the median value among target samples minus the median among non-target sibling samples from the same top-level anatomical compartment. This residual expresses regional differentiation from the surrounding compartment. Coverage varied across donors; donor 12876 contributed one target sample to Cu/Gr.

### Anatomical-order contrast

For each donor, Spearman correlations across the 19 genes were calculated for Cu/Gr with DTLv, DTLv with postcentral gyrus, and Cu/Gr with postcentral gyrus. The first two correlations represent consecutive positions in the predefined order; the third compares the first and third regions. The donor-level anatomical-order contrast was

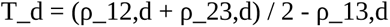

Here, d identifies the donor, and each rho is a Spearman correlation between two 19-gene regional profiles. A positive T_d indicates that the two consecutive pairs are more similar on average than the first-to-third pair. The statistic can be positive when some correlations are negative because it measures relative similarity. The reported group statistic, T, was the arithmetic mean of T_d across the six donors. A one-sided exact sign test summarized the donor-level directions.

### Gene panels matched to the selection process

The matched-panel analysis generated alternative 19-gene panels under constraints designed to approximate the anatomy-guided selection process. It enumerated 8,854 alternative four-region combinations among the 23 regions in the BIGE-derived extract and reapplied the frozen detection, dominance, threshold, and tier-precedence rules. The 23 historical candidates were excluded from control sampling. A common control universe contained 742 genes with complete AHBA regional residuals and broad neuronal-contrast data.

Each control gene was required to have the same exact-symbol AHBA probe count as the target gene it replaced. Matching also used eight prespecified variables: mean log2 expression across the 23 BIGE neural regions; regional standard deviation in BIGE; mean BIGE detection fraction; mean log2(normalized transcripts per million [nTPM]+1) across 193 Human Protein Atlas (HPA) subregions; standard deviation across those HPA subregions; the fraction of HPA subregions with nTPM greater than 1; median all-brain AHBA expression; and robust all-brain AHBA spread, calculated as the median absolute deviation multiplied by 1.4826. The current primary distribution additionally conditioned on the anatomy-screen selection class. The second distribution also matched the neuronal-minus-non-neuronal contrasts in medulla oblongata, thalamus, and cerebral cortex. Each model produced 50,000 accepted panels through an eligible alternative regional combination. These reference distributions test whether the target panel is unusual among panels obtainable under matched parts of the selection process. They do not reproduce every adaptive choice made during discovery and therefore do not estimate an unconditional discovery probability.

The historical technical comparator, retained only as a labeled reference for the cell-class and probe analyses, used 19 candidate-specific pools of 40 controls. Each draw selected one control from every pool, required 19 unique symbols, excluded all 23 original candidate genes, and matched exact AHBA probe count plus the same eight variables. It generated 50,000 panels but did not condition on anatomy-screen selection class or neuronal cell-class contrasts. It is therefore less restrictive than the two current selection-matched distributions and is not interchangeable with them.

### Decomposition of the regional similarity pattern

Four diagnostic components were frozen after recognition of the pattern and before generation of the matched-panel distributions: the mean of the two consecutive-pair correlations, the smaller consecutive-pair correlation within each donor, the number of donors in which both consecutive-pair correlations were positive, and first-to-third similarity. Each component was evaluated under both matched-panel models. Holm correction controlled the four component tests within each model. These tests localize the basis of the aggregate result and retain diagnostic rather than confirmatory status.

### Comparison with alternative regional combinations

The target was compared descriptively with 239 alternative three-region combinations drawn from comparable brainstem, dorsal-thalamic, and parietal compartments. The Cu/Gr-DTLv-postcentral value was ranked in the complete set of 240 combinations. This comparison was designed after inspection of the target pattern and does not provide an inferential test of anatomical specificity.

Two post hoc descriptive analyses assessed whether the target value depended on favorable individual regions or on their particular combination. The local comparison comprised the target and every combination obtained by replacing exactly one region: seven brainstem substitutions, five thalamic substitutions, and four parietal substitutions. An exploratory balanced additive decomposition then predicted each combination’s contrast from the marginal means associated with its three regional identities across the complete 8 by 6 by 5 grid. The difference between the observed and additive expected contrast was ranked across all 240 combinations.

All 240 combinations were also screened for positive correlations at both consecutive pairs in every donor. They were ranked by the anatomical-order contrast and by the consecutive-pair bottleneck, defined as the smaller of the two mean consecutive correlations. The two combinations with composite contrasts above the target were decomposed into their mean first consecutive-pair correlation, second consecutive-pair correlation, first-to-third correlation, and donor counts with both consecutive correlations positive. These local, additive, and component analyses were designed after inspection of the full anatomical ranking. Their ranks and fractions are descriptive and do not carry confirmatory probability interpretations.

### Spatially constrained surrogate maps

Spatial calibration used nuisance-restricted joint Moran spectral randomization adapted from Wagner and Dray [7]. This method generates surrogate expression maps that retain specified spatial properties of the observed data. A common spectral-sign vector was applied to all 19 genes within each fitted block. Up to numerical tolerance, the surrogates preserved fitted nuisance trends, residual Moran spectral power, and residual cross-gene covariance. They did not preserve every higher-order feature of each gene’s distribution.

The spatial decision rule was frozen after recognition of the pattern and before surrogate generation. It used two co-primary models and required both to pass. The within-compartment model fitted 18 donor-by-compartment blocks across 540 samples. It used a symmetric union graph of five nearest neighbors, inverse native-coordinate distance weights, and nuisance terms for the intercept and native x, y, and z coordinates. The whole-donor model fitted six blocks across 2,797 left-hemisphere or unpaired samples. It used a six-nearest-neighbor graph and added top-level anatomical fixed effects. Each co-primary model used 50,000 surrogate datasets. A component passed when its one-sided empirical P value was no greater than 0.05 and the observed statistic was strictly above the model’s 95th percentile. The joint spatial conclusion required both components to pass.

Four sensitivity models, also frozen in the post-discovery protocol, each used 10,000 surrogates. They varied dense-distance weights, whole-donor versus within-compartment scope, and coordinate-axis standardization within donor. Their individual numerical results describe model sensitivity and do not replace the co-primary decision. Leave-one-donor-out analyses recalculated observed and calibrated results after each donor was omitted. The resulting five-donor sets overlap and are treated as sensitivity analyses.

### Broad cell-class evidence

Broad neuronal attribution used Human Protein Atlas (HPA) summaries derived from the adult Human Brain Cell Atlas single-nucleus dataset [14,15]. For each gene, neuronal-minus-non-neuronal expression was summarized in medulla oblongata, thalamus, and cerebral cortex. The panel statistic was the smallest of these three regional contrasts. It was evaluated against the historical technical comparator and against the current, more restrictive selection-matched distribution. The regional labels are broader than Cu/Gr, DTLv, and postcentral gyrus; the analysis therefore addresses general neuronal attribution rather than cell types at the three target regions.

A separate post hoc HPA analysis calculated the anatomical-order contrast from the closest available regional summaries. It provides a cross-platform descriptive comparison and carries no independent decision status.

### Probe and resampling diagnostics

Probe dependence was examined through 50,000 random selections of one AHBA probe per gene. We recorded the proportion of contrasts above zero and the proportion above the historical technical-null 95th percentile. A two-way gene-by-donor bootstrap with 50,000 finite draws quantified directional stability under simultaneous resampling of genes and donors. The percentile range from this bootstrap is a diagnostic uncertainty summary rather than a confirmatory confidence interval.

### Reconstruction of GSE3526

Gene Expression Omnibus (GEO) Series accession GSE3526 belongs to the documented historical source lineage of BIGE [12]. The public series contains 353 Affymetrix Human Genome U133 Plus 2.0 microarrays from 65 normal tissue types and ten donors. The reconstruction used 39 historically mapped probe sets for the 19-gene panel. Linear-scale robust multi-array average intensities were transformed as log2(x+1). Probe values were collapsed separately by median, mean, and the historically selected representative probe. Each gene was then standardized within donor across a frozen neural-tissue background.

Eight donors had complete measurements for medulla, thalamus, parietal lobe, and dorsal root ganglion (DRG) and formed the donor-level analytical set. For a three-region sequence A–B–C, the contrast was T₃ = (ρAB + ρBC)/2 − ρAC. For a four-region sequence A–B–C–D, the contrast was T₄ = (ρAB + ρBC + ρCD)/3 − (ρAC + ρAD + ρBD)/3. In the four-region sequence, A, B, C, and D represented DRG, medulla, thalamus, and parietal lobe, respectively. These broad tissue labels differ from the more specific AHBA mapping. GSE3526 contributed to the historical source lineage of BIGE, and numerical equivalence between the public series and the processed discovery extract has not been established. The reconstruction is therefore a post-selection sensitivity analysis of the historical source cohort.

### Statistical reporting and computational reproducibility

Empirical upper-tail probabilities used the plus-one estimator, (1+b)/(1+N), where b is the number of null values greater than or equal to the observation among N draws. Exact reproduction of the frozen spatial counts used the numerical comparison null greater than or equal to observed minus 10^-15. Null 95th percentiles used linear interpolation.

Protocols, analysis outcomes, tails, random seeds, decision rules, executable hashes, inputs, and outputs were linked by cryptographic file-hash records. The final public release includes the frozen selection, AHBA, spatial-surrogate, probe-resampling, and GSE3526 analyses. Its packaged verification script returned PASS, confirming the archive manifest and the stated release boundary. Computational reproducibility of these analyses remains distinct from biological replication in new donors or a new atlas.

AI-assisted technical support. G.A.B. used OpenAI Codex in ChatGPT Work, with the model identified in the user interface as GPT-5.6 Sol (accessed in 2026), as a technical aid during the development, implementation, debugging, and audit of selected computational analyses and plotting code. The authors established the study questions, anatomical definitions, analytical specifications, inclusion decisions, and interpretation. All reported numerical results were generated from fixed inputs by archived executable scripts and verified through reproducible reruns and file-level integrity checks. The authors reviewed every retained output and take full responsibility for the methods, results, and conclusions.

## Results

### An anatomy-constrained screen defined the fixed 19-gene DCML panel

Among 10,262 genes screened across the three eligible pathways, 23 satisfied the route-wide or balanced multiregional criteria. Nineteen were assigned to the DCML representation, three to descending pain modulation, and one to the basal-ganglia pathway. *ATP2C1*, *RAI2*, and *RAB3GAP2* met the route-wide criterion. The remaining 16 DCML genes met the balanced criterion: *CCT6A*, *MAP1B*, *SH3BGRL2*, *PDHX*, *C7orf25*, *TSHZ2*, *NEFM*, *ATXN3*, *ADAM23*, *HMGCS1*, *SLC16A6*, *NANS*, *IFT57*, *MEF2A*, *BBS7*, and *RAP1GDS1*. All 19 satisfied the common detection and single-region-dominance requirements across four BIGE regional proxies. Their selection did not use disease annotations or AHBA results, and panel membership remained unchanged thereafter.

No gene met the stricter route-exclusive criterion. The fixed panel therefore represents genes with distributed expression across the BIGE representation of DCML, rather than genes unique to that pathway. Subsequent analyses tested whether the panel as a whole carried relational information across narrower, donor-resolved central regions.

#### The fixed panel showed greater similarity between adjacent positions in the anatomical sequence

In the AHBA mapping, the consecutive pairs were Cu/Gr with DTLv and DTLv with postcentral gyrus; Cu/Gr with postcentral gyrus was the first-to-third pair. Across the six donors, the fixed 19-gene panel produced a mean anatomical-order contrast of T=0.345175, with a median donor value of 0.431140. Both consecutive-pair correlations were positive in every donor. Five of six donor contrasts were positive, with values from -0.060526 to 0.685965. The exact one-sided sign test gave P=0.109375. Thus, the profile-level pattern was directionally consistent across both consecutive pairs, while the six donor signs alone supplied limited population-level precision.

The direction was also stable when each donor was omitted in turn. All six leave-one-donor-out estimates remained positive, with values from 0.277018 to 0.426316. No single donor determined the sign of the overall contrast.

**Figure 1.**
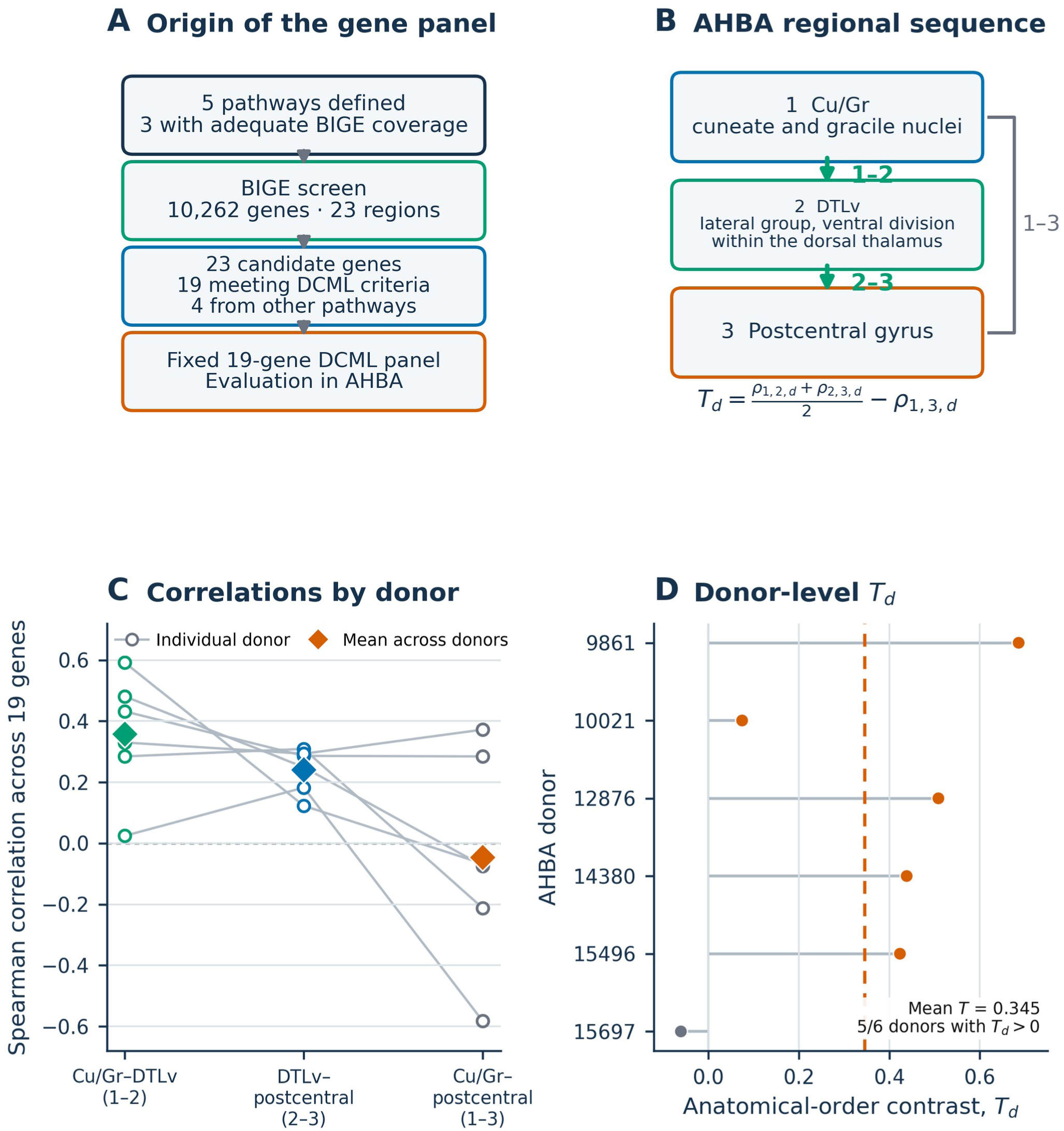
Origin of the panel and regional-order result. Panel A traces five predefined pathways, three with adequate Body Index of Gene Expression (BIGE) coverage, to 23 candidates: 19 genes meeting dorsal column–medial lemniscus (DCML) criteria and four genes meeting criteria for other pathways. The 19 DCML genes formed the fixed panel. Panel B defines the Allen Human Brain Atlas (AHBA) sequence: Cu/Gr, an aggregate of samples assigned to the cuneate and gracile nuclei; DTLv, the lateral group of nuclei, ventral division, within the dorsal thalamus; and postcentral gyrus. Panels C and D show the three Spearman correlations and donor-level contrast T_d. Both adjacent-in-order correlations were positive in all six donors; T_d was positive in five, and mean T was 0.345175. Adjacent-in-order denotes predefined anatomical position rather than physical proximity or demonstrated connectivity.

#### Both adjacent-in-order pairs contributed to the contrast

The mean consecutive-pair correlation across donors was 0.298684. The mean of the smaller consecutive-pair correlation within each donor was 0.209942, which shows that the pattern extended to the weaker of the two regional links. Both consecutive-pair correlations were positive in all six donors.

Each of these three components was unusual under both selection-matched reference distributions after Holm correction. Under the selection-class-matched model, adjusted P values were 0.048719 for the mean consecutive-pair correlation, 0.037679 for the weaker-pair statistic, and 0.048719 for the six-of-six count. The corresponding values under the model that additionally matched broad neuronal contrasts were 0.044999, 0.028399, and 0.044999.

Mean first-to-third similarity was -0.046491. Its lower-tail position was ordinary under both matched-panel models, with Holm-adjusted P values of 0.407052 and 0.521710. The aggregate contrast therefore arose from similarity in both consecutive pairs rather than from unusually low similarity between the first and third regions.

### The panel exceeded gene sets selected under comparable constraints

The selection-matched reference distributions had positive means, which indicates that the anatomy-guided selection process itself favored panels with some positive anatomical-order contrast. The target panel nevertheless extended beyond the upper tails of both distributions.

In the selection-class-matched model, the null mean was 0.138278 and the 95th percentile was 0.318275. Of 50,000 panels, 1,395 equaled or exceeded the observed value, which gave P=0.027919. In the model that additionally matched broad neuronal contrasts, the null mean was 0.175273 and the 95th percentile was 0.321637. A total of 1,328 panels equaled or exceeded the target, which gave P=0.026579. These results show that the target was unusual relative to panels generated under two approximations of the same selection process. Their inferential scope remains conditional on the modeled selection rules and on AHBA, the atlas in which the pattern was recognized.

**Figure 2.**
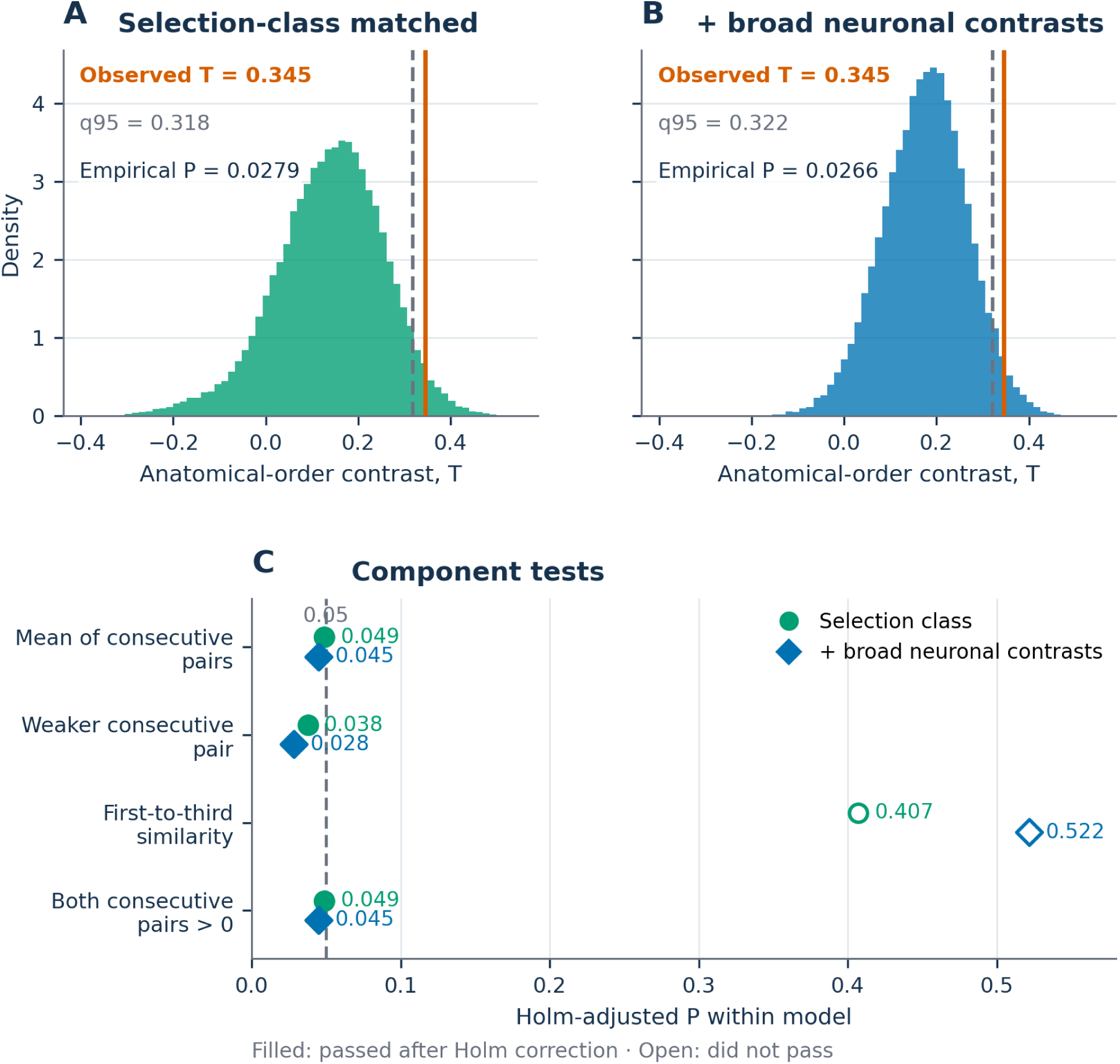
Selection-matched gene panels and component tests. T denotes the mean anatomical-order contrast across the six Allen Human Brain Atlas (AHBA) donors. Panels A and B show the observed T relative to 50,000 selection-class-matched panels and 50,000 panels additionally matched for broad neuronal contrasts. The observation exceeded both 95th percentiles, with empirical upper-tail probabilities of 0.027919 and 0.026579. Panel C decomposes T. The consecutive-pair mean, the weaker consecutive correlation, and the six-of-six donor count were unusual after Holm correction, while first-to-third similarity occupied an ordinary position.

#### The target regional mapping ranked highly in descriptive anatomical comparisons

The Cu/Gr-DTLv-postcentral combination ranked third among 240 comparable brainstem-thalamic-parietal combinations. This complete-set rank is descriptive because the comparison was designed after recognition of the target pattern.

In the post hoc local comparison, the target ranked first among the 17 combinations comprising the target and every single-region substitution. All 16 substitutions had lower contrasts. The highest alternative replaced Cu/Gr with the inferior olivary complex and retained DTLv and postcentral gyrus; its value was T=0.240058, 0.105117 below the target. An additive model based on the marginal means for the three regional identities predicted T=0.006871 for the target. The observed-minus-expected residual was 0.338304 and ranked fourth among all 240 combinations. DTLv ranked fifth of six thalamic regions by its marginal mean, and postcentral gyrus ranked fourth of five parietal regions. The target’s high value therefore reflected the particular three-region combination more than three individually high-ranking regions. This decomposition is descriptive and post hoc.

The two combinations above the target in the complete ranking had larger composite contrasts but a different correlation pattern. For the combination comprising the spinal trigeminal nucleus aggregate (Sp5), rostral intralaminar thalamus (ILr), and supramarginal gyrus, the two mean consecutive correlations were -0.011988 and 0.328655, and the first-to-third correlation was -0.232749. Both consecutive correlations were positive in four of six donors. For Sp5, the medial thalamic group (DTM), and supramarginal gyrus, the corresponding values were -0.018129, 0.314620, and -0.232749, with both consecutive correlations positive in two of six donors. In the target, the two mean consecutive correlations were 0.357018 and 0.240351, the first-to-third correlation was -0.046491, and both consecutive correlations were positive in all six donors. Thus, the higher Sp5 composite values arose mainly from one positive thalamic-cortical correlation combined with lower first-to-third similarity. Among the three highest-ranked combinations, the target alone showed positive correlations for both consecutive pairs in every donor.

Across all 240 combinations, only two showed positive correlations for both consecutive pairs in all six donors: the target and a combination comprising the arcuate nucleus of the medulla (Arc), the dorsal division of the lateral thalamic group (DTLd), and postcentral gyrus. The target ranked first of these two by the anatomical-order contrast. Arc-DTLd-postcentral had stronger mean consecutive correlations, 0.408772 and 0.309649, but also high first-to-third similarity of 0.295906; its contrast was consequently 0.063304, rank 100 of 240. Its consecutive-pair bottleneck was 0.309649, rank 1 of 240, compared with 0.240351 and rank 2 of 240 for the target. No combination exceeded the target contrast while retaining positive correlations at both consecutive pairs in every donor. This all-combination component screen was post hoc, and its counts and ranks are descriptive.

**Table 1.** Decomposition of the three regional combinations with the largest anatomical-order contrast. T denotes the mean anatomical-order contrast across donors. Cu/Gr, analysis-defined aggregate of samples assigned to the cuneate and gracile nuclei; DTLv, pooled samples labeled lateral group of nuclei, ventral division, within the dorsal thalamus; Sp5, spinal trigeminal nucleus aggregate; ILr, rostral intralaminar thalamus; DTM, medial thalamic group.

| Regional combination | Mean correlation 1-2 | Mean correlation 2-3 | Mean first-to-third correlation | Mean within-donor weaker relation | Mean T | Donors with both pairs positive |
| --- | --- | --- | --- | --- | --- | --- |
| Cu/Gr-DTLv-postcentral gyrus | 0.357018 | 0.240351 | -0.046491 | 0.209942 | 0.345175 | 6/6 |
| Sp5-ILr-supramarginal gyrus | -0.011988 | 0.328655 | -0.232749 | -0.014035 | 0.391082 | 4/6 |
| Sp5-DTM-supramarginal gyrus | -0.018129 | 0.314620 | -0.232749 | -0.040351 | 0.380994 | 2/6 |

**Figure 3.**
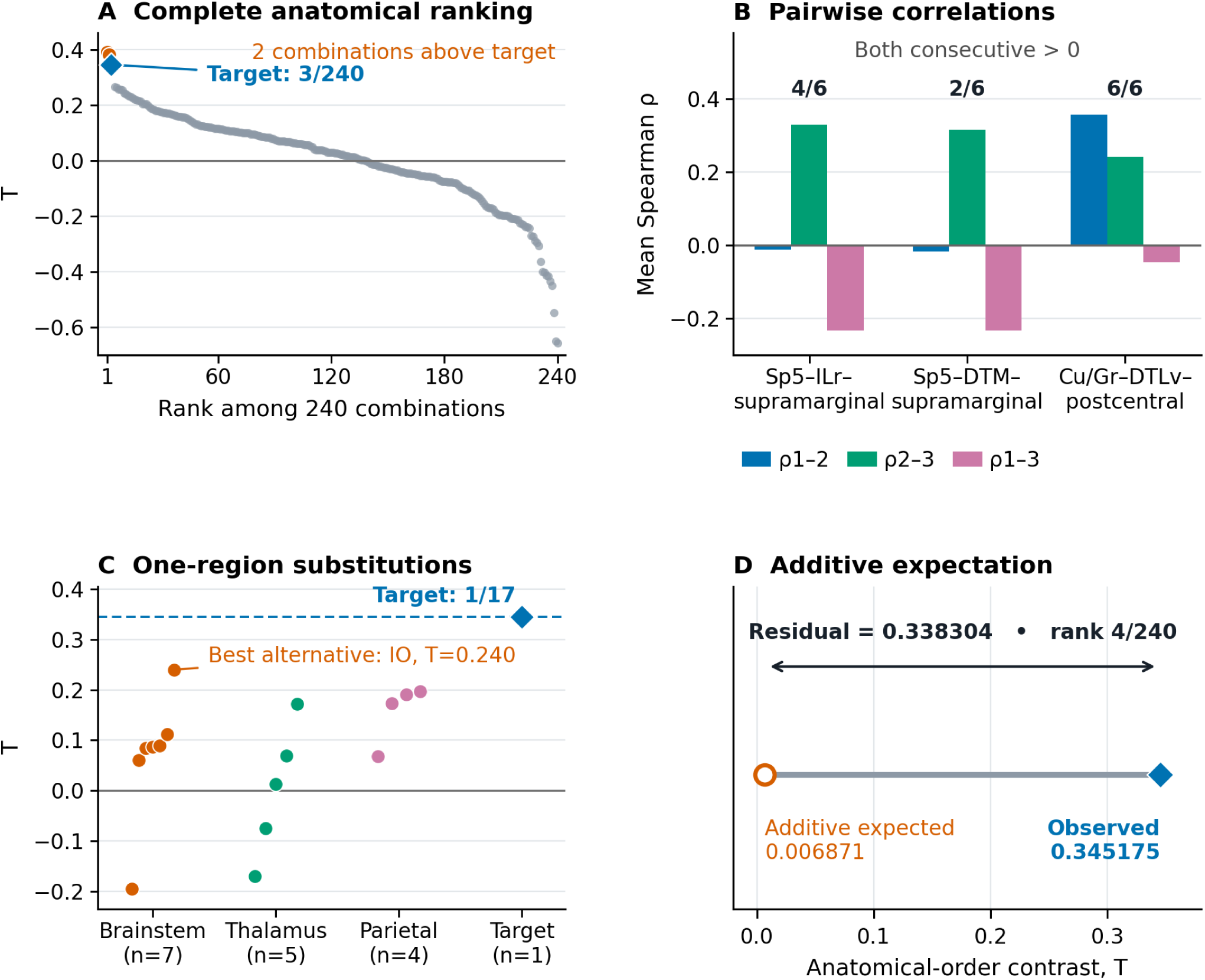
Anatomical comparison of the target combination. T denotes the mean anatomical-order contrast across donors. Panel A places Cu/Gr-DTLv-postcentral gyrus among all 240 comparable regional combinations. Panel B decomposes the two larger Sp5 composite contrasts and the target into their two consecutive correlations and first-to-third correlation. Panel C compares the target with all 16 single-region substitutions; every substitution had a lower contrast, and the strongest reached 0.240058. Panel D compares the observed target contrast with the additive expectation from the marginal means of its three regional identities; the observed-minus-expected residual was 0.338304 and ranked fourth among 240 combinations. The local, additive, and component analyses are post hoc and descriptive. Cu/Gr, aggregate of samples assigned to the cuneate and gracile nuclei; DTLv, pooled samples labeled lateral group of nuclei, ventral division, within the dorsal thalamus; Sp5, spinal trigeminal nucleus aggregate; ILr, rostral intralaminar thalamus; DTM, medial thalamic group. IO, inferior olivary complex.

### Whole-donor spatial calibration favored the target while local calibration was at the boundary

The whole-donor co-primary model placed the observed contrast well beyond its surrogate distribution. Its null mean was 0.006872, its 95th percentile was 0.221499, and 192 of 50,000 surrogate values reached the observation, which gave P=0.003860. Both whole-donor sensitivity models yielded similar results: the dense-distance model gave P=0.005099, and the axis-standardized eight-nearest-neighbor model gave P=0.004500. The observed contrast exceeded the 95th percentile in all three whole-donor formulations.

The within-compartment models produced reference distributions centered closer to the observation. In the primary five-nearest-neighbor model, the null mean was 0.169386 and the 95th percentile was 0.345906. The observed value of 0.345175 fell 0.000731 below that percentile; 2,558 of 50,000 surrogates reached it, which gave P=0.051179. Two within-compartment sensitivity models placed the observation just beyond their nominal thresholds, with P=0.043196 for dense-distance weights and P=0.043996 after coordinate-axis standardization.

The spatial rule, frozen after recognition of the pattern and before surrogate generation, required both co-primary models to pass. Since the primary within-compartment model did not pass, the joint criterion was not met. Across models, whole-donor calibration favored the observation, whereas local structure within medulla, thalamus, and parietal cortex placed it near the decision boundary. The spatial result therefore depended on analytical scale.

Leave-one-donor-out calibration reinforced this distinction. All six whole-donor omissions exceeded their corresponding null 95th percentiles, compared with three of six omissions under the primary within-compartment model. These overlapping subsets are sensitivity tests rather than separate replications.

**Figure 4.**
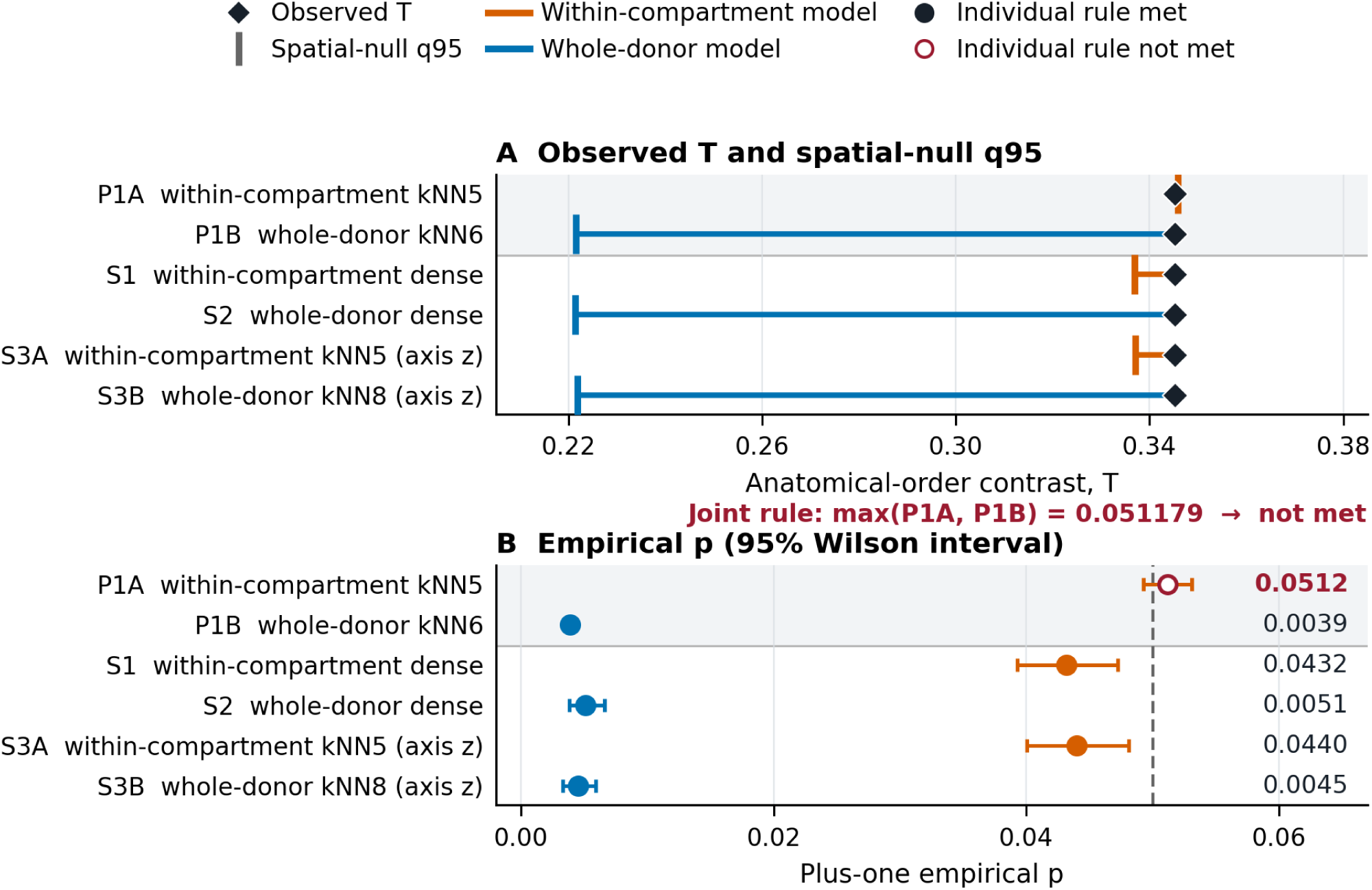
Spatial calibration at two anatomical scales in the Allen Human Brain Atlas (AHBA). T denotes the mean anatomical-order contrast across donors. Panel A compares the observed T with the null 95th percentile for two co-primary models and four frozen post-discovery sensitivities. Panel B shows the corresponding empirical upper-tail probabilities; horizontal whiskers are 95% Wilson intervals for the null exceedance fraction. Panel A uses an expanded horizontal scale spanning the null thresholds and the observed value. Whole-donor calibration favored the target. Within-compartment models placed it near the 95th percentile, and the primary within-compartment model did not pass. The frozen two-model criterion therefore was not met. P1A, co-primary within-compartment model; P1B, co-primary whole-donor model; S1, S2, S3A, and S3B, prespecified sensitivity models; kNN, k-nearest-neighbor graph; q95, 95th percentile of the null distribution; (z), standardized coordinate axes.

### Cell-class and probe analyses supported a panel-level interpretation

Neuronal-minus-non-neuronal expression was positive for the panel in each broad region examined: 0.640672 in medulla oblongata, 0.564917 in thalamus, and 0.604205 in cerebral cortex. Fourteen of the 19 genes had positive neuronal contrasts in all three macroregions. The smallest regional panel contrast, 0.564917, exceeded the historical technical comparator, with 30 of 50,000 sets at or above the observation (P=0.000620). Under the stricter selection-matched comparison, the same statistic yielded P=0.096778, with a 95th percentile of 0.610726. Thus, the panel showed consistent neuronal direction across all three macroregions, while its enrichment relative to similarly selected panels remained uncertain. The broad regional summaries do not resolve cell types at the exact target nuclei.

The post hoc HPA anatomical-order contrast was positive (T=0.780702). Its values under the two matched-panel comparisons were P=0.216636 and P=0.185236. This result provides descriptive cross-platform concordance in direction but does not establish regional specificity.

All 50,000 random choices of one AHBA probe per gene produced a positive anatomical-order contrast. The median was 0.298099, and the 2.5th to 97.5th percentile range was 0.162573 to 0.428363. The direction therefore remained stable across probe choices. Magnitude and threshold-level exceptionality varied: 49.038% of the one-probe values exceeded the historical technical-null 95th percentile.

The two-way gene-by-donor bootstrap gave a median of 0.338036, and 96.114% of draws were positive. Its diagnostic 2.5th to 97.5th percentile range was -0.043287 to 0.631346. These analyses support the 19-gene panel as the appropriate unit of interpretation; the study does not assign the aggregate result to any single gene.

**Supplementary Figure S1.**
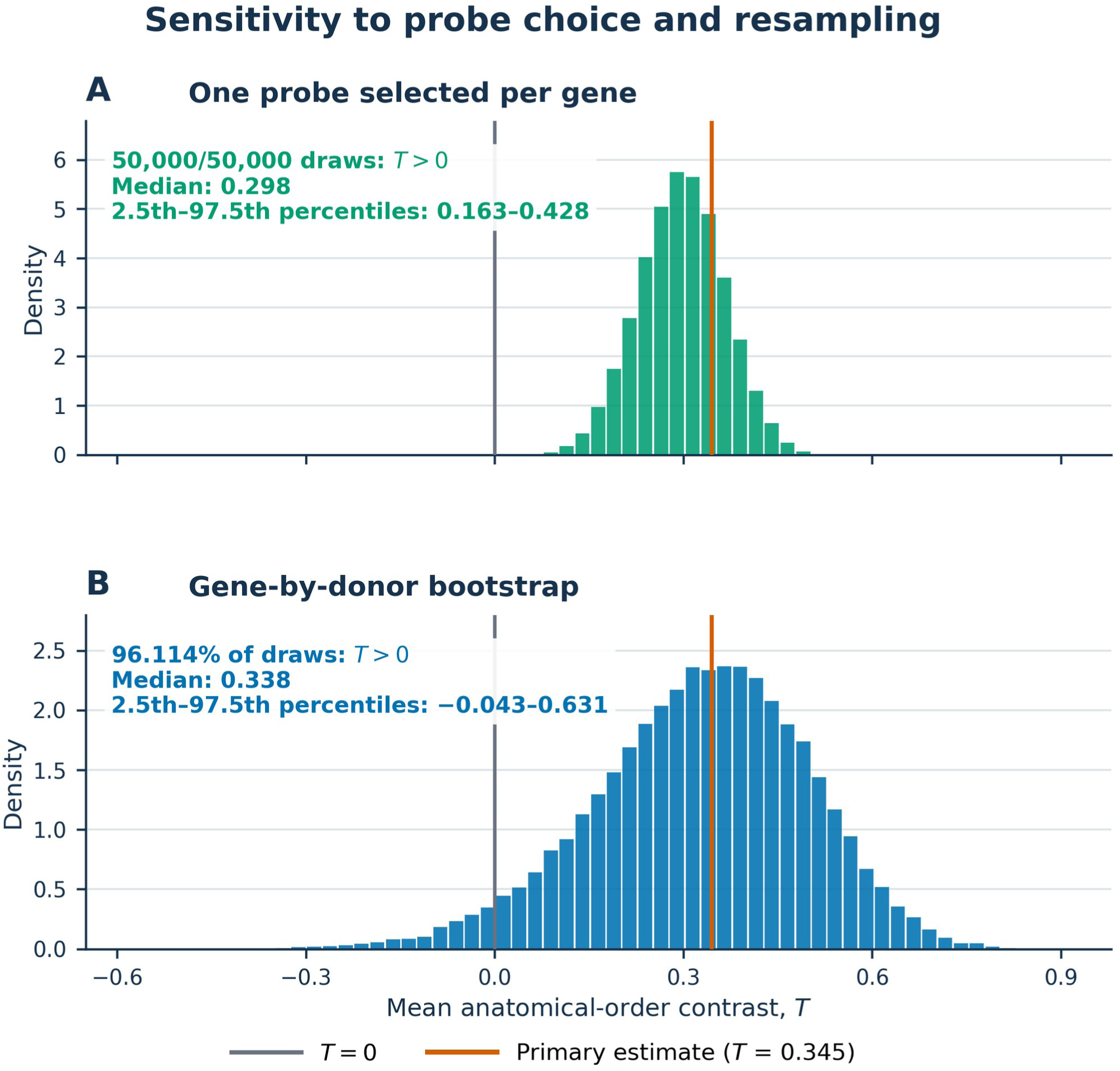
Sensitivity of the anatomical-order contrast to probe choice and resampling. T denotes the mean anatomical-order contrast across donors. Panel A shows the distribution of T in 50,000 random selections of one Allen Human Brain Atlas (AHBA) probe for each of the 19 genes; every value was positive (median 0.298; 2.5th-97.5th percentiles, 0.163-0.428). Panel B shows the distribution of T in 50,000 two-way bootstrap samples of genes and donors; 96.1% were positive (median 0.338; 2.5th-97.5th percentiles, -0.043 to 0.631). The gray line marks T=0 and the orange line the primary estimate obtained with median probe collapse (T=0.345). The bootstrap percentile range is a diagnostic summary, not a confirmatory confidence interval.

### The historical source cohort showed a positive relative-order contrast and donor heterogeneity

The GSE3526 reconstruction yielded positive mean anatomical-order contrasts under every probe-collapse method. For the three-region medulla-thalamus-parietal sequence, mean contrasts were 0.213706 for median collapse, 0.221053 for mean collapse, and 0.191338 for the historical representative probe. Five, six, and five of eight donors, respectively, had positive values. For the four-region DRG-medulla-thalamus-parietal sequence, the corresponding means were 0.152193, 0.165205, and 0.109722, with six of eight donors positive under each method.

The three-region median result reflected relative order rather than uniformly positive coexpression. Mean medulla-thalamus and thalamus-parietal correlations were -0.022149 and -0.028509, while the medulla-parietal first-to-third correlation was -0.239035. The consecutive regions were less dissimilar than the first and third regions. The magnitude also depended on within-donor standardization across the frozen neural-tissue background, as detailed in the supplementary analysis.

The four donors recorded as female had higher three-region contrasts than the four recorded as male under all three probe-collapse methods. Mean female and male values were 0.563158 and -0.135746 for median collapse, 0.568640 and -0.126535 for mean collapse, and 0.529167 and -0.146491 for the representative probe. Among the 70 possible partitions of eight donors into two groups of four, two produce complete separation in either direction (2/70=0.0286). This exploratory pattern was based on four donors per group, key postmortem and technical covariates were unavailable, and the separation did not recur for the four-region median contrast. It identifies donor heterogeneity suitable for prospective study rather than an estimated sex effect.

### Evidence summary

**Table 2.** Summary of the principal findings and their evidential scope. T denotes the mean anatomical-order contrast across six donors; AHBA, Allen Human Brain Atlas; Arc, arcuate nucleus of the medulla; DTLd, dorsal division of the lateral thalamic group.

| Scientific question | Main result | Evidential scope |
| --- | --- | --- |
| Do regional profiles follow the known anatomical order? | Mean T=0.345175; both consecutive correlations positive in 6/6 donors; T>0 in 5/6 | Descriptive AHBA finding; donor sign test P=0.109375 |
| Do both consecutive pairs contribute? | Consecutive mean, weaker-pair statistic, and 6/6 count were unusual after Holm correction; first-to-third similarity was ordinary | Diagnostic decomposition under matched-panel models |
| Is the panel unusual under comparable selection constraints? | P=0.027919 and P=0.026579 in two 50,000-panel comparisons | Conditional post-discovery robustness within AHBA |
| Does the target stand out anatomically? | Rank 3/240; rank 1/17 among itself and all single-region substitutions; additive residual rank 4/240 | Descriptive evidence associated with the complete regional combination |
| How distinctive is positivity at both consecutive pairs across donors? | Only 2/240 regional combinations had both correlations positive in 6/6 donors; the target had the larger contrast of the two, and no higher-contrast combination met this condition | Post hoc descriptive component screen, not an inferential specificity test |
| Does spatial structure account for the result? | Whole-donor co-primary P=0.003860; within-compartment co-primary P=0.051179 | Whole-donor calibration favored the target; local evidence was near the boundary; the joint rule was not met |
| Is the direction technically stable? | 50,000/50,000 one-probe contrasts positive; 96.114% of gene-by-donor bootstrap draws positive | Directional robustness; magnitude remains probe and sampling sensitive |
| Does broad cell-class evidence support neuronal relevance? | Positive neuronal contrasts in medulla, thalamus, and cortex; selection-matched P=0.096778 | Broad neuronal plausibility without target-cell resolution |

## Discussion

The fixed 19-gene panel showed an ordered regional relationship across the represented central DCML sequence. In every AHBA donor, both Cu/Gr–DTLv and DTLv–postcentral correlations were positive, and the mean regional-order contrast was 0.345175. The panel exceeded two selection-matched gene-set distributions and remained prominent in regional-substitution, probe-selection, and donor-resampling analyses. Spatial surrogates showed that the strength of this evidence depended on the anatomical scale used to model background structure.

These findings extend a network view of the brain from macroscopic connectivity to regional molecular organization. Specialized regions retain local identities, and their transcriptomic profiles can also relate according to their positions within a distributed system. The signal observed here arises from the collective relationship among the 19-gene regional profiles. It identifies a molecular pattern aligned with the known DCML order while leaving axonal connectivity between sampled cells as a separate anatomical property.

The component results sharpen the biological meaning of the contrast. First-to-third similarity occupied an ordinary position under both matched-panel distributions. By comparison, the average adjacent-in-order similarity, the weaker of the two adjacent relationships, and the presence of two positive correlations in all six donors each occupied the upper tail of both conditional reference models after multiplicity correction. The regional-order contrast therefore reflected coordinated similarity at both steps of the represented sequence.

The matched-panel analysis also clarifies the role of anatomy-guided selection. Both reference distributions had positive means, which shows that selection rules similar to those used for the target panel can generate some alignment with anatomical order. The target still exceeded the 95th percentile of both distributions. These comparisons ask whether the panel carries more regional order than expected from key features of its own selection process. Their scope remains conditional because the same AHBA dataset contributed to pattern recognition and evaluation, and finite models reproduce only defined parts of the adaptive discovery path.

The anatomical analyses associated the result with the complete regional mapping. Cu/Gr–DTLv–postcentral gyrus ranked first among the target and all 16 single-region substitutions. Its observed contrast also exceeded the expectation from the marginal identities of its three regions by 0.338304, the fourth-largest residual among 240 combinations. DTLv and postcentral gyrus had modest marginal ranks across the full grid. The particular three-region configuration, rather than three independently high-ranking regions, therefore defines the prospective anatomical hypothesis.

Decomposition of the two higher-ranked Sp5 combinations provided an additional interpretive check. Their large composite values combined a positive thalamic–cortical correlation with a relatively low Sp5–cortical correlation, while the Sp5–thalamic correlation remained near zero on average. The target showed positive mean correlations at both adjacent-in-order pairs, and both were positive in every donor. Across all 240 combinations, only the target and Arc– DTLd–postcentral shared this six-donor consistency. Arc–DTLd–postcentral had a stronger adjacent-pair bottleneck but also high first-to-third similarity, which reduced its contrast to 0.063304. A prospective anatomical comparison can now freeze a two-link component rule and include sensory and non-sensory sequences.

Spatial calibration located the result at different levels of anatomical organization. All three whole-donor models placed the observed contrast above their 95th percentiles. Within each major anatomical compartment, the surrogate distributions shifted toward the observation: two sensitivity models crossed their nominal thresholds by small margins, while the primary local model yielded P = 0.051179 and placed the observation just below its 95th percentile. The frozen two-model conjunction was therefore not met. The regional relationship was distinguishable from whole-donor background organization and remained close to local spatial structure within medulla, thalamus, and parietal cortex.

Gene-panel and spatial analyses answer complementary questions. Matched panels vary gene membership while retaining the observed AHBA regional data. Spatial surrogates vary the anatomical maps of the fixed panel while preserving specified trends, spatial power, and cross-gene covariance. Their probabilities describe different reference processes and are interpreted separately. Together, they show favorable selection-aware calibration, favorable whole-donor spatial calibration, and a boundary result under the strict primary local spatial model.

Cell-class and technical analyses support a collective, panel-level interpretation. Neuronal-minus-non-neuronal contrasts were positive in medulla, thalamus, and cortex. Every one of the 50,000 random one-probe-per-gene selections retained a positive contrast, although magnitude varied across probe choices. The gene-by-donor bootstrap favored a positive direction in 96.114% of draws and produced a percentile range that included zero. The defensible unit of inference is the complete 19-gene panel, and sequence-based validation with RNA sequencing or spatial transcriptomics can remove microarray probe ambiguity.

GSE3526 adds a donor-resolved view within the historical source cohort. Its average relative-order contrast was positive across three probe-collapse methods and in both three-and four-region summaries. The component correlations showed that consecutive broad tissues were less dissimilar than the first and last tissues, and the magnitude varied with the neural background used for within-donor standardization. Its shared lineage with BIGE and its broad tissue labels define this reconstruction as a source-cohort sensitivity analysis.

The separation by recorded sex in the three-region GSE3526 contrast is a hypothesis-generating observation. Removal of X-linked *RAI2* preserved complete separation for the median and mean summaries, and every leave-one-gene-out female-minus-male mean difference remained positive. The representative-probe summary after *RAI2* removal and the four-region median contrast did not reproduce complete separation. Four donors per group and the absence of key postmortem and batch covariates limit biological attribution. A future donor-level design can evaluate sex-associated heterogeneity prospectively with balanced sampling and adjustment for age, postmortem interval, RNA integrity, cause of death, batch, and cellular composition.

The study combines several strengths: fixed panel membership during final robustness analyses, donor-resolved evaluation, reference panels that approximate the anatomy-guided selection process, separate assessment of both adjacent relationships, spatial calibration at two scales, and an integrity-checked reproducibility archive. Its anatomical resolution is defined precisely by the source data. Cu/Gr is an analytical aggregate, DTLv is a broad sampled category rather than an isolated ventral posterolateral nucleus, and postcentral gyrus is a gyral-level cortical representation. AHBA contributes six donors with unequal regional coverage, probe choice affects magnitude, and broad cell-class summaries do not deconvolve each target sample. These features delimit the inference and specify the dataset needed for the next test.

The next study should freeze the 19 genes, anatomical mappings, contrast, and spatial models before inspecting a new dataset. Donor-level RNA sequencing, single-nucleus profiling with regional resolution, or spatial transcriptomics could test the pattern while improving cell-type attribution and probe specificity. Sufficient neighboring tissue would support local spatial nulls, and comparisons across sensory and non-sensory sequences would test the breadth of the principle. This design would determine whether the observed regional relationship generalizes beyond the current atlas.

## Conclusion

The anatomy-constrained screen identified a fixed 19-gene panel whose collective expression profile followed the represented central sequence of the DCML system. Across six AHBA donors, both adjacent-in-order regional correlations were consistently positive, the panel outperformed gene sets generated under comparable selection constraints, and its regional combination remained highly ranked across anatomical and technical sensitivity analyses. Spatial calibration showed that the strength of this relationship depends on the anatomical scale used to model background structure. Together, these findings connect pathway anatomy with regional transcriptomic organization and define a specific, reproducible hypothesis for prospective testing in an independent, anatomically resolved dataset.

## Supporting information

Supplementary Archive S1

## Data and code availability

The frozen analysis code, protocols, amendments, software-environment records, aggregate outputs, and checksum manifests are provided with this preprint as Supplementary Archive S1. The public archive passed its packaged file-level integrity checks. AHBA data are available from the Allen Institute [6,16,17], and GSE3526 data and metadata are available from NCBI Gene Expression Omnibus [12].

Aggregate statistics, figures, candidate lists, anatomical mappings, and analytical descriptions derived from the processed BIGE extract are included in the manuscript and supplementary files. The complete processed BIGE matrix and exports that would substantially reconstruct it are not redistributed. This restriction does not affect access to GSE3526 or the other public source resources.

## Author contributions

Gerardo A. Boni: Conceptualization, Methodology, Software, Formal analysis, Investigation, Data curation, Validation, Visualization, Writing – original draft, Writing – review and editing, and Project administration. Alejandro Luque Hernández: Conceptualization, Methodology, Supervision, Validation, and Writing – review and editing. Vivien Iran Bautista Ruedas: Investigation, Validation, and Writing – review and editing. All authors interpreted the findings, reviewed and approved the submitted version, and accept responsibility for the integrity of the work.

## Ethics statement

This study was limited to secondary analysis of previously generated aggregate or deidentified human tissue transcriptomic data. No participants were prospectively recruited, no interventions or new biospecimen collections were performed, and the authors had no access to direct personal identifiers. Ethical oversight and consent procedures for the original tissue collections are described in the source studies and repositories.

## Declaration of generative AI and AI-assisted technologies

During manuscript preparation, G.A.B. used OpenAI Codex in ChatGPT Work, with the model identified in the user interface as GPT-5.6 Sol (accessed in 2026), to improve the grammar, readability, and stylistic consistency of the English-language text. The scientific content and interpretations were supplied and approved by the authors. G.A.B. reviewed and edited all retained suggestions; all authors reviewed and approved the final manuscript and take responsibility for its accuracy, originality, and integrity.

## Supplementary analysis

**Supplementary Table S1.**
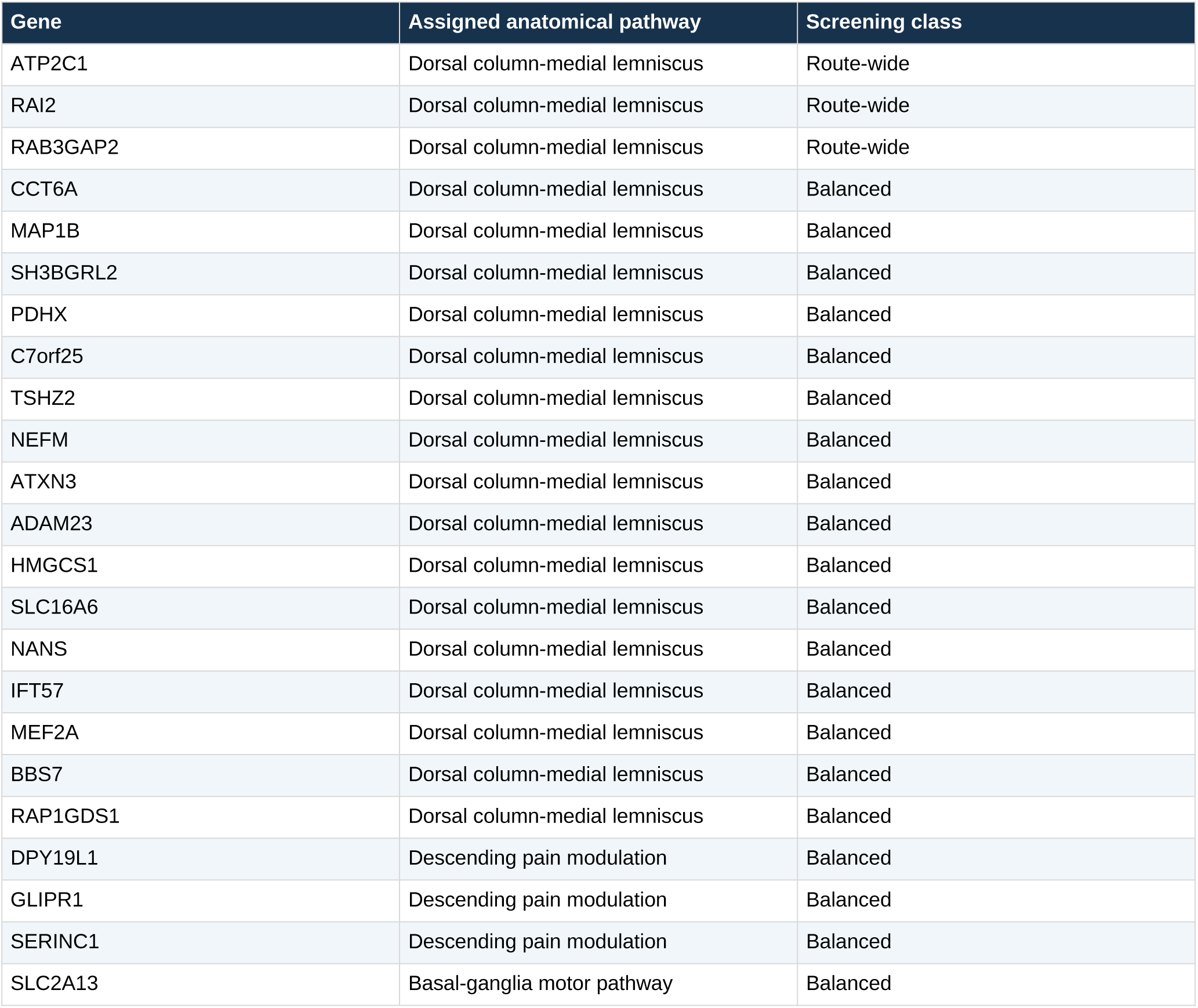
Candidate genes from the anatomy-constrained Body Index of Gene Expression (BIGE) screen. The route-wide class required the weakest represented region to exceed the 90th percentile of off-route regions. The balanced class required a high route minimum and route mean while limiting single-region dominance. Candidate membership was fixed before evaluation in the Allen Human Brain Atlas (AHBA). DCML, dorsal column-medial lemniscus.

| Gene | Assigned anatomical pathway | Screening class |
| --- | --- | --- |
| ATP2C1 | Dorsal column-medial lemniscus | Route-wide |
| RAI2 | Dorsal column-medial lemniscus | Route-wide |
| RAB3GAP2 | Dorsal column-medial lemniscus | Route-wide |
| CCT6A | Dorsal column-medial lemniscus | Balanced |
| MAP1B | Dorsal column-medial lemniscus | Balanced |
| SH3BGRL2 | Dorsal column-medial lemniscus | Balanced |
| PDHX | Dorsal column-medial lemniscus | Balanced |
| C7orf25 | Dorsal column-medial lemniscus | Balanced |
| TSHZ2 | Dorsal column-medial lemniscus | Balanced |
| NEFM | Dorsal column-medial lemniscus | Balanced |
| ATXN3 | Dorsal column-medial lemniscus | Balanced |
| ADAM23 | Dorsal column-medial lemniscus | Balanced |
| HMGCS1 | Dorsal column-medial lemniscus | Balanced |
| SLC16A6 | Dorsal column-medial lemniscus | Balanced |
| NANS | Dorsal column-medial lemniscus | Balanced |
| IFT57 | Dorsal column-medial lemniscus | Balanced |
| MEF2A | Dorsal column-medial lemniscus | Balanced |
| BBS7 | Dorsal column-medial lemniscus | Balanced |
| RAP1GDS1 | Dorsal column-medial lemniscus | Balanced |
| DPY19L1 | Descending pain modulation | Balanced |
| GLIPR1 | Descending pain modulation | Balanced |
| SERINC1 | Descending pain modulation | Balanced |
| SLC2A13 | Basal-ganglia motor pathway | Balanced |

**Supplementary Table S2.** Candidate-circuit catalog and screening decisions. Sixteen literature-defined circuits were cataloged before the gene-expression screen. Five were predefined for possible BIGE screening; only three met the requirement of at least seven source samples for every regional proxy.

| Circuit | Domain | Literature-defined sequence | Screening decision |
| --- | --- | --- | --- |
| Interoceptive ladder | Pain/interoception | Medulla → thalamus → posterior/mid/anterior insula | Conditional; inadequate insula resolution |
| Dorsal-column pathway | Discriminative touch | DRG → dorsal-column nuclei → VPL → S1 | Predefined; entered BIGE screen (DCML) |
| Affective-parabrachial pain | Pain/interoception | Dorsal horn → parabrachial → amygdala/hypothalamus/aMC C | Conditional; incomplete human-resolved chain |
| Descending modulation | Pain/interoception | Forebrain → PAG → RVM → dorsal horn | Predefined; entered BIGE screen |
| Basal-ganglia motor loop | Motor | Cortex → putamen/STN → GPi/SNr → thalamus → cortex | Predefined; entered BIGE screen |
| Cerebro-cerebellar loop | Motor/cognitive | Cortex → pons → cerebellum/dentate → thalamus → cortex | Predefined; excluded because pons n=1 |
| Corticospinal execution | Motor | M1 → internal capsule → brainstem → spinal cord | Negative comparator; internal capsule unavailable |
| Dorsomedial reach | Visuomotor | Medial parietal → dorsal premotor → pons/cerebellum | Deferred; cortical parcels unresolved |
| Dorsolateral grasp | Visuomotor | Anterior parietal → ventral premotor → M1 | Deferred; parcels unresolved |
| Medial initiation | Motor | pre-SMA/SMA → striatum → thalamus → SMA | Secondary; SMA/pre-SMA unresolved |
| Salience handoff | Salience/control | Anterior insula/aMCC → executive/autonomic networks | Deferred; key parcels absent |
| Papez circuit | Memory | Hippocampus → mammillary → anterior thalamus → cingulate → parahippocampus | Secondary; several proxies incomplete |
| Dorsal language stream | Language | Posterior temporal → inferior parietal → inferior frontal | Deferred; lobar resolution insufficient |
| Two visual streams | Visual | Occipital → temporal/parietal → frontal | Secondary; coarse lobar contrast only |
| Reward circuit | Reward | mPFC/VTA → accumbens → ventral pallidum → MD → mPFC | Predefined; excluded because pallidal proxy n=1 |
| Auditory 'what' stream | Auditory | Auditory thalamus → temporal cortex → frontal cortex | Deferred; MGN/auditory areas unavailable |

*Note. BIGE, Body Index of Gene Expression; DCML, dorsal column-medial lemniscus; DRG, dorsal root ganglion; VPL, ventral posterolateral thalamic nucleus; S1, primary somatosensory cortex; aMCC, anterior midcingulate cortex; PAG, periaqueductal gray; RVM, rostral ventromedial medulla; STN, subthalamic nucleus; GPi, internal globus pallidus; SNr, substantia nigra pars reticulata; M1, primary motor cortex; pre-SMA, presupplementary motor area; SMA, supplementary motor area; mPFC, medial prefrontal cortex; VTA, ventral tegmental area; MD, mediodorsal thalamic nucleus; MGN, medial geniculate nucleus. Predefined identifies the five routes frozen before the gene-expression screen; only three entered the BIGE screen.*

### GSE3526 tissue background and relative similarity

The GSE3526 series contains 193 arrays from the frozen neural-tissue set across ten donors. Of these, 180 arrays belonged to the eight donors with complete regional measurements and could contribute to within-donor standardization. The three-region comparison used 24 arrays, and the four-region comparison used 32.

The magnitude of the result depended on the background used to standardize each gene. With the 20 neural tissues common to all eight donors, the median-collapse contrasts were approximately 0.243 for the three-region sequence and 0.150 for the four-region sequence. Values fell to approximately 0.099 and 0.045 when all available tissues per donor from the 65-tissue universe were used, and to 0.003 and 0.011 without within-donor neural standardization. The result therefore represents relative differentiation against the specified tissue background.

Permutation and sign-flip summaries were descriptive because the panel, cohort, and analytical outcome had already been selected, and the anatomical labels do not satisfy a simple exchangeability assumption. These post-selection summaries were not used for confirmatory inference.

### Exploratory donor heterogeneity in GSE3526

Removal of X-linked *RAI2* did not account for the difference between donors recorded as female and male in the three-region analysis. Female-minus-male mean differences remained 0.598, 0.590, and 0.560 under median, mean, and representative-probe collapse. Across leave-one-gene-out analyses, every difference remained positive, with values from 0.549 to 0.768. Joint removal of *RAI2* and *C7orf25* retained differences from 0.422 to 0.490. These checks show that the descriptive separation did not depend on a single gene. They do not resolve confounding among sex, age, postmortem characteristics, RNA quality, and batch.

**Supplementary Figure S2.**
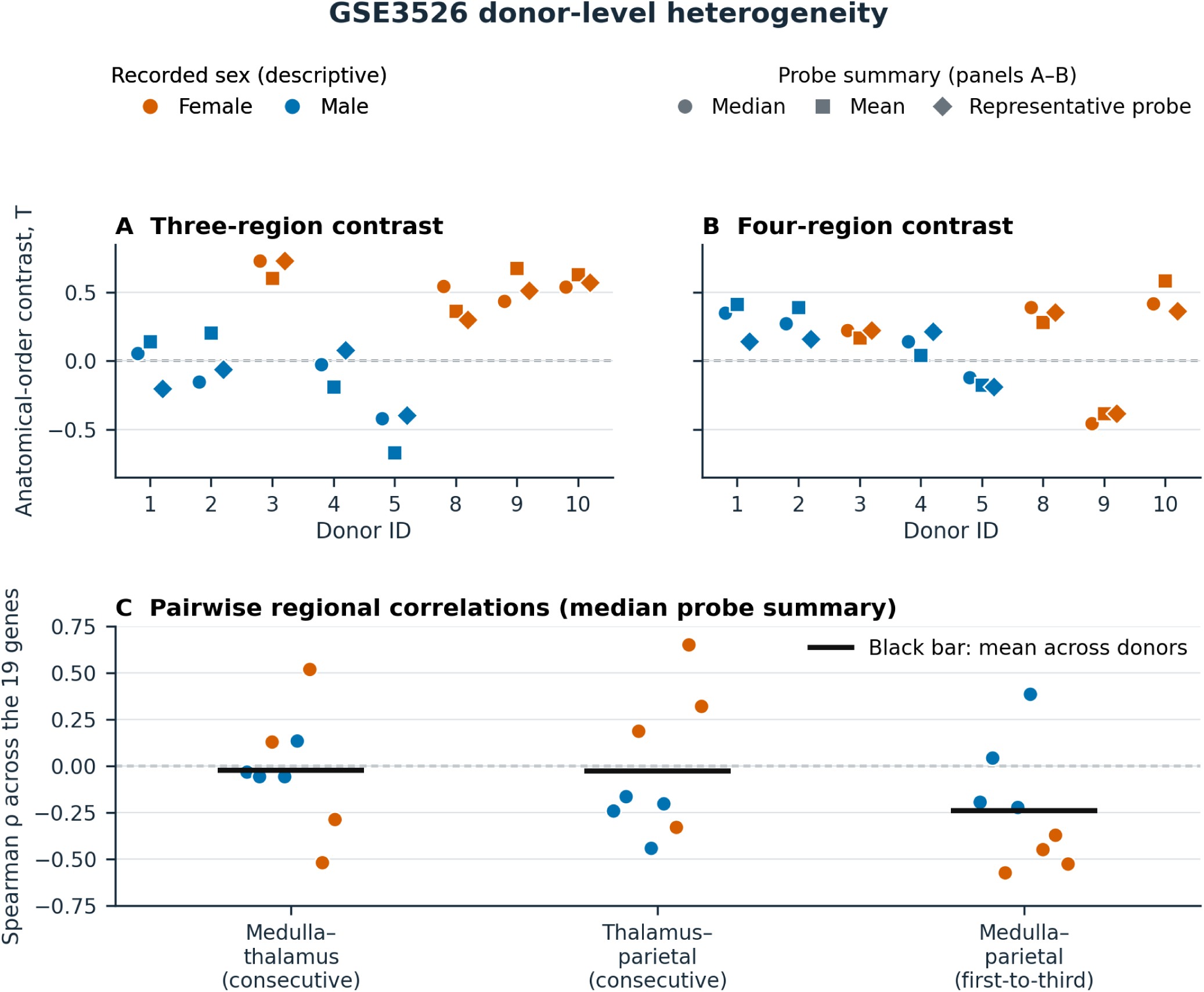
Donor heterogeneity and anatomical-order contrast in Gene Expression Omnibus (GEO) series GSE3526. Panels A and B show three-region and four-region contrasts for eight complete donors by recorded sex and probe-collapse method. Panel C shows the donor-specific pairwise correlations underlying the median-collapse three-region contrast; the x-axis identifies the two consecutive pairs and the first-to-third pair, and black horizontal bars mark the mean across donors. Complete separation by recorded sex occurred in the exploratory three-region comparison; it did not recur for the four-region median-collapse contrast. Each recorded-sex group contained four donors, and relevant postmortem and technical covariates were unavailable.

### GSE3526 probe-annotation sensitivity

The historical Affymetrix Human Genome U133 Plus 2.0 platform annotation (Gene Expression Omnibus accession GPL570) assigns three *RAB3GAP2* probe sets that also map to AURKAPS1 and one *C7orf25* probe set that also maps to PSMA2 [18]. A labeled post hoc sensitivity excluded both genes. Direction remained positive: mean contrasts ranged from 0.187653 to 0.250766 for the three-region analysis and from 0.114890 to 0.203942 for the four-region analysis. Five or six of eight donors were positive in the three-region configurations, and six of eight were positive in every four-region configuration. Contemporary reprocessing of the raw Affymetrix CEL intensity files, the platform’s probe-level data format, with an updated probe definition remains a useful prospective analysis.

### Leave-one-donor-out spatial diagnostics

All six observed AHBA leave-one-donor-out contrasts were positive. Under the technical and neuron-matched gene-panel references, four and three of the six omissions, respectively, exceeded the relevant 95th percentiles. Under the primary spatial models, three of six within-compartment omissions and all six whole-donor omissions exceeded their corresponding 95th percentiles. The five-donor sets share four donors with one another and therefore describe sensitivity rather than independent replication.

**Supplementary Figure S3.**
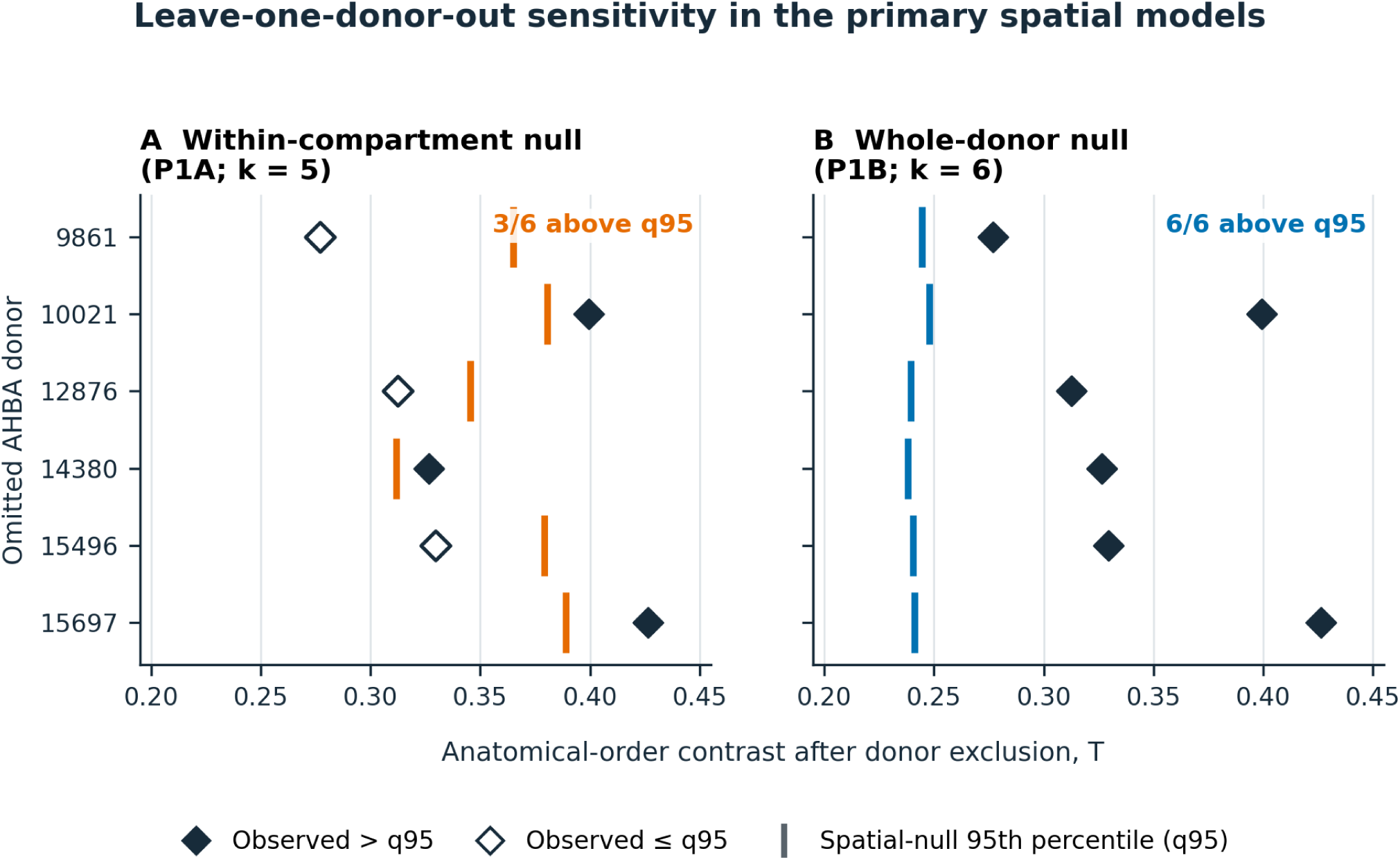
Leave-one-donor-out spatial sensitivity in the Allen Human Brain Atlas (AHBA). T denotes the mean anatomical-order contrast across retained donors. Every observed five-donor T was positive. Three of six within-compartment omissions and all six whole-donor omissions exceeded the corresponding null 95th percentile. Diamonds show observed values; filled diamonds exceeded q95, open diamonds did not, and vertical marks show the corresponding spatial-null 95th percentiles. The overlapping subsets quantify sensitivity to donor omission and are not independent replications.

## References

1. Bassett DS, Sporns O. Network neuroscience. Nature Neuroscience. 2017;20:353–364. doi:10.1038/nn.4502.

2. Richiardi J, Altmann A, Milazzo AC, et al. Correlated gene expression supports synchronous activity in brain networks. Science. 2015;348:1241–1244. doi:10.1126/science.1255905.

3. Burt JB, Demirtas M, Eckner WJ, et al. Hierarchy of transcriptomic specialization across human cortex captured by structural neuroimaging topography. Nature Neuroscience. 2018;21:1251–1259. doi:10.1038/s41593-018-0195-0.

4. Delhaye BP, Long KH, Bensmaia SJ. Neural basis of touch and proprioception in primate cortex. Comprehensive Physiology. 2018;8:1575–1602. doi:10.1002/cphy.c170033.

5. Loutit AJ, Vickery RM, Potas JR. Functional organization and connectivity of the dorsal column nuclei complex reveals a sensorimotor integration and distribution hub. Journal of Comparative Neurology. 2021;529:187–220. doi:10.1002/cne.24942.

6. Hawrylycz MJ, Lein ES, Guillozet-Bongaarts AL, et al. An anatomically comprehensive atlas of the adult human brain transcriptome. Nature. 2012;489:391–399. doi:10.1038/nature11405.

7. Wagner HH, Dray S. Generating spatially constrained null models for irregularly spaced data using Moran spectral randomization methods. Methods in Ecology and Evolution. 2015;6:1169–1178. doi:10.1111/2041-210X.12407.

8. Burt JB, Helmer M, Shinn M, Anticevic A, Murray JD. Generative modeling of brain maps with spatial autocorrelation. NeuroImage. 2020;220:117038. doi:10.1016/j.neuroimage.2020.117038.

9. Markello RD, Misic B. Comparing spatial null models for brain maps. NeuroImage. 2021;236:118052. doi:10.1016/j.neuroimage.2021.118052.

10. Markello RD, Arnatkeviciute A, Poline JB, Fulcher BD, Fornito A, Misic B. Standardizing workflows in imaging transcriptomics with the abagen toolbox. eLife. 2021;10:e72129. doi:10.7554/eLife.72129.

11. Lee J, Hever A, Willhite D, Zlotnik A, Hevezi P. Effects of RNA degradation on gene expression analysis of human postmortem tissues. FASEB Journal. 2005;19:1356–1358. doi:10.1096/fj.04-3552fje.

12. Roth RB, Hevezi P, Lee J, et al. Gene expression analyses reveal molecular relationships among 20 regions of the human CNS. Neurogenetics. 2006;7:67–80. doi:10.1007/s10048-006-0032-6. GEO GSE3526.

13. Ushach I, Burkhardt AM, Martinez C, et al. METEORIN-LIKE is a cytokine associated with barrier tissues and alternatively activated macrophages. Clinical Immunology. 2015;156:119–127. doi:10.1016/j.clim.2014.11.006.

14. Siletti K, Hodge R, Mossi Albiach A, et al. Transcriptomic diversity of cell types across the adult human brain. Science. 2023;382:eadd7046. doi:10.1126/science.add7046.

15. Human Protein Atlas. Brain data version 25.1. Accessed 9 September 2026. https://www.proteinatlas.org/humanproteome/brain/data.

16. Allen Institute. Allen Human Brain Atlas API documentation. Accessed 9 September 2026. https://brain-map.org/support/documentation/human-brain-atlas-api.

17. Allen Institute. Terms of Use and Citation Policy. Accessed 9 September 2026. https://alleninstitute.org/legal/terms-of-use.

18. National Center for Biotechnology Information. Gene Expression Omnibus platform GPL570: Affymetrix Human Genome U133 Plus 2.0 Array. Accessed 9 September 2026. https://www.ncbi.nlm.nih.gov/geo/query/acc.cgi?acc=GPL570.

