## Supplementary Archive S1 for "An anatomy-constrained transcriptomic screen identifies a 19-gene panel with an ordered regional similarity pattern in the human somatosensory system": PUBLIC_RELEASE_AUDIT_A05.md

# File-level release audit — A05

**Audit date:** 9 September 2026  
**Decision:** PASS for the stated derivative-publication boundary; the archive is not represented as public-data-only

## Governing boundary

The investigator authorizes analysis of the internally maintained, processed BIGE extract and publication of aggregate statistics, tables, figures, candidate lists, and other non-reconstructive derivatives. The complete internal matrix and exports that substantially reconstruct it remain restricted.

## Allowlisted content classes

- frozen candidate-gene lists, tiers, route definitions, and anatomical mappings;
- aggregate and donor-level statistics used in the manuscript;
- null-distribution draws and panel-membership ledgers that contain no BIGE expression matrix;
- standardized balance diagnostics and selection-audit summaries;
- figures, captions, and figure source tables;
- executable code, protocols, seeds, schemas, environment records, and hashes;
- AHBA-derived analysis outputs already present in A04;
- the portable GSE3526 source-lineage audit package, which omits its separately downloadable public source matrix.

Some balance files contain standardized summary features whose column labels identify their private-source provenance. They do not contain the underlying regional-by-gene BIGE values and cannot reconstruct the complete internally maintained matrix.

## Denied content classes

- the complete internally maintained BIGE workbook or matrix;
- complete or near-complete transformed regional-by-gene exports;
- serialized objects, caches, or intermediate arrays containing restricted expression values;
- credentials, authentication material, absolute private source paths, and local caches;
- the 85,860,948-byte GSE3526 series matrix, which remains available from GEO and is acquired by a hash-gated downloader.

## Checks required for the sealed ZIP

The finalization script must verify all of the following before the archive is sealed:

1. no spreadsheet, NumPy/R, HDF5, Parquet, database, or other plausible full-matrix container is present;
2. no file exceeds 20 MB;
3. the nested GSE3526 ZIP contains 46 files and no source matrix, absolute path, Python cache, or compiled bytecode;
4. every packaged file except `MANIFEST_SHA256.txt` is represented exactly once in the package manifest;
5. every manifest digest validates after ZIP extraction;
6. frozen v1.0/v1.1 inferential code and result files are byte-identical to A04, excluding only the intentionally replaced manuscript and report;
7. draft v1.2 files are visibly labelled prospective and are not represented as executed evidence.

## Recorded finalization results

- `101/101` inherited scientific and provenance files compared against A04 were identical byte for byte; the superseded manuscript v0.2 and report A04 were intentionally replaced, and package-level release records were regenerated.
- The nested GSE3526 archive passed compressed-data integrity, contained exactly 46 files, and contained no source series matrix, absolute member path, Python cache, or compiled bytecode.
- The nested archive was `gse3526_donor_reconstruction_v2_A02.zip` (SHA-256 `4edd3d126023e730ae8c93bcf6f74bb193237524429c20d5108e63487ffadfae`). Its documentary correction distinguishes 193 neural arrays across all ten donors from the 180 arrays belonging to the eight complete donors; it does not claim numerical equivalence to the internally analyzed BIGE aggregate.
- No packaged file used a denied matrix-like container extension and no packaged file exceeded 20 MiB.
- Text inspection found no credential, user-workspace path, or absolute path to restricted source data. The inherited A04 environment record preserves non-sensitive ephemeral build-toolchain paths under `/opt`, `/project`, and `/tmp`; these are retained byte for byte as computational provenance and do not identify or locate the internal BIGE matrix.
- `protocol_v1_2_draft.json` parsed successfully and the v1.2 records remained explicitly draft and prospective.
- The package manifest contained one entry for every packaged file except the manifest itself; all digests validated.
- `verification/verify_release_A05.py` returned `PASS` after the final package assembly.

Passing this audit establishes conformance to the investigator's stated release boundary for this archive. It is a project governance record, not an independent legal opinion or a grant of rights in third-party source resources.
