## Supplementary Archive S1 for "An anatomy-constrained transcriptomic screen identifies a 19-gene panel with an ordered regional similarity pattern in the human somatosensory system": RELEASE_NOTES.md

# DCML19 transcriptomic topology — v1.1-A05 release candidate

**Inferential analysis frozen:** 8 September 2026  
**Documentary and source-lineage audit update:** 9 September 2026  
**Status:** mixed-provenance analytical release candidate; the complete internally maintained BIGE matrix is excluded

## Scope of A05

A05 is a documentary, governance, and source-lineage-audit update. It does not alter the frozen v1.0 or v1.1 inferential code, inputs, null draws, statistics, or decisions released in A04.

The v1.1 spatial result remains `WHOLE_ONLY_SPATIAL_ROBUSTNESS`: the donor-by-compartment P1A gate failed (`2,558/50,000`, `p=0.0511789764`), the whole-donor P1B gate passed (`192/50,000`, `p=0.0038599228`), and the required conjunction failed with `p_IUT=0.0511789764`. The four secondary spatial sensitivities retain only nominal, unadjusted, model-specific status.

The defensible interpretation is a post-discovery transcriptomic adjacency contrast with favorable selection-aware calibrations and non-uniform spatial support. No result establishes anatomical connectivity, molecular flow, direction, causality, DCML exclusivity, a fibromyalgia or pain mechanism, a stimulation target, or independent confirmation.

## BIGE source and publication boundary

Discovery used **an internally maintained, processed extract of the Body Index of Gene Expression (BIGE)**. Investigator attestation authorizes analysis and publication of aggregate statistics, tables, figures, candidate lists, and other non-reconstructive analytical derivatives. It does not authorize redistribution of the complete internally maintained matrix or an export that substantially reconstructs it.

The provenance record links the resource to Lee et al. (2005), Roth et al. (2006), and Ushach et al. (2015). The 353-array, 65-tissue cohort described by Roth et al., including 20 CNS regions, is publicly deposited as GEO GSE3526. GSE3526 is a public cohort in the documented historical source lineage and a reconstruction source, not the entirety of the internally maintained BIGE extract and not an independent validation cohort. Formal sample- and processing-level reconciliation has not established numerical equivalence between the GSE3526-derived aggregate and the internally analyzed extract.

This archive is deliberately described as mixed-provenance. A file-level audit found no complete or near-complete BIGE matrix, workbook, serialized expression object, restricted raw cache, or reconstructive regional export. See `PUBLIC_RELEASE_AUDIT_A05.md` and `work/governance/BIGE_source_and_governance_record_v1.md`.

## GSE3526 source-lineage audit

A corrected portable package is included at `work/gse3526/gse3526_donor_reconstruction_v2_A02.zip`. It downloads the official series matrix with a fail-closed size and SHA-256 gate, records the computational environment, uses relative manifests, and reproduced 22/22 scientific outputs byte for byte across two clean executions. The 85,860,948-byte public source matrix is not embedded in either ZIP. The series contains 193 arrays from the frozen neural universe across all ten donors; 180 belong to the eight donors with complete endpoints and can influence their within-donor standardized profiles.

Across eight donors with complete broad-region coverage, the frozen 19-gene panel produced positive mean relative serial-order contrasts under three probe-collapse methods. This source-lineage result is descriptive and post-selection. Most three-node pairwise mean correlations were negative; the positive contrast primarily means that selected adjacent pairs were less dissimilar than the selected non-adjacent pair. The result depended materially on the frozen neural background used for within-donor standardization.

The three-node contrast also showed marked post hoc heterogeneity associated with recorded donor sex (`n=4+4`). This cannot be interpreted as a biological sex effect because donor identity is confounded and relevant covariates are unavailable. A 17-gene sensitivity excluding historically ambiguous RAB3GAP2/AURKAPS1 and C7orf25/PSMA2 probe assignments retained the direction but did not resolve probe attribution or create confirmatory evidence.

## New or updated records

- English working manuscript v0.3.
- Spanish integrated scientific report v1.1-A05.
- BIGE provenance, authorization, and release-boundary record.
- Corrected GSE3526 reconstruction package v2_A02.
- Supplementary Figure S1 in PNG, PDF, and SVG, with caption, source data, and plotting script.
- Draft v1.2 protocol, feasibility record, and adversarial root audit. Version 1.2 remains prospective: it is neither frozen nor executed.

## Integrity

From the extracted package root, run:

```bash
sha256sum -c MANIFEST_SHA256.txt
```

The expected result is one `OK` line for every listed file and exit status zero. The package manifest covers every file except itself. The release ZIP has a separate companion SHA-256 record.
