## Supplementary Archive S1 for "An anatomy-constrained transcriptomic screen identifies a 19-gene panel with an ordered regional similarity pattern in the human somatosensory system": verify_release_A05.py

```
#!/usr/bin/env python3
"""Fail-closed integrity and release-boundary checks for the A05 archive."""

from __future__ import annotations

import hashlib
import json
from pathlib import Path, PurePosixPath
import sys
import zipfile


ROOT = Path(__file__).resolve().parents[1]
MANIFEST = ROOT / "MANIFEST_SHA256.txt"
GSE_ZIP = ROOT / "work/gse3526/gse3526_donor_reconstruction_v2_A02.zip"
PROTOCOL = ROOT / "work/route_v1_2_draft/protocol/protocol_v1_2_draft.json"
EXPECTED_GSE_ZIP_SHA256 = (
    "4edd3d126023e730ae8c93bcf6f74bb193237524429c20d5108e63487ffadfae"
)
EXPECTED_GSE_FILES = 46
MAX_FILE_BYTES = 20 * 1024 * 1024
DENIED_SUFFIXES = {
    ".xlsx",
    ".xls",
    ".npz",
    ".npy",
    ".rds",
    ".rdata",
    ".h5",
    ".hdf5",
    ".parquet",
    ".feather",
    ".sqlite",
    ".db",
}


def sha256(path: Path) -> str:
    digest = hashlib.sha256()
    with path.open("rb") as handle:
        for block in iter(lambda: handle.read(1024 * 1024), b""):
            digest.update(block)
    return digest.hexdigest()


def fail(message: str) -> None:
    raise RuntimeError(message)


def verify_manifest() -> None:
    listed: dict[Path, str] = {}
    for line_number, raw in enumerate(MANIFEST.read_text(encoding="utf-8").splitlines(), 1):
        digest, marker, rel = raw.partition("  ")
        if not marker or len(digest) != 64 or not rel.startswith("./"):
            fail(f"malformed manifest line {line_number}")
        path = ROOT / rel[2:]
        if path in listed:
            fail(f"duplicate manifest path: {rel}")
        listed[path] = digest

    actual = {p for p in ROOT.rglob("*") if p.is_file() and p != MANIFEST}
    if set(listed) != actual:
        missing = sorted(str(p.relative_to(ROOT)) for p in actual - set(listed))
        stale = sorted(str(p.relative_to(ROOT)) for p in set(listed) - actual)
        fail(f"manifest coverage mismatch; missing={missing}; stale={stale}")

    for path, expected in listed.items():
        if sha256(path) != expected:
            fail(f"digest mismatch: {path.relative_to(ROOT)}")


def verify_release_boundary() -> None:
    for path in ROOT.rglob("*"):
        if not path.is_file():
            continue
        if path.suffix.lower() in DENIED_SUFFIXES:
            fail(f"denied matrix-like container: {path.relative_to(ROOT)}")
        if path.stat().st_size > MAX_FILE_BYTES:
            fail(f"file exceeds 20 MiB: {path.relative_to(ROOT)}")


def verify_nested_gse_zip() -> None:
    if sha256(GSE_ZIP) != EXPECTED_GSE_ZIP_SHA256:
        fail("GSE3526 package digest mismatch")
    with zipfile.ZipFile(GSE_ZIP) as archive:
        files = [item for item in archive.infolist() if not item.is_dir()]
        if len(files) != EXPECTED_GSE_FILES:
            fail(f"GSE3526 nested file count is {len(files)}, expected {EXPECTED_GSE_FILES}")
        bad: list[str] = []
        for item in files:
            name = item.filename
            path = PurePosixPath(name)
            lower = name.lower()
            if path.is_absolute() or ".." in path.parts:
                bad.append(name)
            elif "__pycache__" in path.parts or lower.endswith(".pyc"):
                bad.append(name)
            elif "series_matrix" in lower:
                bad.append(name)
        if bad:
            fail(f"disallowed nested GSE3526 members: {bad}")


def verify_protocol_status() -> None:
    record = json.loads(PROTOCOL.read_text(encoding="utf-8"))
    serialized = json.dumps(record, ensure_ascii=False).lower()
    if "draft" not in serialized or "prospective" not in serialized:
        fail("v1.2 protocol is not visibly labelled draft and prospective")


def main() -> int:
    verify_manifest()
    verify_release_boundary()
    verify_nested_gse_zip()
    verify_protocol_status()
    print("PASS: A05 manifest, release boundary, nested GSE3526 package, and draft status")
    return 0


if __name__ == "__main__":
    try:
        sys.exit(main())
    except Exception as exc:  # fail closed with a concise audit message
        print(f"FAIL: {exc}", file=sys.stderr)
        sys.exit(1)
```
