## Supplementary Archive S1 for "An anatomy-constrained transcriptomic screen identifies a 19-gene panel with an ordered regional similarity pattern in the human somatosensory system": BIGE_source_and_governance_record_v1.md

# BIGE source, authorization, and release-boundary record

**Record date:** 2026-09-09  
**Status:** analytical and publication use authorized by investigator attestation; full internally maintained matrix restricted from redistribution  
**Applies to:** transcriptomic relay-region project, candidate-discovery layer and its derived outputs

## Investigator attestation

The project lead confirmed that the internally maintained Body Index of Gene Expression (BIGE) matrix may be analyzed and that statistics, tables, and newly generated figures derived from it may be published. The complete internally maintained database must not be redistributed.

This attestation resolves the project's analytical-use and publication-authority gate for non-disclosive derived results. It does not authorize publication of the full workbook, a complete transformed matrix, or another release that would effectively reproduce the restricted database.

## Source provenance

BIGE is an Affymetrix Human Genome U133 Plus 2.0 expression compendium of normal human tissues and cell types. Its construction and use were described in:

1. Lee J, Hever A, Willhite D, Zlotnik A, Hevezi P. Effects of RNA degradation on gene expression analysis of human postmortem tissues. *FASEB Journal*. 2005;19:1356–1358. https://doi.org/10.1096/fj.04-3552fje
2. Roth RB, Hevezi P, Lee J, Willhite D, Lechner SM, Foster AC, Zlotnik A. Gene expression analyses reveal molecular relationships among 20 regions of the human CNS. *Neurogenetics*. 2006;7:67–80. https://doi.org/10.1007/s10048-006-0032-6
3. Ushach I, Burkhardt AM, Martinez C, et al. METEORIN-LIKE is a cytokine associated with barrier tissues and alternatively activated macrophages. *Clinical Immunology*. 2015;156:119–127. https://doi.org/10.1016/j.clim.2014.11.006

Ushach et al. describe BIGE as spanning 105 human tissues and cell types. Roth et al. describe a 353-array component spanning 65 normal tissues, including 20 CNS regions and 45 non-CNS tissues from ten post-mortem donors. That component is publicly archived at NCBI GEO as [GSE3526](https://www.ncbi.nlm.nih.gov/geo/query/acc.cgi?acc=GSE3526).

GSE3526 is therefore a public reconstruction source and discovery audit for the Roth component. It is not an independent biological replication of a panel selected using the internally maintained BIGE-derived regional summaries.

[GSE7307](https://www.ncbi.nlm.nih.gov/geo/query/acc.cgi?acc=GSE7307) is a later public series titled *Human body index — transcriptional profiling*, on the same array platform and from the same submitting group. Its exact identity or overlap with the internally maintained 105-tissue BIGE matrix has not been established in this project and must not be asserted without a sample/manifest-level reconciliation.

## Publicly permitted project outputs

- candidate-gene lists and their frozen selection labels;
- anatomical mappings and route definitions;
- summary statistics and inferential results;
- newly generated tables and figures;
- non-disclosive regional summaries needed to support a reported result;
- analysis code, schemas, protocols, seeds, and synthetic tests that do not embed the complete restricted matrix;
- the public GSE3526 donor-level reconstruction and its derived outputs, subject to NCBI attribution and any applicable third-party rights.

## Restricted outputs

- the complete internally maintained BIGE workbook or matrix;
- complete or near-complete transformed exports that reproduce that matrix;
- raw restricted files, caches, or intermediate objects containing the matrix;
- any package described as fully public-data-only if it includes BIGE-derived restricted material.

## Manuscript wording boundary

The source should be described as an **internally maintained, processed BIGE extract**, not as an anonymous private atlas and not as a wholly public dataset. The manuscript may state that the authors are authorized to publish derived summary statistics and figures while the complete processed matrix is not redistributed.

The manuscript must not state that all BIGE data are available under GSE3526. GSE3526 should be identified specifically as the public 65-tissue, 353-array component described by Roth et al.; the relationship of the expanded 105-tissue resource to other GEO series remains a provenance question.

## Operational consequence

The v1.2 private-input governance blocker is resolved for internal analysis and publication of permitted derived outputs. Before any public package is released, a file-level audit must confirm that the complete internally maintained matrix and reconstructive derivatives are absent. This is a data-governance record for the project and not an independent legal opinion or a substitute for journal-specific disclosure requirements.
