## Supplementary Archive S1 for "An anatomy-constrained transcriptomic screen identifies a 19-gene panel with an ordered regional similarity pattern in the human somatosensory system": README.md

# GSE3526 donor-level source-lineage audit

`gse3526_donor_reconstruction_v2_A02.zip` is a portable, hash-gated reconstruction and sensitivity package for the public GSE3526 cohort.

- Package SHA-256: `4edd3d126023e730ae8c93bcf6f74bb193237524429c20d5108e63487ffadfae`
- Official matrix expected size: `85,860,948` bytes
- Official matrix SHA-256: `77cc18cb606adb8028e0ef8e84db288f26b160308a364089c3876c0335c4b7d0`
- Clean-run reproducibility: `22/22` scientific outputs identical byte for byte across two executions
- Neural-background accounting: `193` arrays across all ten donors; `180` among the eight complete donors can influence the analyzed within-donor profiles

The source matrix is deliberately excluded and is downloaded from the frozen official GEO route by the included helper. This analysis is a donor-level audit of a public cohort in the documented historical source lineage of BIGE. Formal sample- and processing-level reconciliation has not established numerical equivalence to the internally analyzed BIGE aggregate. It is not an independent replication or confirmatory test.
