## Supplementary Archive S1 for "An anatomy-constrained transcriptomic screen identifies a 19-gene panel with an ordered regional similarity pattern in the human somatosensory system": analyze_v1_0_A03_reconstructed.py

```
#!/usr/bin/env python3
"""Evaluate frozen DCML19 v1.0 endpoints against pre-generated null panels.

The panel memberships are generated without reading AHBA topology residuals,
sealed by SHA-256, and consumed here only after protocol-chain verification.
All v1.0 results remain post-v0.11 robustness analyses, not independent
confirmation.
"""

from __future__ import annotations

import gzip
import hashlib
import json
import math
import platform
import sys
from pathlib import Path
from typing import Any

import numpy as np
import pandas as pd
import scipy
from scipy.stats import binomtest, rankdata


HERE = Path(__file__).resolve().parents[1]
PROJECT = HERE.parents[1]
FINAL_OUT = HERE / "analysis"
STAGING_OUT = HERE / "analysis.staging"
NULL_DIR = HERE / "null_design_final"
FAILED_NULL_DIR = HERE / "null_design"
NULL_GENERATOR_PATH = HERE / "scripts" / "generate_conditional_pseudoroute_null_v1_0.py"

PROTOCOL_PATH = HERE / "protocol" / "protocol_v1_0.json"
AMENDMENT1_PATH = HERE / "protocol" / "amendment_01_selection_replay_v1_0.json"
AMENDMENT2_PATH = HERE / "protocol" / "amendment_02_blind_feasibility_v1_0.json"
AMENDMENT3_PATH = HERE / "protocol" / "amendment_03_endpoint_seal_v1_0.json"

V10 = PROJECT / "reconstructed_v010" / "Paquete_reproducible_rutas_transcriptomicas_v0_10"
V11 = (
    PROJECT
    / "reconstructed_v011"
    / "Paquete_reproducible_rutas_transcriptomicas_v0_11"
    / "work"
    / "route_v0_11"
)
V08 = PROJECT / "reconstructed_v008" / "Paquete_reproducible_rutas_transcriptomicas_v0_8"
V04 = PROJECT / "reconstructed_v004"
V06 = PROJECT / "reconstructed_v006" / "Paquete_reproducible_rutas_transcriptomicas_v0_6"

INPUTS = {
    "v10_pools": V10 / "analysis" / "matched_final_pools_v0_10.csv",
    "v10_ahba_residuals": V10
    / "analysis"
    / "primary_node_residuals_candidates_and_controls_v0_10.csv.gz",
    "v11_cell_contrasts": V11
    / "analysis"
    / "all_candidate_control_cell_contrasts_v0_11.csv.gz",
    "v10_triplets": V10 / "analysis" / "alternative_triplet_statistics_v0_10.csv",
    "v10_groups": V10 / "analysis" / "alternative_node_groups_v0_10.csv",
    "v08_sample_metadata": V08 / "analysis" / "ahba_sample_metadata_v0_8.csv",
    "v04_hpa_exact_residuals": V04 / "analysis" / "hpa_node_scores_common_genes.csv",
    "v08_public_raw_manifest": V08 / "protocol" / "public_raw_manifest_v0_8.csv",
    "v08_probe_validation": V08 / "protocol" / "ahba_probe_validation_v0_8.json",
    "v10_primary_tests": V10 / "analysis" / "specificity_primary_tests_v0_10.csv",
}

DESIGN_INPUTS = {
    "v06_private_selection_metrics": V06
    / "analysis"
    / "routewide_gene_metrics_all_v0_6.csv",
    "v06_private_gene_matrix": V06
    / "frozen_inputs"
    / "gene_expression_private_v0_2.npz",
    "v06_private_gene_qc": V06 / "frozen_inputs" / "gene_collapse_qc.csv",
}

PANEL_PATHS = {
    "P1_technical": NULL_DIR
    / "conditional_pseudoroute_panels_P1_technical_v1_0.csv.gz",
    "S1_neuron": NULL_DIR
    / "conditional_pseudoroute_panels_S1_neuron_v1_0.csv.gz",
}

NODES = ["Cu/Gr", "DTLv", "Postcentral"]
DONORS = [9861, 10021, 12876, 14380, 15496, 15697]
CELL_REGIONS = ["Medulla oblongata", "Thalamus", "Cerebral cortex"]
N_DRAWS = 50_000
GENE_COLUMNS = [f"gene_{index:02d}" for index in range(1, 20)]
QUANTILE_METHOD = "linear"


def sha256(path: Path) -> str:
    hasher = hashlib.sha256()
    with path.open("rb") as handle:
        for block in iter(lambda: handle.read(1024 * 1024), b""):
            hasher.update(block)
    return hasher.hexdigest()


def parse_hash_manifest(path: Path) -> dict[str, str]:
    result: dict[str, str] = {}
    for line in path.read_text(encoding="utf-8").splitlines():
        if not line.strip():
            continue
        digest, filename = line.split("  ", 1)
        if len(digest) != 64 or any(character not in "0123456789abcdef" for character in digest):
            raise RuntimeError(f"Malformed SHA-256 in manifest: {filename}")
        if Path(filename).name != filename:
            raise RuntimeError(f"Unsafe path in hash manifest: {filename}")
        if filename in result:
            raise RuntimeError(f"Duplicate hash entry: {filename}")
        result[filename] = digest
    return result


def verify_hash_manifest(directory: Path, manifest_path: Path) -> pd.DataFrame:
    expected = parse_hash_manifest(manifest_path)
    actual = {
        path.name
        for path in directory.iterdir()
        if path.is_file() and path.resolve() != manifest_path.resolve()
    }
    if set(expected) != actual:
        raise RuntimeError(
            "Sealed null directory membership differs from its frozen manifest"
        )
    rows = []
    for filename, wanted in expected.items():
        path = directory / filename
        if not path.is_file():
            raise FileNotFoundError(path)
        observed = sha256(path)
        if observed != wanted:
            raise RuntimeError(f"Hash mismatch for sealed null file {filename}")
        rows.append(
            {
                "source_id": f"sealed_null::{filename}",
                "path": str(path.relative_to(PROJECT)),
                "bytes": path.stat().st_size,
                "sha256": observed,
                "verified_before_endpoint": True,
            }
        )
    return pd.DataFrame(rows)


def verify_inputs(
    protocol: dict[str, Any],
    amendment1: dict[str, Any],
    amendment2: dict[str, Any],
    amendment3: dict[str, Any],
) -> pd.DataFrame:
    chain = {
        "protocol_v1_0": (PROTOCOL_PATH, amendment1["parent_protocol_sha256"]),
        "amendment_v1_0_A01": (
            AMENDMENT1_PATH,
            amendment2["parent_amendment_A01_sha256"],
        ),
        "amendment_v1_0_A02": (
            AMENDMENT2_PATH,
            amendment3["parent_amendment_A02_sha256"],
        ),
    }
    if sha256(PROTOCOL_PATH) != amendment2["parent_protocol_sha256"]:
        raise RuntimeError("A02 protocol parent changed")
    if sha256(PROTOCOL_PATH) != amendment3["parent_protocol_sha256"]:
        raise RuntimeError("A03 protocol parent changed")
    if sha256(AMENDMENT1_PATH) != amendment3["parent_amendment_A01_sha256"]:
        raise RuntimeError("A03 A01 parent changed")
    if sha256(Path(__file__)) != amendment3["endpoint_script_sha256"]:
        raise RuntimeError("Endpoint script changed after A03 seal")
    if amendment3.get("endpoint_read_or_computed_before_seal") is not False:
        raise RuntimeError("A03 does not attest outcome blinding through the seal")

    frozen_input_groups = [
        protocol["input_files"],
        amendment1["additional_input_files"],
        amendment3["additional_input_files"],
    ]
    identifiers = [[str(row["id"]) for row in group] for group in frozen_input_groups]
    for group_ids in identifiers:
        if len(group_ids) != len(set(group_ids)):
            raise RuntimeError("Duplicate input ID within a protocol section")
    all_identifiers = [source_id for group_ids in identifiers for source_id in group_ids]
    if len(all_identifiers) != len(set(all_identifiers)):
        raise RuntimeError("Input ID reused across protocol/amendment sections")
    expected = {
        row["id"]: row for group in frozen_input_groups for row in group
    }
    actual_inputs = {**INPUTS, **DESIGN_INPUTS}
    if set(actual_inputs) != set(expected):
        missing = sorted(set(expected) - set(actual_inputs))
        unexpected = sorted(set(actual_inputs) - set(expected))
        raise RuntimeError(
            f"Frozen input coverage mismatch; missing={missing}, unexpected={unexpected}"
        )
    rows = []
    for source_id, path in actual_inputs.items():
        if not path.is_file():
            raise FileNotFoundError(path)
        observed = sha256(path)
        if observed != expected[source_id]["sha256"]:
            raise RuntimeError(f"Input hash mismatch: {source_id}")
        if "bytes" in expected[source_id] and path.stat().st_size != int(
            expected[source_id]["bytes"]
        ):
            raise RuntimeError(f"Input byte-count mismatch: {source_id}")
        rows.append(
            {
                "source_id": source_id,
                "path": str(path.relative_to(PROJECT)),
                "bytes": path.stat().st_size,
                "sha256": observed,
                "verified_before_endpoint": True,
            }
        )

    for source_id, (path, wanted) in chain.items():
        observed = sha256(path)
        if observed != wanted:
            raise RuntimeError(f"Protocol chain mismatch: {source_id}")
        rows.append(
            {
                "source_id": source_id,
                "path": str(path.relative_to(PROJECT)),
                "bytes": path.stat().st_size,
                "sha256": observed,
                "verified_before_endpoint": True,
            }
        )
    rows.append(
        {
            "source_id": "amendment_v1_0_A03",
            "path": str(AMENDMENT3_PATH.relative_to(PROJECT)),
            "bytes": AMENDMENT3_PATH.stat().st_size,
            "sha256": sha256(AMENDMENT3_PATH),
            "verified_before_endpoint": True,
        }
    )

    null_manifest = NULL_DIR / "null_design_hashes_v1_0.txt"
    if sha256(null_manifest) != amendment3["sealed_null_design"]["hash_manifest_sha256"]:
        raise RuntimeError("Null-design hash manifest changed after A03 seal")
    null_summary = NULL_DIR / "null_design_summary_v1_0.json"
    if sha256(null_summary) != amendment3["sealed_null_design"]["summary_sha256"]:
        raise RuntimeError("Null-design summary changed after A03 seal")
    for model_id, path in PANEL_PATHS.items():
        wanted = amendment3["sealed_null_design"]["panel_sha256"][model_id]
        if sha256(path) != wanted:
            raise RuntimeError(f"Sealed panel changed after A03: {model_id}")
    generator_sha = amendment3["sealed_null_design"]["generator_script_sha256"]
    if sha256(NULL_GENERATOR_PATH) != generator_sha:
        raise RuntimeError("Null-generator script changed after A03 seal")
    rows.extend(verify_hash_manifest(NULL_DIR, null_manifest).to_dict("records"))
    rows.append(
        {
            "source_id": "sealed_null::null_design_hashes_v1_0.txt",
            "path": str(null_manifest.relative_to(PROJECT)),
            "bytes": null_manifest.stat().st_size,
            "sha256": sha256(null_manifest),
            "verified_before_endpoint": True,
        }
    )
    failed_manifest = FAILED_NULL_DIR / "null_design_hashes_v1_0.txt"
    failed_summary = FAILED_NULL_DIR / "null_design_summary_v1_0.json"
    failed_record = amendment2["failed_blind_design_record"]
    for source_id, path, wanted in [
        (
            "failed_blind_design::hash_manifest",
            failed_manifest,
            failed_record["null_design_hash_manifest_sha256"],
        ),
        (
            "failed_blind_design::summary",
            failed_summary,
            failed_record["null_design_summary_sha256"],
        ),
        ("sealed_null::generator_script", NULL_GENERATOR_PATH, generator_sha),
    ]:
        observed = sha256(path)
        if observed != wanted:
            raise RuntimeError(f"Audit-record hash mismatch: {source_id}")
        rows.append(
            {
                "source_id": source_id,
                "path": str(path.relative_to(PROJECT)),
                "bytes": path.stat().st_size,
                "sha256": observed,
                "verified_before_endpoint": True,
            }
        )
    return pd.DataFrame(rows)


def verify_probe_files(
    candidate_order: list[str],
) -> tuple[pd.DataFrame, list[dict[str, Any]]]:
    manifest = pd.read_csv(INPUTS["v08_public_raw_manifest"])
    required_manifest = {"relative_path", "bytes", "sha256"}
    if not required_manifest.issubset(manifest.columns):
        raise RuntimeError("Malformed v0.8 public-raw manifest")
    if manifest["relative_path"].duplicated().any():
        raise RuntimeError("Duplicate path in v0.8 public-raw manifest")
    manifest = manifest.set_index("relative_path", drop=False)

    validation = json.loads(INPUTS["v08_probe_validation"].read_text(encoding="utf-8"))
    if not isinstance(validation, list):
        raise RuntimeError("Malformed AHBA probe-validation inventory")
    wanted_symbols = set(candidate_order)
    inventory: list[dict[str, Any]] = []
    for record in validation:
        symbol = str(record.get("symbol_requested", ""))
        if symbol not in wanted_symbols:
            continue
        if not record.get("api_success") or not record.get("exact_symbol_confirmed"):
            raise RuntimeError(f"Unvalidated frozen probe for {symbol}")
        if str(record.get("returned_symbols")) != symbol:
            raise RuntimeError(f"AHBA returned-symbol mismatch for {symbol}")
        relative = str(record.get("raw_path", ""))
        probe_id = int(record["probe_id"])
        expected_name = f"{symbol}__{probe_id}.json.gz"
        if Path(relative).name != expected_name:
            raise RuntimeError(f"Probe filename mismatch: {relative}")
        inventory.append(
            {
                "symbol": symbol,
                "probe_id": probe_id,
                "relative_path": relative,
                "expected_sample_count": int(record["sample_count"]),
            }
        )
    inventory.sort(key=lambda row: (row["symbol"], row["probe_id"]))
    if len(inventory) != 44:
        raise RuntimeError(f"Expected 44 frozen DCML19 probes, found {len(inventory)}")
    if len({row["relative_path"] for row in inventory}) != len(inventory):
        raise RuntimeError("Duplicate path in probe-validation inventory")
    counts = pd.Series([row["symbol"] for row in inventory]).value_counts()
    if set(counts.index) != wanted_symbols or (counts < 1).any():
        raise RuntimeError(f"Unexpected frozen-probe coverage: {counts.to_dict()}")

    rows = []
    for item in inventory:
        relative = item["relative_path"]
        if relative not in manifest.index:
            raise RuntimeError(f"Probe absent from public-raw manifest: {relative}")
        record = manifest.loc[relative]
        path = V08 / relative
        if not path.is_file():
            raise FileNotFoundError(path)
        observed = sha256(path)
        if observed != str(record["sha256"]) or path.stat().st_size != int(
            record["bytes"]
        ):
            raise RuntimeError(f"Frozen probe file changed: {relative}")
        rows.append(
            {
                "source_id": f"v08_probe::{item['symbol']}::{item['probe_id']}",
                "path": str(path.relative_to(PROJECT)),
                "bytes": path.stat().st_size,
                "sha256": observed,
                "verified_before_endpoint": True,
            }
        )
    return pd.DataFrame(rows), inventory


def load_sealed_panels(
    candidate_order: list[str], amendment3: dict[str, Any]
) -> tuple[dict[str, pd.DataFrame], list[str]]:
    summary = json.loads((NULL_DIR / "null_design_summary_v1_0.json").read_text())
    if summary.get("endpoint_read_or_computed") is not False:
        raise RuntimeError("Sealed null summary does not attest outcome blinding")
    if summary.get("parent_protocol_sha256") != sha256(PROTOCOL_PATH):
        raise RuntimeError("Null summary has wrong protocol parent")
    if summary.get("amendment_A01_sha256") != sha256(AMENDMENT1_PATH):
        raise RuntimeError("Null summary has wrong A01 parent")
    if summary.get("amendment_A02_sha256") != sha256(AMENDMENT2_PATH):
        raise RuntimeError("Null summary has wrong A02 parent")
    if summary.get("generator_script_sha256") != amendment3["sealed_null_design"].get(
        "generator_script_sha256"
    ):
        raise RuntimeError("Null summary and A03 disagree on generator script")
    if summary.get("candidate_symbols") != candidate_order:
        raise RuntimeError("Null summary candidate order differs from protocol")
    if summary.get("eligible_pseudoroute_count") != 134:
        raise RuntimeError("Unexpected eligible-pseudoroute count")
    strict = summary.get("strict_replay_support", {})
    if strict.get("strict_total_A0_B3_C16_pseudoroutes") != 5 or strict.get(
        "strict_total_and_exact_strata_pseudoroutes"
    ) != 0:
        raise RuntimeError("Strict-replay support diagnostic changed")
    if summary.get("target_draws_per_model") != N_DRAWS:
        raise RuntimeError("Unexpected sealed target draw count")
    for model_id in PANEL_PATHS:
        model = summary["models"][model_id]
        if model["status"] != "READY_FOR_ENDPOINT":
            raise RuntimeError(f"Sealed null is not ready: {model_id}")
        if model["endpoint_read_or_computed"]:
            raise RuntimeError("Null design claims prior endpoint access")
        if model.get("accepted_panels") != N_DRAWS:
            raise RuntimeError(f"Wrong accepted-panel count: {model_id}")
        if model.get("structural_qc_pass") is not True or model.get(
            "ensemble_qc_pass"
        ) is not True:
            raise RuntimeError(f"Sealed QC did not pass: {model_id}")
        diversity = model.get("route_diversity", {})
        if diversity.get("diversity_pass") is not True:
            raise RuntimeError(f"Sealed route-diversity QC did not pass: {model_id}")
    s1_identity = summary["models"]["S1_neuron"].get(
        "identical_to_failed_design_where_A02_requires"
    )
    if not isinstance(s1_identity, dict) or not s1_identity or not all(
        value is True for value in s1_identity.values()
    ):
        raise RuntimeError("S1 identity requirement from A02 is not documented")
    controls = list(summary["complete_control_symbols"])
    if (
        summary.get("complete_control_count") != 742
        or len(controls) != 742
        or len(set(controls)) != 742
    ):
        raise RuntimeError("Malformed sealed control universe")
    panels: dict[str, pd.DataFrame] = {}
    candidate_set = set(candidate_order)
    control_set = set(controls)
    expected_columns = ["draw_id", "pseudo_route", "pseudo_route_regions", *GENE_COLUMNS]
    membership = pd.read_csv(
        NULL_DIR / "eligible_pseudoroute_stratum_membership_v1_0.csv.gz",
        keep_default_na=False,
    )
    required_membership = {
        "route_id",
        "tier",
        "exact_ahba_probe_count",
        "control",
    }
    if not required_membership.issubset(membership.columns):
        raise RuntimeError("Malformed sealed pseudoroute-membership table")
    if membership[["route_id", "control"]].duplicated().any():
        raise RuntimeError("Duplicate route/control membership")
    eligible_routes = set(membership["route_id"].astype(str))
    if len(eligible_routes) != 134:
        raise RuntimeError("Membership table does not contain 134 eligible routes")
    if not set(membership["control"].astype(str)).issubset(control_set):
        raise RuntimeError("Membership table contains a symbol outside the control universe")
    membership = membership.copy()
    membership["stratum"] = (
        membership["tier"].astype(str)
        + "|probe"
        + membership["exact_ahba_probe_count"].astype(int).astype(str)
    )
    stratum_by_route_gene = membership.set_index(["route_id", "control"])["stratum"]
    expected_strata = {str(key): int(value) for key, value in summary["candidate_strata"].items()}
    if sum(expected_strata.values()) != 19:
        raise RuntimeError("Sealed candidate strata do not sum to 19")
    for model_id, path in PANEL_PATHS.items():
        if sha256(path) != amendment3["sealed_null_design"]["panel_sha256"][model_id]:
            raise RuntimeError(f"Panel hash differs from A03: {model_id}")
        frame = pd.read_csv(path, keep_default_na=False)
        if frame.columns.tolist() != expected_columns:
            raise RuntimeError(f"Unexpected panel schema: {model_id}")
        if len(frame) != N_DRAWS or frame["draw_id"].tolist() != list(range(1, N_DRAWS + 1)):
            raise RuntimeError(f"Malformed draw IDs: {model_id}")
        if frame[["pseudo_route", "pseudo_route_regions", *GENE_COLUMNS]].eq("").any().any():
            raise RuntimeError(f"Blank value in sealed panels: {model_id}")
        if not set(frame["pseudo_route"].astype(str)).issubset(eligible_routes):
            raise RuntimeError(f"Ineligible pseudoroute in sealed panels: {model_id}")
        if not frame[GENE_COLUMNS].nunique(axis=1).eq(19).all():
            raise RuntimeError(f"Duplicate symbol within a panel: {model_id}")
        symbols = set(frame[GENE_COLUMNS].to_numpy().ravel())
        if not symbols.issubset(control_set) or symbols.intersection(candidate_set):
            raise RuntimeError(f"Invalid symbol in sealed panels: {model_id}")
        expected_route_count = summary["models"][model_id]["route_diversity"][
            "contributing_routes"
        ]
        if frame["pseudo_route"].nunique() != expected_route_count:
            raise RuntimeError(f"Route count differs from sealed summary: {model_id}")
        observed_strata = np.zeros(
            (N_DRAWS, len(expected_strata)), dtype=np.int16
        )
        stratum_names = list(expected_strata)
        route_values = frame["pseudo_route"].astype(str).to_numpy()
        for gene_column in GENE_COLUMNS:
            keys = pd.MultiIndex.from_arrays(
                [route_values, frame[gene_column].astype(str).to_numpy()],
                names=["route_id", "control"],
            )
            mapped = stratum_by_route_gene.reindex(keys)
            if mapped.isna().any():
                raise RuntimeError(
                    f"Panel symbol is not selected by its pseudoroute: {model_id}"
                )
            values = mapped.to_numpy(dtype=str)
            for stratum_index, stratum_name in enumerate(stratum_names):
                observed_strata[:, stratum_index] += values == stratum_name
        expected_matrix = np.asarray(
            [expected_strata[name] for name in stratum_names], dtype=np.int16
        )
        if not (observed_strata == expected_matrix).all():
            raise RuntimeError(f"Panel does not reproduce frozen strata: {model_id}")
        panels[model_id] = frame
    return panels, controls


def panels_to_indices(panels: pd.DataFrame, controls: list[str]) -> np.ndarray:
    index = {symbol: number for number, symbol in enumerate(controls)}
    draws = np.column_stack(
        [panels[column].map(index).to_numpy(dtype=np.int32) for column in GENE_COLUMNS]
    )
    if (draws < 0).any():
        raise RuntimeError("Unmapped sealed control symbol")
    return draws


def residual_tensor(frame: pd.DataFrame, symbols: list[str]) -> np.ndarray:
    pivot = frame.pivot(index="symbol", columns=["donor_id", "node"], values="node_residual_z")
    wanted = pd.MultiIndex.from_product([DONORS, NODES], names=["donor_id", "node"])
    values = pivot.reindex(index=symbols, columns=wanted).to_numpy(dtype=float)
    values = values.reshape(len(symbols), len(DONORS), len(NODES))
    if not np.isfinite(values).all():
        raise RuntimeError("Incomplete AHBA residual tensor")
    return values


def corr_rows(a: np.ndarray, b: np.ndarray) -> np.ndarray:
    a = a - a.mean(axis=1, keepdims=True)
    b = b - b.mean(axis=1, keepdims=True)
    denominator = np.sqrt(np.square(a).sum(axis=1) * np.square(b).sum(axis=1))
    return np.divide(
        (a * b).sum(axis=1),
        denominator,
        out=np.full(len(a), np.nan),
        where=denominator > 0,
    )


def spearman_three_columns(values: np.ndarray) -> tuple[float, float, float]:
    values = np.asarray(values, dtype=float)
    if values.ndim != 2 or values.shape[1] != 3 or not np.isfinite(values).all():
        raise RuntimeError("Spearman kernel requires a finite n-by-3 matrix")
    ranked = rankdata(values, axis=0, method="average")
    r01 = float(corr_rows(ranked[:, 0][None, :], ranked[:, 1][None, :])[0])
    r12 = float(corr_rows(ranked[:, 1][None, :], ranked[:, 2][None, :])[0])
    r02 = float(corr_rows(ranked[:, 0][None, :], ranked[:, 2][None, :])[0])
    if not np.isfinite([r01, r12, r02]).all():
        raise RuntimeError("Undefined Spearman correlation in observed matrix")
    return r01, r12, r02


def topology_observed(tensor: np.ndarray) -> tuple[pd.DataFrame, dict[str, Any]]:
    if tensor.shape != (19, len(DONORS), len(NODES)) or not np.isfinite(tensor).all():
        raise RuntimeError("Malformed observed AHBA tensor")
    rows = []
    for donor_index, donor in enumerate(DONORS):
        values = tensor[:, donor_index, :]
        r01, r12, r02 = spearman_three_columns(values)
        options = np.asarray(
            [
                (r01 + r02) / 2 - r12,
                (r01 + r12) / 2 - r02,
                (r02 + r12) / 2 - r01,
            ]
        )
        rows.append(
            {
                "donor_id": donor,
                "rho_CuGr_DTLv": r01,
                "rho_DTLv_Postcentral": r12,
                "rho_CuGr_Postcentral": r02,
                "adjacent_mean": (r01 + r12) / 2,
                "adjacent_bottleneck": min(r01, r12),
                "endpoint_similarity": r02,
                "adjacency_contrast": (r01 + r12) / 2 - r02,
                "both_adjacent_positive": bool(r01 > 0 and r12 > 0),
                "middle_is_best": bool(int(np.argmax(options)) == 1),
            }
        )
    frame = pd.DataFrame(rows)
    if not np.isfinite(frame.select_dtypes(include=[np.number]).to_numpy()).all():
        raise RuntimeError("Nonfinite observed topology")
    summary = {
        "mean_adjacency_contrast": float(frame["adjacency_contrast"].mean()),
        "median_adjacency_contrast": float(frame["adjacency_contrast"].median()),
        "positive_donor_count": int(frame["adjacency_contrast"].gt(0).sum()),
        "middle_best_donor_count": int(frame["middle_is_best"].sum()),
        "adjacent_mean": float(frame["adjacent_mean"].mean()),
        "adjacent_bottleneck": float(frame["adjacent_bottleneck"].mean()),
        "endpoint_similarity": float(frame["endpoint_similarity"].mean()),
        "both_adjacent_positive_donor_count": int(
            frame["both_adjacent_positive"].sum()
        ),
        "positive_donor_sign_test_one_sided_p": float(
            binomtest(
                int(frame["adjacency_contrast"].gt(0).sum()),
                n=len(DONORS),
                p=0.5,
                alternative="greater",
            ).pvalue
        ),
    }
    return frame, summary


def topology_null(draws: np.ndarray, residuals: np.ndarray) -> pd.DataFrame:
    if draws.shape != (N_DRAWS, 19):
        raise RuntimeError("Malformed sealed draw-index matrix")
    if residuals.shape != (residuals.shape[0], len(DONORS), len(NODES)):
        raise RuntimeError("Malformed control residual tensor")
    n = len(draws)
    donor_contrast = np.empty((n, len(DONORS)), dtype=float)
    donor_adjacent = np.empty((n, len(DONORS)), dtype=float)
    donor_bottleneck = np.empty((n, len(DONORS)), dtype=float)
    donor_endpoint = np.empty((n, len(DONORS)), dtype=float)
    donor_both_positive = np.empty((n, len(DONORS)), dtype=bool)
    donor_middle_best = np.empty((n, len(DONORS)), dtype=bool)
    for donor_index in range(len(DONORS)):
        ranked = [
            rankdata(residuals[draws, donor_index, node], axis=1, method="average")
            for node in range(3)
        ]
        r01 = corr_rows(ranked[0], ranked[1])
        r12 = corr_rows(ranked[1], ranked[2])
        r02 = corr_rows(ranked[0], ranked[2])
        options = np.column_stack(
            [
                (r01 + r02) / 2 - r12,
                (r01 + r12) / 2 - r02,
                (r02 + r12) / 2 - r01,
            ]
        )
        donor_adjacent[:, donor_index] = (r01 + r12) / 2
        donor_bottleneck[:, donor_index] = np.minimum(r01, r12)
        donor_endpoint[:, donor_index] = r02
        donor_contrast[:, donor_index] = options[:, 1]
        donor_both_positive[:, donor_index] = (r01 > 0) & (r12 > 0)
        donor_middle_best[:, donor_index] = np.argmax(options, axis=1) == 1
    frame = pd.DataFrame(
        {
            "draw_id": np.arange(1, n + 1),
            "mean_adjacency_contrast": donor_contrast.mean(axis=1),
            "median_adjacency_contrast": np.median(donor_contrast, axis=1),
            "positive_donor_count": (donor_contrast > 0).sum(axis=1),
            "middle_best_donor_count": donor_middle_best.sum(axis=1),
            "adjacent_mean": donor_adjacent.mean(axis=1),
            "adjacent_bottleneck": donor_bottleneck.mean(axis=1),
            "endpoint_similarity": donor_endpoint.mean(axis=1),
            "both_adjacent_positive_donor_count": donor_both_positive.sum(axis=1),
        }
    )
    for donor_index, donor in enumerate(DONORS):
        frame[f"donor_{donor}_adjacency_contrast"] = donor_contrast[:, donor_index]
    if not np.isfinite(frame.select_dtypes(include=[np.number]).to_numpy()).all():
        raise RuntimeError("Nonfinite topology null")
    return frame


def empirical_summary(
    null: np.ndarray,
    observed: float,
    tail: str = "upper",
    expected_n: int | None = None,
) -> dict[str, Any]:
    values = np.asarray(null, dtype=float)
    if values.ndim != 1:
        raise RuntimeError("Empirical null must be one-dimensional")
    if expected_n is not None and len(values) != expected_n:
        raise RuntimeError(
            f"Empirical null has {len(values)} values; expected {expected_n}"
        )
    if not np.isfinite(observed) or not np.isfinite(values).all():
        raise RuntimeError("Nonfinite empirical endpoint; denominator cannot be changed")
    if not len(values):
        raise RuntimeError("Empty empirical null")
    if tail == "upper":
        count = int((values >= observed).sum())
        threshold = float(np.quantile(values, 0.95, method=QUANTILE_METHOD))
        exceeds = bool(observed > threshold)
        threshold_name = "null_q95"
    elif tail == "lower":
        count = int((values <= observed).sum())
        threshold = float(np.quantile(values, 0.05, method=QUANTILE_METHOD))
        exceeds = bool(observed < threshold)
        threshold_name = "null_q05"
    else:
        raise ValueError(tail)
    return {
        "evaluable": True,
        "tail": tail,
        "observed": float(observed),
        "null_n": int(len(values)),
        "null_mean": float(values.mean()),
        "null_sd": float(values.std(ddof=1)),
        threshold_name: threshold,
        "tail_count": count,
        "empirical_p_one_sided": float((1 + count) / (1 + len(values))),
        "observed_beyond_95pct_threshold": exceeds,
        "quantile_method": QUANTILE_METHOD,
    }


def holm_adjust(pvalues: list[float]) -> list[float]:
    values = np.asarray(pvalues, dtype=float)
    if not np.isfinite(values).all():
        raise RuntimeError("Holm adjustment received nonfinite p-values")
    order = np.argsort(values, kind="stable")
    adjusted_sorted = np.maximum.accumulate(
        np.minimum(1.0, values[order] * (len(values) - np.arange(len(values))))
    )
    adjusted = np.empty_like(adjusted_sorted)
    adjusted[order] = adjusted_sorted
    return adjusted.tolist()


def topology_component_tests(
    observed: dict[str, Any], nulls: dict[str, pd.DataFrame]
) -> pd.DataFrame:
    components = [
        ("adjacent_mean", "upper"),
        ("adjacent_bottleneck", "upper"),
        ("endpoint_similarity", "lower"),
        ("both_adjacent_positive_donor_count", "upper"),
    ]
    rows = []
    for model_id, frame in nulls.items():
        model_rows = []
        for component, tail in components:
            result = empirical_summary(
                frame[component].to_numpy(dtype=float),
                float(observed[component]),
                tail,
                expected_n=N_DRAWS,
            )
            model_rows.append(
                {
                    "model_id": model_id,
                    "component": component,
                    "tail": tail,
                    **result,
                }
            )
        adjusted = holm_adjust(
            [row["empirical_p_one_sided"] for row in model_rows]
        )
        for row, p_adjusted in zip(model_rows, adjusted):
            row["holm_p_within_model"] = p_adjusted
            row["status"] = "DIAGNOSTIC_ONLY"
        rows.extend(model_rows)
    return pd.DataFrame(rows)


def lodo_null_tests(
    observed_donor: pd.DataFrame, nulls: dict[str, pd.DataFrame]
) -> pd.DataFrame:
    rows = []
    observed_contrast = observed_donor.set_index("donor_id")["adjacency_contrast"]
    for model_id, frame in nulls.items():
        donor_columns = [f"donor_{donor}_adjacency_contrast" for donor in DONORS]
        for omitted in DONORS:
            kept_donors = [donor for donor in DONORS if donor != omitted]
            kept_columns = [f"donor_{donor}_adjacency_contrast" for donor in kept_donors]
            observed = float(observed_contrast.loc[kept_donors].mean())
            null = frame[kept_columns].mean(axis=1).to_numpy(dtype=float)
            result = empirical_summary(
                null, observed, "upper", expected_n=N_DRAWS
            )
            rows.append(
                {
                    "model_id": model_id,
                    "omitted_donor": omitted,
                    **result,
                    "observed_positive": bool(observed > 0),
                    "status": "OVERLAPPING_DIAGNOSTIC_ONLY",
                }
            )
    return pd.DataFrame(rows)


def cell_attribution(
    cell: pd.DataFrame,
    candidate_order: list[str],
    controls: list[str],
    draws: np.ndarray,
) -> tuple[pd.DataFrame, dict[str, Any]]:
    pivot = (
        cell[cell["brain_region"].isin(CELL_REGIONS)]
        .pivot(
            index="symbol",
            columns="brain_region",
            values="neuronal_minus_non_neuronal_median",
        )
        .reindex(columns=CELL_REGIONS)
    )
    candidate = pivot.loc[candidate_order].to_numpy(dtype=float)
    control = pivot.loc[controls].to_numpy(dtype=float)
    if not np.isfinite(candidate).all() or not np.isfinite(control).all():
        raise RuntimeError("Cell-attribution matrix is incomplete")
    observed_regions = np.median(candidate, axis=0)
    observed = float(observed_regions.min())
    modules = np.median(control[draws], axis=1)
    statistics = modules.min(axis=1)
    frame = pd.DataFrame(
        {
            "draw_id": np.arange(1, len(draws) + 1),
            "medulla_oblongata_module": modules[:, 0],
            "thalamus_module": modules[:, 1],
            "cerebral_cortex_module": modules[:, 2],
            "weakest_region_statistic": statistics,
        }
    )
    summary = empirical_summary(
        statistics, observed, "upper", expected_n=N_DRAWS
    )
    summary.update(
        {
            "observed_medulla_oblongata": float(observed_regions[0]),
            "observed_thalamus": float(observed_regions[1]),
            "observed_cerebral_cortex": float(observed_regions[2]),
            "all_observed_regions_positive": bool((observed_regions > 0).all()),
            "status": "SUPPORTIVE_NOT_ROUTE_REPLICATION",
        }
    )
    return frame, summary


def geometry_analysis(
    groups: pd.DataFrame,
    triplets: pd.DataFrame,
    metadata: pd.DataFrame,
    expected_observed: float,
) -> tuple[pd.DataFrame, dict[str, Any]]:
    if not pd.api.types.is_bool_dtype(triplets["is_primary_triplet"]):
        raise RuntimeError("is_primary_triplet must be parsed as Boolean")
    if not pd.api.types.is_bool_dtype(groups["eligible"]):
        raise RuntimeError("eligible group flag must be parsed as Boolean")
    if len(triplets) != 240 or triplets["triplet_id"].nunique() != 240:
        raise RuntimeError("Geometry analysis requires 240 unique frozen triplets")
    primary_mask = triplets["is_primary_triplet"].eq(True)
    if int(primary_mask.sum()) != 1 or int((~primary_mask).sum()) != 239:
        raise RuntimeError("Geometry analysis requires one primary and 239 alternatives")
    expected_primary = "MYE_CuGr|THA_DTLv|PAR_postcentral_gyrus"
    if str(triplets.loc[primary_mask, "triplet_id"].iloc[0]) != expected_primary:
        raise RuntimeError("Unexpected primary anatomical triplet")

    eligible_groups = groups[groups["eligible"]].copy()
    if eligible_groups["group_id"].duplicated().any():
        raise RuntimeError("Duplicate eligible anatomical group ID")
    group_compartment = eligible_groups.set_index("group_id")["compartment"].to_dict()
    expected_compartments = {
        "myelencephalon_group": "myelencephalon",
        "thalamus_group": "dorsal thalamus",
        "parietal_group": "parietal lobe",
    }
    for column, compartment in expected_compartments.items():
        identifiers = set(triplets[column].astype(str))
        if not identifiers.issubset(group_compartment):
            raise RuntimeError(f"Triplet references an ineligible group: {column}")
        if any(group_compartment[identifier] != compartment for identifier in identifiers):
            raise RuntimeError(f"Triplet group has wrong compartment: {column}")

    metadata = metadata.copy()
    if metadata["sample_index"].duplicated().any():
        raise RuntimeError("Duplicate AHBA sample_index")
    metadata = metadata.set_index("sample_index", drop=False)
    left = metadata["hemisphere_class"].eq("left_or_unpaired")
    normalized: dict[int, pd.DataFrame] = {}
    for donor in DONORS:
        mask = left & metadata["donor_id"].eq(donor)
        coordinates = metadata.loc[mask, ["mri_x", "mri_y", "mri_z"]].astype(float)
        if coordinates.empty or not np.isfinite(coordinates.to_numpy()).all():
            raise RuntimeError(f"Incomplete left-sample coordinates for donor {donor}")
        scale = coordinates.std(axis=0, ddof=0)
        if not np.isfinite(scale.to_numpy()).all() or (scale == 0).any():
            raise RuntimeError(f"Invalid coordinate scale for donor {donor}")
        normalized[donor] = (coordinates - coordinates.mean(axis=0)) / scale

    centroids: dict[tuple[str, int], np.ndarray] = {}
    for record in eligible_groups.itertuples(index=False):
        indices = [int(value) for value in str(record.sample_indices).split(";") if value]
        if (
            not indices
            or len(indices) != len(set(indices))
            or len(indices) != int(record.total_target_samples)
        ):
            raise RuntimeError(f"Malformed target indices: {record.group_id}")
        if not set(indices).issubset(metadata.index):
            raise RuntimeError(f"Unknown target sample: {record.group_id}")
        selected = metadata.loc[indices]
        if not selected["hemisphere_class"].eq("left_or_unpaired").all():
            raise RuntimeError(f"Right-hemisphere target in group {record.group_id}")
        if not np.isfinite(
            selected[["mri_x", "mri_y", "mri_z"]].to_numpy(dtype=float)
        ).all():
            raise RuntimeError(f"Nonfinite target coordinate: {record.group_id}")
        if set(selected["donor_id"].astype(int)) != set(DONORS):
            raise RuntimeError(f"Target group lacks a frozen donor: {record.group_id}")
        for donor in DONORS:
            donor_indices = [
                index
                for index in indices
                if int(metadata.loc[index, "donor_id"]) == donor
            ]
            if not donor_indices:
                raise RuntimeError(f"Missing centroid samples: {record.group_id}, {donor}")
            centroid = normalized[donor].loc[donor_indices].mean(axis=0).to_numpy(float)
            if not np.isfinite(centroid).all():
                raise RuntimeError("Nonfinite centroid")
            centroids[(record.group_id, donor)] = centroid

    primary = triplets[primary_mask]
    primary = primary.iloc[0]
    primary_counts = np.asarray(
        [
            primary["myelencephalon_samples"],
            primary["thalamus_samples"],
            primary["parietal_samples"],
        ],
        dtype=float,
    )
    rows = []
    for record in triplets.itertuples(index=False):
        expected_counts = [
            int(eligible_groups.set_index("group_id").loc[record.myelencephalon_group, "total_target_samples"]),
            int(eligible_groups.set_index("group_id").loc[record.thalamus_group, "total_target_samples"]),
            int(eligible_groups.set_index("group_id").loc[record.parietal_group, "total_target_samples"]),
        ]
        recorded_counts = [
            int(record.myelencephalon_samples),
            int(record.thalamus_samples),
            int(record.parietal_samples),
        ]
        if recorded_counts != expected_counts:
            raise RuntimeError(f"Triplet sample-count mismatch: {record.triplet_id}")
        distances = []
        for donor in DONORS:
            a = centroids[(record.myelencephalon_group, donor)]
            b = centroids[(record.thalamus_group, donor)]
            c = centroids[(record.parietal_group, donor)]
            distances.append(
                [np.linalg.norm(a - b), np.linalg.norm(b - c), np.linalg.norm(a - c)]
            )
        distance_array = np.asarray(distances)
        counts = np.asarray(
            [
                record.myelencephalon_samples,
                record.thalamus_samples,
                record.parietal_samples,
            ],
            dtype=float,
        )
        rows.append(
            {
                "triplet_id": record.triplet_id,
                "is_primary_triplet": bool(record.is_primary_triplet),
                "mean_adjacency_contrast": float(record.mean_adjacency_contrast),
                "geometry_mean_d01": float(distance_array[:, 0].mean()),
                "geometry_mean_d12": float(distance_array[:, 1].mean()),
                "geometry_mean_d02": float(distance_array[:, 2].mean()),
                "log_sample_ratio_node1": float(np.log(counts[0] / primary_counts[0])),
                "log_sample_ratio_node2": float(np.log(counts[1] / primary_counts[1])),
                "log_sample_ratio_node3": float(np.log(counts[2] / primary_counts[2])),
            }
        )
    frame = pd.DataFrame(rows)
    if not np.isfinite(frame.select_dtypes(include=[np.number]).to_numpy()).all():
        raise RuntimeError("Nonfinite geometry feature or triplet statistic")
    features = [
        "geometry_mean_d01",
        "geometry_mean_d12",
        "geometry_mean_d02",
        "log_sample_ratio_node1",
        "log_sample_ratio_node2",
        "log_sample_ratio_node3",
    ]
    standardized = pd.DataFrame(index=frame.index)
    for feature in features:
        values = frame[feature].to_numpy(float)
        center = float(np.median(values))
        scale = float(np.median(np.abs(values - center)) * 1.4826)
        if not np.isfinite(scale) or scale == 0:
            scale = float(np.std(values, ddof=0))
        if not np.isfinite(scale) or scale == 0:
            raise RuntimeError(f"Cannot scale geometry feature: {feature}")
        standardized[feature] = (values - center) / scale
    primary_index = frame.index[frame["is_primary_triplet"]][0]
    delta = standardized - standardized.loc[primary_index]
    frame["geometry_sampling_distance_to_primary"] = np.sqrt(
        np.square(delta).sum(axis=1)
    )
    alternatives = frame[~frame["is_primary_triplet"]].sort_values(
        ["geometry_sampling_distance_to_primary", "triplet_id"], kind="mergesort"
    )
    observed = float(frame.loc[primary_index, "mean_adjacency_contrast"])
    if not math.isclose(observed, expected_observed, rel_tol=0.0, abs_tol=1e-12):
        raise RuntimeError("Precomputed primary-triplet statistic differs from endpoint")
    summaries = {}
    for size in [50, 25, 75, 100]:
        matched = alternatives.head(size)
        result = empirical_summary(
            matched["mean_adjacency_contrast"].to_numpy(float),
            observed,
            "upper",
            expected_n=size,
        )
        result.update(
            {
                "matched_set_size": size,
                "higher_or_equal_count": int(
                    matched["mean_adjacency_contrast"].ge(observed).sum()
                ),
            }
        )
        summaries[str(size)] = result
    frame["matched_alternative_rank"] = 0
    frame.loc[alternatives.index, "matched_alternative_rank"] = np.arange(1, 240)
    return frame.sort_values(
        ["geometry_sampling_distance_to_primary", "triplet_id"], kind="mergesort"
    ), summaries


def topology_single(values: np.ndarray) -> float:
    r01, r12, r02 = spearman_three_columns(values)
    return float((r01 + r12) / 2 - r02)


def topology_single_draws(values: np.ndarray, draws: np.ndarray) -> np.ndarray:
    ranked = [rankdata(values[draws, node], axis=1, method="average") for node in range(3)]
    return (
        corr_rows(ranked[0], ranked[1]) + corr_rows(ranked[1], ranked[2])
    ) / 2 - corr_rows(ranked[0], ranked[2])


def hpa_topology(
    hpa: pd.DataFrame,
    candidate_order: list[str],
    controls: list[str],
    draws_by_model: dict[str, np.ndarray],
) -> tuple[pd.DataFrame, dict[str, Any]]:
    wanted_nodes = ["Medulla", "Thalamus", "Parietal_lobe"]
    subset = hpa[
        hpa["route_id"].eq("DCML_CORE") & hpa["private_node"].isin(wanted_nodes)
    ]
    pivot = subset.pivot(
        index="private_symbol", columns="private_node", values="exact_residual"
    ).reindex(columns=wanted_nodes)
    values = pivot.loc[candidate_order].to_numpy(float)
    observed = topology_single(values)
    frame = pd.DataFrame({"draw_id": np.arange(1, N_DRAWS + 1)})
    summaries = {}
    control_values = pivot.loc[controls].to_numpy(float)
    if not np.isfinite(control_values).all() or not np.isfinite(values).all():
        raise RuntimeError("HPA exact-node residuals are incomplete")
    for model_id, draws in draws_by_model.items():
        null = topology_single_draws(control_values, draws)
        frame[model_id] = null
        summaries[model_id] = empirical_summary(
            null, observed, "upper", expected_n=N_DRAWS
        )
    return frame, {
        "observed_hpa_exact_node_topology": observed,
        "models": summaries,
        "status": "POSTHOC_CROSS_PLATFORM_CORROBORATION_ONLY",
    }


def optional_int(value: Any) -> int | None:
    if value is None or pd.isna(value):
        return None
    return int(value)


def ahba_sample_key_from_metadata(record: Any) -> tuple[int, int | None, int | None, int, int]:
    return (
        int(record.donor_id),
        optional_int(record.well_id),
        optional_int(record.polygon_id),
        int(record.structure_id),
        int(record.top_level_structure_id),
    )


def ahba_sample_key_from_response(sample: dict[str, Any]) -> tuple[int, int | None, int | None, int, int]:
    donor = sample["donor"]
    structure = sample["structure"]
    top = sample["top_level_structure"]
    sample_meta = sample["sample"]
    return (
        int(donor["id"]),
        optional_int(sample_meta.get("well")),
        optional_int(sample_meta.get("polygon")),
        int(structure["id"]),
        int(top["id"]),
    )


def node_residual_tensor(metadata: pd.DataFrame, values: np.ndarray) -> np.ndarray:
    if len(values) != len(metadata) or not np.isfinite(values).all():
        raise RuntimeError("Invalid aligned probe-expression vector")
    left = metadata["hemisphere_class"].eq("left_or_unpaired")
    top1 = metadata["top_level_structure_name"].eq("myelencephalon")
    top2 = metadata["top_level_structure_name"].eq("dorsal thalamus")
    top3 = metadata["top_level_structure_name"].eq("parietal lobe")
    masks = [
        (top1 & metadata["structure_abbreviation"].isin(["Cu", "Gr"]), top1),
        (top2 & metadata["structure_abbreviation"].eq("DTLv"), top2),
        (
            top3
            & metadata["structure_name"].str.lower().str.startswith("postcentral gyrus"),
            top3,
        ),
    ]
    tensor = np.full((len(DONORS), 3), np.nan)
    for donor_index, donor in enumerate(DONORS):
        donor_mask = metadata["donor_id"].eq(donor) & left
        for node, (target, top) in enumerate(masks):
            target_mask = (donor_mask & target).to_numpy(dtype=bool)
            sibling_mask = (donor_mask & top & ~target).to_numpy(dtype=bool)
            if not target_mask.any() or not sibling_mask.any():
                raise RuntimeError(f"Missing target/sibling probe samples: {donor}, node {node}")
            tensor[donor_index, node] = float(
                np.median(values[target_mask]) - np.median(values[sibling_mask])
            )
    if not np.isfinite(tensor).all():
        raise RuntimeError("Incomplete probe residual tensor")
    return tensor


def probe_residuals(
    metadata: pd.DataFrame,
    candidate_order: list[str],
    inventory: list[dict[str, Any]],
) -> tuple[dict[str, np.ndarray], np.ndarray]:
    metadata = metadata.copy().sort_values("sample_index", kind="mergesort").reset_index(drop=True)
    if metadata["sample_index"].astype(int).tolist() != list(range(len(metadata))):
        raise RuntimeError("Frozen AHBA metadata sample_index is not contiguous")
    reference_keys = [
        ahba_sample_key_from_metadata(record)
        for record in metadata.itertuples(index=False)
    ]
    if len(reference_keys) != len(set(reference_keys)):
        raise RuntimeError("Duplicate sample key in frozen AHBA metadata")

    aligned_by_symbol: dict[str, list[np.ndarray]] = {
        symbol: [] for symbol in candidate_order
    }
    for item in inventory:
        symbol = item["symbol"]
        path = V08 / item["relative_path"]
        with gzip.open(path, "rt", encoding="utf-8") as handle:
            payload = json.load(handle)
        msg = payload.get("msg", {})
        probes = msg.get("probes", [])
        samples = msg.get("samples", [])
        if len(probes) != 1:
            raise RuntimeError(f"Expected one probe in {path.name}")
        probe = probes[0]
        if int(probe["id"]) != item["probe_id"] or str(probe["gene-symbol"]) != symbol:
            raise RuntimeError(f"Probe identity mismatch in {path.name}")
        response_keys = [ahba_sample_key_from_response(sample) for sample in samples]
        if len(response_keys) != item["expected_sample_count"]:
            raise RuntimeError(f"Unexpected sample count in {path.name}")
        if len(response_keys) != len(set(response_keys)):
            raise RuntimeError(f"Duplicate AHBA sample key in {path.name}")
        if set(response_keys) != set(reference_keys):
            raise RuntimeError(f"AHBA sample set differs in {path.name}")
        z_response = pd.to_numeric(
            pd.Series(probe["z-score"]), errors="coerce"
        ).to_numpy(dtype=float)
        if len(z_response) != len(response_keys):
            raise RuntimeError(f"Expression/sample length mismatch in {path.name}")
        if response_keys == reference_keys:
            aligned = z_response
        else:
            value_by_key = dict(zip(response_keys, z_response))
            aligned = np.asarray([value_by_key[key] for key in reference_keys], dtype=float)
        if not np.isfinite(aligned).all():
            raise RuntimeError(f"Nonfinite probe z-score in {path.name}")
        aligned_by_symbol[symbol].append(aligned)

    residual_by_symbol: dict[str, np.ndarray] = {}
    median_tensors = []
    for symbol in candidate_order:
        arrays = aligned_by_symbol[symbol]
        if not arrays:
            raise RuntimeError(f"No sealed probe loaded for {symbol}")
        matrix = np.stack(arrays, axis=0)
        residual_by_symbol[symbol] = np.stack(
            [node_residual_tensor(metadata, row) for row in matrix], axis=0
        )
        collapsed_median = np.median(matrix, axis=0)
        median_tensors.append(node_residual_tensor(metadata, collapsed_median))
    return residual_by_symbol, np.stack(median_tensors, axis=0)


def tensor_topology_batch(tensor: np.ndarray) -> np.ndarray:
    n = tensor.shape[0]
    donor_values = np.empty((n, len(DONORS)), dtype=float)
    for donor in range(len(DONORS)):
        ranked = [
            rankdata(tensor[:, :, donor, node], axis=1, method="average")
            for node in range(3)
        ]
        donor_values[:, donor] = (
            corr_rows(ranked[0], ranked[1]) + corr_rows(ranked[1], ranked[2])
        ) / 2 - corr_rows(ranked[0], ranked[2])
    return donor_values.mean(axis=1)


def probe_choice_diagnostic(
    metadata: pd.DataFrame,
    candidate_order: list[str],
    expected_probe_counts: pd.Series,
    inventory: list[dict[str, Any]],
    candidate_tensor: np.ndarray,
) -> tuple[pd.DataFrame, dict[str, Any]]:
    probes, reconstructed_median_tensor = probe_residuals(
        metadata, candidate_order, inventory
    )
    maximum_difference = float(
        np.max(np.abs(reconstructed_median_tensor - candidate_tensor))
    )
    if not np.allclose(
        reconstructed_median_tensor,
        candidate_tensor,
        rtol=0.0,
        atol=1e-12,
        equal_nan=False,
    ):
        raise RuntimeError(
            "Probe-level reconstruction does not reproduce the v0.10 candidate tensor"
        )
    observed_counts = {symbol: int(len(probes[symbol])) for symbol in candidate_order}
    expected_counts = {
        symbol: int(expected_probe_counts.loc[symbol]) for symbol in candidate_order
    }
    if observed_counts != expected_counts:
        raise RuntimeError("Frozen probe counts differ from v0.10")
    rng = np.random.Generator(np.random.PCG64(20260910))
    choices = np.column_stack(
        [rng.integers(0, len(probes[symbol]), size=N_DRAWS) for symbol in candidate_order]
    )
    statistics = np.empty(N_DRAWS)
    chunk = 2_000
    for start in range(0, N_DRAWS, chunk):
        stop = min(start + chunk, N_DRAWS)
        selected = np.stack(
            [
                probes[symbol][choices[start:stop, gene_index]]
                for gene_index, symbol in enumerate(candidate_order)
            ],
            axis=1,
        )
        statistics[start:stop] = tensor_topology_batch(selected)
    if not np.isfinite(statistics).all():
        raise RuntimeError("Nonfinite fixed-denominator probe-choice diagnostic")
    primary_tests = pd.read_csv(INPUTS["v10_primary_tests"])
    frozen_row = primary_tests[
        primary_tests["test_id"].eq("P1_MATCHED_GENESET_SPECIFICITY")
    ]
    if len(frozen_row) != 1 or int(frozen_row.iloc[0]["null_size"]) != N_DRAWS:
        raise RuntimeError("Cannot recover the unique frozen v0.10 P1 q95")
    original_q95 = float(frozen_row.iloc[0]["null_q95"])
    if not np.isfinite(original_q95):
        raise RuntimeError("Frozen v0.10 P1 q95 is nonfinite")
    frame = pd.DataFrame(
        {
            "draw_id": np.arange(1, N_DRAWS + 1),
            "mean_adjacency_contrast": statistics,
        }
    )
    return frame, {
        "draws": N_DRAWS,
        "rng": "numpy.random.Generator(PCG64)",
        "seed": 20260910,
        "median": float(np.median(statistics)),
        "q025": float(np.quantile(statistics, 0.025, method=QUANTILE_METHOD)),
        "q975": float(np.quantile(statistics, 0.975, method=QUANTILE_METHOD)),
        "quantile_method": QUANTILE_METHOD,
        "proportion_above_zero": float(np.mean(statistics > 0)),
        "proportion_above_original_v10_null_q95": float(
            np.mean(statistics > original_q95)
        ),
        "original_v10_null_q95": original_q95,
        "original_v10_comparison": "strictly_greater_than",
        "all_probe_median_tensor_max_abs_difference_vs_v10": maximum_difference,
        "all_probe_median_tensor_reproduced": True,
        "probe_counts": observed_counts,
        "status": "DIAGNOSTIC_ONLY",
    }


def two_way_bootstrap(
    candidate_tensor: np.ndarray,
) -> tuple[pd.DataFrame, dict[str, Any]]:
    if candidate_tensor.shape != (19, len(DONORS), len(NODES)):
        raise RuntimeError("Unexpected tensor shape for two-way bootstrap")
    if not np.isfinite(candidate_tensor).all():
        raise RuntimeError("Nonfinite tensor for two-way bootstrap")
    rng = np.random.Generator(np.random.PCG64(20260911))
    gene_draws = rng.integers(
        0, candidate_tensor.shape[0], size=(N_DRAWS, candidate_tensor.shape[0])
    )
    donor_draws = rng.integers(
        0, candidate_tensor.shape[1], size=(N_DRAWS, candidate_tensor.shape[1])
    )
    statistics = np.empty(N_DRAWS)
    chunk = 2_000
    for start in range(0, N_DRAWS, chunk):
        stop = min(start + chunk, N_DRAWS)
        genes = gene_draws[start:stop]
        donors = donor_draws[start:stop]
        tensor = np.empty((stop - start, candidate_tensor.shape[0], 6, 3))
        for donor_position in range(6):
            donor_index = donors[:, donor_position][:, None]
            for node in range(3):
                tensor[:, :, donor_position, node] = candidate_tensor[
                    genes, donor_index, node
                ]
        statistics[start:stop] = tensor_topology_batch(tensor)
    finite = np.isfinite(statistics)
    kept = statistics[finite]
    if not len(kept):
        raise RuntimeError("All two-way bootstrap draws were nonfinite")
    frame = pd.DataFrame(
        {
            "draw_id": np.arange(1, N_DRAWS + 1),
            "mean_adjacency_contrast": statistics,
            "finite": finite,
        }
    )
    return frame, {
        "attempted_draws": N_DRAWS,
        "finite_draws": int(finite.sum()),
        "nonfinite_draws": int((~finite).sum()),
        "nonfinite_reason": "undefined Spearman correlation after resampling ties/constant vectors",
        "rng": "numpy.random.Generator(PCG64)",
        "seed": 20260911,
        "q025": float(np.quantile(kept, 0.025, method=QUANTILE_METHOD)),
        "median": float(np.median(kept)),
        "q975": float(np.quantile(kept, 0.975, method=QUANTILE_METHOD)),
        "quantile_method": QUANTILE_METHOD,
        "proportion_above_zero": float(np.mean(kept > 0)),
        "status": "DIAGNOSTIC_ONLY",
    }


def deterministic_gzip_csv(frame: pd.DataFrame, path: Path) -> None:
    payload = frame.to_csv(index=False, lineterminator="\n").encode("utf-8")
    with path.open("wb") as raw:
        with gzip.GzipFile(filename="", mode="wb", fileobj=raw, mtime=0) as compressed:
            compressed.write(payload)


def constant_probe_counts(pools: pd.DataFrame) -> pd.Series:
    candidate_groups = pools.groupby("candidate")["exact_ahba_probe_count"]
    if (candidate_groups.nunique() != 1).any():
        raise RuntimeError("Inconsistent candidate probe count")
    candidate = candidate_groups.first().astype(int)
    control = pools.groupby("control")["exact_ahba_probe_count"].agg(
        lambda x: int(pd.Series(x).drop_duplicates().iloc[0])
        if pd.Series(x).nunique() == 1
        else (_ for _ in ()).throw(RuntimeError("Inconsistent control probe count"))
    )
    values = pd.concat([candidate, control])
    if values.index.has_duplicates:
        if (values.groupby(level=0).nunique() > 1).any():
            raise RuntimeError("Conflicting probe count")
        values = values.groupby(level=0).first()
    return values


def write_output_hashes(output_directory: Path) -> None:
    paths = sorted(
        path
        for path in output_directory.iterdir()
        if path.is_file() and path.name != "analysis_hashes_v1_0.txt"
    )
    with (output_directory / "analysis_hashes_v1_0.txt").open(
        "w", encoding="utf-8"
    ) as handle:
        for path in paths:
            handle.write(f"{sha256(path)}  {path.name}\n")


def verification_snapshot(frame: pd.DataFrame) -> pd.DataFrame:
    columns = ["source_id", "path", "bytes", "sha256"]
    if not set(columns).issubset(frame.columns):
        raise RuntimeError("Malformed verification ledger")
    if frame["source_id"].duplicated().any():
        duplicates = sorted(frame.loc[frame["source_id"].duplicated(), "source_id"])
        raise RuntimeError(f"Duplicate source IDs in verification ledger: {duplicates}")
    return frame[columns].sort_values("source_id", kind="mergesort").reset_index(drop=True)


def main() -> int:
    if FINAL_OUT.exists() or STAGING_OUT.exists():
        raise RuntimeError(
            "Refusing to overwrite an existing final or staging analysis directory"
        )
    protocol = json.loads(PROTOCOL_PATH.read_text(encoding="utf-8"))
    amendment1 = json.loads(AMENDMENT1_PATH.read_text(encoding="utf-8"))
    amendment2 = json.loads(AMENDMENT2_PATH.read_text(encoding="utf-8"))
    amendment3 = json.loads(AMENDMENT3_PATH.read_text(encoding="utf-8"))
    verified = verify_inputs(protocol, amendment1, amendment2, amendment3)
    candidate_order = list(protocol["frozen_gene_order"])
    probe_verification, probe_inventory = verify_probe_files(candidate_order)
    verified = pd.concat([verified, probe_verification], ignore_index=True)
    pre_endpoint_snapshot = verification_snapshot(verified)

    panels, controls = load_sealed_panels(candidate_order, amendment3)
    draws_by_model = {
        model_id: panels_to_indices(frame, controls) for model_id, frame in panels.items()
    }

    # No endpoint-bearing file is parsed above this line. Publish only after every
    # analysis succeeds, so a failure cannot masquerade as a completed v1.0 run.
    STAGING_OUT.mkdir(parents=True, exist_ok=False)

    residual_frame = pd.read_csv(INPUTS["v10_ahba_residuals"])
    candidate_tensor = residual_tensor(residual_frame, candidate_order)
    control_tensor = residual_tensor(residual_frame, controls)
    observed_donor, observed = topology_observed(candidate_tensor)
    if not math.isclose(
        observed["mean_adjacency_contrast"], 0.3451754385964913, abs_tol=1e-12
    ):
        raise RuntimeError("Frozen v0.10 observed topology was not reproduced")

    nulls = {
        model_id: topology_null(draws, control_tensor)
        for model_id, draws in draws_by_model.items()
    }
    primary_tests = {}
    for model_id, null in nulls.items():
        result = empirical_summary(
            null["mean_adjacency_contrast"].to_numpy(float),
            observed["mean_adjacency_contrast"],
            "upper",
            expected_n=N_DRAWS,
        )
        result["pass"] = bool(
            result["empirical_p_one_sided"] <= 0.05
            and result["observed_beyond_95pct_threshold"]
        )
        primary_tests[model_id] = result
        deterministic_gzip_csv(
            null, STAGING_OUT / f"ahba_topology_null_{model_id}_v1_0.csv.gz"
        )
    observed_donor.to_csv(
        STAGING_OUT / "observed_topology_donor_v1_0.csv", index=False
    )

    components = topology_component_tests(observed, nulls)
    components.to_csv(
        STAGING_OUT / "topology_component_tests_v1_0.csv", index=False
    )
    lodo = lodo_null_tests(observed_donor, nulls)
    lodo.to_csv(STAGING_OUT / "lodo_null_tests_v1_0.csv", index=False)

    cell = pd.read_csv(INPUTS["v11_cell_contrasts"])
    cell_null, cell_summary = cell_attribution(
        cell,
        candidate_order,
        controls,
        draws_by_model["P1_technical"],
    )
    deterministic_gzip_csv(
        cell_null,
        STAGING_OUT / "cell_attribution_P1_technical_null_v1_0.csv.gz",
    )
    cell_summary["positive_evaluable_for_overall_label"] = bool(
        cell_summary["evaluable"] and cell_summary["all_observed_regions_positive"]
    )

    groups = pd.read_csv(INPUTS["v10_groups"])
    triplets = pd.read_csv(INPUTS["v10_triplets"])
    metadata = pd.read_csv(INPUTS["v08_sample_metadata"])
    geometry, geometry_summary = geometry_analysis(
        groups,
        triplets,
        metadata,
        expected_observed=observed["mean_adjacency_contrast"],
    )
    geometry.to_csv(
        STAGING_OUT / "geometry_matched_triplets_v1_0.csv", index=False
    )
    geometry_primary = geometry_summary["50"]
    geometry_primary["pass"] = bool(
        geometry_primary["empirical_p_one_sided"] <= 0.05
        and geometry_primary["observed_beyond_95pct_threshold"]
    )

    hpa = pd.read_csv(INPUTS["v04_hpa_exact_residuals"])
    hpa_null, hpa_summary = hpa_topology(
        hpa, candidate_order, controls, draws_by_model
    )
    deterministic_gzip_csv(
        hpa_null, STAGING_OUT / "hpa_exact_node_topology_nulls_v1_0.csv.gz"
    )

    pools = pd.read_csv(INPUTS["v10_pools"])
    probe_counts = constant_probe_counts(pools)
    probe_frame, probe_summary = probe_choice_diagnostic(
        metadata,
        candidate_order,
        probe_counts,
        probe_inventory,
        candidate_tensor,
    )
    deterministic_gzip_csv(
        probe_frame, STAGING_OUT / "probe_choice_diagnostic_v1_0.csv.gz"
    )

    bootstrap_frame, bootstrap_summary = two_way_bootstrap(candidate_tensor)
    deterministic_gzip_csv(
        bootstrap_frame,
        STAGING_OUT / "two_way_bootstrap_diagnostic_v1_0.csv.gz",
    )

    p1 = primary_tests["P1_technical"]
    s1 = primary_tests["S1_neuron"]
    if not p1["pass"]:
        overall = "SELECTION_EXPLAINED_OR_UNRESOLVED"
    elif not geometry_primary["pass"]:
        overall = "GENESET_ROBUST_ANATOMY_NOT_GEOMETRY_ROBUST"
    elif not s1["pass"]:
        overall = "SELECTION_ROBUST_BUT_NEURON_GENERIC"
    elif cell_summary["positive_evaluable_for_overall_label"]:
        overall = "ROBUST_COVERED_TOPOLOGY"
    else:
        overall = None

    overall_status = (
        "FROZEN_LABEL_ASSIGNED"
        if overall is not None
        else "NO_FROZEN_LABEL_DEFINED_FOR_NONPOSITIVE_EVALUABLE_S2"
    )

    summary = {
        "version": "v1.0",
        "protocol_sha256": sha256(PROTOCOL_PATH),
        "amendment_A01_sha256": sha256(AMENDMENT1_PATH),
        "amendment_A02_sha256": sha256(AMENDMENT2_PATH),
        "amendment_A03_sha256": sha256(AMENDMENT3_PATH),
        "endpoint_script_sha256": sha256(Path(__file__)),
        "sealed_null_manifest_sha256": sha256(
            NULL_DIR / "null_design_hashes_v1_0.txt"
        ),
        "observed_ahba_topology": observed,
        "P1_conditional_pseudoroute_technical": p1,
        "S1_conditional_pseudoroute_neuron": s1,
        "S2_conditional_cell_attribution": cell_summary,
        "S3_geometry_matched_anatomical_null": {
            "primary_k50": geometry_primary,
            "sensitivities": {
                key: value for key, value in geometry_summary.items() if key != "50"
            },
        },
        "S4_hpa_exact_node_topology": hpa_summary,
        "S5_probe_choice_diagnostic": probe_summary,
        "S6_two_way_bootstrap_diagnostic": bootstrap_summary,
        "S7_topology_component_decomposition_file": "topology_component_tests_v1_0.csv",
        "S8_leave_one_donor_out_file": "lodo_null_tests_v1_0.csv",
        "overall_label": overall,
        "overall_label_status": overall_status,
        "interpretation_status": "POST_V0_11_ROBUSTNESS_EXTENSION_NOT_INDEPENDENT_CONFIRMATION",
        "unconditional_selection_replay_identifiable": False,
        "spatial_surrogate_completed": False,
        "claim_guardrail": "No connectivity, direction, molecular flow, causality, route exclusivity, pain/fibromyalgia mechanism, or stimulation target is inferred.",
    }

    # Detect any concurrent or accidental mutation between precheck and publish.
    verified_after = verify_inputs(protocol, amendment1, amendment2, amendment3)
    probe_verification_after, probe_inventory_after = verify_probe_files(
        candidate_order
    )
    verified_after = pd.concat(
        [verified_after, probe_verification_after], ignore_index=True
    )
    post_endpoint_snapshot = verification_snapshot(verified_after)
    pd.testing.assert_frame_equal(
        pre_endpoint_snapshot,
        post_endpoint_snapshot,
        check_dtype=True,
        check_exact=True,
    )
    if probe_inventory_after != probe_inventory:
        raise RuntimeError("Probe inventory changed during endpoint execution")
    panels_after, controls_after = load_sealed_panels(candidate_order, amendment3)
    if controls_after != controls or any(
        not panels_after[model_id].equals(panels[model_id]) for model_id in panels
    ):
        raise RuntimeError("Sealed null design changed during endpoint execution")
    verified = verified.copy()
    verified["verified_after_endpoint"] = True

    (STAGING_OUT / "summary_v1_0.json").write_text(
        json.dumps(summary, indent=2, sort_keys=True, ensure_ascii=False) + "\n",
        encoding="utf-8",
    )
    verified.to_csv(
        STAGING_OUT / "verified_endpoint_inputs_v1_0.csv", index=False
    )
    (STAGING_OUT / "environment_versions_analysis_v1_0.txt").write_text(
        "\n".join(
            [
                f"python={platform.python_version()}",
                f"python_implementation={platform.python_implementation()}",
                f"numpy={np.__version__}",
                f"pandas={pd.__version__}",
                f"scipy={scipy.__version__}",
                f"platform={platform.platform()}",
                f"byteorder={sys.byteorder}",
                f"quantile_method={QUANTILE_METHOD}",
            ]
        )
        + "\n",
        encoding="utf-8",
    )
    write_output_hashes(STAGING_OUT)
    STAGING_OUT.replace(FINAL_OUT)
    print(json.dumps(summary, indent=2, sort_keys=True, ensure_ascii=False))
    return 0


if __name__ == "__main__":
    raise SystemExit(main())
```
