## Supplementary Archive S1 for "An anatomy-constrained transcriptomic screen identifies a 19-gene panel with an ordered regional similarity pattern in the human somatosensory system": figure_1_observed_donor_structure_v1_0.pdf

### Observed donor-level correlation structure

Post-v0.11 AHBA robustness analysis; not independent confirmation

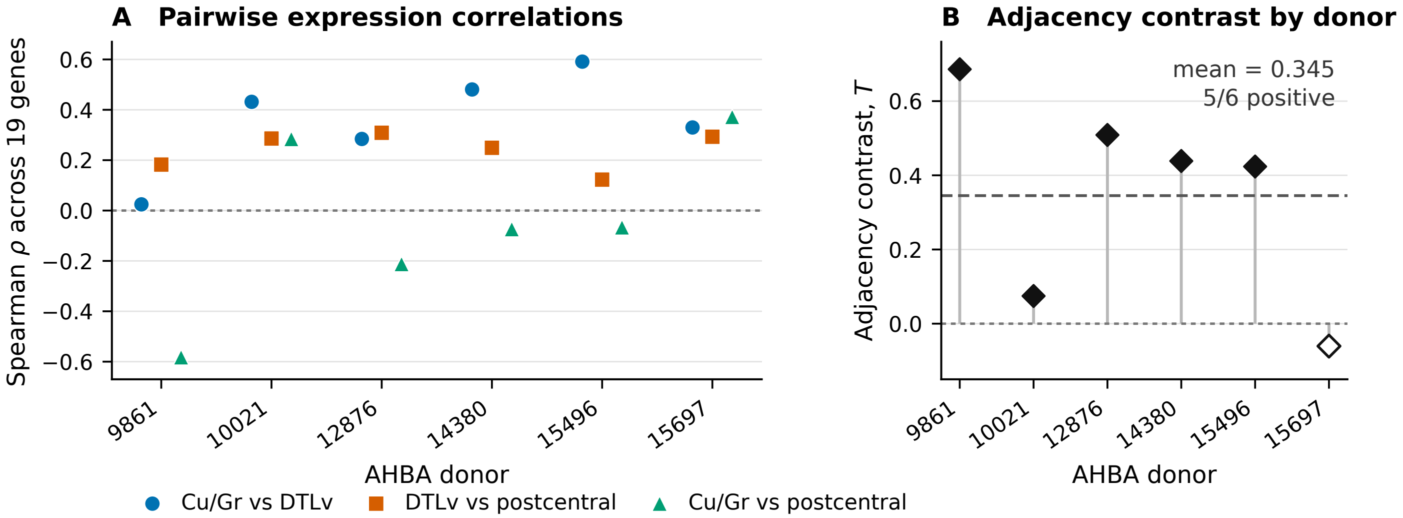
