## Supplementary Archive S1 for "An anatomy-constrained transcriptomic screen identifies a 19-gene panel with an ordered regional similarity pattern in the human somatosensory system": figure_2_conditional_nulls_v1_0.pdf

### Frozen 19-gene topology against conditional pseudoroute nulls

Post-v0.11 AHBA robustness analysis; not independent confirmation

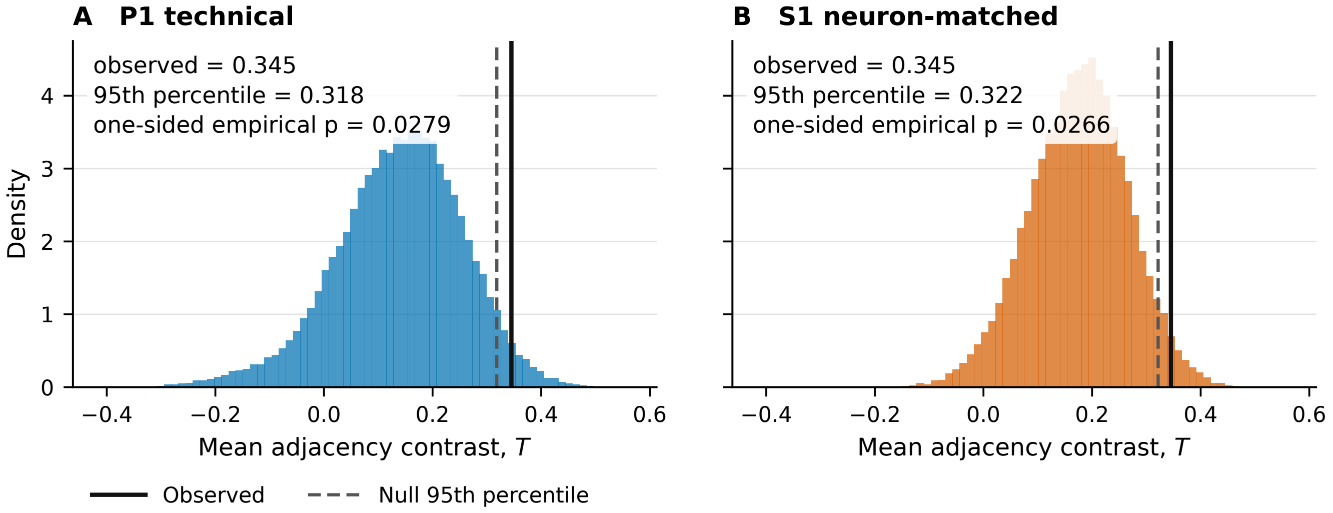
