## Supplementary Archive S1 for "An anatomy-constrained transcriptomic screen identifies a 19-gene panel with an ordered regional similarity pattern in the human somatosensory system": Manuscript_scaffold_v0_1.md

# Selection-aware robustness of a transcriptomic adjacency pattern across human somatosensory relay regions

**Working manuscript — v0.1**  
**Status:** Methods and Results scaffold based on validated v1.0-A04 outputs. Introduction, spatial-control results, independent validation, authorship, affiliations, and final references remain open.

## Abstract

**Background:** Regional transcriptomic organization may reflect shared cellular and anatomical constraints, but spatial autocorrelation, adaptive gene selection, sparse donor coverage, and probe choice can produce apparently pathway-like patterns.  
**Methods:** [Complete after v1.1 spatial protocol is sealed.]  
**Results:** [Complete after v1.1; current results support a selection-aware adjacency pattern but not independent confirmation.]  
**Conclusions:** [Do not finalize before the spatial and validation gates.]

## Introduction

[Draft after the novelty review and v1.1 spatial analysis. The introduction should motivate anatomy-constrained hypothesis generation, explicitly distinguish regional coexpression from connectivity, and avoid fibromyalgia or stimulation as a rationale for the present study.]

## Methods

### Study design and analysis status

We evaluated a fixed 19-gene panel associated with the dorsal column–medial lemniscus (DCML) route in a private 10,262-gene regional atlas: ATP2C1, RAI2, RAB3GAP2, CCT6A, MAP1B, SH3BGRL2, PDHX, C7orf25, TSHZ2, NEFM, ATXN3, ADAM23, HMGCS1, SLC16A6, NANS, IFT57, MEF2A, BBS7, and RAP1GDS1. The panel comprised three `B_ROUTEWIDE_90` and 16 `C_BALANCED` genes selected without disease information. The private source contained regional aggregates rather than donor-level observations.

Version 1.0 was designed after the v0.10–v0.11 findings were known. It was internally time-stamped and hash-sealed before inspection of the new v1.0 endpoints, but was neither externally preregistered nor an independent confirmation. Accordingly, P1 was the primary robustness endpoint within v1.0; S1–S3 were prespecified sensitivity or supportive analyses; S4 was post hoc; and S5–S8 were diagnostic only.

### AHBA residual profiles and topology statistic

Normalized adult-human microarray data were obtained from six Allen Human Brain Atlas donors. For each gene and sample, Allen-provided z scores were collapsed by the median across all exact-symbol HumanMA DNA probes. Analyses used left-hemisphere and unpaired samples. Within each donor and gene, the node residual was the median expression in the target samples minus the median expression in non-target sibling samples belonging to the same top-level anatomical compartment. The targets were the cuneate/gracile nuclei (Cu/Gr; 19 samples), DTLv as the sampled somatosensory-thalamic proxy (20 samples), and postcentral gyrus as the cortical proxy (62 samples). DTLv was not interpreted as an isolated VPL measurement.

For donor $d$, the topology statistic was

$$
T_d=\frac{\rho_{d,\mathrm{Cu/Gr,DTLv}}+\rho_{d,\mathrm{DTLv,Postcentral}}}{2}
-\rho_{d,\mathrm{Cu/Gr,Postcentral}},
$$

where each $\rho$ was a Spearman correlation across the 19 genes. The primary statistic was the arithmetic mean of $T_d$ across the six donors. Observed and null correlations used the identical average-rank/Pearson-on-ranks implementation.

### Conditional pseudoroute nulls

All 8,854 non-DCML four-region combinations from the frozen 23-region private atlas were enumerated, and the original detection, dominance, threshold, and tier-precedence rules were reapplied. All 23 original candidates were excluded from control sampling. A common universe of 742 controls with complete AHBA residuals and broad-region neuronal contrasts was used.

The null conditioned on panel size and the observed tier-by-exact-AHBA-probe-count composition. Of the pseudoroutes, 134 contained enough eligible controls in all six required strata. For each accepted panel, one eligible pseudoroute was sampled and 19 unique genes were drawn within those strata. P1 matched eight technical variables: private-atlas mean expression, regional dispersion and detection; HPA mean expression, dispersion and detection fraction; and AHBA all-brain median expression and robust spread. S1 additionally matched neuronal-minus-non-neuronal contrasts in medulla oblongata, thalamus, and cerebral cortex. Each null contained 50,000 accepted panels.

One-sided Monte Carlo probabilities used $(1+b)/(1+50{,}000)$, where $b$ was the number of null statistics greater than or equal to the observed statistic. Quantiles used the linear method. P1 and S1 passed only if the empirical probability was no greater than 0.05 and the observed statistic strictly exceeded the corresponding null 95th percentile.

This null was conditional rather than a reconstruction of the complete adaptive selection path. Five pseudoroutes reproduced the total tier counts A=0, B=3, C=16, but none also reproduced the exact tier-by-probe table; an unconditional selection-replay probability was therefore not identifiable.

### Additional analyses

S2 reused the P1 panels and defined cell attribution as the minimum, across medulla oblongata, thalamus, and cerebral cortex, of the median neuronal-minus-non-neuronal contrast across genes. Its frozen role in the overall label required only evaluability and positive observed medians in all three regions.

S3 compared the observed triplet with the 50 closest of 239 frozen alternative triplets. Matching used three donor-averaged distances between regional centroids in within-donor standardized MRI coordinates and three log sample-count ratios. Features were robustly standardized across all 240 triplets. Because the nearest-neighbor set was deterministic and asymmetric, $(1+b)/(1+k)$ was treated as a geometry-matched empirical tail/rank score, not an exact randomization probability. Sensitivities used $k=25,75,$ and $100$.

S4 applied the same topology statistic to previously inspected HPA exact-node regional residuals and was post hoc. S5 randomly selected one of 44 exact-symbol AHBA probes per gene in 50,000 draws. S6 used 50,000 two-way bootstrap draws resampling genes and donors independently. S7 decomposed the topology into adjacent mean, donor-wise adjacent bottleneck, endpoint similarity, and the number of donors with both adjacent correlations positive; empirical probabilities were Holm-adjusted across four components within each null model. S8 repeated P1 and S1 after omitting each donor. S5–S8 had no decision role.

### Blinded amendments and numerical-lineage correction

A covariate-only feasibility design initially failed frozen P1 distributional quality control without opening the AHBA topology endpoint. Amendment A02 added a per-panel variance-ratio caliper of 0.25–4.00 while retaining all endpoint definitions, final quality-control thresholds, seeds, and tails.

Endpoint attempt 01 subsequently stopped at the prespecified probe-reconstruction check and produced no final directory or summary; archived partial endpoints were not opened or interpreted. The discrepancy arose because the v0.10 lineage parsed, realigned, and collapsed probe values in `float32`, whereas the first v1.0 consumer reconstructed them in `float64`. The latter differed from the frozen $19\times6\times3$ tensor by at most $2.27\times10^{-7}$ across all 342 cells. Amendment A04 restored the historical `float32` lineage without relaxing the $10^{-12}$ tolerance; the reconstructed tensor then differed by at most $2.22\times10^{-16}$. Genes, samples, probes, panels, statistics, thresholds, tails, quantiles, seeds, and decision rules were unchanged. Final analyses used Python 3.12.13, NumPy 2.3.5, pandas 2.3.3, and SciPy 1.18.1.

### Spatial autocorrelation analysis

[Reserved for v1.1. A cortical spin is not applicable to the mixed brainstem–thalamic–cortical design. The method must be frozen before the corresponding endpoint is read.]

## Results

### Discovery sequence and negative route-wide findings

[Insert the exact v0.3–v0.8 selection flow and the negative route-wide result. This is essential context and must precede the post-discovery topology result.]

### Conditional robustness of the AHBA topology

The observed donor-averaged topology was $T=0.3451754386$ (donor median, 0.4311403509). Five of six donor-specific contrasts were positive, and the prespecified middle node was optimal in five of six donors; the one-sided donor sign test was not significant ($P=0.109375$). Both adjacent correlations were nevertheless positive in all six donors. The adjacent mean was 0.2986842105, the donor-wise bottleneck mean was 0.2099415205, and the mean endpoint similarity was −0.0464912281. [Figure 1]

Both null ensembles passed all structural, balance, variance, distributional, and route-diversity checks. Under P1, 1,395 of 50,000 conditional pseudoroute panels equalled or exceeded the observed statistic, giving $P=0.02791944$; the null mean was 0.13827817 and its 95th percentile was 0.31827485. Under the stricter neuron-conditioned S1 null, 1,328 panels equalled or exceeded the observation ($P=0.02657947$); the null mean was 0.17527322 and its 95th percentile was 0.32163743. P1 and S1 therefore met their frozen dual pass criteria. These probabilities quantify conditional post-discovery robustness and are not independent replication probabilities. [Figure 2]

### Cell attribution and anatomical sensitivity

The observed median neuronal-minus-non-neuronal contrasts were positive in medulla oblongata (0.64067193), thalamus (0.56491715), and cerebral cortex (0.60420511). The weakest-region statistic was 0.56491715, below the P1-null 95th percentile of 0.61072560 ($P=0.09677806$). S2 therefore met only its frozen positive-and-evaluable condition; it did not demonstrate significant conditional neuronal enrichment or route replication.

None of the 50 geometry-nearest regional alternatives reached the observed AHBA topology. The geometry-matched empirical tail/rank score was $1/51=0.01960784$, and the matched-set 95th percentile was 0.23499269. The result met the frozen operational S3 criterion and was unchanged qualitatively for $k=25$, 75, and 100, with zero exceedances and scores of 0.03846154, 0.01315789, and 0.00990099, respectively. Across the full set of 240 triplets, however, the primary triplet ranked third: two geometrically distant Sp5–thalamus–supramarginal combinations produced larger statistics. The result therefore does not establish DCML exclusivity.

Because P1, S1, and S3 met their frozen criteria and S2 was positive and evaluable, the decision algorithm assigned `ROBUST_COVERED_TOPOLOGY`. This is an operational robustness label for AHBA-covered nodes, not confirmation of an end-to-end DCML route or an external biological validation.

### Diagnostic analyses

The post hoc HPA exact-node statistic was 0.78070175, but it was not exceptional under either conditioned null: $P=0.21663567$, 95th percentile 0.91052632 for P1; and $P=0.18523630$, 95th percentile 0.87280702 for S1.

Across 50,000 one-probe-per-gene draws, the median topology was 0.29809942 and the 2.5th–97.5th percentile range was 0.16257310–0.42836257. Every draw was positive, but only 49.038% exceeded the historical v0.10 technical-null 95th percentile of 0.29985380. These are probe-choice distribution percentiles, not confidence intervals or a formal test.

The two-way gene-by-donor bootstrap had median 0.33803645 and a percentile range of −0.04328732 to 0.63134618; 96.114% of draws were positive. Because the genes were selected and only six donors were available, this was diagnostic and the interval does not provide confirmatory coverage.

Component decomposition showed that the signal arose from strong and donor-consistent adjacent-node similarities. Holm-adjusted probabilities for adjacent mean, adjacent bottleneck, and six-of-six positive adjacent pairs were 0.04871903, 0.03767925, and 0.04871903 under P1, and 0.04499910, 0.02839943, and 0.04499910 under S1. Endpoint similarity was not unusually low under either model (Holm-adjusted $P=0.40705186$ and 0.52170957). Thus, the topology should be described as enhanced adjacent-relay residual-profile similarity, not as exceptional endpoint divergence. [Figure 3]

All 12 leave-one-donor-out effects remained positive. Empirical probabilities ranged from 0.00998 to 0.10478 under P1 and from 0.00766 to 0.13402 under S1, so donor-direction stability was strong but statistical calibration was not uniform. These overlapping analyses are not six independent replications.

## Discussion

[Complete only after v1.1 and the independent-atlas feasibility review. Preserve the following boundary: regional expression similarity is not evidence of anatomical connectivity, direction, molecular flow, causality, route exclusivity, a pain/fibromyalgia mechanism, or a stimulation target.]

## Data and code availability

[Pending institutional authorization for the private-source derivatives and a final review of Allen Institute terms and citation requirements. Do not release raw Allen content or private data in a public package until this review is complete.]

## References

1. Markello RD, Arnatkeviciute A, Poline J-B, Fulcher BD, Fornito A, Misic B. Standardizing workflows in imaging transcriptomics with the abagen toolbox. *eLife*. 2021;10:e72129. https://doi.org/10.7554/eLife.72129
2. Burt JB, Helmer M, Shinn M, Anticevic A, Murray JD. Generative modeling of brain maps with spatial autocorrelation. *NeuroImage*. 2020;220:117038. https://doi.org/10.1016/j.neuroimage.2020.117038
3. Wagner HH, Dray S. Generating spatially constrained null models for irregularly spaced data using Moran spectral randomization methods. *Methods in Ecology and Evolution*. 2015;6:1169–1178. https://doi.org/10.1111/2041-210X.12407
