## Supplementary Archive S1 for "An anatomy-constrained transcriptomic screen identifies a 19-gene panel with an ordered regional similarity pattern in the human somatosensory system": README.md

# Route transcriptomics v1.0-A04

This directory contains the frozen post-v0.11 robustness extension for the 19-gene AHBA topology analysis. It is an internal scientific checkpoint, not a public data release and not an external preregistration.

## Scientific scope

The analysis tests whether residual-expression profiles are more similar between adjacent covered somatosensory relay regions (Cu/Gr–DTLv and DTLv–postcentral gyrus) than between Cu/Gr and postcentral gyrus. It does not test anatomical connectivity, direction, molecular flow, causality, route exclusivity, fibromyalgia, or a stimulation target.

## Directory map

- `protocol/`: frozen protocol and amendments A01–A04.
- `null_design_final/`: sealed P1 and S1 conditional-pseudoroute panels and quality-control records.
- `scripts/`: null generation, endpoint analysis, output validation, and figure scripts.
- `analysis/`: the validated final v1.0-A04 outputs.
- `verification/`: reproducibility and provenance records.
- `failed_runs/`: retained prepublication failure evidence; never promote these files as results.
- `verification_runs/`: one valid complete reference copy plus excluded incomplete/orphan execution directories.
- `report/`: Spanish scientific decision report.
- `manuscript/`: English manuscript scaffold.
- `figures/`: manuscript-figure drafts.

## Authoritative outputs

- `analysis/summary_v1_0.json`
- `analysis/analysis_hashes_v1_0.txt`
- `verification/execution_reproducibility_v1_0.json`

The internal frozen decision label is `ROBUST_COVERED_TOPOLOGY`, with interpretation status `POST_V0_11_ROBUSTNESS_EXTENSION_NOT_INDEPENDENT_CONFIRMATION`.

## Validation

From the project root:

```bash
python work/route_v1_0/scripts/verify_analysis_outputs_v1_0.py \
  work/route_v1_0/analysis \
  --reference work/route_v1_0/verification_runs/endpoint_attempt_02_run2
```

The expected terminal status is `VALID_COMPLETE_OUTPUT_SET`, with 13 non-manifest artifacts, 11 row-count checks, and 14 files identical to the preserved complete reference copy.

## Rerun requirements

`scripts/analyze_v1_0.py` verifies hashes before and after endpoint calculation. An exact rerun also requires the recovered upstream v0.4, v0.6, v0.8, v0.10, and v0.11 inputs at the paths encoded in the script. These upstream data are intentionally not duplicated here. The private source and Allen-derived content must not be redistributed until data-governance review is complete.

## Known audit notes

- Endpoint attempt 01 stopped at the frozen S5 tensor check because its consumer used `float64` instead of the historical v0.10 `float32` lineage. A04 restored that lineage without changing any endpoint or decision rule.
- `verification_runs/endpoint_attempt_02_run1` contains an empty bootstrap CSV and is an excluded, incomplete copy.
- The final `analysis/` directory and `verification_runs/endpoint_attempt_02_run2` are complete and byte-identical.
- Recalculation from decimal CSV exports can move a few mathematically tied S7/S8 values across a strict comparison threshold. The sealed summary is authoritative; a later version should retain binary precision or specify a tie tolerance.
