## Supplementary Archive S1 for "An anatomy-constrained transcriptomic screen identifies a 19-gene panel with an ordered regional similarity pattern in the human somatosensory system": Informe_avance_cientifico_v1_0_A04.md

# Informe de avance científico v1.0-A04

**Proyecto:** topología transcriptómica restringida por anatomía en relés somatosensoriales humanos  
**Fecha de cierre:** 8 de septiembre de 2026  
**Estado:** extensión de robustez interna completa; control espacial integral y confirmación independiente pendientes

## Veredicto ejecutivo

El proyecto contiene una señal publicable y ahora dispone de una cadena analítica auditable. El conjunto congelado de 19 genes seleccionado en la ruta DCML mostró, en los seis donantes de AHBA, una coordinación transcriptómica mayor entre los pares de regiones adyacentes Cu/Gr–DTLv y DTLv–giro postcentral que entre los extremos Cu/Gr–giro postcentral. El contraste medio observado fue 0.34518.

La señal superó dos nulos condicionales de 50,000 paneles que preservan el mecanismo de selección, propiedades técnicas y, en el segundo modelo, atribución neuronal regional amplia. También fue mayor que las 50 alternativas anatómicas más cercanas por geometría y cobertura muestral. Sin embargo, estas pruebas se diseñaron después de conocer los resultados v0.10–v0.11 y reutilizan AHBA; por ello establecen **robustez frente a explicaciones alternativas concretas**, no confirmación independiente.

La formulación central defendible es:

> Un panel congelado de 19 genes, seleccionado mediante restricciones anatómicas, presenta en AHBA una coordinación transcriptómica inusualmente alta entre regiones somatosensoriales de relevo adyacentes, comparado con paneles condicionados por selección y con alternativas regionales geométricamente comparables. Este patrón no demuestra conectividad, dirección, flujo molecular, exclusividad de la ruta ni causalidad.

No se debe vincular este resultado, por ahora, con un mecanismo de fibromialgia ni con un blanco de estimulación.

## Qué se analizó

La secuencia científica completa debe conservarse en el artículo:

1. Se catalogaron 16 circuitos y se congelaron cinco rutas.
2. Tres rutas se evaluaron en la base privada agregada.
3. Se obtuvieron 23 candidatos: 19 asignados a DCML, tres a modulación descendente del dolor y uno a ganglios basales.
4. La hipótesis inicial de continuidad transcriptómica global de ruta no se confirmó.
5. En AHBA se identificó posteriormente, para DCML19, una topología de similitud entre los tres relés centrales cubiertos.
6. v1.0 sometió esa observación ya conocida a controles más estrictos de selección, neuronalidad, geometría, sondas y estabilidad.

La unidad analítica primaria de la topología fue el donante AHBA. Para cada donante se calcularon correlaciones de Spearman entre los perfiles de los 19 genes en Cu/Gr, DTLv y giro postcentral. El estadístico congelado fue:

$$
T_d=\frac{\rho_{\mathrm{Cu/Gr,DTLv}}+\rho_{\mathrm{DTLv,Postcentral}}}{2}
-\rho_{\mathrm{Cu/Gr,Postcentral}},
$$

y el resultado primario fue la media de $T_d$ entre seis donantes.

## Resultados v1.0

| Análisis | Resultado exacto | Interpretación permitida |
|---|---:|---|
| Topología observada | $T=0.34518$; mediana por donante 0.43114; 5/6 donantes positivos | Patrón agregado positivo; la prueba de signo por donante no fue significativa ($p=0.10938$) |
| P1: nulo técnico y de selección | $p=0.02792$; q95=0.31827; 1,395/50,000 nulos ≥ observado | La señal excede paneles derivados de pseudorrutas y balanceados en selección y variables técnicas |
| S1: P1 más neuronalidad amplia | $p=0.02658$; q95=0.32164; 1,328/50,000 ≥ observado | La señal no se explica únicamente por atribución neuronal regional amplia |
| S2: atribución celular condicionada | bulbo 0.64067; tálamo 0.56492; corteza 0.60421; $p=0.09678$; q95=0.61073 | Dirección positiva en las tres regiones, pero sin enriquecimiento significativo frente al nulo condicionado |
| S3: alternativas anatómicas | 0/50 alternativas cercanas ≥ observado; score $1/51=0.01961$; q95=0.23499 | Sensibilidad anatómica favorable; es un score de rango determinista, no un valor p de randomización exacta |
| S4: HPA en nodos exactos | observado 0.78070; $p=0.21664$ (P1), $p=0.18524$ (S1) | Concordancia absoluta descriptiva, sin especificidad estadística del panel ni confirmación independiente |
| S5: elección de una sonda por gen | mediana 0.29810; 95% [0.16257, 0.42836]; 100% >0; 49.04% > q95 antiguo | El signo es estable; la excepcionalidad respecto del umbral previo depende de la sonda |
| S6: bootstrap gen × donante | mediana 0.33804; 95% [−0.04329, 0.63135]; 96.11% >0 | Estabilidad direccional alta, con incertidumbre que todavía incluye cero |
| S8: dejar un donante fuera | los seis efectos fueron positivos, 0.27702–0.42632 | Ningún donante determina el signo; la significación no se conserva en todas las omisiones |

El rótulo interno congelado fue `ROBUST_COVERED_TOPOLOGY`. No conviene utilizarlo literalmente como conclusión editorial: representa el cumplimiento de una regla operativa v1.0, no una etiqueta biológica.

## Qué genera el contraste

La descomposición del estadístico cambia de forma importante la narrativa. El resultado no procede de una separación o anticorrelación excepcional entre los extremos. Procede de la semejanza elevada y consistente de los dos pares adyacentes:

- media Cu/Gr–DTLv: $\rho=0.35702$;
- media DTLv–postcentral: $\rho=0.24035$;
- Cu/Gr–postcentral: $\rho=-0.04649$;
- ambos enlaces adyacentes fueron positivos en 6/6 donantes.

Después de corrección de Holm dentro de cada modelo, fueron favorables la media de adyacencias, el enlace adyacente más débil y el recuento 6/6 de enlaces positivos. La similitud baja entre extremos no fue excepcional.

| Componente | P1, $p_{Holm}$ | S1, $p_{Holm}$ | Lectura |
|---|---:|---:|---|
| Media de adyacencias | 0.04872 | 0.04500 | Favorable |
| Cuello de botella adyacente | 0.03768 | 0.02840 | Favorable |
| Ambos pares positivos en 6/6 | 0.04872 | 0.04500 | Favorable |
| Similitud baja de extremos | 0.40705 | 0.52171 | No excepcional |

La expresión apropiada es **“coordinación transcriptómica entre relés adyacentes”**. No son apropiadas “flujo molecular”, “transmisión”, “direccionalidad” ni “conectividad”.

## Especificidad anatómica

La tríada objetivo fue superior a las primeras 100 alternativas más cercanas por geometría y cobertura. En el universo completo de 240 tríadas ocupó el tercer lugar. Dos tríadas lejanas que incluyen Sp5 y giro supramarginal tuvieron contrastes mayores (0.39108 y 0.38099).

Por tanto, el resultado es compatible con una organización somatosensorial central más amplia y no prueba exclusividad de DCML. Las dos tríadas Sp5 deben mostrarse, no ocultarse, porque delimitan con precisión la inferencia.

## Donantes, sondas y plataforma

Cinco de seis donantes tuvieron $T_d>0$. El sexto donante tuvo $T_d=-0.06053$ aun cuando ambos pares adyacentes fueron positivos; en él, la similitud entre extremos fue todavía mayor. Todos los análisis de omisión mantuvieron signo positivo, pero P1 superó el umbral sólo en 4/6 omisiones y S1 en 3/6. Esto demuestra estabilidad cualitativa, no seis replicaciones independientes.

Las 50,000 elecciones aleatorias de una sonda por gen conservaron signo positivo, pero únicamente 49.04% superaron el q95 del nulo v0.10. La mediana y la media multisonde respaldan el patrón, mientras que la significación no es invariante a la sonda. Una validación RNA-seq es prioritaria.

HPA mostró una topología absoluta alta, pero muchos paneles condicionados también la muestran. No debe describirse como replicación positiva. De forma análoga, la atribución neuronal v0.11 pierde significación al condicionar por el proceso de selección en v1.0; queda como apoyo direccional, no como validación celular.

## Trazabilidad e incidente A04

El protocolo v1.0, las enmiendas A01–A03, el diseño nulo y el código de endpoints se sellaron antes de la primera ejecución de resultados v1.0. Esa primera ejecución se detuvo en el control duro S5 porque la reconstrucción empleaba `float64`, mientras que el tensor histórico v0.10 había seguido un linaje `float32`. La diferencia máxima era $2.27\times10^{-7}$.

La enmienda A04 cambió exclusivamente tres conversiones de tipo en la reconstrucción S5. No cambió genes, muestras, sondas, estadísticos, nulos, umbrales, colas, semillas ni reglas de decisión. Con el linaje histórico, el tensor se reprodujo con diferencia máxima $2.22\times10^{-16}$.

Una copia posterior de resultados conservó un archivo bootstrap vacío y fue excluida. El conjunto final contiene 13 artefactos declarados, todos no vacíos y con hashes válidos; sus 11 conteos estructurales son correctos y es idéntico byte por byte a una copia completa preservada. Los intentos fallidos o incompletos permanecen separados para auditoría.

Hay una consideración menor para la futura liberación: al recalcular algunos conteos de cola desde los CSV decimales de S7/S8, empates matemáticos muy próximos pueden cambiar por redondeo en unas pocas observaciones de 50,000. El resumen sellado es autoritativo; v1.1 debe conservar valores binarios o fijar una tolerancia explícita de empate.

## Estructura recomendada del manuscrito

Título provisional:

> *Selection-aware robustness of a transcriptomic adjacency pattern across human somatosensory relay regions*

Orden narrativo:

1. Descubrimiento anatómicamente restringido y resultado negativo de continuidad génica global.
2. Congelación de DCML19 y topología observada por donante.
3. Nulos P1/S1 que reproducen la selección y controlan propiedades técnicas/neuronalidad.
4. Especificidad anatómica emparejada por geometría y descomposición del contraste.
5. Resultados que limitan generalización: atribución celular, HPA, sondas, bootstrap y LODO.
6. Discusión de una organización transcriptómica de adyacencia, sin inferencia de tracto o mecanismo clínico.

En el cuerpo principal deben aparecer los resultados negativos y las sensibilidades que moderan la conclusión. Los archivos completos de 50,000 paneles, las 240 tríadas, los protocolos, hashes e incidente de precisión numérica pueden ir al suplemento.

## Gates pendientes

Antes de aspirar a una revista exigente deben resolverse, en este orden:

1. **Control espacial integral.** La combinación tronco–tálamo–corteza no admite un simple *spin* cortical. Se necesita un nulo para muestras irregulares que preserve autocorrelación espacial o, como mínimo, paneles génicos emparejados explícitamente por autocorrelación de los mapas completos. BrainSMASH genera mapas sustitutos con autocorrelación espacial comparable; la randomización espectral de Moran admite muestras irregulares y randomización conjunta de variables. Véanse [Burt et al., 2020](https://doi.org/10.1016/j.neuroimage.2020.117038) y [Wagner & Dray, 2015](https://doi.org/10.1111/2041-210X.12407).
2. **Coexpresión basal.** Emparejar paneles también por coexpresión global fuera de los nodos objetivo, para separar adyacencia anatómica de una correlación genérica del panel.
3. **Confirmación no inspeccionada.** Buscar una plataforma RNA-seq o single-nucleus con resolución comparable de bulbo/núcleos de columnas dorsales, VPL y corteza somatosensorial. Si no existe cobertura exacta, declarar el límite y diseñar una validación modular, no forzar equivalencias.
4. **Procesamiento AHBA.** Replicar una sensibilidad con una canalización estandarizada y documentada; las decisiones de normalización pueden modificar de forma material los resultados AHBA. Véase [Markello et al., 2021](https://doi.org/10.7554/eLife.72129).
5. **Gobernanza.** Confirmar autorización para publicar derivados de la base privada y separar el paquete académico de cualquier uso comercial. Los términos actuales de Allen permiten investigación no comercial y una cantidad limitada de contenido en publicaciones académicas con cita, pero restringen la redistribución comercial; esto requiere revisión institucional dada la relación con iVenta. Véanse [términos de uso](https://alleninstitute.org/legal/terms-of-use) y [política de citación](https://alleninstitute.org/legal/citation-policy).

## Decisión de avance

**GO para manuscrito exploratorio sólido; GO condicionado para una aspiración alta.** La fortaleza actual es una topología de adyacencia internamente reproducible y resistente a controles de selección razonables. La debilidad principal es la ausencia de un control espacial completo y de una confirmación verdaderamente independiente. La siguiente versión debe resolver primero esos dos puntos; no conviene ampliar la historia hacia fibromialgia o estimulación antes de hacerlo.
