## Supplementary Archive S1 for "An anatomy-constrained transcriptomic screen identifies a 19-gene panel with an ordered regional similarity pattern in the human somatosensory system": figure_captions_v1_0.md

# Draft figure captions — route v1.0-A04

These figures describe post-v0.11 robustness analyses in the same AHBA dataset. They do not constitute independent confirmation and do not establish anatomical connectivity, direction, molecular flow, causality, route exclusivity, a pain or fibromyalgia mechanism, or a stimulation target.

## Figure 1. Observed donor-level correlation structure

(A) Donor-specific Spearman correlations across the frozen 19-gene set between each pair of AHBA regional aggregates: Cu/Gr versus DTLv, DTLv versus postcentral gyrus, and Cu/Gr versus postcentral gyrus. Both adjacent-region correlations were positive in every donor. (B) Donor-specific adjacency contrast, \(T_d=[(\rho_{CuGr,DTLv}+\rho_{DTLv,Postcentral})/2]-\rho_{CuGr,Postcentral}\). Five of six donor values were positive; the mean was \(0.345175\), the median was \(0.431140\), and the donor range was \(-0.060526\) to \(0.685965\). The dashed line marks the six-donor mean. These points are descriptive observations from the same AHBA analysis and are not independent confirmation of the post-selected result.

## Figure 2. Frozen 19-gene topology against conditional pseudoroute nulls

The histograms show 50,000 conditional pseudoroute draws for the mean adjacency-contrast statistic, \(T\), under (A) the P1 technical model and (B) the S1 neuron-matched model. The solid black line marks the observed value (\(T=0.345175\)); the dashed gray line marks the corresponding conditional-null 95th percentile. The observed statistic exceeded the threshold under both models (P1: 1,395 of 50,000 draws at least as large, plus-one empirical upper-tail \(p=0.027919\), null 95th percentile \(=0.318275\); S1: 1,328 of 50,000, \(p=0.026579\), null 95th percentile \(=0.321637\)). These are conditioned, post-selection robustness tests in AHBA, not independent replications.

## Figure 3. Diagnostic component decomposition relative to the conditional nulls

Thick colored segments show the central 5th–95th percentile interval of each 50,000-draw conditional null; open circles show null means and black diamonds show observed values. The mean donor adjacent correlation (observed \(=0.298684\)) and the mean of the donor-wise minimum adjacent correlations (observed \(=0.209942\)) exceeded their upper 95th-percentile thresholds under P1 and S1. Their within-model Holm-adjusted values were, respectively, \(p_{Holm}=0.048719\) and \(0.037679\) under P1 and \(0.044999\) and \(0.028399\) under S1. Both adjacent correlations were positive in all six donors (upper 95th percentile \(=5\) donors under both models; \(p_{Holm}=0.048719\) for P1 and \(0.044999\) for S1). In contrast, mean donor endpoint similarity (observed \(\rho=-0.046491\)) remained within both conditional-null intervals and was not unusually low (lower-tail \(p_{Holm}=0.407052\) for P1 and \(0.521710\) for S1). This frozen diagnostic decomposition shows that the aggregate departure is attributable to the adjacent-correlation components and not to exceptional endpoint dissimilarity. The component tests are marked `DIAGNOSTIC_ONLY` in the endpoint output and do not replace the primary aggregate tests.

## Figure 4. Leave-one-donor-out sensitivity

The mean adjacency-contrast statistic was recomputed after omitting each AHBA donor. Diamonds show the observed leave-one-donor-out (LODO) effect and colored ticks show the model-specific conditional-null 95th percentile. Filled diamonds mark omissions for which the observed effect exceeded that threshold; open diamonds mark effects at or below it. The effect remained positive for all six omissions under both models, but threshold exceedance was not uniform: 4 of 6 omissions under P1 (empirical upper-tail \(p\) range, 0.009980–0.104778) and 3 of 6 under S1 (0.007660–0.134017). Because the six analyses use overlapping five-donor subsets and the same selected gene set, this is a sensitivity diagnostic rather than six independent replications.

## Abbreviations and visual encoding

- **AHBA:** Allen Human Brain Atlas.
- **Cu/Gr:** cuneate and gracile nuclei aggregate.
- **DTLv:** ventral division of the dorsal thalamus aggregate.
- **P1:** technical conditional pseudoroute model.
- **S1:** neuron-matched conditional pseudoroute model.
- **LODO:** leave one donor out.
- Figure 1 colors identify the three regional comparisons as shown in its legend. In Figures 2–4, blue denotes P1 and vermilion denotes S1. These colors were chosen from a color-vision-accessible palette.
