## Supplementary Archive S1 for "An anatomy-constrained transcriptomic screen identifies a 19-gene panel with an ordered regional similarity pattern in the human somatosensory system": generate_conditional_pseudoroute_null_v1_0.py

```
#!/usr/bin/env python3
"""Blind conditional-pseudoroute null generator for frozen DCML19 v1.0.

This program deliberately does NOT read AHBA topology residuals and does NOT
compute the endpoint. It reapplies v0.6 eligibility rules to pseudo-routes and
then conditions on the observed panel size and tier-by-probe composition. It is
not an exact replay of the total number of genes selected by each pseudo-route.

The frozen candidate composition is reproduced jointly by v0.6 tier and AHBA
DNA-probe count.  All original candidate symbols are excluded from controls.
"""

from __future__ import annotations

import gzip
import hashlib
import itertools
import json
import platform
import sys
from collections import Counter
from pathlib import Path
from typing import Any

import numpy as np
import pandas as pd


SEED = 20260908
TARGET_DRAWS = 50_000
MAX_PROPOSALS = 10_000_000
PANEL_MAX_ABS_SMD = 0.25
ENSEMBLE_TARGET_ABS_SMD = 0.10
ENSEMBLE_MAX_ABS_SMD = 0.15
ENSEMBLE_MAX_KS = 0.20
VARIANCE_RATIO_RANGE = (0.50, 2.00)
MIN_CONTRIBUTING_ROUTES = 20
MIN_EFFECTIVE_ROUTES = 15.0
MAX_ROUTE_SHARE = 0.20

PRIMARY_ROUTE = (
    "Dorsal_root_ganglia",
    "Medulla",
    "Thalamus",
    "Parietal_lobe",
)

TECHNICAL_FEATURES = [
    "private_mean_log2",
    "private_sd_log2",
    "private_mean_detection",
    "hpa_mean_log2_ntpm1",
    "hpa_sd_log2_ntpm1",
    "hpa_fraction_ntpm_gt1",
    "ahba_allbrain_median_expression",
    "ahba_allbrain_mad_expression",
]

CELL_REGIONS = ["Medulla oblongata", "Thalamus", "Cerebral cortex"]
CELL_FEATURES = [
    f"neuronal_minus_non_neuronal_median__{region.replace(' ', '_')}"
    for region in CELL_REGIONS
]
BALANCE_FEATURES = TECHNICAL_FEATURES + CELL_FEATURES

HERE = Path(__file__).resolve().parents[1]
PROJECT = HERE.parents[1]
PROTOCOL_PATH = HERE / "protocol" / "protocol_v1_0.json"
AMENDMENT_PATH = HERE / "protocol" / "amendment_01_selection_replay_v1_0.json"
AMENDMENT2_PATH = HERE / "protocol" / "amendment_02_blind_feasibility_v1_0.json"
OUT = HERE / "null_design_final"

V06 = PROJECT / "reconstructed_v006" / "Paquete_reproducible_rutas_transcriptomicas_v0_6"
V10 = PROJECT / "reconstructed_v010" / "Paquete_reproducible_rutas_transcriptomicas_v0_10"
V11 = (
    PROJECT
    / "reconstructed_v011"
    / "Paquete_reproducible_rutas_transcriptomicas_v0_11"
    / "work"
    / "route_v0_11"
)

INPUTS = {
    "v06_private_gene_matrix": V06 / "frozen_inputs" / "gene_expression_private_v0_2.npz",
    "v06_private_gene_qc": V06 / "frozen_inputs" / "gene_collapse_qc.csv",
    "v06_private_selection_metrics": V06 / "analysis" / "routewide_gene_metrics_all_v0_6.csv",
    "v10_pools": V10 / "analysis" / "matched_final_pools_v0_10.csv",
    "v11_cell_contrasts": V11 / "analysis" / "all_candidate_control_cell_contrasts_v0_11.csv.gz",
}


def sha256(path: Path) -> str:
    h = hashlib.sha256()
    with path.open("rb") as fh:
        for chunk in iter(lambda: fh.read(1024 * 1024), b""):
            h.update(chunk)
    return h.hexdigest()


def verify_design_inputs(
    protocol: dict[str, Any], amendment: dict[str, Any], amendment2: dict[str, Any]
) -> pd.DataFrame:
    parent_hash = sha256(PROTOCOL_PATH)
    if parent_hash != amendment["parent_protocol_sha256"]:
        raise RuntimeError(
            f"Parent protocol hash changed: {parent_hash} != "
            f"{amendment['parent_protocol_sha256']}"
        )
    amendment_hash = sha256(AMENDMENT_PATH)
    if parent_hash != amendment2["parent_protocol_sha256"]:
        raise RuntimeError("A02 parent protocol hash changed")
    if amendment_hash != amendment2["parent_amendment_A01_sha256"]:
        raise RuntimeError("A02 parent amendment hash changed")
    failed_design = HERE / "null_design"
    failed_paths = {
        "failed_blind_design_hash_manifest": failed_design
        / "null_design_hashes_v1_0.txt",
        "failed_blind_design_summary": failed_design / "null_design_summary_v1_0.json",
    }
    failed_expected = {
        "failed_blind_design_hash_manifest": amendment2["failed_blind_design_record"][
            "null_design_hash_manifest_sha256"
        ],
        "failed_blind_design_summary": amendment2["failed_blind_design_record"][
            "null_design_summary_sha256"
        ],
    }
    for source_id, path in failed_paths.items():
        if not path.is_file() or sha256(path) != failed_expected[source_id]:
            raise RuntimeError(f"Failed blind-design audit record changed: {path}")
    expected = {
        row["id"]: row
        for row in protocol["input_files"] + amendment["additional_input_files"]
    }
    rows = []
    for source_id, path in INPUTS.items():
        if source_id not in expected:
            raise RuntimeError(f"Input {source_id} is not frozen")
        if not path.is_file():
            raise FileNotFoundError(path)
        observed = sha256(path)
        wanted = expected[source_id]["sha256"]
        if observed != wanted:
            raise RuntimeError(f"Hash mismatch for {source_id}: {observed} != {wanted}")
        rows.append(
            {
                "source_id": source_id,
                "path": str(path.relative_to(PROJECT)),
                "bytes": path.stat().st_size,
                "sha256": observed,
                "verified_before_null_generation": True,
            }
        )
    rows.extend(
        [
            {
                "source_id": "protocol_v1_0",
                "path": str(PROTOCOL_PATH.relative_to(PROJECT)),
                "bytes": PROTOCOL_PATH.stat().st_size,
                "sha256": parent_hash,
                "verified_before_null_generation": True,
            },
            {
                "source_id": "amendment_v1_0_A01",
                "path": str(AMENDMENT_PATH.relative_to(PROJECT)),
                "bytes": AMENDMENT_PATH.stat().st_size,
                "sha256": sha256(AMENDMENT_PATH),
                "verified_before_null_generation": True,
            },
            {
                "source_id": "amendment_v1_0_A02",
                "path": str(AMENDMENT2_PATH.relative_to(PROJECT)),
                "bytes": AMENDMENT2_PATH.stat().st_size,
                "sha256": sha256(AMENDMENT2_PATH),
                "verified_before_null_generation": True,
            },
        ]
    )
    for source_id, path in failed_paths.items():
        rows.append({
            "source_id": source_id,
            "path": str(path.relative_to(PROJECT)),
            "bytes": path.stat().st_size,
            "sha256": sha256(path),
            "verified_before_null_generation": True,
        })
    return pd.DataFrame(rows)


def weighted_ks(candidate: np.ndarray, controls: np.ndarray, weights: np.ndarray) -> float:
    """Two-sample KS distance with normalized weights on the control sample."""
    candidate = np.asarray(candidate, float)
    controls = np.asarray(controls, float)
    weights = np.asarray(weights, float)
    weights = weights / weights.sum()
    grid = np.unique(np.concatenate([candidate, controls]))
    fc = np.searchsorted(np.sort(candidate), grid, side="right") / len(candidate)
    order = np.argsort(controls)
    xs, ws = controls[order], weights[order]
    cum = np.cumsum(ws)
    ix = np.searchsorted(xs, grid, side="right") - 1
    fw = np.where(ix >= 0, cum[np.maximum(ix, 0)], 0.0)
    return float(np.max(np.abs(fc - fw)))


def _constant_side_features(pools: pd.DataFrame, side: str) -> pd.DataFrame:
    columns = [f"{side}_{feature}" for feature in TECHNICAL_FEATURES]
    for column in columns:
        spread = pools.groupby(side)[column].agg(
            lambda x: float(np.nanmax(x) - np.nanmin(x))
        )
        if (spread > 1e-10).any():
            bad = spread[spread > 1e-10].index.tolist()[:10]
            raise RuntimeError(f"Inconsistent {column} for {bad}")
    out = pools[[side, *columns]].drop_duplicates(side).set_index(side)
    out.columns = TECHNICAL_FEATURES
    return out


def _constant_probe_counts(pools: pd.DataFrame, side: str) -> pd.Series:
    counts = pools.groupby(side)["exact_ahba_probe_count"].agg(
        lambda x: int(pd.Series(x).drop_duplicates().iloc[0])
        if pd.Series(x).nunique() == 1
        else (_ for _ in ()).throw(
            RuntimeError(f"Inconsistent exact AHBA probe count for {side}")
        )
    )
    return counts.astype(int)


def load_inputs(protocol: dict[str, Any], amendment: dict[str, Any]):
    npz = np.load(INPUTS["v06_private_gene_matrix"], allow_pickle=False)
    symbols = npz["symbols"].astype(str)
    regions = npz["regions"].astype(str)
    gene_z = npz["gene_z"].astype(float)
    detection = npz["detection_fraction"].astype(float)
    if list(regions) != amendment["private_region_universe"]:
        raise RuntimeError("Private region order differs from the frozen amendment")
    if gene_z.shape != (10262, 23) or detection.shape != gene_z.shape:
        raise RuntimeError(f"Unexpected private matrix shape: {gene_z.shape}")
    if len(set(symbols)) != len(symbols):
        raise RuntimeError("Private symbols are not unique")

    v6_all = pd.read_csv(INPUTS["v06_private_selection_metrics"])
    v6 = (
        v6_all[v6_all["route_id"].eq("DCML_CORE")]
        .drop_duplicates("symbol")
        .set_index("symbol")
    )
    pools = pd.read_csv(INPUTS["v10_pools"])

    cell = pd.read_csv(INPUTS["v11_cell_contrasts"])
    cell = cell[cell["brain_region"].isin(CELL_REGIONS)]
    cell = cell.pivot(
        index="symbol",
        columns="brain_region",
        values="neuronal_minus_non_neuronal_median",
    ).dropna(subset=CELL_REGIONS)

    candidates = list(protocol["frozen_gene_order"])
    if set(candidates) != set(pools["candidate"].astype(str)):
        raise RuntimeError("Pool candidates differ from the frozen 19-gene panel")
    original_candidates = set(
        v6_all.loc[v6_all["private_tier"].ne("NOT_CANDIDATE"), "symbol"].astype(str)
    )
    if len(original_candidates) != 23 or not set(candidates).issubset(original_candidates):
        raise RuntimeError("v0.6 does not recover the 23 frozen original candidates")

    controls = sorted(
        (set(pools["control"].astype(str)) & set(cell.index) & set(v6.index))
        - original_candidates
    )
    if len(candidates) != 19:
        raise RuntimeError(f"Expected 19 frozen genes, found {len(candidates)}")
    if len(controls) != 742:
        raise RuntimeError(f"Expected 742 complete controls, found {len(controls)}")
    if not set(candidates).issubset(cell.index):
        raise RuntimeError("A frozen candidate lacks a central cell contrast")

    return (
        symbols,
        regions,
        gene_z,
        detection,
        v6,
        pools,
        cell,
        candidates,
        controls,
        original_candidates,
    )


def make_feature_table(
    pools: pd.DataFrame,
    cell: pd.DataFrame,
    candidates: list[str],
    controls: list[str],
) -> pd.DataFrame:
    symbols = candidates + controls
    out = pd.DataFrame(index=symbols, columns=BALANCE_FEATURES, dtype=float)
    candidate_values = _constant_side_features(pools, "candidate")
    control_values = _constant_side_features(pools, "control")
    out.loc[candidates, TECHNICAL_FEATURES] = candidate_values.loc[
        candidates, TECHNICAL_FEATURES
    ].to_numpy(float)
    out.loc[controls, TECHNICAL_FEATURES] = control_values.loc[
        controls, TECHNICAL_FEATURES
    ].to_numpy(float)

    for region, feature in zip(CELL_REGIONS, CELL_FEATURES):
        out.loc[symbols, feature] = cell.loc[symbols, region].to_numpy(float)

    if not np.isfinite(out.to_numpy(float)).all():
        raise RuntimeError("Non-finite matching covariates")
    return out


def replay_tiers(
    route_indices: tuple[int, int, int, int],
    gene_z: np.ndarray,
    detection: np.ndarray,
) -> np.ndarray:
    """Literal v0.6 rules, including final precedence A > B > C."""
    route_idx = np.asarray(route_indices, dtype=int)
    off_idx = np.setdiff1d(np.arange(gene_z.shape[1]), route_idx)
    route_z = gene_z[:, route_idx]
    off_z = gene_z[:, off_idx]

    route_floor = route_z.min(axis=1)
    route_mean = route_z.mean(axis=1)
    off_median = np.median(off_z, axis=1)
    off_p90 = np.quantile(off_z, 0.90, axis=1, method="linear")
    off_max = off_z.max(axis=1)
    all_node_detection = detection[:, route_idx].min(axis=1) >= 0.50

    positive_contrasts = np.maximum(route_z - off_median[:, None], 0.0)
    contrast_sum = positive_contrasts.sum(axis=1)
    dominance = np.divide(
        positive_contrasts.max(axis=1),
        contrast_sum,
        out=np.ones_like(contrast_sum),
        where=contrast_sum > 0,
    )

    c_mask = (
        all_node_detection
        & ((route_floor - off_median) >= 0.50)
        & ((route_mean - off_p90) > 0)
        & (dominance <= 0.50)
    )
    b_mask = (
        all_node_detection
        & ((route_floor - off_p90) > 0)
        & (dominance <= 0.50)
    )
    a_mask = (
        all_node_detection
        & ((route_floor - off_max) > 0)
        & (dominance <= 0.50)
    )

    tier = np.full(len(gene_z), "NOT_CANDIDATE", dtype=object)
    tier[c_mask] = "C_BALANCED"
    tier[b_mask] = "B_ROUTEWIDE_90"
    tier[a_mask] = "A_EXCLUSIVE"  # last assignment enforces A > B > C
    return tier


def candidate_strata(
    candidates: list[str], v6: pd.DataFrame, probe_counts: pd.Series
) -> Counter[tuple[str, int]]:
    strata = Counter(
        (str(v6.loc[s, "private_tier"]), int(probe_counts.loc[s]))
        for s in candidates
    )
    expected = Counter(
        {
            ("B_ROUTEWIDE_90", 2): 2,
            ("B_ROUTEWIDE_90", 4): 1,
            ("C_BALANCED", 1): 1,
            ("C_BALANCED", 2): 11,
            ("C_BALANCED", 3): 3,
            ("C_BALANCED", 4): 1,
        }
    )
    if strata != expected:
        raise RuntimeError(f"Frozen tier x probe strata changed: {strata}")
    return strata


def enumerate_pseudoroutes(
    regions: np.ndarray,
    control_gene_z: np.ndarray,
    control_detection: np.ndarray,
    control_probe_counts: np.ndarray,
    strata: Counter[tuple[str, int]],
):
    primary_set = frozenset(PRIMARY_ROUTE)
    eligible = []
    audit_rows = []
    membership_rows = []
    strict_total_count = 0
    strict_strata_count = 0
    for quartet_number, route_idx in enumerate(
        itertools.combinations(range(len(regions)), 4), start=1
    ):
        route_names = tuple(regions[list(route_idx)])
        if frozenset(route_names) == primary_set:
            continue
        route_id = f"Q{quartet_number:05d}"
        tiers = replay_tiers(route_idx, control_gene_z, control_detection)
        stratum_indices = {}
        sufficient = True
        for (tier, probe_count), need in sorted(strata.items()):
            idx = np.flatnonzero((tiers == tier) & (control_probe_counts == probe_count))
            stratum_indices[(tier, probe_count)] = idx
            sufficient &= len(idx) >= need
        tier_counts = Counter(tiers[tiers != "NOT_CANDIDATE"])
        strict_total = bool(
            tier_counts.get("A_EXCLUSIVE", 0) == 0
            and tier_counts.get("B_ROUTEWIDE_90", 0) == 3
            and tier_counts.get("C_BALANCED", 0) == 16
        )
        strict_strata = bool(
            strict_total
            and all(
                len(stratum_indices[key]) == need for key, need in sorted(strata.items())
            )
        )
        strict_total_count += int(strict_total)
        strict_strata_count += int(strict_strata)
        audit_rows.append(
            {
                "quartet_number": quartet_number,
                "route_id": route_id,
                "regions": "|".join(route_names),
                "eligible": bool(sufficient),
                "strict_total_A0_B3_C16": strict_total,
                "strict_total_and_exact_strata": strict_strata,
                "total_A_EXCLUSIVE": int(tier_counts.get("A_EXCLUSIVE", 0)),
                "total_B_ROUTEWIDE_90": int(tier_counts.get("B_ROUTEWIDE_90", 0)),
                "total_C_BALANCED": int(tier_counts.get("C_BALANCED", 0)),
                **{
                    f"available_{tier}_probe{probe_count}": len(stratum_indices[(tier, probe_count)])
                    for tier, probe_count in sorted(strata)
                },
            }
        )
        if sufficient:
            eligible.append(
                {
                    "route_id": route_id,
                    "route_names": route_names,
                    "stratum_indices": stratum_indices,
                }
            )
            for (tier, probe_count), indices in sorted(stratum_indices.items()):
                for control_index in indices:
                    membership_rows.append(
                        {
                            "route_id": route_id,
                            "regions": "|".join(route_names),
                            "tier": tier,
                            "exact_ahba_probe_count": probe_count,
                            "control_index": int(control_index),
                        }
                    )
    strict = {
        "strict_total_A0_B3_C16_pseudoroutes": strict_total_count,
        "strict_total_and_exact_strata_pseudoroutes": strict_strata_count,
    }
    return eligible, pd.DataFrame(audit_rows), pd.DataFrame(membership_rows), strict


def generate_panels(
    eligible_routes,
    controls: list[str],
    strata: Counter[tuple[str, int]],
    features: pd.DataFrame,
    candidates: list[str],
    balance_features: list[str],
    panel_variance_ratio_range: tuple[float, float] | None,
    target_draws: int,
    max_proposals: int,
    seed: int,
):
    rng = np.random.default_rng(seed)
    control_x = features.loc[controls, balance_features].to_numpy(float)
    candidate_x = features.loc[candidates, balance_features].to_numpy(float)
    target_mean = candidate_x.mean(axis=0)
    reference_sd_raw = np.sqrt(
        (candidate_x.var(axis=0, ddof=1) + control_x.var(axis=0, ddof=1)) / 2.0
    )
    zero_reference_sd = reference_sd_raw == 0.0
    reference_sd = np.where(zero_reference_sd, 1.0, reference_sd_raw)
    candidate_variance = candidate_x.var(axis=0, ddof=1)
    nonconstant_candidate = np.ptp(candidate_x, axis=0) > 0.0

    accepted_indices = []
    accepted_routes = []
    accepted_smd = []
    accepted_min_variance_ratio = []
    accepted_max_variance_ratio = []
    proposals = 0
    accepted_count = 0
    batch_size = 25_000
    stratum_items = sorted(strata.items())
    while accepted_count < target_draws and proposals < max_proposals:
        proposed = min(batch_size, max_proposals - proposals)
        route_choice = rng.integers(0, len(eligible_routes), size=proposed)
        selected = np.empty((proposed, 19), dtype=np.int32)
        for route_number in np.unique(route_choice):
            rows = np.flatnonzero(route_choice == route_number)
            route_pools = eligible_routes[int(route_number)]["stratum_indices"]
            offset = 0
            for key, need in stratum_items:
                pool = route_pools[key]
                if need == len(pool):
                    chosen = np.broadcast_to(pool, (len(rows), len(pool)))
                elif need == 1:
                    chosen = pool[rng.integers(0, len(pool), size=(len(rows), 1))]
                else:
                    random_keys = rng.random((len(rows), len(pool)))
                    positions = np.argpartition(random_keys, need - 1, axis=1)[:, :need]
                    chosen = pool[positions]
                selected[rows, offset : offset + need] = chosen
                offset += need
            if offset != 19:
                raise RuntimeError("Frozen strata do not total 19")
        panel_values = control_x[selected]
        means = panel_values.mean(axis=1)
        smd = (means - target_mean) / reference_sd
        if zero_reference_sd.any():
            equal = np.isclose(
                means[:, zero_reference_sd],
                target_mean[zero_reference_sd],
                rtol=1e-12,
                atol=1e-12,
            )
            smd[:, zero_reference_sd] = np.where(equal, 0.0, np.inf)
        acceptance = np.max(np.abs(smd), axis=1) <= PANEL_MAX_ABS_SMD
        if panel_variance_ratio_range is not None:
            if not nonconstant_candidate.any():
                raise RuntimeError("No nonconstant feature is available for the variance caliper")
            panel_variance = panel_values.var(axis=1, ddof=1)
            variance_ratio = (
                panel_variance[:, nonconstant_candidate]
                / candidate_variance[nonconstant_candidate]
            )
            min_variance_ratio = variance_ratio.min(axis=1)
            max_variance_ratio = variance_ratio.max(axis=1)
            acceptance &= (
                (min_variance_ratio >= panel_variance_ratio_range[0])
                & (max_variance_ratio <= panel_variance_ratio_range[1])
            )
        else:
            min_variance_ratio = np.full(proposed, np.nan)
            max_variance_ratio = np.full(proposed, np.nan)
        accepted_in_batch = np.flatnonzero(acceptance)
        remaining = target_draws - accepted_count
        if len(accepted_in_batch) >= remaining and remaining > 0:
            accepted_in_batch = accepted_in_batch[:remaining]
            consumed = int(accepted_in_batch[-1]) + 1
        else:
            consumed = proposed
        if len(accepted_in_batch):
            accepted_indices.append(selected[accepted_in_batch].copy())
            accepted_routes.append(route_choice[accepted_in_batch].copy())
            accepted_smd.append(smd[accepted_in_batch].copy())
            accepted_min_variance_ratio.append(
                min_variance_ratio[accepted_in_batch].copy()
            )
            accepted_max_variance_ratio.append(
                max_variance_ratio[accepted_in_batch].copy()
            )
            accepted_count += len(accepted_in_batch)
        proposals += consumed
        if proposals % 250_000 < batch_size or accepted_count == target_draws:
            print(
                f"seed={seed} proposals={proposals} accepted={accepted_count}",
                flush=True,
            )

    if accepted_indices:
        selected_all = np.concatenate(accepted_indices, axis=0)
        routes_all = np.concatenate(accepted_routes, axis=0)
        smd_all = np.concatenate(accepted_smd, axis=0)
        min_variance_ratio_all = np.concatenate(accepted_min_variance_ratio, axis=0)
        max_variance_ratio_all = np.concatenate(accepted_max_variance_ratio, axis=0)
    else:
        selected_all = np.empty((0, 19), dtype=np.int32)
        routes_all = np.empty(0, dtype=np.int32)
        smd_all = np.empty((0, len(balance_features)), dtype=float)
        min_variance_ratio_all = np.empty(0, dtype=float)
        max_variance_ratio_all = np.empty(0, dtype=float)
    if len(selected_all) and np.any(np.diff(np.sort(selected_all, axis=1), axis=1) == 0):
        raise RuntimeError("A stratified panel contains a duplicate symbol")

    route_ids = [eligible_routes[int(index)]["route_id"] for index in routes_all]
    route_regions = [
        "|".join(eligible_routes[int(index)]["route_names"]) for index in routes_all
    ]
    panels = pd.DataFrame(
        {
            "draw_id": np.arange(1, len(selected_all) + 1),
            "pseudo_route": route_ids,
            "pseudo_route_regions": route_regions,
        }
    )
    control_array = np.asarray(controls, dtype=object)
    for index in range(19):
        panels[f"gene_{index + 1:02d}"] = control_array[selected_all[:, index]]
    balances = pd.DataFrame(
        {
            "draw_id": np.arange(1, len(selected_all) + 1),
            "pseudo_route": route_ids,
            "max_abs_smd": np.max(np.abs(smd_all), axis=1)
            if len(smd_all)
            else np.asarray([], dtype=float),
        }
    )
    if panel_variance_ratio_range is not None:
        balances["min_panel_variance_ratio"] = min_variance_ratio_all
        balances["max_panel_variance_ratio"] = max_variance_ratio_all
    for index, feature in enumerate(balance_features):
        balances[f"smd_{feature}"] = smd_all[:, index]
    return panels, balances, proposals, reference_sd, zero_reference_sd


def ensemble_qc(
    panels: pd.DataFrame,
    features: pd.DataFrame,
    candidates: list[str],
    controls: list[str],
    reference_sd: np.ndarray,
    balance_features: list[str],
):
    gene_columns = [f"gene_{i:02d}" for i in range(1, 20)]
    usage = panels[gene_columns].stack().value_counts().reindex(controls, fill_value=0).astype(float)
    weights = usage.to_numpy(float)
    weights /= weights.sum()
    candidate_x = features.loc[candidates, balance_features].to_numpy(float)
    control_x = features.loc[controls, balance_features].to_numpy(float)
    candidate_mean = candidate_x.mean(axis=0)
    full_control_var = control_x.var(axis=0, ddof=1)
    weighted_mean = np.average(control_x, axis=0, weights=weights)
    weighted_var_numerator = np.average(
        (control_x - weighted_mean) ** 2, axis=0, weights=weights
    )
    weight_correction = 1.0 - float(np.square(weights).sum())
    if weight_correction <= 0:
        raise RuntimeError("Weighted variance is not estimable")
    weighted_var = weighted_var_numerator / weight_correction
    candidate_var = candidate_x.var(axis=0, ddof=1)
    zero_reference_sd = (candidate_var + full_control_var) == 0.0

    rows = []
    for j, feature in enumerate(balance_features):
        smd = (weighted_mean[j] - candidate_mean[j]) / reference_sd[j]
        if zero_reference_sd[j]:
            smd = (
                0.0
                if np.isclose(
                    weighted_mean[j], candidate_mean[j], rtol=1e-12, atol=1e-12
                )
                else np.inf
            )
        candidate_constant = bool(np.ptp(candidate_x[:, j]) == 0.0)
        control_constant = bool(np.ptp(control_x[weights > 0, j]) == 0.0)
        if not candidate_constant:
            variance_ratio = weighted_var[j] / candidate_var[j]
            variance_ok = VARIANCE_RATIO_RANGE[0] <= variance_ratio <= VARIANCE_RATIO_RANGE[1]
        elif control_constant:
            variance_ratio = 1.0
            variance_ok = True
        else:
            variance_ratio = np.inf
            variance_ok = False
        ks = weighted_ks(candidate_x[:, j], control_x[:, j], weights)
        rows.append(
            {
                "feature": feature,
                "candidate_mean": candidate_mean[j],
                "weighted_control_mean": weighted_mean[j],
                "weighted_smd": smd,
                "meets_ideal_abs_smd_le_0_10": bool(abs(smd) <= ENSEMBLE_TARGET_ABS_SMD),
                "variance_ratio": variance_ratio,
                "weighted_ks": ks,
                "mean_balance_pass": abs(smd) <= ENSEMBLE_MAX_ABS_SMD,
                "variance_pass": bool(variance_ok),
                "ks_pass": ks <= ENSEMBLE_MAX_KS,
            }
        )
    balance = pd.DataFrame(rows)

    route_usage = panels["pseudo_route"].value_counts().rename_axis("pseudo_route").reset_index(name="draw_count")
    route_usage["share"] = route_usage["draw_count"] / len(panels)
    effective_routes = float(1.0 / np.square(route_usage["share"]).sum())
    diversity_pass = bool(
        len(route_usage) >= MIN_CONTRIBUTING_ROUTES
        and effective_routes >= MIN_EFFECTIVE_ROUTES
        and route_usage["share"].max() <= MAX_ROUTE_SHARE
    )
    overall = bool(
        balance[["mean_balance_pass", "variance_pass", "ks_pass"]].all().all()
        and diversity_pass
    )
    diversity = {
        "contributing_routes": int(len(route_usage)),
        "effective_routes_inverse_simpson": effective_routes,
        "maximum_route_share": float(route_usage["share"].max()),
        "diversity_pass": diversity_pass,
    }
    return balance, route_usage, diversity, overall


def deterministic_gzip_csv(frame: pd.DataFrame, path: Path) -> None:
    payload = frame.to_csv(index=False, lineterminator="\n").encode("utf-8")
    with path.open("wb") as raw:
        with gzip.GzipFile(filename="", mode="wb", fileobj=raw, mtime=0) as gz:
            gz.write(payload)


def validate_panel_structure(
    panels: pd.DataFrame,
    eligible_routes: list[dict[str, Any]],
    controls: list[str],
    strata: Counter[tuple[str, int]],
) -> bool:
    lookup: dict[tuple[str, str], tuple[str, int]] = {}
    for route in eligible_routes:
        for key, indices in route["stratum_indices"].items():
            for index in indices:
                lookup[(route["route_id"], controls[int(index)])] = key
    expected_slots = [
        key for key, count in sorted(strata.items()) for _ in range(int(count))
    ]
    if len(expected_slots) != 19:
        return False
    for slot, expected in enumerate(expected_slots, start=1):
        gene_column = f"gene_{slot:02d}"
        observed = [
            lookup.get((route, gene))
            for route, gene in zip(panels["pseudo_route"], panels[gene_column])
        ]
        if any(value != expected for value in observed):
            return False
    return bool(
        panels[[f"gene_{i:02d}" for i in range(1, 20)]].nunique(axis=1).eq(19).all()
    )


def write_output_hashes() -> None:
    paths = sorted(
        path
        for path in OUT.iterdir()
        if path.is_file() and path.name != "null_design_hashes_v1_0.txt"
    )
    with (OUT / "null_design_hashes_v1_0.txt").open("w", encoding="utf-8") as handle:
        for path in paths:
            handle.write(f"{sha256(path)}  {path.name}\n")


def main() -> int:
    OUT.mkdir(parents=True, exist_ok=True)
    protocol = json.loads(PROTOCOL_PATH.read_text(encoding="utf-8"))
    amendment = json.loads(AMENDMENT_PATH.read_text(encoding="utf-8"))
    amendment2 = json.loads(AMENDMENT2_PATH.read_text(encoding="utf-8"))
    input_manifest = verify_design_inputs(protocol, amendment, amendment2)

    (
        symbols,
        regions,
        gene_z,
        detection,
        v6,
        pools,
        cell,
        candidates,
        controls,
        original_candidates,
    ) = load_inputs(protocol, amendment)

    symbol_to_row = {symbol: i for i, symbol in enumerate(symbols)}
    if set(candidates + controls) - set(symbol_to_row):
        raise RuntimeError("Candidate/control symbol absent from the frozen private matrix")
    control_rows = np.asarray([symbol_to_row[s] for s in controls], dtype=int)
    features = make_feature_table(pools, cell, candidates, controls)
    probe_counts_all = pd.concat(
        [
            _constant_probe_counts(pools, "candidate"),
            _constant_probe_counts(pools, "control"),
        ]
    )
    if probe_counts_all.index.has_duplicates:
        duplicate_counts = probe_counts_all.groupby(level=0).nunique()
        if (duplicate_counts > 1).any():
            raise RuntimeError("A symbol has conflicting probe counts across sides")
        probe_counts_all = probe_counts_all.groupby(level=0).first()
    strata = candidate_strata(candidates, v6, probe_counts_all)
    control_probe_counts = probe_counts_all.loc[controls].astype(int).to_numpy()

    eligible, route_audit, membership, strict_support = enumerate_pseudoroutes(
        regions,
        gene_z[control_rows],
        detection[control_rows],
        control_probe_counts,
        strata,
    )
    if len(route_audit) != 8854:
        raise RuntimeError(f"Expected 8,854 nonprimary quartets, found {len(route_audit)}")
    if len(eligible) != 134:
        raise RuntimeError(f"Expected 134 eligible pseudoroutes, found {len(eligible)}")
    if strict_support != {
        "strict_total_A0_B3_C16_pseudoroutes": 5,
        "strict_total_and_exact_strata_pseudoroutes": 0,
    }:
        raise RuntimeError(f"Strict replay support changed: {strict_support}")
    control_array = np.asarray(controls, dtype=object)
    membership["control"] = control_array[membership["control_index"].to_numpy(int)]

    route_audit.to_csv(OUT / "pseudoroute_audit_v1_0.csv", index=False)
    deterministic_gzip_csv(
        membership,
        OUT / "eligible_pseudoroute_stratum_membership_v1_0.csv.gz",
    )
    input_manifest.to_csv(OUT / "verified_null_design_inputs_v1_0.csv", index=False)

    model_specs = {
        "P1_technical": {
            "seed": 20260908,
            "features": TECHNICAL_FEATURES,
            "panel_variance_ratio_range": (0.25, 4.00),
        },
        "S1_neuron": {
            "seed": 20260909,
            "features": BALANCE_FEATURES,
            "panel_variance_ratio_range": None,
        },
    }
    model_summaries: dict[str, Any] = {}
    all_ready = True
    for model_id, spec in model_specs.items():
        model_features = list(spec["features"])
        panels, panel_balance, proposals, reference_sd, zero_reference_sd = generate_panels(
            eligible,
            controls,
            strata,
            features,
            candidates,
            model_features,
            spec["panel_variance_ratio_range"],
            TARGET_DRAWS,
            MAX_PROPOSALS,
            int(spec["seed"]),
        )
        enough = len(panels) == TARGET_DRAWS
        structural = enough and validate_panel_structure(
            panels, eligible, controls, strata
        )
        if enough:
            balance, route_usage, diversity, ensemble_pass = ensemble_qc(
                panels,
                features,
                candidates,
                controls,
                reference_sd,
                model_features,
            )
        else:
            balance = pd.DataFrame()
            route_usage = pd.DataFrame()
            diversity = {}
            ensemble_pass = False
        qc_pass = bool(enough and structural and ensemble_pass)
        status = "READY_FOR_ENDPOINT" if qc_pass else "TECHNICALLY_INCONCLUSIVE"
        all_ready &= qc_pass

        deterministic_gzip_csv(
            panels, OUT / f"conditional_pseudoroute_panels_{model_id}_v1_0.csv.gz"
        )
        deterministic_gzip_csv(
            panel_balance,
            OUT / f"conditional_pseudoroute_panel_balance_{model_id}_v1_0.csv.gz",
        )
        if not balance.empty:
            balance.to_csv(
                OUT / f"conditional_pseudoroute_ensemble_balance_{model_id}_v1_0.csv",
                index=False,
            )
            route_usage.to_csv(
                OUT / f"conditional_pseudoroute_route_usage_{model_id}_v1_0.csv",
                index=False,
            )
        reference_frame = pd.DataFrame(
            {
                "feature": model_features,
                "fixed_reference_sd_used": reference_sd,
                "raw_reference_sd_was_zero": zero_reference_sd,
            }
        )
        reference_frame.to_csv(
            OUT / f"conditional_pseudoroute_reference_sd_{model_id}_v1_0.csv",
            index=False,
        )
        historical_identity: dict[str, bool] | None = None
        if model_id == "S1_neuron":
            historical_identity = {}
            historical_dir = HERE / "null_design"
            filenames = [
                f"conditional_pseudoroute_panels_{model_id}_v1_0.csv.gz",
                f"conditional_pseudoroute_panel_balance_{model_id}_v1_0.csv.gz",
                f"conditional_pseudoroute_ensemble_balance_{model_id}_v1_0.csv",
                f"conditional_pseudoroute_route_usage_{model_id}_v1_0.csv",
                f"conditional_pseudoroute_reference_sd_{model_id}_v1_0.csv",
            ]
            for filename in filenames:
                historical_path = historical_dir / filename
                final_path = OUT / filename
                same = bool(
                    historical_path.is_file()
                    and final_path.is_file()
                    and sha256(historical_path) == sha256(final_path)
                )
                historical_identity[filename] = same
            if not all(historical_identity.values()):
                raise RuntimeError(
                    f"S1 changed after A02 despite unchanged rules: {historical_identity}"
                )
        model_summaries[model_id] = {
            "status": status,
            "endpoint_read_or_computed": False,
            "seed": int(spec["seed"]),
            "balance_features": model_features,
            "accepted_panels": int(len(panels)),
            "proposals_consumed": int(proposals),
            "acceptance_fraction": float(len(panels) / proposals) if proposals else 0.0,
            "panel_max_abs_smd_threshold": PANEL_MAX_ABS_SMD,
            "panel_variance_ratio_caliper": spec["panel_variance_ratio_range"],
            "panel_max_abs_smd_observed": float(panel_balance["max_abs_smd"].max())
            if len(panel_balance)
            else None,
            "minimum_panel_variance_ratio_observed": float(
                panel_balance["min_panel_variance_ratio"].min()
            )
            if spec["panel_variance_ratio_range"] is not None and len(panel_balance)
            else None,
            "maximum_panel_variance_ratio_observed": float(
                panel_balance["max_panel_variance_ratio"].max()
            )
            if spec["panel_variance_ratio_range"] is not None and len(panel_balance)
            else None,
            "ensemble_qc_pass": bool(ensemble_pass),
            "structural_qc_pass": bool(structural),
            "route_diversity": diversity,
            "identical_to_failed_design_where_A02_requires": historical_identity,
            "maximum_marginal_abs_smd": float(balance["weighted_smd"].abs().max())
            if not balance.empty
            else None,
            "all_marginal_abs_smd_at_ideal_0_10": bool(
                balance["meets_ideal_abs_smd_le_0_10"].all()
            )
            if not balance.empty
            else False,
            "maximum_weighted_ks": float(balance["weighted_ks"].max())
            if not balance.empty
            else None,
            "minimum_variance_ratio": float(balance["variance_ratio"].min())
            if not balance.empty
            else None,
            "maximum_variance_ratio": float(balance["variance_ratio"].max())
            if not balance.empty
            else None,
        }

    design = {
        "protocol_id": "CONDITIONAL_TIER_MATCHED_PSEUDOROUTE_NULL_v1.0",
        "endpoint_read_or_computed": False,
        "parent_protocol_sha256": sha256(PROTOCOL_PATH),
        "amendment_A01_sha256": sha256(AMENDMENT_PATH),
        "amendment_A02_sha256": sha256(AMENDMENT2_PATH),
        "failed_blind_design_hash_manifest_sha256": amendment2[
            "failed_blind_design_record"
        ]["null_design_hash_manifest_sha256"],
        "generator_script_sha256": sha256(Path(__file__)),
        "target_draws_per_model": TARGET_DRAWS,
        "maximum_proposals_per_model": MAX_PROPOSALS,
        "panel_max_abs_smd": PANEL_MAX_ABS_SMD,
        "ensemble_target_abs_smd": ENSEMBLE_TARGET_ABS_SMD,
        "ensemble_hard_max_abs_smd": ENSEMBLE_MAX_ABS_SMD,
        "ensemble_max_weighted_ks": ENSEMBLE_MAX_KS,
        "variance_ratio_range": VARIANCE_RATIO_RANGE,
        "candidate_strata": {
            f"{key[0]}|probe{key[1]}": value for key, value in sorted(strata.items())
        },
        "candidate_symbols": candidates,
        "excluded_original_candidates": sorted(original_candidates),
        "complete_control_symbols": controls,
        "complete_control_count": len(controls),
        "eligible_pseudoroute_count": len(eligible),
        "strict_replay_support": strict_support,
        "conditioning_statement": "Pseudoroutes may contain A genes or surplus B/C genes. The null conditions on panel size 19 and the observed tier-by-probe composition; it is not an unconditional replay of total candidate count.",
        "fixed_smd_denominator": "sqrt((candidate sample variance + full 742-control sample variance)/2), fixed before panel generation",
        "models": model_summaries,
    }
    (OUT / "null_design_summary_v1_0.json").write_text(
        json.dumps(design, indent=2, sort_keys=True, ensure_ascii=False) + "\n",
        encoding="utf-8",
    )
    (OUT / "environment_versions_null_design_v1_0.txt").write_text(
        "\n".join(
            [
                f"python={platform.python_version()}",
                f"python_implementation={platform.python_implementation()}",
                f"numpy={np.__version__}",
                f"pandas={pd.__version__}",
                f"platform={platform.platform()}",
                f"byteorder={sys.byteorder}",
            ]
        )
        + "\n",
        encoding="utf-8",
    )
    write_output_hashes()
    print(json.dumps(design, indent=2, sort_keys=True, ensure_ascii=False))
    return 0 if all_ready else 2


if __name__ == "__main__":
    raise SystemExit(main())
```
