## Supplementary Archive S1 for "An anatomy-constrained transcriptomic screen identifies a 19-gene panel with an ordered regional similarity pattern in the human somatosensory system": plot_v1_0_results.py

```
#!/usr/bin/env python3
"""Create deterministic draft figures for the sealed v1.0-A04 results.

The script is intentionally read-only with respect to the endpoint analysis and
protocol directories.  It writes figures and a provenance manifest only to the
requested output directory.
"""

from __future__ import annotations

import argparse
import hashlib
import json
import os
import platform
from pathlib import Path
import tempfile
import xml.etree.ElementTree as ET

import matplotlib as mpl
mpl.use("Agg")
import matplotlib.pyplot as plt
from matplotlib import font_manager
from matplotlib.lines import Line2D
import numpy as np
import pandas as pd
from PIL import Image, __version__ as pillow_version


ANALYSIS_FILES = (
    "summary_v1_0.json",
    "ahba_topology_null_P1_technical_v1_0.csv.gz",
    "ahba_topology_null_S1_neuron_v1_0.csv.gz",
    "observed_topology_donor_v1_0.csv",
    "topology_component_tests_v1_0.csv",
    "lodo_null_tests_v1_0.csv",
)

MODEL_CONFIG = {
    "P1_technical": {
        "short": "P1 technical",
        "color": "#0072B2",
        "summary_key": "P1_conditional_pseudoroute_technical",
        "null_file": "ahba_topology_null_P1_technical_v1_0.csv.gz",
    },
    "S1_neuron": {
        "short": "S1 neuron-matched",
        "color": "#D55E00",
        "summary_key": "S1_conditional_pseudoroute_neuron",
        "null_file": "ahba_topology_null_S1_neuron_v1_0.csv.gz",
    },
}

COMPONENT_CONFIG = (
    ("adjacent_mean", "Adjacent mean", r"Mean donor adjacent $\rho$"),
    (
        "adjacent_bottleneck",
        "Adjacent bottleneck",
        r"Mean donor-wise minimum adjacent $\rho$",
    ),
    ("endpoint_similarity", "Endpoint similarity", r"Mean donor endpoint $\rho$"),
    (
        "both_adjacent_positive_donor_count",
        "Both adjacent positive",
        r"Donors with both adjacent $\rho>0$",
    ),
)


def sha256(path: Path) -> str:
    digest = hashlib.sha256()
    with path.open("rb") as handle:
        for block in iter(lambda: handle.read(1024 * 1024), b""):
            digest.update(block)
    return digest.hexdigest()


def atomic_write_bytes(destination: Path, payload: bytes) -> None:
    descriptor, temporary_name = tempfile.mkstemp(
        prefix=f".{destination.stem}.", suffix=destination.suffix, dir=destination.parent
    )
    temporary_path = Path(temporary_name)
    try:
        with os.fdopen(descriptor, "wb") as handle:
            handle.write(payload)
            handle.flush()
            os.fsync(handle.fileno())
        os.replace(temporary_path, destination)
    finally:
        temporary_path.unlink(missing_ok=True)


def verify_figure_file(path: Path) -> None:
    if not path.is_file() or path.stat().st_size == 0:
        raise RuntimeError(f"Missing or empty figure output: {path}")
    if path.suffix == ".png":
        with Image.open(path) as image:
            image.verify()
    elif path.suffix == ".svg":
        ET.parse(path)
    elif path.suffix == ".pdf":
        payload = path.read_bytes()
        if not payload.startswith(b"%PDF-") or b"%%EOF" not in payload[-1024:]:
            raise RuntimeError(f"Invalid PDF envelope: {path}")
    else:
        raise ValueError(f"Unsupported figure extension: {path.suffix}")


def configure_matplotlib() -> None:
    mpl.rcParams.update(
        {
            "font.family": "DejaVu Sans",
            "font.size": 8.5,
            "axes.titlesize": 9.5,
            "axes.labelsize": 9,
            "axes.linewidth": 0.75,
            "axes.spines.top": False,
            "axes.spines.right": False,
            "xtick.labelsize": 8,
            "ytick.labelsize": 8,
            "xtick.major.width": 0.7,
            "ytick.major.width": 0.7,
            "legend.fontsize": 8,
            "figure.dpi": 120,
            "savefig.dpi": 600,
            "savefig.facecolor": "white",
            "svg.hashsalt": "route-v1.0-A04",
            "pdf.fonttype": 42,
            "ps.fonttype": 42,
        }
    )


def add_status_note(fig: mpl.figure.Figure) -> None:
    fig.text(
        0.99,
        0.965,
        "Post-v0.11 AHBA robustness analysis; not independent confirmation",
        ha="right",
        va="top",
        fontsize=6.5,
        color="#555555",
    )


def save_figure(fig: mpl.figure.Figure, base: Path) -> list[Path]:
    outputs: list[Path] = []
    metadata_by_suffix = {
        ".png": {"Software": "matplotlib; route v1.0-A04"},
        ".svg": {"Creator": "matplotlib; route v1.0-A04", "Date": None},
        ".pdf": {
            "Creator": "matplotlib; route v1.0-A04",
            "CreationDate": None,
            "ModDate": None,
        },
    }
    for suffix in (".png", ".svg", ".pdf"):
        destination = base.with_suffix(suffix)
        descriptor, temporary_name = tempfile.mkstemp(
            prefix=f".{base.name}.", suffix=suffix, dir=base.parent
        )
        os.close(descriptor)
        temporary_path = Path(temporary_name)
        try:
            fig.savefig(
                temporary_path,
                format=suffix.removeprefix("."),
                bbox_inches="tight",
                pad_inches=0.035,
                metadata=metadata_by_suffix[suffix],
            )
            if temporary_path.stat().st_size == 0:
                raise RuntimeError(f"Empty temporary figure output: {temporary_path}")
            os.replace(temporary_path, destination)
        finally:
            temporary_path.unlink(missing_ok=True)
        outputs.append(destination)
    plt.close(fig)
    return outputs


def verify_inputs(analysis_dir: Path) -> dict[str, Path]:
    paths = {name: analysis_dir / name for name in ANALYSIS_FILES}
    missing = [str(path) for path in paths.values() if not path.is_file()]
    if missing:
        raise FileNotFoundError("Missing required analysis files: " + ", ".join(missing))
    return paths


def load_results(
    analysis_dir: Path,
) -> tuple[dict, dict[str, pd.DataFrame], pd.DataFrame, pd.DataFrame, pd.DataFrame]:
    with (analysis_dir / "summary_v1_0.json").open("r", encoding="utf-8") as handle:
        summary = json.load(handle)
    if summary.get("execution_revision") != "v1.0-A04_FLOAT32_LINEAGE_FIX":
        raise ValueError("Unexpected endpoint revision")
    if summary.get("interpretation_status") != "POST_V0_11_ROBUSTNESS_EXTENSION_NOT_INDEPENDENT_CONFIRMATION":
        raise ValueError("Unexpected interpretation status")

    nulls: dict[str, pd.DataFrame] = {}
    for model, config in MODEL_CONFIG.items():
        frame = pd.read_csv(analysis_dir / config["null_file"])
        if len(frame) != 50_000:
            raise ValueError(f"{model}: expected 50,000 null draws; observed {len(frame):,}")
        nulls[model] = frame

    components = pd.read_csv(analysis_dir / "topology_component_tests_v1_0.csv")
    lodo = pd.read_csv(analysis_dir / "lodo_null_tests_v1_0.csv")
    donors = pd.read_csv(analysis_dir / "observed_topology_donor_v1_0.csv")
    if len(components) != 8 or len(lodo) != 12 or len(donors) != 6:
        raise ValueError("Unexpected component, LODO, or donor row count")

    required_donor_columns = {
        "donor_id",
        "rho_CuGr_DTLv",
        "rho_DTLv_Postcentral",
        "rho_CuGr_Postcentral",
        "adjacency_contrast",
        "both_adjacent_positive",
    }
    if not required_donor_columns.issubset(donors.columns) or donors["donor_id"].duplicated().any():
        raise ValueError("Invalid observed donor table")
    recomputed_contrast = (
        (donors["rho_CuGr_DTLv"] + donors["rho_DTLv_Postcentral"]) / 2
        - donors["rho_CuGr_Postcentral"]
    )
    if not np.allclose(
        recomputed_contrast.to_numpy(float),
        donors["adjacency_contrast"].to_numpy(float),
        atol=1e-12,
        rtol=0,
    ):
        raise ValueError("Donor adjacency contrasts do not reproduce from pairwise correlations")
    observed_summary = summary["observed_ahba_topology"]
    donor_contrasts = donors["adjacency_contrast"].to_numpy(float)
    if not np.isclose(
        donor_contrasts.mean(),
        float(observed_summary["mean_adjacency_contrast"]),
        atol=1e-12,
        rtol=0,
    ):
        raise ValueError("Donor mean contrast does not reproduce the endpoint summary")
    if not np.isclose(
        np.median(donor_contrasts),
        float(observed_summary["median_adjacency_contrast"]),
        atol=1e-12,
        rtol=0,
    ):
        raise ValueError("Donor median contrast does not reproduce the endpoint summary")
    computed_both_positive = (
        (donors["rho_CuGr_DTLv"] > 0) & (donors["rho_DTLv_Postcentral"] > 0)
    )
    if not np.array_equal(
        computed_both_positive.to_numpy(bool),
        donors["both_adjacent_positive"].astype(bool).to_numpy(),
    ):
        raise ValueError("Donor adjacent-positive indicators do not reproduce")
    if int(computed_both_positive.sum()) != int(
        observed_summary["both_adjacent_positive_donor_count"]
    ):
        raise ValueError("Adjacent-positive donor count does not match the endpoint summary")
    return summary, nulls, components, lodo, donors


def figure_1_observed_donor_structure(
    output_dir: Path,
    summary: dict,
    donors: pd.DataFrame,
) -> list[Path]:
    donor_labels = donors["donor_id"].astype(str).tolist()
    x = np.arange(len(donor_labels), dtype=float)
    fig, axes = plt.subplots(
        1,
        2,
        figsize=(7.15, 3.25),
        gridspec_kw={"width_ratios": [1.6, 1.0]},
    )

    pairwise_series = (
        ("rho_CuGr_DTLv", "Cu/Gr vs DTLv", "#0072B2", "o", -0.18),
        ("rho_DTLv_Postcentral", "DTLv vs postcentral", "#D55E00", "s", 0.0),
        ("rho_CuGr_Postcentral", "Cu/Gr vs postcentral", "#009E73", "^", 0.18),
    )
    for column, label, color, marker, offset in pairwise_series:
        axes[0].scatter(
            x + offset,
            donors[column].to_numpy(float),
            s=34,
            marker=marker,
            facecolor=color,
            edgecolor="white",
            linewidth=0.45,
            label=label,
            zorder=3,
        )
    axes[0].axhline(0, color="#777777", linestyle=(0, (2, 2)), linewidth=0.9)
    axes[0].set_ylim(-0.67, 0.67)
    axes[0].set_title("A   Pairwise expression correlations", loc="left", fontweight="bold")
    axes[0].set_ylabel(r"Spearman $\rho$ across 19 genes")
    axes[0].legend(
        frameon=False,
        loc="upper left",
        bbox_to_anchor=(0.0, -0.28),
        ncol=3,
        columnspacing=1.0,
        handletextpad=0.35,
    )

    contrasts = donors["adjacency_contrast"].to_numpy(float)
    positive = contrasts > 0
    axes[1].vlines(x, 0, contrasts, color="#B8B8B8", linewidth=1.1, zorder=1)
    axes[1].scatter(
        x[positive],
        contrasts[positive],
        s=35,
        marker="D",
        facecolor="#111111",
        edgecolor="#111111",
        linewidth=0.7,
        zorder=3,
    )
    axes[1].scatter(
        x[~positive],
        contrasts[~positive],
        s=35,
        marker="D",
        facecolor="white",
        edgecolor="#111111",
        linewidth=1.0,
        zorder=3,
    )
    mean_contrast = float(summary["observed_ahba_topology"]["mean_adjacency_contrast"])
    axes[1].axhline(mean_contrast, color="#555555", linestyle=(0, (4, 2)), linewidth=1.05)
    axes[1].axhline(0, color="#777777", linestyle=(0, (2, 2)), linewidth=0.9)
    axes[1].text(
        0.97,
        0.95,
        f"mean = {mean_contrast:.3f}\n{int(positive.sum())}/6 positive",
        transform=axes[1].transAxes,
        ha="right",
        va="top",
        linespacing=1.3,
        color="#333333",
    )
    axes[1].set_ylim(-0.15, 0.76)
    axes[1].set_title("B   Adjacency contrast by donor", loc="left", fontweight="bold")
    axes[1].set_ylabel(r"Adjacency contrast, $T$")

    for axis in axes:
        axis.set_xticks(x, donor_labels, rotation=35, ha="right")
        axis.set_xlabel("AHBA donor")
        axis.grid(axis="y", color="#D8D8D8", linewidth=0.55, alpha=0.7)
        axis.set_axisbelow(True)

    fig.suptitle(
        "Observed donor-level correlation structure",
        x=0.07,
        y=1.015,
        ha="left",
        fontsize=11,
        fontweight="bold",
    )
    add_status_note(fig)
    fig.subplots_adjust(left=0.095, right=0.995, top=0.85, bottom=0.31, wspace=0.34)
    return save_figure(fig, output_dir / "figure_1_observed_donor_structure_v1_0")


def figure_2_null_distributions(
    output_dir: Path,
    summary: dict,
    nulls: dict[str, pd.DataFrame],
) -> list[Path]:
    all_values = np.concatenate(
        [frame["mean_adjacency_contrast"].to_numpy(float) for frame in nulls.values()]
    )
    bins = np.linspace(all_values.min(), all_values.max(), 75)
    fig, axes = plt.subplots(1, 2, figsize=(7.15, 3.05), sharex=True, sharey=True)

    for panel, (axis, (model, config)) in enumerate(zip(axes, MODEL_CONFIG.items()), start=1):
        values = nulls[model]["mean_adjacency_contrast"].to_numpy(float)
        result = summary[config["summary_key"]]
        observed = float(result["observed"])
        q95 = float(np.quantile(values, 0.95, method="linear"))
        tail_count = int(np.count_nonzero(values >= observed))
        p_value = (tail_count + 1) / (len(values) + 1)
        if not np.isclose(q95, float(result["null_q95"]), atol=1e-12, rtol=0):
            raise ValueError(f"Null 95th-percentile mismatch for {model}")
        if tail_count != int(result["tail_count"]):
            raise ValueError(f"Upper-tail count mismatch for {model}")
        if not np.isclose(
            p_value, float(result["empirical_p_one_sided"]), atol=1e-15, rtol=0
        ):
            raise ValueError(f"Empirical p-value mismatch for {model}")

        axis.hist(
            values,
            bins=bins,
            density=True,
            color=config["color"],
            alpha=0.72,
            linewidth=0,
            rasterized=False,
        )
        axis.axvline(q95, color="#555555", linestyle=(0, (4, 2)), linewidth=1.2)
        axis.axvline(observed, color="#111111", linewidth=1.6)
        axis.set_title(f"{chr(64 + panel)}   {config['short']}", loc="left", fontweight="bold")
        axis.set_xlabel(r"Mean adjacency contrast, $T$")
        axis.text(
            0.035,
            0.96,
            f"observed = {observed:.3f}\n95th percentile = {q95:.3f}\none-sided empirical p = {p_value:.4f}",
            transform=axis.transAxes,
            ha="left",
            va="top",
            linespacing=1.35,
            bbox={"boxstyle": "round,pad=0.28", "fc": "white", "ec": "none", "alpha": 0.88},
        )
        axis.grid(axis="y", color="#D8D8D8", linewidth=0.55, alpha=0.7)
        axis.set_axisbelow(True)

    axes[0].set_ylabel("Density")
    axes[0].legend(
        handles=[
            Line2D([0], [0], color="#111111", lw=1.6, label="Observed"),
            Line2D([0], [0], color="#555555", lw=1.2, ls=(0, (4, 2)), label="Null 95th percentile"),
        ],
        frameon=False,
        loc="upper left",
        bbox_to_anchor=(0.0, -0.23),
        ncol=2,
        handlelength=2.5,
    )
    fig.suptitle(
        "Frozen 19-gene topology against conditional pseudoroute nulls",
        x=0.07,
        y=1.015,
        ha="left",
        fontsize=11,
        fontweight="bold",
    )
    add_status_note(fig)
    fig.subplots_adjust(left=0.09, right=0.995, top=0.86, bottom=0.27, wspace=0.13)
    return save_figure(fig, output_dir / "figure_2_conditional_nulls_v1_0")


def figure_3_component_decomposition(
    output_dir: Path,
    nulls: dict[str, pd.DataFrame],
    components: pd.DataFrame,
) -> list[Path]:
    fig, axes = plt.subplots(2, 2, figsize=(7.15, 4.75))
    y_positions = {"P1_technical": 1.0, "S1_neuron": 0.0}

    for panel, (axis, (component, title, xlabel)) in enumerate(
        zip(axes.ravel(), COMPONENT_CONFIG), start=1
    ):
        observed_values: list[float] = []
        interval_values: list[float] = []
        for model, config in MODEL_CONFIG.items():
            subset = components.loc[
                (components["model_id"] == model) & (components["component"] == component)
            ]
            if len(subset) != 1:
                raise ValueError(f"Expected one row for {model}/{component}")
            row = subset.iloc[0]
            values = nulls[model][component].to_numpy(float)
            q05, q95 = np.quantile(values, [0.05, 0.95], method="linear")
            null_mean = float(values.mean())
            observed = float(row["observed"])
            if not np.isclose(null_mean, float(row["null_mean"]), atol=1e-12, rtol=0):
                raise ValueError(f"Null mean mismatch for {model}/{component}")
            recorded_quantile = row["null_q05"] if row["tail"] == "lower" else row["null_q95"]
            calculated_quantile = q05 if row["tail"] == "lower" else q95
            if not np.isclose(calculated_quantile, float(recorded_quantile), atol=1e-12, rtol=0):
                raise ValueError(f"Tail quantile mismatch for {model}/{component}")

            y = y_positions[model]
            axis.plot([q05, q95], [y, y], color=config["color"], lw=4.2, solid_capstyle="butt")
            axis.scatter(
                [null_mean],
                [y],
                s=30,
                facecolor="white",
                edgecolor=config["color"],
                linewidth=1.25,
                zorder=3,
            )
            axis.scatter(
                [observed],
                [y],
                s=34,
                marker="D",
                facecolor="#111111",
                edgecolor="white",
                linewidth=0.45,
                zorder=4,
            )
            observed_values.append(observed)
            interval_values.extend([q05, q95])

        values_for_limits = np.asarray(interval_values + observed_values, dtype=float)
        span = float(np.ptp(values_for_limits))
        padding = 0.12 * span if span else 0.25
        if component == "both_adjacent_positive_donor_count":
            axis.set_xlim(-0.3, 6.35)
            axis.set_xticks(np.arange(0, 7, 1))
        else:
            axis.set_xlim(values_for_limits.min() - padding, values_for_limits.max() + padding)
        axis.set_ylim(-0.55, 1.55)
        axis.set_yticks([1.0, 0.0], ["P1 technical", "S1 neuron-matched"])
        if panel in (2, 4):
            axis.tick_params(axis="y", labelleft=False)
        axis.set_title(f"{chr(64 + panel)}   {title}", loc="left", fontweight="bold")
        axis.set_xlabel(xlabel)
        axis.grid(axis="x", color="#D8D8D8", linewidth=0.55, alpha=0.7)
        axis.set_axisbelow(True)

    fig.legend(
        handles=[
            Line2D([0], [0], color="#777777", lw=4.2, label="Conditional-null 5th–95th percentiles"),
            Line2D(
                [0],
                [0],
                color="#777777",
                marker="o",
                markerfacecolor="white",
                lw=0,
                label="Null mean",
            ),
            Line2D(
                [0],
                [0],
                color="#111111",
                marker="D",
                markerfacecolor="#111111",
                lw=0,
                label="Observed",
            ),
        ],
        frameon=False,
        loc="lower center",
        bbox_to_anchor=(0.5, -0.005),
        ncol=3,
        columnspacing=1.5,
        handlelength=2.2,
    )
    fig.suptitle(
        "Component decomposition relative to conditional nulls",
        x=0.075,
        y=1.012,
        ha="left",
        fontsize=11,
        fontweight="bold",
    )
    add_status_note(fig)
    fig.subplots_adjust(left=0.17, right=0.99, top=0.88, bottom=0.18, hspace=0.48, wspace=0.38)
    return save_figure(fig, output_dir / "figure_3_component_decomposition_v1_0")


def figure_4_lodo_sensitivity(output_dir: Path, lodo: pd.DataFrame) -> list[Path]:
    donors = lodo.loc[lodo["model_id"] == "P1_technical", "omitted_donor"].astype(str).tolist()
    if len(donors) != 6 or set(lodo["omitted_donor"].astype(str)) != set(donors):
        raise ValueError("Unexpected donor membership in LODO results")

    fig, axes = plt.subplots(1, 2, figsize=(7.15, 3.45), sharex=True, sharey=True)
    for panel, (axis, (model, config)) in enumerate(zip(axes, MODEL_CONFIG.items()), start=1):
        subset = (
            lodo.loc[lodo["model_id"] == model]
            .assign(omitted_donor=lambda frame: frame["omitted_donor"].astype(str))
            .set_index("omitted_donor")
            .loc[donors]
        )
        y = np.arange(len(donors))[::-1]
        observed = subset["observed"].to_numpy(float)
        q95 = subset["null_q95"].to_numpy(float)
        recorded_beyond = subset["observed_beyond_95pct_threshold"].astype(bool).to_numpy()
        beyond = observed > q95
        if not np.array_equal(beyond, recorded_beyond):
            raise ValueError(f"LODO threshold-indicator mismatch for {model}")

        for yi, obs, threshold in zip(y, observed, q95):
            axis.plot(
                [min(obs, threshold), max(obs, threshold)],
                [yi, yi],
                color="#B8B8B8",
                linewidth=1.0,
                zorder=1,
            )
        axis.scatter(
            q95,
            y,
            marker="|",
            s=145,
            linewidth=2.0,
            color=config["color"],
            label="Conditional-null 95th percentile",
            zorder=2,
        )
        axis.scatter(
            observed,
            y,
            marker="D",
            s=33,
            facecolor=np.where(beyond, "#111111", "white"),
            edgecolor="#111111",
            linewidth=0.9,
            label="Observed LODO effect",
            zorder=3,
        )
        axis.axvline(0, color="#777777", linestyle=(0, (2, 2)), linewidth=0.9)
        axis.set_title(f"{chr(64 + panel)}   {config['short']}", loc="left", fontweight="bold")
        axis.set_xlabel(r"LODO mean adjacency contrast, $T$")
        axis.grid(axis="x", color="#D8D8D8", linewidth=0.55, alpha=0.7)
        axis.set_axisbelow(True)
        axis.text(
            0.04,
            0.04,
            f"{int(beyond.sum())}/6 above null 95th percentile",
            transform=axis.transAxes,
            ha="left",
            va="bottom",
            color="#333333",
        )

    axes[0].set_yticks(np.arange(len(donors))[::-1], donors)
    axes[0].set_ylabel("Omitted AHBA donor")
    axes[0].set_xlim(-0.015, 0.465)
    axes[0].legend(
        handles=[
            Line2D(
                [0], [0], color="#111111", marker="D", markerfacecolor="#111111", lw=0,
                label="Observed above null 95th percentile",
            ),
            Line2D(
                [0], [0], color="#111111", marker="D", markerfacecolor="white", lw=0,
                label="Observed at or below null 95th percentile",
            ),
            Line2D(
                [0], [0], color="#777777", marker="|", markersize=10, markeredgewidth=2, lw=0,
                label="Null 95th percentile",
            ),
        ],
        frameon=False,
        loc="upper left",
        bbox_to_anchor=(0.0, -0.22),
        ncol=3,
        columnspacing=1.2,
        handletextpad=0.45,
    )
    fig.suptitle(
        "Leave-one-donor-out sensitivity (overlapping diagnostic)",
        x=0.07,
        y=1.015,
        ha="left",
        fontsize=11,
        fontweight="bold",
    )
    add_status_note(fig)
    fig.subplots_adjust(left=0.12, right=0.995, top=0.87, bottom=0.27, wspace=0.14)
    return save_figure(fig, output_dir / "figure_4_lodo_sensitivity_v1_0")


def main() -> None:
    parser = argparse.ArgumentParser(description=__doc__)
    parser.add_argument(
        "--analysis-dir",
        type=Path,
        default=Path(__file__).resolve().parents[1] / "analysis",
    )
    parser.add_argument(
        "--output-dir",
        type=Path,
        default=Path(__file__).resolve().parents[1] / "figures",
    )
    parser.add_argument(
        "--caption-source",
        type=Path,
        default=Path(__file__).resolve().with_name("figure_captions_v1_0.md"),
    )
    args = parser.parse_args()
    analysis_dir = args.analysis_dir.resolve()
    output_dir = args.output_dir.resolve()
    caption_source = args.caption_source.resolve()
    input_paths = verify_inputs(analysis_dir)
    if not caption_source.is_file():
        raise FileNotFoundError(f"Missing caption source: {caption_source}")
    output_dir.mkdir(parents=True, exist_ok=True)

    configure_matplotlib()
    summary, nulls, components, lodo, donors = load_results(analysis_dir)

    obsolete_bases = (
        "figure_1_conditional_nulls_v1_0",
        "figure_2_component_decomposition_v1_0",
        "figure_3_lodo_sensitivity_v1_0",
    )
    for base in obsolete_bases:
        for suffix in (".png", ".svg", ".pdf"):
            (output_dir / f"{base}{suffix}").unlink(missing_ok=True)

    outputs: list[Path] = []
    outputs += figure_1_observed_donor_structure(output_dir, summary, donors)
    outputs += figure_2_null_distributions(output_dir, summary, nulls)
    outputs += figure_3_component_decomposition(output_dir, nulls, components)
    outputs += figure_4_lodo_sensitivity(output_dir, lodo)

    caption_path = output_dir / "figure_captions_v1_0.md"
    if caption_source != caption_path.resolve():
        atomic_write_bytes(caption_path, caption_source.read_bytes())

    for output in outputs:
        verify_figure_file(output)

    manifest = {
        "version": "v1.0",
        "execution_revision": summary["execution_revision"],
        "interpretation_status": summary["interpretation_status"],
        "claim_guardrail": summary["claim_guardrail"],
        "generator_file": {
            "path": "scripts/plot_v1_0_results.py",
            "bytes": Path(__file__).resolve().stat().st_size,
            "sha256": sha256(Path(__file__).resolve()),
        },
        "caption_source": {
            "path": "scripts/figure_captions_v1_0.md",
            "bytes": caption_source.stat().st_size,
            "sha256": sha256(caption_source),
        },
        "runtime": {
            "python": platform.python_version(),
            "matplotlib": mpl.__version__,
            "numpy": np.__version__,
            "pandas": pd.__version__,
            "pillow": pillow_version,
            "backend": str(mpl.get_backend()),
            "font_family": "DejaVu Sans",
            "font_file_sha256": sha256(Path(font_manager.findfont("DejaVu Sans"))),
        },
        "input_files": {
            name: {"bytes": path.stat().st_size, "sha256": sha256(path)}
            for name, path in sorted(input_paths.items())
        },
        "output_files": {
            output.name: {"bytes": output.stat().st_size, "sha256": sha256(output)}
            for output in sorted(outputs)
        },
    }
    manifest["caption_file"] = {
        "path": caption_path.name,
        "bytes": caption_path.stat().st_size,
        "sha256": sha256(caption_path),
    }
    manifest_path = output_dir / "figure_manifest_v1_0.json"
    manifest_payload = (json.dumps(manifest, indent=2, sort_keys=True) + "\n").encode("utf-8")
    atomic_write_bytes(manifest_path, manifest_payload)
    print(f"Generated and verified {len(outputs)} figure files in {output_dir}")
    print(f"Manifest: {manifest_path} ({sha256(manifest_path)})")


if __name__ == "__main__":
    main()
```
