## Supplementary Archive S1 for "An anatomy-constrained transcriptomic screen identifies a 19-gene panel with an ordered regional similarity pattern in the human somatosensory system": verify_analysis_outputs_v1_0.py

```
#!/usr/bin/env python3
"""Structural and byte-reproducibility checks for published v1.0 outputs.

This validator does not calculate or reinterpret any endpoint. It rejects
missing, empty, malformed, hash-inconsistent or row-incomplete artifacts and
can compare a retained execution with an independent rerun byte for byte.
"""

from __future__ import annotations

import argparse
import hashlib
import json
from pathlib import Path

import pandas as pd


HERE = Path(__file__).resolve().parents[1]
DEFAULT_ANALYSIS = HERE / "analysis"

EXPECTED_ROWS = {
    "ahba_topology_null_P1_technical_v1_0.csv.gz": 50_000,
    "ahba_topology_null_S1_neuron_v1_0.csv.gz": 50_000,
    "cell_attribution_P1_technical_null_v1_0.csv.gz": 50_000,
    "geometry_matched_triplets_v1_0.csv": 240,
    "hpa_exact_node_topology_nulls_v1_0.csv.gz": 50_000,
    "lodo_null_tests_v1_0.csv": 12,
    "observed_topology_donor_v1_0.csv": 6,
    "probe_choice_diagnostic_v1_0.csv.gz": 50_000,
    "topology_component_tests_v1_0.csv": 8,
    "two_way_bootstrap_diagnostic_v1_0.csv.gz": 50_000,
    "verified_endpoint_inputs_v1_0.csv": 84,
}


def sha256(path: Path) -> str:
    digest = hashlib.sha256()
    with path.open("rb") as handle:
        for block in iter(lambda: handle.read(1024 * 1024), b""):
            digest.update(block)
    return digest.hexdigest()


def read_manifest(path: Path) -> dict[str, str]:
    entries: dict[str, str] = {}
    for line in path.read_text(encoding="utf-8").splitlines():
        digest, filename = line.split("  ", 1)
        if filename in entries or Path(filename).name != filename:
            raise RuntimeError(f"Malformed or duplicate manifest entry: {filename}")
        entries[filename] = digest
    return entries


def verify_directory(directory: Path) -> dict[str, object]:
    if not directory.is_dir():
        raise FileNotFoundError(directory)
    manifest_path = directory / "analysis_hashes_v1_0.txt"
    entries = read_manifest(manifest_path)
    actual = {
        path.name
        for path in directory.iterdir()
        if path.is_file() and path.name != manifest_path.name
    }
    if set(entries) != actual:
        raise RuntimeError("Output-directory membership differs from hash manifest")
    for filename, expected_hash in entries.items():
        path = directory / filename
        if path.stat().st_size <= 0:
            raise RuntimeError(f"Empty published artifact: {filename}")
        if sha256(path) != expected_hash:
            raise RuntimeError(f"Published-artifact hash mismatch: {filename}")
    for filename, expected_rows in EXPECTED_ROWS.items():
        frame = pd.read_csv(directory / filename)
        if len(frame) != expected_rows:
            raise RuntimeError(
                f"Wrong row count for {filename}: {len(frame)} != {expected_rows}"
            )
    summary = json.loads((directory / "summary_v1_0.json").read_text(encoding="utf-8"))
    if summary.get("execution_revision") != "v1.0-A04_FLOAT32_LINEAGE_FIX":
        raise RuntimeError("Unexpected execution revision")
    ledger = pd.read_csv(directory / "verified_endpoint_inputs_v1_0.csv")
    if not ledger["verified_before_endpoint"].eq(True).all() or not ledger[
        "verified_after_endpoint"
    ].eq(True).all():
        raise RuntimeError("Pre/post input verification ledger is incomplete")
    return {
        "directory": str(directory),
        "manifest_sha256": sha256(manifest_path),
        "summary_sha256": sha256(directory / "summary_v1_0.json"),
        "artifact_count_excluding_manifest": len(entries),
        "row_count_checks": len(EXPECTED_ROWS),
        "status": "VALID_COMPLETE_OUTPUT_SET",
    }


def compare_directories(first: Path, second: Path) -> dict[str, object]:
    first_files = {path.name for path in first.iterdir() if path.is_file()}
    second_files = {path.name for path in second.iterdir() if path.is_file()}
    if first_files != second_files:
        raise RuntimeError("Compared output directories have different membership")
    differences = [
        filename
        for filename in sorted(first_files)
        if sha256(first / filename) != sha256(second / filename)
    ]
    if differences:
        raise RuntimeError(f"Byte-level rerun differences: {differences}")
    return {
        "reference_directory": str(second),
        "compared_file_count": len(first_files),
        "byte_identical": True,
    }


def main() -> int:
    parser = argparse.ArgumentParser()
    parser.add_argument("analysis", nargs="?", type=Path, default=DEFAULT_ANALYSIS)
    parser.add_argument("--reference", type=Path)
    arguments = parser.parse_args()
    report = verify_directory(arguments.analysis)
    if arguments.reference is not None:
        verify_directory(arguments.reference)
        report["rerun_comparison"] = compare_directories(
            arguments.analysis, arguments.reference
        )
    print(json.dumps(report, indent=2, sort_keys=True))
    return 0


if __name__ == "__main__":
    raise SystemExit(main())
```
