## Supplementary Archive S1 for "An anatomy-constrained transcriptomic screen identifies a 19-gene panel with an ordered regional similarity pattern in the human somatosensory system": analyze_spatial_v1_1_A02_failed.py

```
#!/usr/bin/env python3
"""Run the frozen v1.1 joint Moran-spectral spatial robustness analysis.

This is a post-discovery robustness extension.  It preserves fitted spatial
trends and jointly randomizes the residual fields of all 19 genes.  It cannot
establish connectivity, direction, route exclusivity, causality, disease
mechanism, a stimulation target, or independent replication.
"""

from __future__ import annotations

import gzip
import hashlib
import json
import math
import os
import platform
from dataclasses import dataclass
from pathlib import Path
from typing import Any

import numpy as np
import pandas as pd
import scipy
from scipy.linalg import eigh, qr
from scipy.stats import rankdata


HERE = Path(__file__).resolve().parents[1]
PROJECT = HERE.parents[1]
PROTOCOL_PATH = HERE / "protocol" / "protocol_v1_1.json"
AMENDMENT1_PATH = (
    HERE / "protocol" / "amendment_01_double_gate_and_trend_preservation_v1_1.json"
)
AMENDMENT2_PATH = HERE / "protocol" / "amendment_02_endpoint_seal_v1_1.json"
OUT = HERE / "analysis"
STAGING = HERE / "analysis.staging"

V08 = PROJECT / "reconstructed_v008" / "Paquete_reproducible_rutas_transcriptomicas_v0_8"
V10 = PROJECT / "reconstructed_v010" / "Paquete_reproducible_rutas_transcriptomicas_v0_10"
V1 = PROJECT / "work" / "route_v1_0"

GENES = [
    "ATP2C1",
    "RAI2",
    "RAB3GAP2",
    "CCT6A",
    "MAP1B",
    "SH3BGRL2",
    "PDHX",
    "C7orf25",
    "TSHZ2",
    "NEFM",
    "ATXN3",
    "ADAM23",
    "HMGCS1",
    "SLC16A6",
    "NANS",
    "IFT57",
    "MEF2A",
    "BBS7",
    "RAP1GDS1",
]
DONORS = [9861, 10021, 12876, 14380, 15496, 15697]
NODES = ["Cu/Gr", "DTLv", "Postcentral"]
COMPARTMENTS = ["myelencephalon", "dorsal thalamus", "parietal lobe"]
EXPECTED_SOURCE_SAMPLES = 3702
EXPECTED_ELIGIBLE_SAMPLES = 2797
EXPECTED_ELIGIBLE_BY_DONOR = {
    9861: 481,
    10021: 453,
    12876: 363,
    14380: 529,
    15496: 470,
    15697: 501,
}
EXPECTED_TARGET_COUNTS = {
    9861: [3, 5, 12],
    10021: [5, 4, 12],
    12876: [1, 3, 13],
    14380: [5, 2, 10],
    15496: [2, 3, 5],
    15697: [3, 3, 10],
}
EXPECTED_BLOCK_COUNTS = {
    9861: [39, 22, 39],
    10021: [46, 19, 26],
    12876: [16, 21, 37],
    14380: [44, 23, 42],
    15496: [35, 16, 32],
    15697: [23, 19, 41],
}
MODEL_SPECS = [
    {
        "model_id": "P1A_BLOCKWISE_KNN5_TREND_PRESERVED",
        "scope": "block",
        "weight": "knn",
        "k": 5,
        "coordinate": "native",
        "draws": 50_000,
        "root_seed": 20260912,
        "primary_component": True,
    },
    {
        "model_id": "P1B_WHOLE_DONOR_KNN6_FIXED_EFFECTS",
        "scope": "whole",
        "weight": "knn",
        "k": 6,
        "coordinate": "native",
        "draws": 50_000,
        "root_seed": 20260913,
        "primary_component": True,
    },
    {
        "model_id": "S1_BLOCKWISE_DENSE_NATIVE",
        "scope": "block",
        "weight": "dense",
        "k": None,
        "coordinate": "native",
        "draws": 10_000,
        "root_seed": 20260914,
        "primary_component": False,
    },
    {
        "model_id": "S2_WHOLE_DONOR_DENSE_NATIVE",
        "scope": "whole",
        "weight": "dense",
        "k": None,
        "coordinate": "native",
        "draws": 10_000,
        "root_seed": 20260915,
        "primary_component": False,
    },
    {
        "model_id": "S3A_BLOCKWISE_KNN5_AXIS_Z",
        "scope": "block",
        "weight": "knn",
        "k": 5,
        "coordinate": "axis_z",
        "draws": 10_000,
        "root_seed": 20260916,
        "primary_component": False,
    },
    {
        "model_id": "S3B_WHOLE_DONOR_KNN8_AXIS_Z",
        "scope": "whole",
        "weight": "knn",
        "k": 8,
        "coordinate": "axis_z",
        "draws": 10_000,
        "root_seed": 20260917,
        "primary_component": False,
    },
]
BATCH_SIZE = 200
AUDIT_DRAWS = 100
TAIL_TOLERANCE = 1e-15


def sha256(path: Path) -> str:
    digest = hashlib.sha256()
    with path.open("rb") as handle:
        for block in iter(lambda: handle.read(1024 * 1024), b""):
            digest.update(block)
    return digest.hexdigest()


def write_json(path: Path, obj: Any) -> None:
    path.write_text(json.dumps(obj, indent=2, sort_keys=True) + "\n", encoding="utf-8")


def csv_bytes(frame: pd.DataFrame) -> bytes:
    return frame.to_csv(
        index=False,
        float_format="%.17g",
        lineterminator="\n",
    ).encode("utf-8")


def write_csv(path: Path, frame: pd.DataFrame) -> None:
    path.write_bytes(csv_bytes(frame))


def write_csv_gzip(path: Path, frame: pd.DataFrame) -> None:
    with path.open("wb") as raw:
        with gzip.GzipFile(filename="", mode="wb", fileobj=raw, mtime=0) as handle:
            handle.write(csv_bytes(frame))


def verify_static_inputs(
    protocol: dict[str, Any], amendment1: dict[str, Any], amendment2: dict[str, Any]
) -> pd.DataFrame:
    if sha256(PROTOCOL_PATH) != amendment1["parent_protocol_sha256"]:
        raise RuntimeError("A01 parent protocol hash mismatch")
    if sha256(PROTOCOL_PATH) != amendment2["parent_protocol_sha256"]:
        raise RuntimeError("A02 parent protocol hash mismatch")
    if sha256(AMENDMENT1_PATH) != amendment2["parent_amendment_A01_sha256"]:
        raise RuntimeError("A02 parent amendment hash mismatch")
    if sha256(Path(__file__).resolve()) != amendment2["endpoint_script_sha256"]:
        raise RuntimeError("Endpoint script differs from A02 seal")

    paths = {
        "v08_sample_metadata": V08 / "analysis" / "ahba_sample_metadata_v0_8.csv",
        "v08_public_raw_manifest": V08 / "protocol" / "public_raw_manifest_v0_8.csv",
        "v08_probe_validation": V08 / "protocol" / "ahba_probe_validation_v0_8.json",
        "v10_frozen_residuals": V10
        / "analysis"
        / "primary_node_residuals_candidates_and_controls_v0_10.csv.gz",
        "v10_lineage_script": V10 / "scripts" / "analyze_specificity_v0_10.py",
        "v10_protocol": V1 / "protocol" / "protocol_v1_0.json",
        "v10_A04": V1 / "protocol" / "amendment_04_float32_lineage_fix_v1_0.json",
        "v10_result_summary": V1 / "analysis" / "summary_v1_0.json",
    }
    expected = {item["id"]: item for item in protocol["input_files"]}
    if set(paths) != set(expected):
        raise RuntimeError("Protocol input identifiers differ from implementation")
    rows: list[dict[str, Any]] = []
    for input_id, path in paths.items():
        if not path.is_file():
            raise FileNotFoundError(path)
        observed = sha256(path)
        if observed != expected[input_id]["sha256"]:
            raise RuntimeError(f"Input hash mismatch: {input_id}")
        rows.append(
            {
                "source_id": input_id,
                "path": str(path.relative_to(PROJECT)),
                "bytes": path.stat().st_size,
                "sha256": observed,
                "verified_before_endpoint": True,
            }
        )
    for input_id, path in [
        ("protocol_v1_1", PROTOCOL_PATH),
        ("amendment_v1_1_A01", AMENDMENT1_PATH),
        ("amendment_v1_1_A02", AMENDMENT2_PATH),
        ("endpoint_script_v1_1", Path(__file__).resolve()),
    ]:
        rows.append(
            {
                "source_id": input_id,
                "path": str(path.relative_to(PROJECT)),
                "bytes": path.stat().st_size,
                "sha256": sha256(path),
                "verified_before_endpoint": True,
            }
        )
    return pd.DataFrame(rows)


def validate_metadata(metadata: pd.DataFrame) -> pd.DataFrame:
    required = {
        "sample_index",
        "donor_id",
        "well_id",
        "mri_x",
        "mri_y",
        "mri_z",
        "structure_name",
        "structure_abbreviation",
        "top_level_structure_name",
        "hemisphere_class",
    }
    if not required.issubset(metadata.columns):
        raise RuntimeError("Frozen metadata is missing required columns")
    if len(metadata) != EXPECTED_SOURCE_SAMPLES:
        raise RuntimeError("Unexpected frozen source-sample count")
    if metadata["sample_index"].tolist() != list(range(EXPECTED_SOURCE_SAMPLES)):
        raise RuntimeError("Frozen sample_index is not the expected ordered range")
    if metadata[["donor_id", "well_id"]].duplicated().any():
        raise RuntimeError("Duplicate donor/well sample keys")
    if not np.isfinite(metadata[["mri_x", "mri_y", "mri_z"]].to_numpy(float)).all():
        raise RuntimeError("Nonfinite MRI coordinate")
    eligible = metadata[metadata["hemisphere_class"].eq("left_or_unpaired")]
    if len(eligible) != EXPECTED_ELIGIBLE_SAMPLES:
        raise RuntimeError("Unexpected left-or-unpaired fitted-sample count")
    counts = eligible.groupby("donor_id").size().to_dict()
    if counts != EXPECTED_ELIGIBLE_BY_DONOR:
        raise RuntimeError(f"Unexpected eligible donor counts: {counts}")
    if sorted(metadata["donor_id"].unique().tolist()) != DONORS:
        raise RuntimeError("Unexpected donor identities")
    return eligible


def load_expression_maps(
    metadata: pd.DataFrame, ledger: pd.DataFrame
) -> tuple[np.ndarray, pd.DataFrame]:
    validation_path = V08 / "protocol" / "ahba_probe_validation_v0_8.json"
    validation = json.loads(validation_path.read_text(encoding="utf-8"))
    selected = [
        row
        for row in validation
        if row["symbol_requested"] in GENES
        and bool(row["api_success"])
        and bool(row["exact_symbol_confirmed"])
    ]
    selected.sort(key=lambda row: (GENES.index(row["symbol_requested"]), int(row["probe_id"])))
    if len(selected) != 44:
        raise RuntimeError(f"Expected 44 frozen candidate probes, found {len(selected)}")
    observed_gene_order = list(dict.fromkeys(row["symbol_requested"] for row in selected))
    if observed_gene_order != GENES:
        raise RuntimeError("Probe-validation genes differ from frozen order")

    manifest = pd.read_csv(V08 / "protocol" / "public_raw_manifest_v0_8.csv")
    manifest = manifest.set_index("relative_path", verify_integrity=True)
    expected_keys = list(
        zip(metadata["donor_id"].astype(int).tolist(), metadata["well_id"].astype(int).tolist())
    )
    probe_values: dict[str, list[np.ndarray]] = {gene: [] for gene in GENES}
    raw_rows: list[dict[str, Any]] = []
    for row in selected:
        relative = str(row["raw_path"])
        if relative not in manifest.index:
            raise RuntimeError(f"Raw probe path absent from v0.8 manifest: {relative}")
        path = V08 / relative
        wanted_hash = str(manifest.loc[relative, "sha256"])
        wanted_bytes = int(manifest.loc[relative, "bytes"])
        if path.stat().st_size != wanted_bytes or sha256(path) != wanted_hash:
            raise RuntimeError(f"Raw probe file failed manifest verification: {relative}")
        with gzip.open(path, "rt", encoding="utf-8") as handle:
            obj = json.load(handle)
        if not obj.get("success"):
            raise RuntimeError(f"Allen response unsuccessful: {relative}")
        samples = obj["msg"]["samples"]
        probes = obj["msg"]["probes"]
        if len(samples) != EXPECTED_SOURCE_SAMPLES or len(probes) != 1:
            raise RuntimeError(f"Unexpected Allen response dimensions: {relative}")
        probe = probes[0]
        if int(probe["id"]) != int(row["probe_id"]):
            raise RuntimeError(f"Probe id mismatch: {relative}")
        if str(probe.get("gene-symbol") or "") != row["symbol_requested"]:
            raise RuntimeError(f"Probe symbol mismatch: {relative}")
        api_keys = [
            (int(sample["donor"]["id"]), int(sample["sample"]["well"])) for sample in samples
        ]
        if len(set(api_keys)) != len(api_keys) or set(api_keys) != set(expected_keys):
            raise RuntimeError(f"Probe sample identities differ: {relative}")
        api_position = {key: index for index, key in enumerate(api_keys)}
        reorder = np.asarray([api_position[key] for key in expected_keys], dtype=int)
        z = np.asarray(probe["z-score"], dtype=np.float32)[reorder]
        if z.shape != (EXPECTED_SOURCE_SAMPLES,) or not np.isfinite(z).all():
            raise RuntimeError(f"Invalid z-score vector: {relative}")
        probe_values[row["symbol_requested"]].append(z)
        raw_rows.append(
            {
                "source_id": f"raw_probe::{row['symbol_requested']}::{int(row['probe_id'])}",
                "path": str(path.relative_to(PROJECT)),
                "bytes": path.stat().st_size,
                "sha256": wanted_hash,
                "verified_before_endpoint": True,
            }
        )

    collapsed = []
    for gene in GENES:
        values = np.median(np.stack(probe_values[gene], axis=0), axis=0).astype(np.float32)
        collapsed.append(values)
    expression = np.column_stack(collapsed).astype(np.float64)
    if expression.shape != (EXPECTED_SOURCE_SAMPLES, len(GENES)):
        raise RuntimeError("Unexpected collapsed expression shape")
    combined_ledger = pd.concat([ledger, pd.DataFrame(raw_rows)], ignore_index=True)
    return expression, combined_ledger


def target_mask(metadata: pd.DataFrame, node: str) -> pd.Series:
    if node == "Cu/Gr":
        return metadata["structure_abbreviation"].isin(["Cu", "Gr"])
    if node == "DTLv":
        return metadata["structure_abbreviation"].eq("DTLv")
    if node == "Postcentral":
        return metadata["structure_name"].str.lower().str.startswith("postcentral gyrus")
    raise ValueError(node)


def observed_tensor(metadata: pd.DataFrame, expression: np.ndarray) -> np.ndarray:
    tensor = np.full((len(GENES), len(DONORS), len(NODES)), np.nan, dtype=float)
    left = metadata["hemisphere_class"].eq("left_or_unpaired")
    for donor_index, donor in enumerate(DONORS):
        for node_index, (node, compartment) in enumerate(zip(NODES, COMPARTMENTS)):
            block = left & metadata["donor_id"].eq(donor) & metadata[
                "top_level_structure_name"
            ].eq(compartment)
            target = block & target_mask(metadata, node)
            sibling = block & ~target_mask(metadata, node)
            block_count = int(block.sum())
            target_count = int(target.sum())
            if block_count != EXPECTED_BLOCK_COUNTS[donor][node_index]:
                raise RuntimeError("Unexpected macrocompartment block count")
            if target_count != EXPECTED_TARGET_COUNTS[donor][node_index]:
                raise RuntimeError("Unexpected target sample count")
            if not target_count or not int(sibling.sum()):
                raise RuntimeError("Empty target or sibling set")
            tensor[:, donor_index, node_index] = np.median(
                expression[target.to_numpy(), :], axis=0
            ) - np.median(expression[sibling.to_numpy(), :], axis=0)
    if not np.isfinite(tensor).all():
        raise RuntimeError("Nonfinite observed residual tensor")
    return tensor


def validate_observed_replay(tensor: np.ndarray) -> tuple[float, pd.DataFrame, float]:
    frozen = pd.read_csv(
        V10 / "analysis" / "primary_node_residuals_candidates_and_controls_v0_10.csv.gz"
    )
    frozen = frozen[frozen["set_role"].eq("frozen_candidate")].copy()
    expected_rows = []
    for gene_index, gene in enumerate(GENES):
        for donor_index, donor in enumerate(DONORS):
            for node_index, node in enumerate(NODES):
                match = frozen[
                    frozen["symbol"].eq(gene)
                    & frozen["donor_id"].eq(donor)
                    & frozen["node"].eq(node)
                ]
                if len(match) != 1:
                    raise RuntimeError("Frozen candidate residual lookup is not unique")
                expected_rows.append(
                    abs(
                        tensor[gene_index, donor_index, node_index]
                        - float(match.iloc[0]["node_residual_z"])
                    )
                )
    max_residual_difference = float(max(expected_rows))
    if max_residual_difference > 1e-12:
        raise RuntimeError("Observed residual tensor does not reproduce v0.10")
    donor_frame = donor_topology_from_tensor(tensor)
    v1_donor = pd.read_csv(V1 / "analysis" / "observed_topology_donor_v1_0.csv")
    columns = [
        "rho_CuGr_DTLv",
        "rho_DTLv_Postcentral",
        "rho_CuGr_Postcentral",
        "adjacency_contrast",
    ]
    if v1_donor["donor_id"].astype(int).tolist() != DONORS:
        raise RuntimeError("v1.0 donor table order differs")
    max_endpoint_difference = float(
        np.max(np.abs(donor_frame[columns].to_numpy() - v1_donor[columns].to_numpy()))
    )
    if max_endpoint_difference > 1e-12:
        raise RuntimeError("Observed endpoint does not reproduce v1.0 donor table")
    observed = float(donor_frame["adjacency_contrast"].mean())
    v1_summary = json.loads((V1 / "analysis" / "summary_v1_0.json").read_text())
    expected_observed = float(
        v1_summary["observed_ahba_topology"]["mean_adjacency_contrast"]
    )
    if abs(observed - expected_observed) > 1e-12:
        raise RuntimeError("Observed aggregate endpoint does not reproduce v1.0")
    return max_residual_difference, donor_frame, max_endpoint_difference


def centered_ranks(values: np.ndarray) -> np.ndarray:
    ranks = rankdata(values, axis=-1, method="average").astype(float)
    ranks -= ranks.mean(axis=-1, keepdims=True)
    return ranks


def row_correlation(first: np.ndarray, second: np.ndarray) -> np.ndarray:
    denominator = np.sqrt(np.square(first).sum(axis=-1) * np.square(second).sum(axis=-1))
    return np.divide(
        (first * second).sum(axis=-1),
        denominator,
        out=np.full(first.shape[:-1], np.nan, dtype=float),
        where=denominator > 0,
    )


def donor_topology_from_tensor(tensor: np.ndarray) -> pd.DataFrame:
    rows = []
    for donor_index, donor in enumerate(DONORS):
        profiles = tensor[:, donor_index, :].T
        ranks = centered_ranks(profiles)
        r01 = float(row_correlation(ranks[0], ranks[1]))
        r12 = float(row_correlation(ranks[1], ranks[2]))
        r02 = float(row_correlation(ranks[0], ranks[2]))
        rows.append(
            {
                "donor_id": donor,
                "rho_CuGr_DTLv": r01,
                "rho_DTLv_Postcentral": r12,
                "rho_CuGr_Postcentral": r02,
                "adjacency_contrast": (r01 + r12) / 2 - r02,
            }
        )
    return pd.DataFrame(rows)


def distance_matrix(coordinates: np.ndarray) -> np.ndarray:
    delta = coordinates[:, None, :] - coordinates[None, :, :]
    distances = np.sqrt(np.square(delta).sum(axis=2))
    if not np.isfinite(distances).all():
        raise RuntimeError("Nonfinite spatial distance")
    off_diagonal = ~np.eye(len(coordinates), dtype=bool)
    if np.any(distances[off_diagonal] <= 0):
        raise RuntimeError("Duplicate coordinates within fitted spatial block")
    return distances


def weight_matrix(
    coordinates: np.ndarray,
    sample_indices: np.ndarray,
    kind: str,
    k: int | None,
) -> tuple[np.ndarray, int, bool]:
    distances = distance_matrix(coordinates)
    n = len(coordinates)
    if kind == "dense":
        weights = np.divide(
            1.0,
            distances,
            out=np.zeros_like(distances),
            where=distances > 0,
        )
    elif kind == "knn":
        if k is None or not (1 <= k < n):
            raise RuntimeError("Invalid frozen k for spatial block")
        directed = np.zeros((n, n), dtype=bool)
        for row in range(n):
            order = np.lexsort((sample_indices, distances[row]))
            neighbours = [index for index in order if index != row][:k]
            directed[row, neighbours] = True
        adjacency = directed | directed.T
        weights = np.divide(
            1.0,
            distances,
            out=np.zeros_like(distances),
            where=adjacency,
        )
    else:
        raise ValueError(kind)
    np.fill_diagonal(weights, 0.0)
    if not np.allclose(weights, weights.T, atol=0, rtol=0):
        raise RuntimeError("Spatial weight matrix is not exactly symmetric")
    adjacency = weights > 0
    visited = {0}
    stack = [0]
    while stack:
        current = stack.pop()
        for neighbour in np.flatnonzero(adjacency[current]):
            neighbour = int(neighbour)
            if neighbour not in visited:
                visited.add(neighbour)
                stack.append(neighbour)
    connected = len(visited) == n
    edge_count = int(np.triu(adjacency, 1).sum())
    return weights, edge_count, connected


def standardize_coordinates(coordinates: np.ndarray) -> np.ndarray:
    scale = coordinates.std(axis=0, ddof=0)
    if np.any(scale <= 0) or not np.isfinite(scale).all():
        raise RuntimeError("Coordinate axis cannot be standardized")
    return (coordinates - coordinates.mean(axis=0)) / scale


def block_design(coordinates_native: np.ndarray) -> tuple[np.ndarray, list[str]]:
    xyz = standardize_coordinates(coordinates_native)
    return np.column_stack([np.ones(len(xyz)), xyz]), ["intercept", "x", "y", "z"]


def whole_design(
    metadata_block: pd.DataFrame, coordinates_native: np.ndarray
) -> tuple[np.ndarray, list[str], list[str]]:
    xyz = standardize_coordinates(coordinates_native)
    levels = sorted(metadata_block["top_level_structure_name"].astype(str).unique())
    reference = levels[0]
    columns = [np.ones(len(metadata_block)), xyz[:, 0], xyz[:, 1], xyz[:, 2]]
    names = ["intercept", "x", "y", "z"]
    observed = metadata_block["top_level_structure_name"].astype(str).to_numpy()
    for level in levels[1:]:
        columns.append((observed == level).astype(float))
        names.append(f"top::{level}")
    return np.column_stack(columns), names, [reference, *levels[1:]]


def canonicalize_columns(vectors: np.ndarray) -> np.ndarray:
    result = vectors.copy()
    for column in range(result.shape[1]):
        pivot = int(np.argmax(np.abs(result[:, column])))
        if result[pivot, column] < 0:
            result[:, column] *= -1
    return result


@dataclass
class SpatialFit:
    model_id: str
    block_id: str
    donor_id: int
    sample_indices: np.ndarray
    expression: np.ndarray
    design: np.ndarray
    design_names: list[str]
    factor_levels: list[str]
    weights: np.ndarray
    fixed: np.ndarray
    residual: np.ndarray
    vectors: np.ndarray
    eigenvalues: np.ndarray
    coefficients: np.ndarray
    edge_count: int
    connected: bool


def fit_spatial_block(
    model_id: str,
    block_id: str,
    donor_id: int,
    metadata_block: pd.DataFrame,
    expression_block: np.ndarray,
    scope: str,
    weight_kind: str,
    k: int | None,
    coordinate_mode: str,
    donor_axis_z: np.ndarray | None,
) -> SpatialFit:
    sample_indices = metadata_block["sample_index"].to_numpy(int)
    coordinates_native = metadata_block[["mri_x", "mri_y", "mri_z"]].to_numpy(float)
    if coordinate_mode == "native":
        coordinates_weight = coordinates_native
    elif coordinate_mode == "axis_z":
        if donor_axis_z is None:
            raise RuntimeError("Missing donor-standardized coordinates")
        coordinates_weight = donor_axis_z
    else:
        raise ValueError(coordinate_mode)
    if scope == "block":
        design, design_names = block_design(coordinates_native)
        factor_levels: list[str] = []
    elif scope == "whole":
        design, design_names, factor_levels = whole_design(metadata_block, coordinates_native)
    else:
        raise ValueError(scope)
    rank = int(np.linalg.matrix_rank(design))
    if rank != design.shape[1]:
        raise RuntimeError(f"Spatial design is rank deficient in {model_id}/{block_id}")
    weights, edge_count, connected = weight_matrix(
        coordinates_weight, sample_indices, weight_kind, k
    )
    if not connected:
        raise RuntimeError(f"Disconnected graph in {model_id}/{block_id}")

    q_full, _ = qr(design, mode="full", pivoting=False, check_finite=True)
    q_residual = q_full[:, rank:]
    if q_residual.shape[1] != len(metadata_block) - rank:
        raise RuntimeError("Wrong design-orthogonal basis dimension")
    reduced = q_residual.T @ weights @ q_residual
    eigenvalues, reduced_vectors = eigh(reduced, check_finite=True)
    order = np.argsort(eigenvalues, kind="stable")[::-1]
    eigenvalues = eigenvalues[order]
    vectors = q_residual @ reduced_vectors[:, order]
    vectors = canonicalize_columns(vectors)
    coefficients_fixed, *_ = np.linalg.lstsq(design, expression_block, rcond=None)
    fixed = design @ coefficients_fixed
    residual = expression_block - fixed
    coefficients = vectors.T @ residual
    arrays = [
        expression_block,
        design,
        weights,
        fixed,
        residual,
        vectors,
        eigenvalues,
        coefficients,
    ]
    if any(not np.isfinite(array).all() for array in arrays):
        raise RuntimeError(f"Nonfinite spatial fit in {model_id}/{block_id}")
    residual_variance = residual.var(axis=0, ddof=1)
    if np.any(residual_variance <= 0):
        raise RuntimeError(f"Nonpositive residual gene variance in {model_id}/{block_id}")
    return SpatialFit(
        model_id=model_id,
        block_id=block_id,
        donor_id=donor_id,
        sample_indices=sample_indices,
        expression=expression_block,
        design=design,
        design_names=design_names,
        factor_levels=factor_levels,
        weights=weights,
        fixed=fixed,
        residual=residual,
        vectors=vectors,
        eigenvalues=eigenvalues,
        coefficients=coefficients,
        edge_count=edge_count,
        connected=connected,
    )


def moran_i(values: np.ndarray, weights: np.ndarray) -> np.ndarray:
    centered = values - values.mean(axis=0, keepdims=True)
    numerator = np.sum(centered * (weights @ centered), axis=0)
    denominator = np.sum(np.square(centered), axis=0)
    scale = len(values) / weights.sum()
    if np.any(denominator <= 0) or weights.sum() <= 0:
        raise RuntimeError("Moran I is undefined")
    return scale * numerator / denominator


def fit_qc(fit: SpatialFit) -> dict[str, Any]:
    vectors = fit.vectors
    design = fit.design
    residual = fit.residual
    reconstruction = fit.fixed + vectors @ fit.coefficients
    residual_variance = residual.var(axis=0, ddof=1)
    orthogonality = float(
        np.max(np.abs(vectors.T @ vectors - np.eye(vectors.shape[1])))
    )
    design_orthogonality = float(np.max(np.abs(design.T @ vectors)))
    reconstruction_difference = float(np.max(np.abs(reconstruction - fit.expression)))
    if vectors.shape[1] != len(fit.sample_indices) - design.shape[1]:
        raise RuntimeError("Unexpected retained Moran mode count")
    if orthogonality > 1e-9 or design_orthogonality > 1e-9:
        raise RuntimeError("Moran basis failed frozen orthogonality gate")
    if reconstruction_difference > 1e-9:
        raise RuntimeError("All-plus reconstruction failed frozen gate")
    return {
        "model_id": fit.model_id,
        "block_id": fit.block_id,
        "donor_id": fit.donor_id,
        "sample_count": len(fit.sample_indices),
        "design_rank": fit.design.shape[1],
        "moran_mode_count": vectors.shape[1],
        "edge_count": fit.edge_count,
        "connected": fit.connected,
        "design_columns": ";".join(fit.design_names),
        "factor_levels_reference_first": ";".join(fit.factor_levels),
        "max_basis_orthogonality_error": orthogonality,
        "max_design_orthogonality_error": design_orthogonality,
        "max_all_plus_reconstruction_error": reconstruction_difference,
        "minimum_residual_gene_variance_ddof1": float(residual_variance.min()),
        "maximum_residual_gene_variance_ddof1": float(residual_variance.max()),
        "minimum_positive_edge_distance": float(1.0 / fit.weights[fit.weights > 0].max()),
        "maximum_positive_edge_distance": float(1.0 / fit.weights[fit.weights > 0].min()),
        "minimum_eigenvalue": float(fit.eigenvalues.min()),
        "maximum_eigenvalue": float(fit.eigenvalues.max()),
        "minimum_adjacent_sorted_eigenvalue_gap": float(
            np.min(np.abs(np.diff(fit.eigenvalues)))
        )
        if len(fit.eigenvalues) > 1
        else np.nan,
    }


def audit_surrogates(fit: SpatialFit, signs: np.ndarray) -> dict[str, float]:
    audit_n = min(AUDIT_DRAWS, len(signs))
    design_pseudoinverse = np.linalg.pinv(fit.design)
    original_sd = fit.residual.std(axis=0, ddof=1)
    original_covariance = np.cov(fit.residual, rowvar=False, ddof=1)
    original_corr = np.corrcoef(fit.residual, rowvar=False)
    original_moran = moran_i(fit.residual, fit.weights)
    if not all(
        np.isfinite(array).all()
        for array in [original_sd, original_covariance, original_corr, original_moran]
    ):
        raise RuntimeError(f"Nonfinite original invariant in {fit.model_id}/{fit.block_id}")
    max_mean = 0.0
    max_fitted_nuisance = 0.0
    max_sd = 0.0
    max_covariance = 0.0
    max_corr = 0.0
    max_moran = 0.0
    for draw in range(audit_n):
        surrogate_residual = fit.vectors @ (
            fit.coefficients * signs[draw, :, None]
        )
        surrogate_sd = surrogate_residual.std(axis=0, ddof=1)
        surrogate_covariance = np.cov(surrogate_residual, rowvar=False, ddof=1)
        surrogate_corr = np.corrcoef(surrogate_residual, rowvar=False)
        surrogate_moran = moran_i(surrogate_residual, fit.weights)
        surrogate_fitted_nuisance = fit.design @ (
            design_pseudoinverse @ (fit.fixed + surrogate_residual)
        )
        if not all(
            np.isfinite(array).all()
            for array in [
                surrogate_residual,
                surrogate_sd,
                surrogate_covariance,
                surrogate_corr,
                surrogate_moran,
                surrogate_fitted_nuisance,
            ]
        ):
            raise RuntimeError(
                f"Nonfinite surrogate invariant in {fit.model_id}/{fit.block_id}"
            )
        max_mean = max(max_mean, float(np.max(np.abs(surrogate_residual.mean(axis=0)))))
        max_fitted_nuisance = max(
            max_fitted_nuisance,
            float(np.max(np.abs(surrogate_fitted_nuisance - fit.fixed))),
        )
        max_sd = max(
            max_sd,
            float(np.max(np.abs(surrogate_sd - original_sd))),
        )
        max_covariance = max(
            max_covariance,
            float(np.max(np.abs(surrogate_covariance - original_covariance))),
        )
        max_corr = max(
            max_corr,
            float(np.max(np.abs(surrogate_corr - original_corr))),
        )
        max_moran = max(
            max_moran,
            float(np.max(np.abs(surrogate_moran - original_moran))),
        )
    if (
        max_mean > 1e-8
        or max_fitted_nuisance > 1e-9
        or max_sd > 1e-8
        or max_covariance > 1e-8
        or max_corr > 1e-8
        or max_moran > 1e-8
    ):
        raise RuntimeError(f"Surrogate invariant audit failed in {fit.model_id}/{fit.block_id}")
    return {
        "audit_draws": audit_n,
        "max_residual_mean_abs": max_mean,
        "max_fitted_nuisance_difference": max_fitted_nuisance,
        "max_residual_sd_difference_ddof1": max_sd,
        "max_residual_gene_covariance_difference_ddof1": max_covariance,
        "max_residual_gene_correlation_difference": max_corr,
        "max_residual_moran_i_difference": max_moran,
    }


def signs_for_fit(seed_sequence: np.random.SeedSequence, draws: int, modes: int) -> np.ndarray:
    rng = np.random.Generator(np.random.PCG64(seed_sequence))
    return rng.integers(0, 2, size=(draws, modes), dtype=np.int8) * 2 - 1


def project_values(
    fit: SpatialFit,
    signs: np.ndarray,
    local_rows: np.ndarray,
) -> np.ndarray:
    draws = len(signs)
    output = np.empty((draws, len(local_rows), len(GENES)), dtype=np.float64)
    vectors = fit.vectors[local_rows, :]
    fixed = fit.fixed[local_rows, :]
    for start in range(0, draws, BATCH_SIZE):
        stop = min(start + BATCH_SIZE, draws)
        signed_coefficients = signs[start:stop, :, None] * fit.coefficients[None, :, :]
        output[start:stop] = fixed[None, :, :] + vectors @ signed_coefficients
    if not np.isfinite(output).all():
        raise RuntimeError("Nonfinite surrogate expression")
    return output


def generate_node_residuals(
    fit: SpatialFit,
    signs: np.ndarray,
    projected_local_rows: np.ndarray,
    node_pairs: list[tuple[np.ndarray, np.ndarray]],
) -> list[np.ndarray]:
    """Project in batches and retain only target-minus-sibling gene profiles."""
    draws = len(signs)
    vectors = fit.vectors[projected_local_rows, :]
    fixed = fit.fixed[projected_local_rows, :]
    outputs = [np.empty((draws, len(GENES)), dtype=np.float64) for _ in node_pairs]
    for start in range(0, draws, BATCH_SIZE):
        stop = min(start + BATCH_SIZE, draws)
        signed_coefficients = (
            signs[start:stop, :, None] * fit.coefficients[None, :, :]
        )
        values = fixed[None, :, :] + vectors @ signed_coefficients
        for output, (target_rows, sibling_rows) in zip(outputs, node_pairs):
            output[start:stop] = np.median(values[:, target_rows, :], axis=1) - np.median(
                values[:, sibling_rows, :], axis=1
            )
    if any(not np.isfinite(output).all() for output in outputs):
        raise RuntimeError("Nonfinite surrogate node residual")
    return outputs


def donor_axis_z_lookup(metadata_donor: pd.DataFrame) -> dict[int, np.ndarray]:
    coordinates = metadata_donor[["mri_x", "mri_y", "mri_z"]].to_numpy(float)
    standardized = standardize_coordinates(coordinates)
    return {
        int(sample_index): standardized[row]
        for row, sample_index in enumerate(metadata_donor["sample_index"].to_numpy(int))
    }


def topology_from_draw_profiles(profiles: np.ndarray) -> tuple[np.ndarray, dict[str, np.ndarray]]:
    # profiles shape: draws x 3 nodes x 19 genes
    ranks = centered_ranks(profiles)
    r01 = row_correlation(ranks[:, 0, :], ranks[:, 1, :])
    r12 = row_correlation(ranks[:, 1, :], ranks[:, 2, :])
    r02 = row_correlation(ranks[:, 0, :], ranks[:, 2, :])
    contrast = (r01 + r12) / 2 - r02
    return contrast, {"rho01": r01, "rho12": r12, "rho02": r02}


def run_model(
    spec: dict[str, Any],
    metadata: pd.DataFrame,
    expression: np.ndarray,
) -> tuple[pd.DataFrame, list[dict[str, Any]], list[dict[str, Any]]]:
    model_id = spec["model_id"]
    draws = int(spec["draws"])
    left = metadata[metadata["hemisphere_class"].eq("left_or_unpaired")].copy()
    block_descriptors: list[tuple[int, str, pd.DataFrame]] = []
    if spec["scope"] == "block":
        for donor in DONORS:
            for compartment in COMPARTMENTS:
                frame = left[
                    left["donor_id"].eq(donor)
                    & left["top_level_structure_name"].eq(compartment)
                ].copy()
                block_descriptors.append((donor, compartment, frame))
    else:
        for donor in DONORS:
            frame = left[left["donor_id"].eq(donor)].copy()
            block_descriptors.append((donor, "whole_donor", frame))
    child_sequences = np.random.SeedSequence(int(spec["root_seed"])).spawn(
        len(block_descriptors)
    )
    donor_node_profiles: dict[int, list[np.ndarray]] = {donor: [] for donor in DONORS}
    qc_rows: list[dict[str, Any]] = []
    invariant_rows: list[dict[str, Any]] = []

    for descriptor_index, (donor, block_id, frame) in enumerate(block_descriptors):
        frame = frame.sort_values("sample_index", kind="stable")
        global_rows = frame["sample_index"].to_numpy(int)
        donor_frame = left[left["donor_id"].eq(donor)].sort_values(
            "sample_index", kind="stable"
        )
        axis_lookup = donor_axis_z_lookup(donor_frame)
        axis_z = np.vstack([axis_lookup[int(index)] for index in global_rows])
        fit = fit_spatial_block(
            model_id=model_id,
            block_id=block_id,
            donor_id=donor,
            metadata_block=frame,
            expression_block=expression[global_rows, :],
            scope=spec["scope"],
            weight_kind=spec["weight"],
            k=spec["k"],
            coordinate_mode=spec["coordinate"],
            donor_axis_z=axis_z,
        )
        base_qc = fit_qc(fit)
        signs = signs_for_fit(child_sequences[descriptor_index], draws, fit.vectors.shape[1])
        invariant = audit_surrogates(fit, signs[:AUDIT_DRAWS])
        qc_rows.append(base_qc)
        invariant_rows.append({**base_qc, **invariant})

        if spec["scope"] == "block":
            node_index = COMPARTMENTS.index(block_id)
            node = NODES[node_index]
            target_global = frame.loc[target_mask(frame, node), "sample_index"].to_numpy(int)
            target_set = set(target_global.tolist())
            local_target = np.asarray(
                [index for index, sample in enumerate(global_rows) if int(sample) in target_set],
                dtype=int,
            )
            local_sibling = np.asarray(
                [index for index in range(len(global_rows)) if index not in set(local_target.tolist())],
                dtype=int,
            )
            residuals = generate_node_residuals(
                fit,
                signs,
                np.arange(len(global_rows), dtype=int),
                [(local_target, local_sibling)],
            )[0]
            donor_node_profiles[donor].append(residuals)
        else:
            relevant = frame["top_level_structure_name"].isin(COMPARTMENTS).to_numpy()
            local_relevant = np.flatnonzero(relevant)
            relevant_frame = frame.iloc[local_relevant].copy()
            node_pairs: list[tuple[np.ndarray, np.ndarray]] = []
            for node, compartment in zip(NODES, COMPARTMENTS):
                in_compartment = relevant_frame["top_level_structure_name"].eq(compartment)
                in_target = in_compartment & target_mask(relevant_frame, node)
                in_sibling = in_compartment & ~target_mask(relevant_frame, node)
                local_target = np.flatnonzero(in_target.to_numpy())
                local_sibling = np.flatnonzero(in_sibling.to_numpy())
                if not len(local_target) or not len(local_sibling):
                    raise RuntimeError("Whole-donor target/sibling projection is empty")
                node_pairs.append((local_target, local_sibling))
            residual_profiles = generate_node_residuals(
                fit, signs, local_relevant, node_pairs
            )
            full_audit_signs = signs[:10]
            full_values = project_values(
                fit, full_audit_signs, np.arange(len(frame), dtype=int)
            )[:, local_relevant, :]
            targeted_values = project_values(fit, full_audit_signs, local_relevant)
            projection_error = float(np.max(np.abs(full_values - targeted_values)))
            if projection_error > 1e-9:
                raise RuntimeError("Targeted whole-donor projection failed audit")
            invariant_rows[-1]["targeted_projection_max_difference"] = projection_error
            donor_node_profiles[donor].extend(residual_profiles)

    donor_contrasts = np.empty((draws, len(DONORS)), dtype=float)
    component_arrays: dict[int, dict[str, np.ndarray]] = {}
    for donor_index, donor in enumerate(DONORS):
        if len(donor_node_profiles[donor]) != 3:
            raise RuntimeError(f"Wrong node-profile count for donor {donor}")
        profiles = np.stack(donor_node_profiles[donor], axis=1)
        contrast, components = topology_from_draw_profiles(profiles)
        donor_contrasts[:, donor_index] = contrast
        component_arrays[donor] = components
    if not np.isfinite(donor_contrasts).all():
        raise RuntimeError(f"Nonfinite topology in {model_id}")
    output = pd.DataFrame(
        {
            "draw_id": np.arange(1, draws + 1),
            "mean_adjacency_contrast": donor_contrasts.mean(axis=1),
            **{
                f"donor_{donor}_adjacency_contrast": donor_contrasts[:, index]
                for index, donor in enumerate(DONORS)
            },
        }
    )
    return output, qc_rows, invariant_rows


def wilson_interval(successes: int, total: int, z: float = 1.959963984540054) -> tuple[float, float]:
    proportion = successes / total
    denominator = 1 + z * z / total
    centre = (proportion + z * z / (2 * total)) / denominator
    half = z * math.sqrt(
        proportion * (1 - proportion) / total + z * z / (4 * total * total)
    ) / denominator
    return centre - half, centre + half


def summarize_model(frame: pd.DataFrame, observed: float) -> dict[str, Any]:
    values = frame["mean_adjacency_contrast"].to_numpy(float)
    donor_columns = [f"donor_{donor}_adjacency_contrast" for donor in DONORS]
    donor_values = frame[donor_columns].to_numpy(float)
    if not np.isfinite(values).all() or not np.isfinite(donor_values).all():
        raise RuntimeError("Model summary received nonfinite values")
    null_variance = float(values.var(ddof=1))
    unique_statistic_count = int(np.unique(values).size)
    donor_sd = donor_values.std(axis=0, ddof=1)
    if null_variance <= 0 or unique_statistic_count <= 1 or np.any(donor_sd <= 0):
        raise RuntimeError("Spatial null failed the frozen endpoint-diversity gate")
    exceedances = int(np.sum(values >= observed - TAIL_TOLERANCE))
    empirical_p = (1 + exceedances) / (1 + len(values))
    q95 = float(np.quantile(values, 0.95, method="linear"))
    wilson_low, wilson_high = wilson_interval(exceedances, len(values))
    passes = bool(empirical_p <= 0.05 and observed > q95)
    return {
        "observed": observed,
        "null_n": len(values),
        "null_mean": float(values.mean()),
        "null_sd_ddof1": math.sqrt(null_variance),
        "null_variance_ddof1": null_variance,
        "unique_statistic_count": unique_statistic_count,
        "donor_contrast_sd_ddof1": {
            str(donor): float(donor_sd[index]) for index, donor in enumerate(DONORS)
        },
        "null_q95_linear": q95,
        "tail": "upper",
        "tail_tolerance": TAIL_TOLERANCE,
        "exceedance_count": exceedances,
        "empirical_p_plus_one": empirical_p,
        "exceedance_fraction": exceedances / len(values),
        "exceedance_fraction_wilson95": [wilson_low, wilson_high],
        "monte_carlo_borderline_at_0_05": bool(wilson_low <= 0.05 <= wilson_high),
        "observed_beyond_q95": bool(observed > q95),
        "numeric_pass": passes,
    }


def lodo_table(
    model_id: str,
    frame: pd.DataFrame,
    observed_donor: pd.DataFrame,
) -> pd.DataFrame:
    rows = []
    for donor in DONORS:
        retained = [candidate for candidate in DONORS if candidate != donor]
        null_columns = [f"donor_{candidate}_adjacency_contrast" for candidate in retained]
        null_values = frame[null_columns].mean(axis=1).to_numpy(float)
        observed = float(
            observed_donor[
                observed_donor["donor_id"].isin(retained)
            ]["adjacency_contrast"].mean()
        )
        exceedances = int(np.sum(null_values >= observed - TAIL_TOLERANCE))
        p_value = (1 + exceedances) / (1 + len(null_values))
        q95 = float(np.quantile(null_values, 0.95, method="linear"))
        low, high = wilson_interval(exceedances, len(null_values))
        rows.append(
            {
                "model_id": model_id,
                "omitted_donor": donor,
                "observed": observed,
                "null_n": len(null_values),
                "null_q95_linear": q95,
                "exceedance_count": exceedances,
                "empirical_p_plus_one": p_value,
                "exceedance_fraction_wilson95_low": low,
                "exceedance_fraction_wilson95_high": high,
                "observed_positive": observed > 0,
                "observed_beyond_q95": observed > q95,
                "status": "OVERLAPPING_DIAGNOSTIC_ONLY",
            }
        )
    return pd.DataFrame(rows)


def write_environment(path: Path) -> None:
    lines = [
        f"python={platform.python_version()}",
        f"numpy={np.__version__}",
        f"pandas={pd.__version__}",
        f"scipy={scipy.__version__}",
        f"platform={platform.platform()}",
    ]
    path.write_text("\n".join(lines) + "\n", encoding="utf-8")


def verify_ledger_after(ledger: pd.DataFrame) -> pd.DataFrame:
    after = []
    for row in ledger.itertuples(index=False):
        path = PROJECT / row.path
        ok = path.is_file() and path.stat().st_size == int(row.bytes) and sha256(path) == row.sha256
        after.append(bool(ok))
    result = ledger.copy()
    result["verified_after_endpoint"] = after
    if not result["verified_after_endpoint"].all():
        raise RuntimeError("An input changed during endpoint execution")
    return result


def main() -> int:
    if OUT.exists():
        raise RuntimeError(f"Refusing to overwrite existing final directory: {OUT}")
    if STAGING.exists():
        raise RuntimeError(f"Staging directory already exists: {STAGING}")
    protocol = json.loads(PROTOCOL_PATH.read_text(encoding="utf-8"))
    amendment1 = json.loads(AMENDMENT1_PATH.read_text(encoding="utf-8"))
    amendment2 = json.loads(AMENDMENT2_PATH.read_text(encoding="utf-8"))
    ledger = verify_static_inputs(protocol, amendment1, amendment2)
    metadata = pd.read_csv(V08 / "analysis" / "ahba_sample_metadata_v0_8.csv")
    validate_metadata(metadata)
    expression, ledger = load_expression_maps(metadata, ledger)
    observed = observed_tensor(metadata, expression)
    residual_difference, observed_donor, endpoint_difference = validate_observed_replay(
        observed
    )
    observed_statistic = float(observed_donor["adjacency_contrast"].mean())

    STAGING.mkdir(parents=True)
    model_frames: dict[str, pd.DataFrame] = {}
    summaries: dict[str, dict[str, Any]] = {}
    all_qc: list[dict[str, Any]] = []
    all_invariants: list[dict[str, Any]] = []
    for spec in MODEL_SPECS:
        print(f"Running {spec['model_id']} ({spec['draws']} draws)", flush=True)
        frame, qc_rows, invariant_rows = run_model(spec, metadata, expression)
        model_frames[spec["model_id"]] = frame
        summaries[spec["model_id"]] = summarize_model(frame, observed_statistic)
        all_qc.extend(qc_rows)
        all_invariants.extend(invariant_rows)
        filename = f"spatial_null_{spec['model_id']}_v1_1.csv.gz"
        write_csv_gzip(STAGING / filename, frame)

    p1a = summaries["P1A_BLOCKWISE_KNN5_TREND_PRESERVED"]
    p1b = summaries["P1B_WHOLE_DONOR_KNN6_FIXED_EFFECTS"]
    if p1a["numeric_pass"] and p1b["numeric_pass"]:
        classification = "SPATIAL_ROBUST_DOUBLE_GATE"
    elif p1a["numeric_pass"]:
        classification = "BLOCK_ONLY_SPATIAL_ROBUSTNESS"
    elif p1b["numeric_pass"]:
        classification = "WHOLE_ONLY_SPATIAL_ROBUSTNESS"
    else:
        classification = "SPATIAL_NOT_ROBUST"
    iut_p = max(p1a["empirical_p_plus_one"], p1b["empirical_p_plus_one"])
    all_models_numeric_pass = all(item["numeric_pass"] for item in summaries.values())
    all_models_nonborderline = all(
        item["exceedance_fraction_wilson95"][1] < 0.05
        for item in summaries.values()
    )
    if p1a["numeric_pass"] and p1b["numeric_pass"]:
        if all_models_numeric_pass and all_models_nonborderline:
            model_stability_classification = (
                "SPATIAL_ROBUST_ACROSS_CHECKED_MODELS_NONBORDERLINE"
            )
        else:
            model_stability_classification = (
                "SPATIAL_ROBUST_DOUBLE_GATE_MODEL_SENSITIVE_OR_BORDERLINE"
            )
    else:
        model_stability_classification = "NOT_APPLICABLE_PRIMARY_DOUBLE_GATE_NOT_PASSED"

    lodo = pd.concat(
        [
            lodo_table(
                model_id,
                model_frames[model_id],
                observed_donor,
            )
            for model_id in [
                "P1A_BLOCKWISE_KNN5_TREND_PRESERVED",
                "P1B_WHOLE_DONOR_KNN6_FIXED_EFFECTS",
            ]
        ],
        ignore_index=True,
    )
    write_csv(STAGING / "observed_topology_donor_v1_1.csv", observed_donor)
    write_csv(STAGING / "spatial_block_qc_v1_1.csv", pd.DataFrame(all_qc))
    write_csv(STAGING / "spatial_invariant_audit_v1_1.csv", pd.DataFrame(all_invariants))
    write_csv(STAGING / "spatial_lodo_v1_1.csv", lodo)

    summary = {
        "version": "v1.1",
        "execution_revision": "v1.1-A02_ENDPOINT_SEAL",
        "protocol_sha256": sha256(PROTOCOL_PATH),
        "amendment_A01_sha256": sha256(AMENDMENT1_PATH),
        "amendment_A02_sha256": sha256(AMENDMENT2_PATH),
        "endpoint_script_sha256": sha256(Path(__file__).resolve()),
        "observed": {
            "mean_adjacency_contrast": observed_statistic,
            "maximum_residual_replay_difference_vs_v0_10": residual_difference,
            "maximum_endpoint_replay_difference_vs_v1_0": endpoint_difference,
        },
        "models": summaries,
        "primary_double_gate": {
            "component_A": "P1A_BLOCKWISE_KNN5_TREND_PRESERVED",
            "component_B": "P1B_WHOLE_DONOR_KNN6_FIXED_EFFECTS",
            "intersection_union_p_max": iut_p,
            "both_observed_beyond_q95": bool(
                p1a["observed_beyond_q95"] and p1b["observed_beyond_q95"]
            ),
            "both_numeric_pass": bool(p1a["numeric_pass"] and p1b["numeric_pass"]),
        },
        "classification": classification,
        "model_stability_classification": model_stability_classification,
        "all_six_models_numeric_pass": all_models_numeric_pass,
        "all_six_models_wilson_upper_below_0_05": all_models_nonborderline,
        "interpretation_status": "POST_DISCOVERY_SPATIAL_ROBUSTNESS_NOT_INDEPENDENT_CONFIRMATION",
        "implementation_note": "Custom float64 nuisance-restricted joint singleton MSR adapted from Wagner and Dray (2015); not a bitwise reproduction of BrainSpace.",
        "claim_guardrail": "No connectivity, direction, molecular flow, causality, route exclusivity, pain/fibromyalgia mechanism, stimulation target, or independent replication is inferred.",
        "marginal_limitation": "Joint singleton MSR preserves fitted trends, residual Moran spectral power and residual gene covariance, but not higher-order univariate marginals.",
    }
    write_json(STAGING / "summary_v1_1.json", summary)
    write_environment(STAGING / "environment_versions_spatial_v1_1.txt")
    ledger = verify_ledger_after(ledger)
    write_csv(STAGING / "verified_spatial_inputs_v1_1.csv", ledger)

    output_files = sorted(path for path in STAGING.iterdir() if path.is_file())
    manifest_lines = [f"{sha256(path)}  {path.name}" for path in output_files]
    (STAGING / "spatial_analysis_hashes_v1_1.txt").write_text(
        "\n".join(manifest_lines) + "\n", encoding="utf-8"
    )
    os.replace(STAGING, OUT)
    print(json.dumps(summary, indent=2, sort_keys=True), flush=True)
    return 0


if __name__ == "__main__":
    try:
        raise SystemExit(main())
    except Exception:
        # Preserve staging for audit; never promote it after a failure.
        raise
```
