## Supplementary Archive S1 for "An anatomy-constrained transcriptomic screen identifies a 19-gene panel with an ordered regional similarity pattern in the human somatosensory system": figure_S_gse3526_donor_heterogeneity_caption.md

**Supplementary Figure S1. Donor heterogeneity in the public GSE3526 source-lineage reconstruction.** (A) Three-node medulla–thalamus–parietal-lobe relative serial-order contrast and (B) four-node dorsal-root-ganglion–medulla–thalamus–parietal-lobe adjacency-minus-nonadjacency contrast for the eight donors with complete broad-region coverage. Each donor is shown under median, mean, and historically selected delegate probe-set summaries. Recorded sex and age are public GEO series metadata. The three-node contrast separated by recorded sex in this small post hoc comparison, whereas the four-node contrast did not show the same pattern. This observation is descriptive and cannot establish a sex effect because each stratum contains four donors and sex may be confounded by age and unavailable cause-of-death, postmortem-interval, RNA-integrity, or processing metadata. (C) Donor-specific pairwise Spearman correlations underlying the median-collapse three-node statistic. The mean relative-order contrast is positive because the two selected adjacent pairs are less dissimilar than the nonadjacent medulla–parietal pair; it does not imply uniformly positive coexpression. GSE3526 is a public cohort in the documented historical source lineage of BIGE, but formal sample- and processing-level reconciliation has not established numerical equivalence to the internally analyzed aggregate. Accordingly, all panels are source-lineage, post-selection sensitivities rather than independent replication or confirmatory inference.

Source: NCBI GEO GSE3526; Roth et al., *Neurogenetics* 2006, doi:10.1007/s10048-006-0032-6. The plotted data are in `figure_S_gse3526_donor_heterogeneity_source_data.csv`.
