## Supplementary Archive S1 for "An anatomy-constrained transcriptomic screen identifies a 19-gene panel with an ordered regional similarity pattern in the human somatosensory system": figure_S_gse3526_donor_heterogeneity_v1.pdf

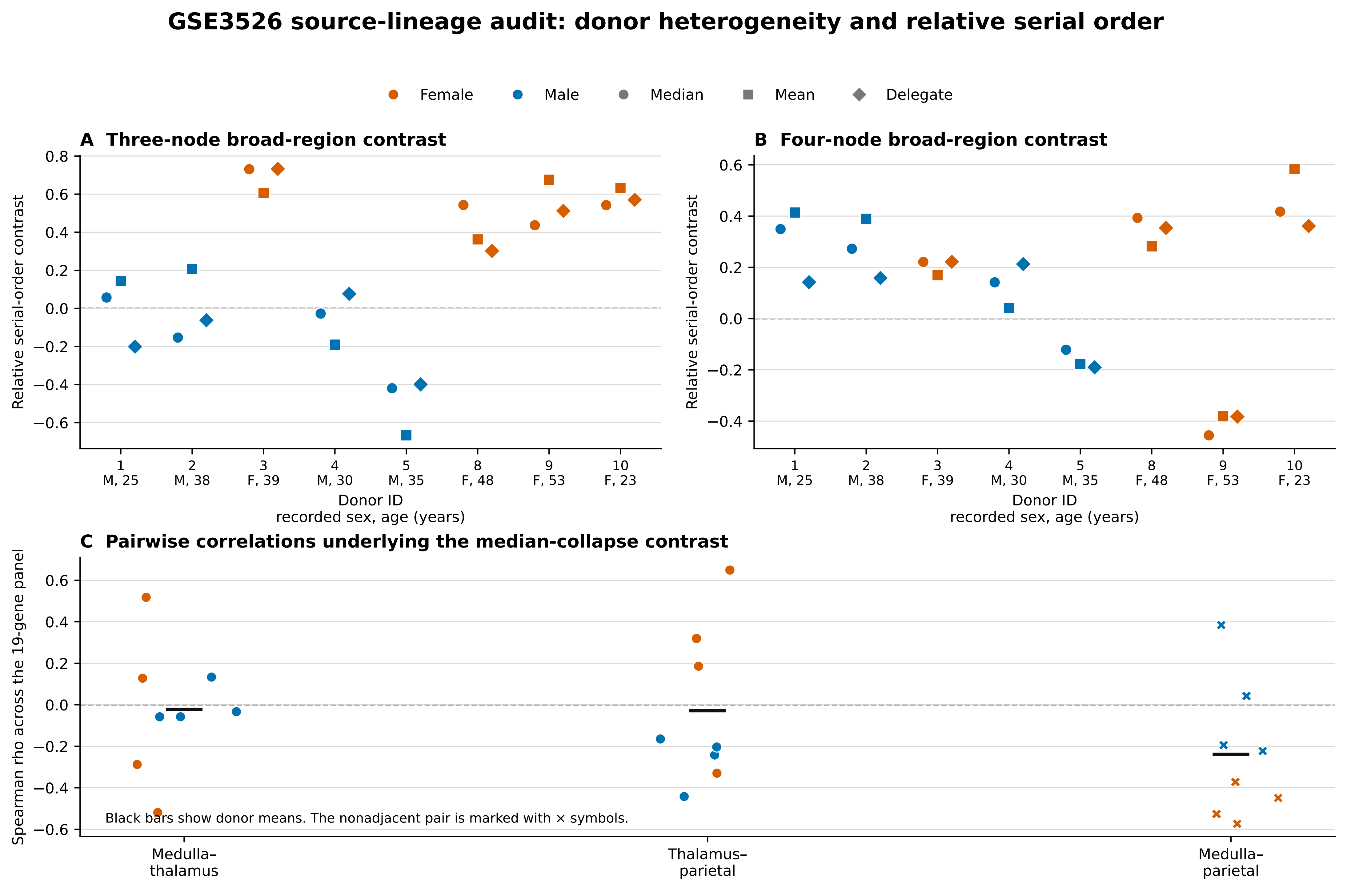

Descriptive post-selection analysis of eight complete donors. Recorded sex was not a prespecified hypothesis and cannot be interpreted causally.
