## Supplementary Archive S1 for "An anatomy-constrained transcriptomic screen identifies a 19-gene panel with an ordered regional similarity pattern in the human somatosensory system": plot_gse3526_donor_heterogeneity.py

```
#!/usr/bin/env python3
"""Create a publication-style descriptive figure for the GSE3526 source-lineage audit."""

from __future__ import annotations

import argparse
from pathlib import Path

import matplotlib.pyplot as plt
from matplotlib.lines import Line2D
import numpy as np
import pandas as pd


DONOR_METADATA = {
    1: ("Male", 25),
    2: ("Male", 38),
    3: ("Female", 39),
    4: ("Male", 30),
    5: ("Male", 35),
    8: ("Female", 48),
    9: ("Female", 53),
    10: ("Female", 23),
}

SEX_COLOR = {"Male": "#0072B2", "Female": "#D55E00"}
METHOD_STYLE = {
    "median": ("o", -0.20, "Median"),
    "mean": ("s", 0.00, "Mean"),
    "delegate": ("D", 0.20, "Delegate"),
}


def parse_args() -> argparse.Namespace:
    here = Path(__file__).resolve().parent
    default_analysis = here.parent / "gse3526" / "unpacked" / "analysis"
    parser = argparse.ArgumentParser(description=__doc__)
    parser.add_argument("--analysis", type=Path, default=default_analysis)
    parser.add_argument("--output", type=Path, default=here)
    return parser.parse_args()


def add_metadata(frame: pd.DataFrame) -> pd.DataFrame:
    result = frame.copy()
    result["recorded_sex"] = result["donor_id"].map(lambda x: DONOR_METADATA[int(x)][0])
    result["age_years"] = result["donor_id"].map(lambda x: DONOR_METADATA[int(x)][1])
    return result


def donor_panel(ax: plt.Axes, frame: pd.DataFrame, value: str, title: str) -> None:
    donors = sorted(frame["donor_id"].astype(int).unique())
    donor_position = {donor: position for position, donor in enumerate(donors)}
    for method, (marker, offset, _label) in METHOD_STYLE.items():
        subset = frame.loc[frame["collapse_method"].eq(method)].sort_values("donor_id")
        for row in subset.itertuples(index=False):
            donor = int(row.donor_id)
            sex, _age = DONOR_METADATA[donor]
            ax.scatter(
                donor_position[donor] + offset,
                getattr(row, value),
                marker=marker,
                s=48,
                facecolor=SEX_COLOR[sex],
                edgecolor="white",
                linewidth=0.7,
                zorder=3,
            )
    ax.axhline(0, color="#333333", linewidth=0.9, linestyle="--", zorder=1)
    ax.set_xticks(range(len(donors)))
    ax.set_xticklabels(
        [f"{d}\n{DONOR_METADATA[d][0][0]}, {DONOR_METADATA[d][1]}" for d in donors],
        fontsize=8,
    )
    ax.set_xlabel("Donor ID\nrecorded sex, age (years)")
    ax.set_ylabel("Relative serial-order contrast")
    ax.set_title(title, loc="left", fontweight="bold")
    ax.grid(axis="y", color="#D9D9D9", linewidth=0.6)


def correlation_panel(ax: plt.Axes, three: pd.DataFrame) -> None:
    median = three.loc[three["collapse_method"].eq("median")].copy()
    pairs = [
        ("Medulla–\nthalamus", "rho_medulla_thalamus", True),
        ("Thalamus–\nparietal", "rho_thalamus_parietal", True),
        ("Medulla–\nparietal", "rho_medulla_parietal", False),
    ]
    rng = np.random.default_rng(20260909)
    for position, (label, column, adjacent) in enumerate(pairs):
        values = median[column].to_numpy(dtype=float)
        jitter = rng.uniform(-0.10, 0.10, len(values))
        colors = [SEX_COLOR[DONOR_METADATA[int(d)][0]] for d in median["donor_id"]]
        ax.scatter(
            np.full(len(values), position) + jitter,
            values,
            s=38,
            c=colors,
            marker="o" if adjacent else "X",
            edgecolor="white",
            linewidth=0.6,
            zorder=3,
        )
        ax.scatter(
            position,
            values.mean(),
            marker="_",
            s=500,
            color="#111111",
            linewidth=2.0,
            zorder=4,
        )
    ax.axhline(0, color="#333333", linewidth=0.9, linestyle="--", zorder=1)
    ax.set_xticks(range(len(pairs)))
    ax.set_xticklabels([item[0] for item in pairs])
    ax.set_ylabel("Spearman rho across the 19-gene panel")
    ax.set_title("C  Pairwise correlations underlying the median-collapse contrast", loc="left", fontweight="bold")
    ax.grid(axis="y", color="#D9D9D9", linewidth=0.6)
    ax.text(
        0.02,
        0.04,
        "Black bars show donor means. The nonadjacent pair is marked with × symbols.",
        transform=ax.transAxes,
        fontsize=8,
        va="bottom",
    )


def main() -> int:
    args = parse_args()
    args.output.mkdir(parents=True, exist_ok=True)
    three = add_metadata(pd.read_csv(args.analysis / "donor_topology_three_node.csv"))
    four = add_metadata(pd.read_csv(args.analysis / "donor_topology_four_node.csv"))

    three["analysis"] = "three_node"
    four["analysis"] = "four_node"
    figure_source = pd.concat(
        [
            three[
                [
                    "analysis",
                    "collapse_method",
                    "donor_id",
                    "recorded_sex",
                    "age_years",
                    "anatomical_contrast",
                    "rho_medulla_thalamus",
                    "rho_thalamus_parietal",
                    "rho_medulla_parietal",
                ]
            ].rename(columns={"anatomical_contrast": "serial_order_contrast"}),
            four[
                [
                    "analysis",
                    "collapse_method",
                    "donor_id",
                    "recorded_sex",
                    "age_years",
                    "anatomical_adjacency_minus_nonadjacency",
                    "rho_medulla_thalamus",
                    "rho_thalamus_parietal",
                    "rho_medulla_parietal",
                ]
            ].rename(
                columns={"anatomical_adjacency_minus_nonadjacency": "serial_order_contrast"}
            ),
        ],
        ignore_index=True,
    )
    figure_source.to_csv(
        args.output / "figure_S_gse3526_donor_heterogeneity_source_data.csv",
        index=False,
        lineterminator="\n",
        float_format="%.12g",
    )

    plt.rcParams.update(
        {
            "font.family": "DejaVu Sans",
            "font.size": 9,
            "axes.spines.top": False,
            "axes.spines.right": False,
            "axes.linewidth": 0.8,
        }
    )
    fig = plt.figure(figsize=(11.8, 8.0))
    grid = fig.add_gridspec(2, 2, height_ratios=[1.05, 1.0])
    ax_a = fig.add_subplot(grid[0, 0])
    ax_b = fig.add_subplot(grid[0, 1])
    ax_c = fig.add_subplot(grid[1, :])

    donor_panel(ax_a, three, "anatomical_contrast", "A  Three-node broad-region contrast")
    donor_panel(
        ax_b,
        four,
        "anatomical_adjacency_minus_nonadjacency",
        "B  Four-node broad-region contrast",
    )
    correlation_panel(ax_c, three)

    sex_handles = [
        Line2D(
            [0],
            [0],
            marker="o",
            linestyle="none",
            markerfacecolor=SEX_COLOR[sex],
            markeredgecolor="white",
            markersize=7,
            label=sex,
        )
        for sex in ["Female", "Male"]
    ]
    method_handles = [
        Line2D(
            [0],
            [0],
            marker=marker,
            linestyle="none",
            color="#555555",
            markerfacecolor="#777777",
            markeredgecolor="white",
            markersize=7,
            label=label,
        )
        for marker, _offset, label in METHOD_STYLE.values()
    ]
    fig.legend(
        handles=sex_handles + method_handles,
        loc="upper center",
        bbox_to_anchor=(0.5, 0.905),
        ncol=5,
        frameon=False,
    )
    fig.suptitle(
        "GSE3526 source-lineage audit: donor heterogeneity and relative serial order",
        fontsize=14,
        fontweight="bold",
        y=0.972,
    )
    fig.text(
        0.5,
        0.018,
        "Descriptive post-selection analysis of eight complete donors. Recorded sex was not a prespecified hypothesis and cannot be interpreted causally.",
        ha="center",
        fontsize=8,
        color="#444444",
    )
    fig.subplots_adjust(left=0.08, right=0.98, bottom=0.10, top=0.82, wspace=0.16, hspace=0.38)

    stem = args.output / "figure_S_gse3526_donor_heterogeneity_v1"
    fig.savefig(stem.with_suffix(".png"), dpi=320, bbox_inches="tight")
    fig.savefig(stem.with_suffix(".pdf"), bbox_inches="tight")
    fig.savefig(stem.with_suffix(".svg"), bbox_inches="tight")
    plt.close(fig)
    return 0


if __name__ == "__main__":
    raise SystemExit(main())
```
