## Supplementary Archive S1 for "An anatomy-constrained transcriptomic screen identifies a 19-gene panel with an ordered regional similarity pattern in the human somatosensory system": Manuscript_draft_v0_3.md

# Selection-aware and spatial-null evaluation of a transcriptomic adjacency contrast across human somatosensory relay-region proxies

**Working manuscript — v0.3**  
**Analysis version:** v1.1-A04; documentary release A05  
**Authors and affiliations:** [To be completed]  
**Corresponding author:** [To be completed]

## Abstract

Regional transcriptomic similarity can appear pathway-like because of adaptive gene selection, shared cellular composition, probe choice, and spatial autocorrelation. We evaluated a frozen 19-gene panel selected in a dorsal column–medial lemniscus (DCML)-constrained screen across three Allen Human Brain Atlas (AHBA) relay-region proxies: cuneate/gracile nuclei, the ventral division of the dorsal thalamus aggregate (DTLv), and postcentral gyrus. The donor-level statistic contrasted the mean Spearman similarity of adjacent regions against endpoint similarity. The observed six-donor mean was 0.3451754386; five of six donor contrasts were positive, although the one-sided sign test was not significant ($P=0.109375$). In post-discovery analyses, the observation was evaluated against two 50,000-draw selection-aware conditional pseudoroute nulls and met both frozen criteria: 1,395 technical-null panels equalled or exceeded it ($P=0.02791944$), as did 1,328 neuron-conditioned panels ($P=0.02657947$). We then applied nuisance-restricted joint Moran spectral randomization to sample-level AHBA maps. The prespecified spatial test was a conjunction of a donor-by-compartment 5-nearest-neighbour model (v1.1-P1A) and a whole-donor 6-nearest-neighbour model with top-level anatomical fixed effects (v1.1-P1B). v1.1-P1A did not pass (2,558/50,000 exceedances; $P=0.05117898$; observed below q95=0.34590643), whereas v1.1-P1B passed (192/50,000; $P=0.00385992$; q95=0.22149854). Thus, the intersection-union probability was 0.05117898 and the double gate failed. Four dense-distance or axis-standardized sensitivities met their nominal, unadjusted individual criteria ($P=0.04319568$, 0.00509949, 0.04399560, and 0.00449955); these sensitivities had no familywise inferential or primary decision role. Leave-one-donor-out results remained positive but differed by model. A public donor-level reconstruction from GSE3526, a cohort in the documented source lineage of the historical BIGE resource, produced positive mean relative serial-order contrasts across eight complete donors (five or six positive donors, depending on probe collapse), but most three-node pairwise correlations were negative and the descriptive tail fraction depended on calculation (minimum 0.0487755). Formal sample- and processing-level reconciliation with the internally analyzed BIGE aggregate has not been completed. This post-selection, nonexchangeable-label, source-lineage sensitivity has no confirmatory interpretation and is not an independent replication. The evidence supports a post-discovery adjacency contrast under conditional selection calibrations and several spatial specifications, but the prespecified spatial conjunction failed; support is model-dependent and not independently confirmed. It does not establish connectivity, direction, route exclusivity, causality, a fibromyalgia or pain mechanism, or a stimulation target.

**Keywords:** Allen Human Brain Atlas; imaging transcriptomics; Moran spectral randomization; somatosensory relays; spatial autocorrelation; conditional null models

## Introduction

Spatially resolved transcriptomic atlases make it possible to ask whether anatomically related regions share molecular organization. Such analyses are difficult to interpret, however, because nearby samples tend to have similar expression, broad anatomical compartments differ in cellular composition, and gene panels are often selected after multiple data-dependent steps. A pathway-shaped pattern can therefore arise without pathway-specific biology. Spatial autocorrelation and the selection process must be represented explicitly in the null model rather than treated as secondary caveats [1–5].

The present study originated in an anatomy-constrained screen of regional transcriptomic patterns. The initial hypothesis—continuous expression across complete candidate routes—was not supported. A subsequent analysis identified a topology for 19 genes selected in a DCML-constrained screen: the adjacency contrast was positive in five of six donors, while both adjacent correlations were positive in all six; the donor-level sign test for the contrast was not significant. Residual expression profiles were, on average, more similar between adjacent sampled relay-region proxies than between the two endpoint regions. This observation was generated after inspection of the underlying data and is treated here as a post-discovery signal, not a prospectively confirmed hypothesis.

We evaluated two distinct alternative explanations. First, selection-aware conditional nulls asked whether the 19-gene result exceeded panels obtainable through analogous anatomical and technical selection rules. Second, joint spatial surrogate models asked whether the topology exceeded patterns expected after preserving fitted spatial trends, residual spatial spectral power, and cross-gene covariance. Because the design spans brainstem, thalamus, and cortex, a cortical surface rotation was not appropriate; spatial tests were instead defined over irregularly sampled AHBA coordinates.

Our aim was not to infer anatomical connectivity or molecular transmission. We asked a narrower question: does the observed adjacent-relay transcriptomic topology remain unusual under explicit models of selection and space, and how stable is that conclusion across defensible spatial formulations?

## Methods

### Study design and inferential status

The analysis sequence comprised discovery in **an internally maintained, processed extract of the Body Index of Gene Expression (BIGE)** followed by AHBA evaluation. BIGE has been described as a body-wide Affymetrix Human Genome U133 Plus 2.0 microarray resource spanning more than 105 human cells and tissues [13]. The processed extract used here contained regional means for 10,262 genes rather than individual-level or donor-level observations. Its documented source lineage includes the postmortem-tissue quality analysis of Lee et al. [11], the 353-array survey of 65 normal human tissues reported by Roth et al. and deposited as GSE3526 [12], and the later published description of BIGE by Ushach et al. [13]. Five anatomical routes were frozen, of which three primary routes were evaluated in this extract. Twenty-three candidates emerged: 19 from the DCML-constrained screen, three from descending pain modulation, and one from basal ganglia. The route-wide continuity hypothesis was negative. The present analysis concerns only the subsequently identified topology of the fixed 19-gene panel and is scientifically separate from fibromyalgia and stimulation hypotheses.

Version 1.0 was designed after the topology had been observed in earlier AHBA analyses and tested selection-conditioned and diagnostic alternatives. Version 1.1 was likewise designed after the AHBA observation was known and evaluated spatial autocorrelation. Protocols, amendments, executable hashes, random seeds, sample rules, endpoints, tails, and decision criteria were sealed before the corresponding new endpoint was generated. These procedures reduce analytic flexibility but do not make either version an independent confirmation or an external preregistration.

### Gene panel

The fixed panel contained ATP2C1, RAI2, RAB3GAP2, CCT6A, MAP1B, SH3BGRL2, PDHX, C7orf25, TSHZ2, NEFM, ATXN3, ADAM23, HMGCS1, SLC16A6, NANS, IFT57, MEF2A, BBS7, and RAP1GDS1. Three genes had the frozen selection tier `B_ROUTEWIDE_90` and 16 had `C_BALANCED`. Disease annotations were not used to select this panel.

### Public GSE3526 source-lineage donor reconstruction

We reconstructed donor-level profiles from the public GSE3526 series matrix, which contains 353 Affymetrix HG-U133 Plus 2.0 arrays from 65 normal tissue types (20 central-nervous-system and 45 non-CNS tissues) across ten donors [12]. GSE3526 is a documented public component in the historical source lineage of BIGE, but formal reconciliation has not established numerical equivalence between the GSE3526-derived aggregate and the internally analyzed BIGE extract. It can recover donor heterogeneity within that source lineage but cannot serve as independent replication or external validation. The official matrix contained linear-scale RMA intensities (range 4.538–33,730.54), so values were transformed as $\log_2(x+1)$; replacing this with $\log_2(x)$ changed the endpoint only minimally. The reconstruction used 39 GPL570 probe sets representing the frozen 19-gene panel. Probes were collapsed separately by median, mean, and the historically selected delegate probe, and each gene was standardized within donor across the 21–23 tissues available to that donor from the frozen 23-tissue neural universe. The series contained 193 neural arrays across all ten donors; 180 belonged to the eight donors with complete endpoints and could influence their within-donor standardized profiles. The endpoints themselves used 24 arrays for the three-node analysis or 32 for the four-node analysis. No donor–tissue duplicates were present. Eight donors (1, 2, 3, 4, 5, 8, 9, and 10) had complete broad medulla, thalamus, parietal-lobe, and dorsal-root-ganglion (DRG) observations and constituted the donor-level analytical units ($n=8$). These broad tissues are not equivalent to the AHBA Cu/Gr, DTLv, and postcentral-gyrus proxies; effect magnitudes across the two resources are therefore not directly comparable and cannot establish relay- or route-specificity.

The three-node analysis applied the adjacency contrast to medulla–thalamus–parietal-lobe profiles. The four-node analysis added DRG and contrasted the mean of the three anatomically adjacent correlations with the mean of the three nonadjacent correlations. Three-node middle-label assignments were enumerated exactly ($3^8=6{,}561$); four-node orders were sampled in 100,000 donor-stratified permutations for each probe-collapse method; and all $2^8$ sign assignments were enumerated. A deterministic convolution over the finite per-donor four-node label distributions then calculated the exact tail counts across $24^8$ joint assignments. Because the gene panel, tissues, and source lineage were already historically linked to discovery and anatomical labels are not exchangeable, these outputs were defined as post-selection descriptive tail fractions, not confirmatory $P$ values.

Background dependence was assessed post hoc by replacing the donor-specific available neural background with the 20 neural tissues common to the eight complete donors, all tissues available to each donor from the 65-tissue universe (29–58 tissues per donor), or no within-donor gene standardization. Recorded-sex heterogeneity was also inspected post hoc by comparing the four female and four male donors; complete rank separation was summarized by the two-sided partition fraction $2/\binom{8}{4}$. To assess whether this separation was driven by one gene, we repeated the three-node female-minus-male contrast after removing each gene in turn, after removing X-linked RAI2, and after jointly removing RAI2 and C7orf25. These are heterogeneity diagnostics rather than tests of a sex effect.

Review of the NCBI GPL570 platform annotation accessed on 9 September 2026 identified an unresolved probe-specificity issue [15]: all three probe sets assigned in the frozen map to RAB3GAP2 are also annotated to AURKAPS1, and the sole C7orf25 probe set is also annotated to PSMA2. The primary reconstruction retained the historical map for provenance. A labelled post hoc sensitivity excluded both historically assigned genes, leaving 17 genes; contemporary CEL-level reprocessing with an updated custom definition file remains prospective.

### AHBA expression data and regional residuals

Adult-human microarray data from six AHBA donors were analyzed [1]. The source comprised 3,702 samples. Each whole-donor model used all 2,797 eligible left-hemisphere or unpaired samples; the 18 donor-by-compartment models used the 540 samples belonging to myelencephalon, dorsal thalamus, or parietal lobe. Forty-four exact-symbol HumanMA DNA probes represented the 19 genes. Allen-provided probe z-score vectors were parsed, aligned to frozen sample order, and collapsed by the median across probes for each gene. To reproduce the historical v0.10 lineage, parsing, alignment, probe medians, and the observed regional median subtraction were evaluated in `float32`; spatial regression and surrogate generation used the same collapsed maps converted to `float64`.

The three target proxies were cuneate/gracile nuclei (Cu/Gr; 19 target samples), the ventral division of the dorsal thalamus aggregate (DTLv; 20 target samples), and postcentral gyrus (62 target samples). DTLv was treated as the sampled somatosensory-thalamic proxy and not as an isolated ventral posterolateral nucleus (VPL) measurement. Within each donor, region, and gene, the node residual was the median expression in target samples minus the median in non-target sibling samples from the same top-level macrocompartment. Target counts were unequal across donors; notably, donor 12876 had one Cu/Gr target and 15 sibling samples.

### Topology statistic

For donor $d$, Spearman correlations across the 19 genes were calculated for the three pairs of regional residual profiles. The frozen adjacency-contrast statistic was

$$
T_d=\frac{\rho_{d,\mathrm{Cu/Gr,DTLv}}+\rho_{d,\mathrm{DTLv,Postcentral}}}{2}
-\rho_{d,\mathrm{Cu/Gr,Postcentral}}.
$$

The primary observed statistic was the arithmetic mean of $T_d$ across all six donors. Correlations in the observed and null data used the same average-rank and Pearson-on-ranks implementation.

### Selection-aware conditional nulls (v1.0)

All 8,854 non-DCML four-region combinations in the frozen 23-region BIGE-derived regional extract were enumerated, and the original detection, dominance, threshold, and tier-precedence rules were reapplied. All 23 original candidates were excluded from control sampling. A common universe of 742 controls with complete AHBA residual and broad-region neuronal-contrast data was used.

The v1.0-P1 technical null matched panel size and the exact tier-by-AHBA-probe-count composition, together with eight expression and technical variables from the BIGE-derived extract, Human Protein Atlas (HPA) [14], and AHBA. The v1.0-S1 null additionally matched neuronal-minus-non-neuronal contrasts in medulla oblongata, thalamus, and cerebral cortex. One hundred thirty-four pseudoroutes had enough controls in all six required strata. Each null comprised 50,000 accepted 19-gene panels sampled through an eligible pseudoroute. These models condition on important parts of the selection mechanism; they do not reproduce the entire adaptive history and do not yield an unconditional selection probability.

Additional v1.0 analyses evaluated broad-region cell attribution, geometry-matched anatomical triplets, HPA exact-node residuals, random choice of one probe per gene, a gene-by-donor bootstrap, component decomposition, and leave-one-donor-out (LODO) calibration. Their prespecified or post hoc roles were retained in interpretation.

Before v1.0, v0.11 had tested the minimum across medulla-oblongata, thalamic, and cortical neuronal-minus-non-neuronal contrasts in the adult Human Brain Cell Atlas single-nucleus dataset [6]. Its technically matched 50,000-gene-set null gave 30 exceedances and a plus-one empirical $P=0.000620$. v1.0-S2 evaluated the same observed minimum against the more restrictive v1.0-P1 pseudoroute panels, thereby additionally conditioning on the route-selection mechanism. These tests answer different questions and were reported together.

### Joint spatial surrogate design (v1.1)

Spatial nulls used a custom `float64` nuisance-restricted joint singleton Moran spectral randomization (MSR) adapted from Wagner and Dray [2]. This implementation was not intended as a bitwise reproduction of BrainSpace or BrainSMASH [3,4,7].

For a fitted sample block with $n\times19$ expression matrix $X$ and full-rank nuisance design $Z$, ordinary least squares gave fitted values $ZB$ and residual matrix $R=X-ZB$. Let $Q$ be an orthonormal basis for the null space of $Z'$, and let $V=QE$, where $E$ contains the eigenvectors of $Q'WQ$ for spatial-weight matrix $W$. All $n-\mathrm{rank}(Z)$ modes were retained. For each surrogate, a single vector of spectral signs was shared across all 19 genes and the map was reconstructed as

$$
X^*=ZB+V(SV'R),
$$

where $S$ is diagonal with entries in $\{-1,+1\}$. Joint signs preserve the complete residual Moran spectral power and residual gene covariance within the fitted block, up to numerical tolerance. The fitted nuisance component was identical in every surrogate. Higher-order univariate marginal distributions were not exactly preserved.

The primary spatial endpoint was a conservative intersection-union double gate:

- **v1.1-P1A (`P1A_BLOCKWISE_KNN5_TREND_PRESERVED`):** 18 donor-by-macrocompartment blocks, each containing the target and sibling samples in myelencephalon, dorsal thalamus, or parietal lobe; symmetric union 5-nearest-neighbour graph with inverse native-coordinate Euclidean-distance weights; nuisance design comprising intercept and native MRI $x,y,z$ coordinates; 50,000 draws; root seed 20260912.
- **v1.1-P1B (`P1B_WHOLE_DONOR_KNN6_FIXED_EFFECTS`):** six whole-donor blocks containing all eligible samples; symmetric union 6-nearest-neighbour graph with inverse native-coordinate distance weights; nuisance design comprising intercept, native $x,y,z$, and top-level-structure fixed effects; 50,000 draws; root seed 20260913.

The Allen API sample MRI coordinates [8] were interpreted as locations in each donor's individual MR volume in millimetres, not MNI coordinates. Nearest-neighbour ties were resolved deterministically by distance and frozen sample index. Graphs were symmetrized by union, were not row-normalized, and were required to be connected.

Four v1.1 sensitivity models were frozen. v1.1-S1 used dense inverse-distance weights within the 18 blocks (10,000 draws; seed 20260914). v1.1-S2 used dense weights within whole donors (10,000; seed 20260915). v1.1-S3A repeated the blockwise 5-NN model after donor-wise standardization of each coordinate axis (10,000; seed 20260916). v1.1-S3B used axis-standardized whole-donor coordinates and an 8-NN graph (10,000; seed 20260917). The pre-endpoint coordinate-only connectivity check showed that $k=8$ was the smallest common value connecting all v1.1-S3B donor graphs.

### Statistical criteria

All primary and sensitivity spatial tests used the upper tail. With $b$ surrogate statistics greater than or equal to the observed statistic among $N$ draws, the empirical probability was $(1+b)/(1+N)$. The 95th percentile used linear interpolation. A component passed numerically only if $P\leq0.05$ and the observation strictly exceeded q95. A two-sided 95% Wilson interval for $b/N$ flagged Monte Carlo-borderline results when it contained 0.05.

The primary double gate passed only if both v1.1-P1A and v1.1-P1B passed all technical and numerical criteria. Its intersection-union probability was $\max(P_{P1A},P_{P1B})$. Sensitivities did not replace either co-primary component. LODO analyses reused draw-wise v1.1-P1A and v1.1-P1B donor contrasts and were overlapping diagnostics with no independent decision role.

### Reproducibility and quality control

All protocols, amendments, executables, inputs, and outputs were linked by SHA-256. Spatial fits were required to have finite values, connected graphs, full-rank nuisance designs, positive residual gene variances, and valid residual bases. Fixed audit draws tested reconstruction, fitted nuisance invariance, residual variance, covariance, correlation, Moran's $I$, endpoint diversity, and targeted-versus-full projection.

An initial v1.1 attempt stopped before surrogate generation because the observed residual replay used `float64` medians rather than the historical `float32` subtraction. No spatial surrogate statistic was produced or inspected. Amendment A03 restored the historical observed-replay arithmetic; genes, samples, graphs, models, seeds, endpoints, and decision rules were unchanged. Amendment A04 added dependency-ledger and audit outputs without changing inferential operations. Final analyses used Python 3.12.13, NumPy 2.3.5, pandas 2.3.3, and SciPy 1.18.1.

## Results

### Observed topology

The donor-averaged topology was $T=0.3451754386$ and the donor median was 0.4311403509. Five of six donor contrasts were positive; the one-sided sign test was not significant ($P=0.109375$). The donor-specific contrasts were 0.6859649123, 0.0745614035, 0.5087719298, 0.4385964912, 0.4236842105, and −0.0605263158.

Both adjacent correlations were positive in all six donors. The adjacent mean was 0.2986842105 and the donor-wise adjacent bottleneck was 0.2099415205. Mean endpoint similarity was −0.0464912281. In the diagnostic v1.0 component decomposition, the adjacent mean, bottleneck, and six-of-six positive-pair count were conditionally extreme after within-model Holm correction, whereas endpoint dissimilarity was not. The conditional extremeness of the aggregate statistic was therefore concentrated in adjacent-profile components rather than endpoint separation. This diagnostic decomposition had no confirmatory or independent decision role.

### Selection-conditioned robustness

Under the v1.0-P1 technical and selection-conditioned null, 1,395 of 50,000 panels equalled or exceeded the observation ($P=0.02791944$); the null mean was 0.13827817 and q95 was 0.31827485. Under the v1.0-S1 neuron-conditioned null, 1,328 of 50,000 panels equalled or exceeded the observation ($P=0.02657947$); the null mean was 0.17527322 and q95 was 0.32163743. Both met their frozen v1.0 criteria. These probabilities quantify post-discovery conditional robustness rather than independent confirmation.

Broad regional neuronal-minus-non-neuronal contrasts were positive in medulla oblongata (0.64067193), thalamus (0.56491715), and cerebral cortex (0.60420511). Against the earlier technically matched v0.11 single-nucleus null, 30 of 50,000 null sets equalled or exceeded the observed minimum contrast (plus-one $P=0.000620$). Against the stricter v1.0-S2 pseudoroute-conditioned comparator, the same weakest-region statistic did not exceed q95 ($P=0.09677806$; q95=0.61072560). Thus, neuronal attribution was directionally supported in broad regional proxies, but its apparent enrichment attenuated after conditioning on the selection mechanism and did not constitute route-specific cellular validation. None of the 50 geometry-nearest triplets exceeded the observed AHBA topology, yielding a deterministic tail/rank score of $1/51=0.01960784$. Across all 240 triplets, however, the target ranked third; two more distant triplets containing the spinal trigeminal nucleus aggregate (Sp5) had larger values. The result therefore did not establish DCML exclusivity.

The post hoc HPA exact-node statistic [14] was not exceptional under either conditional panel null ($P=0.21663567$ and 0.18523630). Across 50,000 one-probe-per-gene draws, every topology value was positive, but only 49.038% exceeded the historical technical-null q95. The gene-by-donor bootstrap median was 0.33803645 and its 2.5th–97.5th percentile range was −0.04328732 to 0.63134618.

### Public GSE3526 source-lineage sensitivity

All three probe-collapse choices yielded a positive mean relative serial-order contrast in the eight donors with complete public GSE3526 profiles. The archived descriptive results were:

| Node analysis | Probe collapse | Mean contrast | Positive donors | Descriptive label-tail fraction | Post-selection sign-flip tail fraction |
|---|---|---:|---:|---:|---:|
| Medulla–thalamus–parietal | Median | 0.213706140351 | 5/8 | 0.0996799268404 | 0.09765625 |
| Medulla–thalamus–parietal | Mean | 0.221052631579 | 6/8 | 0.110653863740 | 0.10937500 |
| Medulla–thalamus–parietal | Delegate | 0.191337719298 | 5/8 | 0.144032921811 | 0.11328125 |
| DRG–medulla–thalamus–parietal | Median | 0.152192982456 | 6/8 | 0.0576994230058 | 0.09765625 |
| DRG–medulla–thalamus–parietal | Mean | 0.165204678363 | 6/8 | 0.0506094939051 | 0.08593750 |
| DRG–medulla–thalamus–parietal | Delegate | 0.109722222222 | 6/8 | 0.132368676313 | 0.14062500 |

These positive contrasts must not be read as positive coexpression or continuous expression across the route. Under median collapse, the donor-mean pairwise correlations for the three-node analysis were −0.022149122807 (medulla–thalamus), −0.028508771930 (thalamus–parietal), and −0.239035087719 (medulla–parietal): the statistic was positive because the selected adjacent pairs were less dissimilar than the endpoint pair. In the four-node analysis, only the donor-mean DRG–medulla correlation was positive under median collapse (0.203289473684); the other five pairwise means ranged from −0.239035087719 to −0.010526315789.

The endpoint was also background-conditioned. In a median-collapse audit, replacing the donor-specific available neural background with the 20 neural tissues common to all eight donors gave approximate three- and four-node contrasts of 0.243 and 0.150, respectively; using all tissues available to each donor from the 65-tissue universe (29–58 tissues per donor) reduced them to approximately 0.099 and 0.045, and omitting within-donor neural standardization reduced them to approximately 0.003 and 0.011. Thus, the GSE3526 statistic describes relative serial order after normalization to the frozen neural-tissue background, not general cross-tissue similarity.

The three-node donor contrasts also showed strong exploratory metadata heterogeneity (Supplementary Figure S1). The four female donors had higher contrasts than all four male donors under every collapse method. Female-versus-male means were 0.563157894737 versus −0.135745614035 for median collapse, 0.568640350877 versus −0.126535087719 for mean collapse, and 0.529166666666 versus −0.146491228070 for the delegate. The minimum exact two-sided separation fraction was $2/70=0.0286$, but this is descriptive: sex was not a prespecified endpoint, each group contained only four donors, and sex is potentially confounded by age and by unrecorded cause of death, postmortem interval, RNA integrity, or array batch. The series summary provides age and sex but not those latter variables. RAI2 was the only X-linked candidate, but its removal did not account for the separation: female-minus-male differences remained 0.598, 0.590, and 0.560 under median, mean, and delegate collapse, respectively, with complete separation retained for median and mean. Leave-one-gene-out differences remained positive for every gene (0.549–0.768); jointly excluding RAI2 and C7orf25 left differences of 0.422–0.490, with strict separation retained only for the median. These post hoc checks argue that the pattern is not solely RAI2-driven but do not make it a sex effect. The four-node median contrast did not show an analogous separation (female 0.144005847953; male 0.160380116959). These observations establish donor heterogeneity and motivate a metadata-aware sensitivity; they do not establish a sex effect.

All six archived label-tail fractions and all six exact sign-flip fractions in the table were at least 0.05. A subsequent exact convolution over the finite four-node label space gave label-tail fractions of 0.0578575, 0.0487755, and 0.1311227 for median, mean, and delegate collapse, respectively, compared with the archived Monte Carlo values of 0.0576994, 0.0506095, and 0.1323687. The mean-collapse crossing of the conventional 0.05 boundary is itself evidence against a threshold-based narrative; the labels are not anatomically exchangeable and every fraction remains post-selection and descriptive, not confirmatory.

A post hoc exclusion of RAB3GAP2 and C7orf25 preserved the direction of the source-lineage result: across probe-collapse methods, mean contrasts ranged from 0.187653 to 0.250766 for three nodes and from 0.114890 to 0.203942 for four nodes, with five to six and six of eight positive donors, respectively. Because this 17-gene sensitivity was chosen after the annotation audit and remains tied to the historically dependent cohort, its tail fractions are descriptive and it does not establish gene-specific validity.

### Spatial surrogate results

The co-primary spatial components produced different conclusions. Under v1.1-P1A, 2,558 of 50,000 surrogate statistics were greater than or equal to the observation. The empirical probability was 0.0511789764 (Wilson 95% interval for $b/N$, 0.04926306–0.05312590), the null mean was 0.16938577, and q95 was 0.34590643. The observed statistic did not strictly exceed q95. v1.1-P1A therefore failed both numerical requirements and was flagged Monte Carlo borderline.

Under v1.1-P1B, 192 of 50,000 surrogates reached the observation ($P=0.0038599228$; Wilson interval, 0.00333468–0.00442155). The null mean was 0.00687241 and q95 was 0.22149854. v1.1-P1B passed.

Because the frozen primary rule required both components, the intersection-union probability was 0.0511789764 and the double gate did not pass. The resulting classification was `WHOLE_ONLY_SPATIAL_ROBUSTNESS`, and the separate across-model-stability classification was not applicable because the primary conjunction failed.

All four spatial sensitivities met their nominal, unadjusted individual numerical criteria. v1.1-S1 blockwise-dense yielded $P=0.0431956804$ and q95=0.33684942; v1.1-S2 whole-donor-dense yielded $P=0.0050994901$ and q95=0.22134503; v1.1-S3A blockwise axis-standardized yielded $P=0.0439956004$ and q95=0.33713450; and v1.1-S3B whole-donor axis-standardized yielded $P=0.0044995500$ and q95=0.22179094. No multiplicity-adjusted familywise inference was assigned to these four sensitivity models. They show that v1.1-P1A was near, but did not cross, its frozen boundary under one local spatial specification; they do not override the failed primary conjunction.

The three whole-donor null distributions were centred near zero, whereas the three blockwise distributions had means of 0.15796655–0.16938577. All whole-donor models met their individual numerical criteria, whereas the blockwise models lay near the decision boundary; this contrast demonstrates dependence on the spatial formulation.

### Donor sensitivity

All six LODO observed effects were positive, ranging from 0.27701754 to 0.42631579. For v1.1-P1A, 3/6 omissions exceeded the corresponding q95 and empirical probabilities ranged from 0.02305954 to 0.18661627. For v1.1-P1B, all 6/6 omissions exceeded q95 and probabilities ranged from 0.00099998 to 0.02953941. The sign was not driven by a single donor, but calibration under the blockwise model was not uniformly stable. LODO samples overlap and are not independent replications.

### Numerical audit and deterministic re-execution

The observed residual tensor reproduced v0.10 with maximum absolute difference $2.2204\times10^{-16}$, and the donor endpoints reproduced v1.0 with maximum difference $8.3267\times10^{-17}$. Across 72 fitted spatial units, the maximum basis-orthogonality error was $7.4096\times10^{-13}$, the maximum all-plus reconstruction error was $5.3091\times10^{-13}$, the maximum residual covariance difference was $4.7518\times10^{-14}$, and the maximum residual Moran-$I$ difference was $2.9410\times10^{-14}$. All technical gates passed.

The input ledger contained 59 verified rows. A second clean deterministic execution with the same sealed code and recorded numerical environment generated 16 files identical byte for byte to the preserved first complete A04 run. This demonstrates computational repeatability, not an independent implementation or biological replication. The frozen v1.1-A04 analysis output-manifest SHA-256 was `eb90201948f278e15e42f4b70f0b42d27693649acb6f03573c47b1165e56cecd`, and the v1.1-A04 summary SHA-256 was `2455156e44c2e2946ee3e8d28c368ac7359a829afddcfe5c8a6577f907ed089e`; neither is the package-level A05 release manifest.

## Discussion

The frozen 19-gene panel showed a positive donor-averaged transcriptomic adjacency contrast across the six previously inspected AHBA donors; the donor-specific contrast was positive in five of six. Selection-aware nulls indicated that the signal was unusual relative to panels obtainable through analogous anatomical and technical filters. The spatial analysis added a more demanding result: all three whole-donor models met their individual numerical criteria, whereas results under the intracompartimental models were near the decision boundary and the predeclared v1.1-P1A component failed just above 0.05 and just below its 95th percentile. The primary double gate therefore failed.

The selection-aware and spatial analyses address different null hypotheses and were not combined into a single joint randomization. Version 1.0 varied the gene panel while holding the observed AHBA regional residuals fixed; version 1.1 randomized sample-level spatial fields for the fixed 19-gene panel. Their probabilities cannot be multiplied, combined, or described as a joint test showing that the result "survived selection and space." In addition, the v1.0 gene null did not explicitly match within-panel coexpression or spatial-autocorrelation profiles, whereas v1.1 preserved the covariance of this already-selected panel. A coexpression- and spatially matched selection null remains a prospective analysis.

This divergence is scientifically informative. Whole-donor models preserve linear coordinates and broad anatomical fixed effects while randomizing a global residual field. Blockwise models randomize within the three target macrocompartments, where the null topology itself was shifted upward. The data support an adjacency contrast under the whole-donor spatial specification, but do not show that it is uniformly exceptional after local within-compartment spatial organization is preserved. Describing the outcome simply as “spatially significant” would conceal the principal inferential result.

Several observations nevertheless argue against a trivial single-donor artifact. Five of six donor contrasts were positive, both adjacent links were positive in every donor, and every LODO effect remained positive. The whole-donor spatial calibration passed in all six omissions. Conversely, the donor sign test was not significant, v1.1-P1A passed in only three omissions, and one donor contributed only a single Cu/Gr target sample. These facts support qualitative consistency but not donor-level replication.

Anatomical specificity is also unresolved. The target triplet was high-ranking among regional alternatives, but two Sp5-containing triplets were stronger in the full search. This may reflect a broader somatosensory molecular organization rather than a DCML-specific route. DTLv and postcentral measurements are atlas sampling proxies, and regional expression similarity does not establish synaptic or axonal continuity.

Probe and platform limitations remain material. Random one-probe selections preserved the direction of the effect but not consistent exceedance of the historical null threshold. In the public GSE3526 reconstruction, the historical RAB3GAP2 probe sets are also annotated to AURKAPS1 and the sole C7orf25 probe set is also annotated to PSMA2. Excluding both historically assigned genes in a labelled post hoc sensitivity preserved the mean contrast direction, but it did not validate the remaining probe annotations or confer confirmatory status; CEL-level reprocessing with an updated definition remains desirable. HPA and broad neuronal contrasts provided descriptive corroboration without significant conditional specificity. An uninspected RNA-seq, single-nucleus, or spatial-transcriptomic dataset with suitable brainstem, thalamic, and cortical coverage is required for independent confirmation.

The GSE3526 reconstruction adds a useful but limited view of donor heterogeneity from a public cohort in the documented source lineage of BIGE. Formal sample- and processing-level reconciliation with the internally analyzed BIGE aggregate remains outstanding. Its positive relative-order contrast arose largely because adjacent pairs were less negative than the selected nonadjacent pairs, not because all adjacent tissues were positively correlated, and its magnitude depended strongly on the tissue background used for within-donor standardization. The broad GSE3526 medulla, thalamus, parietal lobe, and DRG are not anatomically equivalent to the AHBA relay proxies, so the magnitudes are not directly comparable and do not demonstrate route specificity. Its complete separation by recorded sex in the three-node contrast further argues against a simple, generally consistent cohort-level interpretation and requires metadata-aware analysis; the pattern was not explained solely by the X-linked candidate RAI2, but the available metadata cannot resolve major confounders. Because the cohort is in the same historical source lineage, and because the gene panel and tissue order were already selected, it is a source-lineage post-selection sensitivity rather than replication or validation.

The study's main strengths are the preserved negative discovery history, frozen gene set, conditional reconstruction of prespecified parts of the selection process, conservative double spatial gate, joint surrogates that retain residual cross-gene covariance, auditable hash-linked provenance trail, and byte-identical re-execution. Its main weaknesses are post-discovery use of AHBA, separate rather than joint selection-and-space nulls, lack of explicit coexpression matching in the gene-panel null, sparse and unequal regional sampling, dependence on spatial model, sensitivity to probe choice, imperfect preservation of higher-order marginal distributions by singleton MSR, and the absence of independent atlas confirmation. Broad external cell-class contrasts do not constitute AHBA sample-level adjustment or deconvolution, so residual confounding by cellular composition remains formally uncontrolled. Irregular sampling also produced a maximum retained v1.1-P1A nearest-neighbour edge of 49.13 mm, motivating prospective distance-bin and variogram diagnostics.

Accordingly, the present results support a model-dependent transcriptomic adjacency pattern. They do not support connectivity, direction, molecular flow, causality, route exclusivity, a pain or fibromyalgia mechanism, or selection of a stimulation target. Clinical or interventional hypotheses should be developed only in a separate, prospectively designed study.

## Data and code availability

The analysis code, frozen protocols, amendments, manifests, aggregate result tables, and reproducibility records are being prepared for release at [repository/DOI placeholder]. AHBA access, attribution, and redistribution will follow Allen Institute terms and citation requirements [9,10].

The discovery source was an internally maintained, processed extract of BIGE. Investigator authorization permits publication of non-disclosive aggregate statistics, result tables, figures, and analytical descriptions derived from that extract. The complete internally maintained matrix and exports that would reconstruct it will not be redistributed. A file-level release audit excluded the matrix and reconstructive derivatives from the release candidate. The public GSE3526 series matrix and sample metadata are available from GEO (https://www.ncbi.nlm.nih.gov/geo/query/acc.cgi?acc=GSE3526). GSE3526 is in the documented source lineage, but its numerical equivalence to the internally analyzed aggregate has not been established and it is not designated as independent validation. Data-access contact and conditions for the internally maintained extract: [placeholder].

## Author contributions

[CRediT statement placeholder]

## Funding

[Placeholder]

## Competing interests

[Placeholder; include any relevant institutional or commercial relationships after author review.]

## Ethics statement

[Confirm the applicable secondary-use and source-dataset statements before submission.]

## References

1. Hawrylycz MJ, Lein ES, Guillozet-Bongaarts AL, et al. An anatomically comprehensive atlas of the adult human brain transcriptome. *Nature*. 2012;489:391–399. https://doi.org/10.1038/nature11405
2. Wagner HH, Dray S. Generating spatially constrained null models for irregularly spaced data using Moran spectral randomization methods. *Methods in Ecology and Evolution*. 2015;6:1169–1178. https://doi.org/10.1111/2041-210X.12407
3. Burt JB, Helmer M, Shinn M, Anticevic A, Murray JD. Generative modeling of brain maps with spatial autocorrelation. *NeuroImage*. 2020;220:117038. https://doi.org/10.1016/j.neuroimage.2020.117038
4. Markello RD, Misic B. Comparing spatial null models for brain maps. *NeuroImage*. 2021;236:118052. https://doi.org/10.1016/j.neuroimage.2021.118052
5. Markello RD, Arnatkeviciute A, Poline J-B, Fulcher BD, Fornito A, Misic B. Standardizing workflows in imaging transcriptomics with the abagen toolbox. *eLife*. 2021;10:e72129. https://doi.org/10.7554/eLife.72129
6. Siletti K, Hodge R, Mossi Albiach A, et al. Transcriptomic diversity of cell types across the adult human brain. *Science*. 2023;382(6667):eadd7046. https://doi.org/10.1126/science.add7046
7. Vos de Wael R, Benkarim O, Paquola C, et al. BrainSpace: a toolbox for the analysis of macroscale gradients in neuroimaging and connectomics datasets. *Communications Biology*. 2020;3:103. https://doi.org/10.1038/s42003-020-0794-7
8. Allen Institute. Allen Human Brain Atlas API documentation. https://brain-map.org/support/documentation/human-brain-atlas-api
9. Allen Institute. Terms of Use. https://alleninstitute.org/legal/terms-of-use
10. Allen Institute. Citation Policy. https://alleninstitute.org/legal/citation-policy
11. Lee J, Hever A, Willhite D, Zlotnik A, Hevezi P. Effects of RNA degradation on gene expression analysis of human postmortem tissues. *FASEB Journal*. 2005;19:1356–1358. https://doi.org/10.1096/fj.04-3552fje
12. Roth RB, Hevezi P, Lee J, et al. Gene expression analyses reveal molecular relationships among 20 regions of the human CNS. *Neurogenetics*. 2006;7:67–80. https://doi.org/10.1007/s10048-006-0032-6. GEO: GSE3526.
13. Ushach I, Burkhardt AM, Martinez C, et al. METEORIN-LIKE is a cytokine associated with barrier tissues and alternatively activated macrophages. *Clinical Immunology*. 2015;156:119–127. https://doi.org/10.1016/j.clim.2014.11.006
14. Human Protein Atlas. Brain data, version 25.1. Accessed 9 September 2026. https://www.proteinatlas.org/humanproteome/brain/data
15. National Center for Biotechnology Information. Gene Expression Omnibus platform GPL570: Affymetrix Human Genome U133 Plus 2.0 Array. Accessed 9 September 2026. https://www.ncbi.nlm.nih.gov/geo/query/acc.cgi?acc=GPL570
