## Supplementary Archive S1 for "An anatomy-constrained transcriptomic screen identifies a 19-gene panel with an ordered regional similarity pattern in the human somatosensory system": README.md

# Route transcriptomics v1.1-A04

This directory contains the frozen spatial-autocorrelation extension of the DCML19 regional transcriptomic-topology analysis. It is an internal, post-discovery scientific checkpoint; it is not an external preregistration, an independent replication, or a public-data release.

## Scientific scope

For a fixed 19-gene panel, the analysis asks whether residual-expression profiles are more similar between the two adjacent covered region pairs (Cu/Gr–DTLv and DTLv–postcentral cortex) than between the non-adjacent endpoints (Cu/Gr–postcentral cortex). The observed donor-mean adjacency contrast is `T = 0.3451754385964913`.

The v1.1 extension evaluates that already-observed statistic with joint Moran spectral randomization (MSR). The custom float64 implementation preserves the fitted nuisance component, residual Moran spectral power, and residual gene covariance. It is adapted from Wagner and Dray (2015), is not a bitwise reproduction of BrainSpace, and does not preserve higher-order univariate marginals. Allen `sample.mri` coordinates are treated as millimetre coordinates in each donor's native MR-volume space, not as MNI coordinates.

The analysis does **not** establish anatomical connectivity, direction, molecular flow, causality, route exclusivity, a pain or fibromyalgia mechanism, a stimulation target, or independent replication.

## Frozen endpoint and exact result

The primary decision was an intersection–union double gate: both P1A and P1B had to satisfy `p <= 0.05` and place the observation beyond their linearly interpolated 95th null percentile. That conjunction did not pass.

| Model | Scope / weights | Draws | Empirical p (+1) | Null q95 | Result |
|---|---|---:|---:|---:|---|
| P1A | blockwise, native-coordinate 5-NN | 50,000 | 0.0511789764 | 0.3459064327 | FAIL |
| P1B | whole donor, native-coordinate 6-NN | 50,000 | 0.0038599228 | 0.2214985380 | PASS |
| S1 | blockwise, dense native-distance | 10,000 | 0.0431956804 | 0.3368494152 | PASS |
| S2 | whole donor, dense native-distance | 10,000 | 0.0050994901 | 0.2213450292 | PASS |
| S3A | blockwise, axis-standardized 5-NN | 10,000 | 0.0439956004 | 0.3371345029 | PASS |
| S3B | whole donor, axis-standardized 8-NN | 10,000 | 0.0044995500 | 0.2217909357 | PASS |

P1A had 2,558 exceedances in 50,000 draws; the observation was `0.0007309942` below its q95 and its binomial Wilson interval for the exceedance fraction was 0.04926–0.05313. P1B had 192 exceedances. The frozen intersection–union p-value is therefore `max(p_P1A, p_P1B) = 0.0511789764`.

The prespecified machine label is:

```text
WHOLE_ONLY_SPATIAL_ROBUSTNESS
```

This label means that P1B passed while P1A did not. Five of the six evaluated formulations passed, including both secondary blockwise sensitivity models, but the sensitivities do not override the failed primary double gate. The defensible interpretation is **convergent but non-uniform, model-dependent spatial support**.

## Directory map

- `protocol/`: base v1.1 protocol, hash-linked amendments A01–A04, and the preserved A03 pre-audit executable.
- `scripts/`: frozen endpoint, independent validator, and figure-generation code.
- `analysis/`: authoritative final A04 output set from the second execution.
- `verification/`: independent validation report for output hashes, endpoint calculations, QC, ledger, and repeat execution.
- `verification_runs/`: preserved first complete A04 execution; intentionally omitted from the curated release because it duplicates `analysis/` byte for byte.
- `failed_runs/`: failed-closed pre-surrogate incident and its exact A02 executable; no v1.1 surrogate statistic was generated in that attempt.
- `figures/`: manuscript-ready spatial-result figures and their manifest.
- `report/`: Spanish scientific decision report integrating v1.0 and v1.1.
- `manuscript/`: English manuscript draft integrating v1.0 and v1.1.

## Authoritative records

- `analysis/summary_v1_1.json`
- `analysis/spatial_analysis_hashes_v1_1.txt`
- `verification/spatial_validation_v1_1.json`
- `protocol/protocol_v1_1.json`
- `protocol/amendment_01_double_gate_and_trend_preservation_v1_1.json`
- `protocol/amendment_02_endpoint_seal_v1_1.json`
- `protocol/amendment_03_float32_residual_replay_v1_1.json`
- `protocol/amendment_04_dependency_and_audit_seal_v1_1.json`

The final summary SHA-256 is `2455156e44c2e2946ee3e8d28c368ac7359a829afddcfe5c8a6577f907ed089e`; the analysis-manifest SHA-256 is `eb90201948f278e15e42f4b70f0b42d27693649acb6f03573c47b1165e56cecd`.

## Reproducibility and quality control

The independent validator reports `PASS`:

- all 16 final analysis files were byte-identical across two complete A04 executions;
- all 72 fitted spatial graphs were connected;
- all 72 invariant-audit rows passed their frozen numerical gates;
- the input ledger contains 59 unique, pre/post-verified sources, including 44 AHBA probe files;
- the maximum residual-correlation invariant discrepancy was approximately `1.01e-13`, well below its `1e-8` gate.

The recorded execution environment used Python 3.12.13, NumPy 2.3.5, pandas 2.3.3, SciPy 1.18.1, and OpenBLAS 0.3.30. See `analysis/environment_versions_spatial_v1_1.txt` for the complete build record.

## Validation and rerun commands

From the project root, the authoritative full workspace can be revalidated with:

```bash
python work/route_v1_1/scripts/verify_spatial_outputs_v1_1.py
```

Expected status: `PASS`, with 15 manifest entries, 59 ledger rows, and 16 files byte-identical to `verification_runs/spatial_A04_run1`.

An exact endpoint re-execution must be performed in a **separate clean copy** of the complete project, with neither `work/route_v1_1/analysis` nor `work/route_v1_1/analysis.staging` present:

```bash
python work/route_v1_1/scripts/analyze_spatial_v1_1.py
python work/route_v1_1/scripts/verify_spatial_outputs_v1_1.py
```

The endpoint deliberately refuses to overwrite an existing final or staging directory. It also verifies all frozen inputs before and after execution. Re-execution requires the upstream v0.8 and v0.10 reconstructed workspaces at the exact paths encoded in the script, plus `work/route_v1_0`. The curated release is an audited result snapshot and does not include those upstream dependencies or the duplicate first execution.

After validation, the spatial figures can be regenerated atomically with:

```bash
python work/route_v1_1/scripts/plot_spatial_v1_1.py
```

The two consecutive final figure builds were byte-identical. Their generator, source-data hashes, output hashes, software versions, and font hash are recorded in `figures/figure_manifest_spatial_v1_1.json` (SHA-256 `7d51707d54209f7e4a608dc19aadf2780a3abb2f39494e0d0df3b5a9652dceea`).

## Data exclusions and release status

The curated release excludes the 154 MB private source database, private frozen matrices and selection metrics, reconstructed Allen public-raw files, bulky duplicate executions, caches, zero-byte temporary plotting files, and other transient artifacts. It retains hashes and dependency ledgers for auditability.

Some v1.0 outputs and null-design derivatives originated downstream of the private discovery database. Inclusion in this internal package does not itself establish permission for external redistribution. Public release of private-derived tables, Allen-derived content, or other source-dependent artifacts requires institutional/data-governance and applicable licence review. The manuscript can report aggregate results while that review is pending, but the package must not be represented as a cleared public dataset.
