## Supplementary Archive S1 for "An anatomy-constrained transcriptomic screen identifies a 19-gene panel with an ordered regional similarity pattern in the human somatosensory system": Informe_avance_cientifico_v1_1_A05.md

# Informe científico del análisis v1.1-A04 — liberación documental A05

**Proyecto:** contraste transcriptómico restringido por anatomía en proxies regionales de relés somatosensoriales humanos  
**Cierre del análisis AHBA v1.1-A04:** 8 de septiembre de 2026  
**Auditoría GSE3526 y actualización documental A05:** 9 de septiembre de 2026  
**Estado:** liberación documental A05; los resultados inferenciales v1.1-A04 permanecen sin cambios; control espacial post-descubrimiento completo, auditado y reproducido; reconstrucción pública GSE3526 integrada como sensibilidad del linaje fuente documentado; el doble *gate* espacial primario no se superó

## Veredicto ejecutivo

El proyecto sigue siendo publicable, pero la conclusión debe ser más estrecha que la propuesta al cierre de v1.0. En los residuales regionales AHBA, el panel congelado de 19 genes mantuvo un contraste positivo de adyacencia transcriptómica y superó los controles condicionales de selección de v1.0. Sin embargo, en v1.1 la evidencia espacial dependió de cómo se definió el campo espacial.

El contraste observado fue $T=0.3451754386$. En el modelo espacial primario intracompartimental P1A, 2,558 de 50,000 sustitutos igualaron o superaron el valor observado: $p=0.0511789764$ y q95=0.3459064327. Por tanto, el observado quedó por debajo de q95 y **P1A no pasó**. En el modelo co-primario de donante completo P1B, 192 de 50,000 sustitutos alcanzaron el observado: $p=0.0038599228$ y q95=0.2214985380; **P1B sí pasó**. El *p* de intersección-unión fue el máximo de ambos, $p_{IUT}=0.0511789764$.

La regla congelada exigía que P1A y P1B pasaran simultáneamente. Esa conjunción **falló** y la clasificación resultante es:

> `WHOLE_ONLY_SPATIAL_ROBUSTNESS`

Las cuatro sensibilidades espaciales también superaron sus criterios numéricos individuales nominales y no ajustados; no recibieron una inferencia familiar ni sustituyen el *gate* primario fallido. El resultado es, en consecuencia, **robustez espacial dependiente del modelo dentro del mismo AHBA**, no robustez espacial general ni confirmación independiente.

La reconstrucción donante por donante en GSE3526 añade información sobre heterogeneidad de una cohorte pública perteneciente al linaje fuente documentado de BIGE. La equivalencia numérica entre el agregado reconstruido y el extracto BIGE efectivamente analizado no se ha establecido mediante una reconciliación de muestras y procesamiento. No es replicación: reutiliza el panel y el orden anatómico ya seleccionados, y la menor fracción de cola descriptiva varió entre 0.0506095 en el Monte Carlo archivado y 0.0487755 en la convolución exacta posterior, sin interpretación confirmatoria. Además, mostró una separación exploratoria completa por sexo en el contraste de tres nodos, y el contraste positivo refleja principalmente que los pares adyacentes fueron menos disímiles que los no adyacentes seleccionados, no coexpresión positiva general ni continuidad.

La afirmación central defendible es:

> En datos AHBA previamente inspeccionados, un panel congelado de 19 genes mostró un contraste residual positivo en cinco de seis donantes y ambas correlaciones adyacentes positivas en los seis. La prueba de signo del contraste no fue significativa. El patrón fue favorable frente a nulos condicionados por selección y frente a varios modelos espaciales conjuntos, pero la evidencia espacial no fue uniforme: pasó el modelo de donante completo y falló el componente intracompartimental primario preespecificado.

No se infiere conectividad, dirección, flujo molecular, causalidad, exclusividad de DCML, mecanismo de dolor o fibromialgia ni un blanco de estimulación.

## Trayectoria científica que debe conservarse

La historia no comienza con una hipótesis clínica confirmada. La secuencia documentada fue:

1. Se catalogaron 16 circuitos y se congelaron cinco rutas.
2. Tres rutas se examinaron en **un extracto procesado y mantenido internamente del Body Index of Gene Expression (BIGE)**, compuesto por promedios regionales.
3. Surgieron 23 candidatos: 19 seleccionados en el cribado restringido por DCML, tres en modulación descendente del dolor y uno en ganglios basales.
4. La hipótesis original de continuidad transcriptómica global a lo largo de las rutas fue negativa.
5. Después se observó en AHBA una topología entre tres proxies regionales para el panel de 19 genes.
6. v1.0 evaluó esa observación conocida con nulos condicionales de selección, neuronalidad, anatomía, sonda y donante.
7. v1.1 evaluó autocorrelación y tendencias espaciales con randomización espectral de Moran conjunta.

Los análisis v1.0 y v1.1 son extensiones de robustez diseñadas después de conocer la señal. Aunque sus protocolos fueron congelados antes de abrir sus nuevos endpoints, no son una preregistración externa ni una replicación en datos no inspeccionados.

## Proveniencia de BIGE, autorización y reconstrucción GSE3526

BIGE se ha descrito en la literatura como una base de microarreglos Affymetrix Human Genome U133 Plus 2.0 con expresión de más de 105 células y tejidos humanos. La cadena de procedencia relevante incluye el estudio de degradación de RNA post mortem de Lee et al. (2005), el análisis de 353 arreglos y 65 tejidos normales de Roth et al. (2006), depositado como GEO GSE3526, y la descripción publicada de BIGE por Ushach et al. (2015). El insumo de descubrimiento debe describirse como **un extracto procesado y mantenido internamente de BIGE**, no como un atlas privado sin nombre.

Existe autorización del investigador para publicar estadísticas agregadas, tablas, gráficos, descripciones metodológicas y otros resultados no reconstructivos derivados del extracto. La matriz interna completa y exportaciones que permitan reconstruirla no se publicarán ni redistribuirán. Esto resuelve el bloqueo para analizar y comunicar resultados agregados. El inventario archivo por archivo de A05 ya excluyó la matriz y los derivados reconstructivos; cualquier liberación futura deberá repetir ese control.

La serie pública GSE3526 contiene 353 muestras, 65 tipos de tejido y diez donantes. La matriz oficial contiene intensidades RMA en escala lineal (rango 4.538–33,730.54), por lo que la transformación $\log_2(x+1)$ es apropiada y no una segunda transformación logarítmica; sustituirla por $\log_2(x)$ apenas cambió el endpoint. La reconstrucción usó 39 sondas GPL570 para el panel congelado de 19 genes, colapso por mediana, media y sonda delegada histórica, y estandarización por gen dentro de donante a través de los 21–23 tejidos disponibles para ese donante dentro del universo neural congelado de 23 tejidos. La serie contiene 193 arreglos neurales en los diez donantes; 180 pertenecen a los ocho donantes con endpoint completo y pueden influir en sus perfiles estandarizados. El endpoint utilizó 24 arreglos para tres nodos o 32 para cuatro nodos. No hubo duplicados donante–tejido. Ocho donantes (1, 2, 3, 4, 5, 8, 9 y 10) tuvieron los cuatro tejidos requeridos y constituyen las unidades analíticas a nivel de donante ($n=8$). Médula, tálamo, lóbulo parietal y ganglio de la raíz dorsal (DRG) son tejidos amplios y no equivalen a Cu/Gr, DTLv y giro postcentral de AHBA; sus magnitudes no son directamente comparables ni demuestran especificidad de relé o de ruta. GSE3526 pertenece al linaje fuente documentado de BIGE, pero la equivalencia numérica con el agregado interno no está reconciliada. Por ello esta reconstrucción es una **sensibilidad postselección del mismo linaje fuente**, no replicación o validación independiente.

| Análisis | Colapso | Contraste medio | Donantes positivos | Fracción de cola por etiquetas | Fracción de cola por signos |
|---|---|---:|---:|---:|---:|
| Médula–tálamo–parietal | Mediana | 0.213706140351 | 5/8 | 0.0996799268404 | 0.09765625 |
| Médula–tálamo–parietal | Media | 0.221052631579 | 6/8 | 0.110653863740 | 0.10937500 |
| Médula–tálamo–parietal | Delegada | 0.191337719298 | 5/8 | 0.144032921811 | 0.11328125 |
| DRG–médula–tálamo–parietal | Mediana | 0.152192982456 | 6/8 | 0.0576994230058 | 0.09765625 |
| DRG–médula–tálamo–parietal | Media | 0.165204678363 | 6/8 | 0.0506094939051 | 0.08593750 |
| DRG–médula–tálamo–parietal | Delegada | 0.109722222222 | 6/8 | 0.132368676313 | 0.14062500 |

Ninguna fracción de la tabla es un valor $p$ confirmatorio. Las permutaciones de etiquetas presuponen intercambiabilidad anatómica que aquí no se sostiene, y las enumeraciones de signos se ejecutaron después de seleccionar panel, cohorte y endpoint. Una convolución exacta posterior del espacio finito de etiquetas de cuatro nodos produjo fracciones 0.0578575, 0.0487755 y 0.1311227 para mediana, media y delegada, frente a 0.0576994, 0.0506095 y 0.1323687 en el Monte Carlo archivado. Que el resultado de media cruce 0.05 según el cálculo demuestra precisamente por qué no admite una narrativa de umbral: ambos cálculos son postselección y las etiquetas no son intercambiables. Bajo el colapso por mediana, las correlaciones medias de tres nodos fueron −0.022149122807 para médula–tálamo, −0.028508771930 para tálamo–parietal y −0.239035087719 para médula–parietal. Por tanto, $T>0$ significa **menor disimilitud relativa entre pares adyacentes**, no expresión compartida positiva ni continuidad transcriptómica. En cuatro nodos sólo DRG–médula tuvo una correlación media positiva con colapso por mediana (0.203289473684); las otras cinco fueron pequeñas o negativas. El resultado está además condicionado por el fondo de normalización: con los 20 tejidos neurales comunes a los ocho donantes, $T$ fue aproximadamente 0.243/0.150 para tres/cuatro nodos; al usar todos los tejidos disponibles para cada donante dentro del universo de 65 (29–58 tejidos por donante), fue 0.099/0.045; y sin estandarización neural, 0.003/0.011. Debe describirse como desviación relativa al fondo neural congelado, no como similitud general.

La heterogeneidad más llamativa fue post hoc: las cuatro donantes mujeres superaron a los cuatro donantes hombres en el contraste de tres nodos con los tres métodos. Las medias mujer frente a hombre fueron 0.563157894737 frente a −0.135745614035 (mediana), 0.568640350877 frente a −0.126535087719 (media) y 0.529166666666 frente a −0.146491228070 (delegada). La fracción exacta bilateral mínima para una separación completa es $2/70=0.0286$, pero es sólo descriptiva: hay cuatro donantes por grupo y sexo puede estar confundido con edad y con causa de muerte, intervalo post mortem, integridad de RNA o lote, que no aparecen en el resumen de la serie; éste sólo proporciona edad y sexo. RAI2 es el único candidato ligado al cromosoma X, pero retirarlo no explicó la separación: las diferencias mujer−hombre fueron 0.598, 0.590 y 0.560 con mediana, media y delegada, y la separación completa persistió para mediana y media. Al retirar cada gen por separado, todas las diferencias siguieron positivas (0.549–0.768); al excluir conjuntamente RAI2 y C7orf25 quedaron en 0.422–0.490, con separación estricta sólo bajo mediana. El endpoint de cuatro nodos no mostró una separación análoga bajo mediana (mujeres 0.144005847953; hombres 0.160380116959). El patrón no depende sólo de RAI2, pero no permite inferir un efecto sexual; se requiere una sensibilidad prospectiva que incorpore metadatos.

La anotación vigente de GPL570 introduce otra limitación: las tres sondas atribuidas históricamente a RAB3GAP2 también mapean a AURKAPS1, y la única sonda de C7orf25 también mapea a PSMA2. Una sensibilidad post hoc que excluyó ambos genes conservó la dirección: los contrastes medios fueron 0.187653–0.250766 en tres nodos y 0.114890–0.203942 en cuatro, con cinco a seis y seis de ocho donantes positivos, respectivamente. Esto reduce la posibilidad de que la dirección dependa sólo de esas asignaciones, pero no valida las sondas restantes ni convierte el análisis en confirmatorio; sigue siendo deseable reprocesar los archivos CEL con una definición contemporánea.

## Estadístico y señal observada

Para cada donante AHBA se calcularon correlaciones de Spearman entre los perfiles residuales de los 19 genes en cuneado/grácil (Cu/Gr), la división ventral agregada del tálamo dorsal (DTLv) y giro postcentral. DTLv no equivale a una medición aislada del núcleo ventral posterolateral (VPL). El estadístico congelado fue:

$$
T_d=\frac{\rho_{d,\mathrm{Cu/Gr,DTLv}}+\rho_{d,\mathrm{DTLv,Postcentral}}}{2}
-\rho_{d,\mathrm{Cu/Gr,Postcentral}},
$$

y el resultado agregado fue la media de $T_d$ en seis donantes.

El valor observado fue $T=0.3451754386$ y la mediana por donante fue 0.4311403509. Cinco de seis contrastes por donante fueron positivos; la prueba de signo unilateral no fue significativa ($p=0.109375$). Los contrastes individuales fueron 0.6859649123, 0.0745614035, 0.5087719298, 0.4385964912, 0.4236842105 y −0.0605263158. Ambas correlaciones adyacentes fueron positivas en los seis donantes, pero en el último donante la semejanza entre extremos fue todavía mayor.

En una descomposición diagnóstica v1.0, la excepcionalidad condicional se concentró en los componentes de semejanza adyacente y no en una separación excepcional entre extremos:

| Componente | Observado | Resultado condicionado |
|---|---:|---|
| Media de las dos adyacencias | 0.2986842105 | $p_{Holm}=0.04871903$ en P1 y 0.04499910 en S1 |
| Cuello de botella adyacente | 0.2099415205 | $p_{Holm}=0.03767925$ en P1 y 0.02839943 en S1 |
| Ambos pares positivos | 6/6 donantes | $p_{Holm}=0.04871903$ en P1 y 0.04499910 en S1 |
| Similitud entre extremos | −0.0464912281 | no excepcional: $p_{Holm}=0.40705186$ y 0.52170957 |

Esta descomposición estaba sellada como diagnóstica: sus valores ajustados localizan qué componente concentra la excepcionalidad condicional, pero no tienen papel confirmatorio ni constituyen una decisión independiente.

La expresión correcta es **similitud residual o coordinación transcriptómica entre relés adyacentes**. “Conectividad”, “transmisión” y “flujo” exceden los datos.

## Resultados integrados de v1.0

| Análisis | Resultado exacto | Alcance |
|---|---:|---|
| v0.11, atribución neuronal en single-nucleus | estadístico mínimo=0.56491715; 30/50,000 ≥ observado; $p_{+1}=0.000620$ | Favorable frente al nulo técnico emparejado de v0.11, en proxies regionales amplios |
| P1, pseudorrutas condicionadas técnicamente | 1,395/50,000; $p=0.02791944$; q95=0.31827485 | Favorable frente a paneles que aproximan partes prespecificadas del mecanismo de selección y reproducen los estratos técnicos |
| S1, pseudorrutas más neuronalidad amplia | 1,328/50,000; $p=0.02657947$; q95=0.32163743 | Favorable al añadir contrastes neuronales regionales amplios |
| S2, atribución celular condicionada | observado=0.56491715; $p=0.09677806$; q95=0.61072560 | Positiva en las tres macrorregiones, pero no enriquecida significativamente |
| S3, 50 alternativas anatómicas cercanas | 0/50 ≥ observado; score de rango $1/51=0.01960784$ | Favorable como comparación determinista; no es un *p* exacto de randomización |
| Universo completo de tríadas | tercer lugar de 240 | Dos combinaciones distantes con el agregado del núcleo trigeminal espinal (Sp5) fueron mayores; no hay exclusividad de DCML |
| HPA exact-node post hoc | observado=0.78070175; $p=0.21663567$ y 0.18523630 | Corroboración descriptiva, no especificidad ni replicación |
| Selección de una sonda por gen | mediana=0.29809942; percentiles 2.5–97.5% [0.16257310, 0.42836257]; 49.038% sobre q95 histórico | El signo fue estable, pero la excepcionalidad dependió de la sonda; no es un IC |
| Bootstrap gen × donante | mediana=0.33803645; intervalo percentil diagnóstico 2.5–97.5% [−0.04328732, 0.63134618]; 96.114% >0 | Apoyo direccional con incertidumbre que incluye cero; no es un IC confirmatorio |

El rótulo v1.0 `ROBUST_COVERED_TOPOLOGY` fue una clasificación operativa para controles condicionales no espaciales. No prevalece sobre el resultado espacial más específico de v1.1 ni debe utilizarse como una conclusión biológica.

La atribución neuronal requiere una lectura histórica explícita. En v0.11, 30 de 50,000 conjuntos del nulo técnicamente emparejado igualaron o superaron el resultado observado en datos humanos de núcleo único ($p=0.000620$). Al comparar el mismo estadístico con los paneles v1.0 que además aproximan partes prespecificadas de la selección por pseudorruta, la evidencia se atenuó a $p=0.09677806$. No es una contradicción: los comparadores responden preguntas diferentes. La conclusión integrada es que el panel tiene atribución neuronal direccional en proxies amplios, pero no demuestra enriquecimiento neuronal específico frente al comparador condicional más estricto.

Los controles v1.0 y v1.1 tampoco constituyen una sola prueba conjunta. v1.0 varía paneles génicos sobre los residuales regionales AHBA observados; v1.1 randomiza campos espaciales para el panel fijo de 19 genes. Sus probabilidades no deben multiplicarse, combinarse ni resumirse diciendo que la señal “sobrevivió conjuntamente selección y espacio”. Además, el nulo de paneles v1.0 no emparejó explícitamente la coexpresión interna o la autocorrelación espacial de cada panel. v1.1 preserva la covarianza del panel ya seleccionado, pero no corrige retrospectivamente esa limitación de selección.

## Diseño espacial v1.1

El análisis reconstruyó los mapas a partir de 44 sondas AHBA de símbolo exacto. Se conservaron las conversiones históricas `float32` para la lectura, reordenamiento, mediana entre sondas y replay de los residuales observados; la regresión y la randomización espacial se ejecutaron en `float64`. De 3,702 muestras fuente, las 2,797 muestras izquierdas o no pareadas se usaron en cada modelo de donante completo. Los 18 bloques intracompartimentales reunieron 540 muestras relevantes de mielencéfalo, tálamo dorsal y lóbulo parietal.

Los sustitutos se generaron mediante una implementación propia de randomización espectral de Moran conjunta y restringida por variables de nuisance, adaptada de Wagner y Dray. Primero se ajustó una tendencia espacial, luego se randomizó el residuo en la base espectral de Moran. El mismo vector de signos espectrales se aplicó a los 19 genes en cada réplica. Así se preservaron, hasta tolerancia numérica, la tendencia ajustada, la potencia espectral residual de Moran y la covarianza residual entre genes. No se preservaron exactamente los marginales univariados de orden superior.

- **P1A:** 18 bloques donante × macrocompartimento; grafo unión de 5 vecinos más cercanos con peso inverso a la distancia; ajuste por intercepto y coordenadas MRI nativas $x,y,z$; 50,000 réplicas.
- **P1B:** seis donantes completos; grafo unión de 6 vecinos; ajuste por intercepto, $x,y,z$ y efectos fijos de estructura superior; 50,000 réplicas.
- **S1:** versión intracompartimental con pesos densos de distancia nativa; 10,000 réplicas.
- **S2:** versión de donante completo con pesos densos de distancia nativa; 10,000 réplicas.
- **S3A:** versión intracompartimental 5-NN con ejes estandarizados por donante; 10,000 réplicas.
- **S3B:** versión de donante completo 8-NN con ejes estandarizados; 10,000 réplicas. El valor $k=8$ fue el mínimo que conectó los seis grafos en el preflight de coordenadas, antes de producir endpoints.

Los valores empíricos unilaterales usaron $(1+b)/(1+N)$, donde $b$ fue el número de sustitutos mayores o iguales al observado. El criterio numérico de cada modelo exigió $p\leq0.05$ y observado estrictamente mayor que q95 lineal. El doble *gate* exigió que P1A y P1B pasaran.

## Resultados espaciales v1.1

| Modelo | Excedencias | $p$ empírico +1 | IC Wilson 95% de $b/N$ | Media nula | q95 | ¿Observado > q95? | Resultado |
|---|---:|---:|---:|---:|---:|:---:|:---:|
| P1A, bloques 5-NN | 2,558/50,000 | 0.0511789764 | [0.04926306, 0.05312590] | 0.16938577 | 0.34590643 | No | **No pasa** |
| P1B, donante completo 6-NN | 192/50,000 | 0.0038599228 | [0.00333468, 0.00442155] | 0.00687241 | 0.22149854 | Sí | Pasa |
| S1, bloques densos | 431/10,000 | 0.0431956804 | [0.03929201, 0.04725889] | 0.15796655 | 0.33684942 | Sí | Pasa |
| S2, donante completo denso | 50/10,000 | 0.0050994901 | [0.00379490, 0.00658526] | 0.00246019 | 0.22134503 | Sí | Pasa |
| S3A, bloques 5-NN, ejes estandarizados | 439/10,000 | 0.0439956004 | [0.04005666, 0.04809362] | 0.16394393 | 0.33713450 | Sí | Pasa |
| S3B, donante completo 8-NN, ejes estandarizados | 44/10,000 | 0.0044995500 | [0.00327944, 0.00590118] | 0.00692848 | 0.22179094 | Sí | Pasa |

P1A fue además marcado como Monte Carlo borderline porque su intervalo Wilson contiene 0.05. No obstante, la decisión no depende de esa etiqueta: falla tanto porque $p>0.05$ como porque $T<q95$. El *p* de intersección-unión fue 0.0511789764 y el doble *gate* no pasó.

El contraste entre familias de modelos es informativo. Las tres formulaciones de donante completo produjeron nulos centrados cerca de cero y cumplieron sus criterios numéricos individuales. Las formulaciones intracompartimentales produjeron medias nulas de 0.158–0.169 y resultados cercanos al umbral; la versión primaria 5-NN no pasó, mientras que las variantes densa y con ejes estandarizados sí pasaron por márgenes pequeños. Esto impide hablar de robustez uniforme frente a la autocorrelación espacial.

## Diagnóstico dejar-un-donante-fuera

El efecto observado permaneció positivo en las seis omisiones, con rango 0.27701754–0.42631579. En P1A, sólo 3/6 omisiones quedaron por encima de q95; los *p* empíricos oscilaron entre 0.02305954 y 0.18661627. En P1B, 6/6 omisiones pasaron, con $p$ entre 0.00099998 y 0.02953941.

Esto muestra que ningún donante determina el signo y que la familia de donante completo es estable a las omisiones. No convierte las omisiones solapadas en seis replicaciones independientes y tampoco rescata el *gate* P1A. Debe señalarse además que la cobertura local fue desigual: el bloque Cu/Gr del donante 12876 tuvo una sola muestra objetivo, aunque contó con 15 muestras hermanas y su omisión no cambió el signo agregado.

## Auditoría y reproducibilidad

La cadena v1.1 quedó fijada por el protocolo y cuatro enmiendas enlazadas por SHA-256. A01 sustituyó el modelo único inicial por el doble *gate* conservador antes de generar endpoints. A02 selló las convenciones numéricas y cambió únicamente S3B de 6-NN a 8-NN porque el grafo estandarizado de un donante no era conexo con $k\leq7$; esto se decidió con coordenadas, sin estadísticas sustitutas.

El primer intento A02 se detuvo antes de iniciar cualquier modelo espacial porque el replay residual histórico no superó la tolerancia de $10^{-12}$. No se produjo ni leyó un sustituto, un *p* o un cuantil v1.1. A03 restauró la aritmética `float32` exacta del linaje histórico para el replay observado, sin cambiar el modelo espacial. La diferencia máxima frente a v0.10 pasó de $1.2666\times10^{-7}$ a $2.2204\times10^{-16}$; la diferencia máxima del endpoint frente a v1.0 fue $8.3267\times10^{-17}$. A04 agregó dependencias y salidas de auditoría, sin cambiar operaciones inferenciales.

La validación final fue `PASS`:

- los 59 insumos del ledger se verificaron antes y después del endpoint;
- los 72 ajustes espaciales cumplieron conectividad, rango, varianza e invariantes;
- el error máximo de reconstrucción fue $5.3091\times10^{-13}$;
- el error máximo de ortogonalidad de la base fue $7.4096\times10^{-13}$;
- la diferencia máxima de covarianza residual fue $4.7518\times10^{-14}$;
- la diferencia máxima de correlación residual fue $1.0147\times10^{-13}$;
- la diferencia máxima de Moran $I$ residual fue $2.9410\times10^{-14}$;
- dos ejecuciones completas A04 produjeron 16 archivos idénticos byte por byte;
- el manifiesto de outputs del análisis v1.1-A04 contiene 15 entradas y tiene SHA-256 `eb90201948f278e15e42f4b70f0b42d27693649acb6f03573c47b1165e56cecd`;
- el resumen del análisis v1.1-A04 tiene SHA-256 `2455156e44c2e2946ee3e8d28c368ac7359a829afddcfe5c8a6577f907ed089e`; ninguno de los dos es el manifiesto de liberación A05.

La reproducibilidad computacional es una fortaleza real. No resuelve por sí misma el carácter post-descubrimiento ni la dependencia del modelo espacial.

## Interpretación y publicabilidad

**Diagnóstico: GO para un artículo exploratorio transparente y metodológicamente interesante; NO-GO, en el estado actual, para una afirmación biológica fuerte. Esta última sólo podría reconsiderarse tras una validación prospectiva independiente.**

El manuscrito puede aportar tres elementos valiosos: una estrategia anatómicamente restringida, nulos que aproximan de forma condicional partes prespecificadas del proceso de selección y una evaluación espacial conjunta que preserva coexpresión residual. El resultado cercano al umbral de P1A no debe ocultarse ni presentarse como “casi positivo”. Es un resultado negativo bajo una especificación primaria y convierte la dependencia del modelo en parte central de la contribución.

La narrativa editorial más sólida es:

1. El descubrimiento original de continuidad global fue negativo.
2. El patrón posdescubrimiento tuvo $T>0$ en cinco de seis donantes y ambas adyacencias positivas en seis de seis, pero sin una prueba de signo significativa; fue favorable frente a nulos condicionales de selección.
3. Los controles espaciales produjeron evidencia no uniforme: los tres modelos de donante completo cumplieron sus criterios numéricos individuales, mientras que los modelos intracompartimentales quedaron cerca del umbral y el componente primario P1A falló.
4. La especificidad de DCML no está demostrada y dos tríadas somatosensoriales con Sp5 fueron más fuertes.
5. La selección de sondas y la ausencia de un atlas independiente siguen siendo límites materiales.
6. GSE3526 recupera heterogeneidad del mismo linaje fuente, incluida una separación exploratoria completa por sexo en tres nodos, pero no aporta replicación; sus contrastes positivos reflejan menor disimilitud relativa y no coexpresión positiva general.

Un título provisional honesto sería:

> *Selection-aware and spatially constrained evaluation of a transcriptomic adjacency pattern across human somatosensory relay-region proxies*

## Próximos gates

1. **Confirmación no inspeccionada.** Localizar un conjunto RNA-seq, single-nucleus o espacial con cobertura suficiente de núcleos de columnas dorsales, tálamo somatosensorial y corteza postcentral. Si no existe equivalencia exacta, especificar una validación modular antes de examinar resultados.
2. **Nulo conjunto de selección y espacio.** Evaluar prospectivamente paneles que reproduzcan la selección y que también estén emparejados por coexpresión basal y autocorrelación espacial; no combinar aritméticamente los valores $p$ separados de v1.0 y v1.1.
3. **Sensibilidades espaciales adicionales.** Auditar variogramas y distancias de arista —P1A alcanzó 49.13 mm por el muestreo irregular— y congelar una sensibilidad que preserve marginales o variogramas. Descomponer también el endpoint espacial en adyacencia y extremos, con control de multiplicidad.
4. **Coexpresión basal fuera de los nodos.** Evaluar si el panel posee una coexpresión global excepcional que explique la adyacencia aun después de MSR. El análisis debe congelarse antes de abrir su endpoint.
5. **Sensibilidad de procesamiento AHBA.** Replicar con una canalización estandarizada, documentando normalización, selección de sondas, mapeo regional y hemisferio. Las decisiones de procesamiento pueden cambiar asociaciones de transcriptómica de imagen.
6. **Dependencia de plataforma, sonda, gen, metadatos y composición celular.** Priorizar RNA-seq; el diagnóstico de una sonda mostró estabilidad del signo, no estabilidad del umbral. En GSE3526 ya se ejecutó una sensibilidad post hoc del endpoint excluyendo RAB3GAP2/C7orf25; el dejar-un-gen-fuera se limitó al QC de la diferencia mujer–hombre del contraste de tres nodos y no es una sensibilidad general del endpoint. El siguiente nivel debe reprocesar CEL con una definición contemporánea y congelar un análisis que incorpore sexo y edad. Causa de muerte, intervalo post mortem, integridad de RNA y lote sólo podrán incorporarse si se recuperan de otra fuente. El patrón post hoc por sexo no debe presentarse como hipótesis confirmada. Ajustar o estratificar por composición celular a nivel de muestra cuando sea posible.
7. **Gobernanza y paquete público.** La autorización para analizar el extracto BIGE y publicar resultados agregados queda registrada. A05 superó una auditoría de liberación archivo por archivo que excluye la matriz interna completa y cualquier exportación reconstructiva; cada paquete futuro deberá repetirla. Verificar además los términos de Allen/GEO y separar reproducción académica de cualquier uso comercial.
8. **Congelación editorial.** Distinguir en texto, tablas y figuras descubrimiento, análisis prespecificados post-descubrimiento, sensibilidades y análisis post hoc. Publicar también el intento fallido pre-sustituto y las enmiendas técnicas.

## Límites inferenciales no negociables

- AHBA v1.0 y v1.1 reutilizan los datos en los que se reconoció la topología; no son confirmación independiente.
- Las regiones son proxies anatómicos muestreados: DTLv no equivale a una medición aislada de VPL.
- La similitud de expresión regional no demuestra conexiones axonales ni dirección de señal.
- La randomización conjunta preserva tendencias y segundos momentos residuales, no todos los marginales ni todas las propiedades biológicas de los mapas.
- Los contrastes celulares externos son proxies regionales amplios; no ajustan ni deconvolucionan la composición celular muestra por muestra en AHBA, por lo que permanece confusión residual posible.
- P1A falló; el resultado espacial es dependiente del modelo y no debe resumirse como un control espacial “positivo”.
- En la atribución neuronal v0.11, 30 de 50,000 conjuntos del nulo técnico igualaron o superaron el observado ($p=0.000620$), mientras que el comparador v1.0 condicionado por selección produjo $p=0.09677806$; HPA fue descriptivo. Ninguno constituye validación independiente ni específica de ruta.
- El extracto procesado y mantenido internamente de BIGE contiene agregados regionales, no observaciones individuales con covariables; pueden publicarse resultados agregados, pero no la matriz completa ni derivados reconstructivos.
- GSE3526 pertenece al mismo linaje fuente que el agregado BIGE, fue inspeccionado después de seleccionar el panel y no es una replicación independiente; sus fracciones de cola son descriptivas.
- El contraste GSE3526 de tres nodos separó post hoc a cuatro mujeres de cuatro hombres, con posibles confusores, y contiene sondas con anotación ambigua; no admite una conclusión sobre sexo ni una interpretación biológica firme sin nuevas sensibilidades.
- No existe evidencia aquí para un mecanismo de fibromialgia, una fisiopatología del dolor o un blanco/parámetro de estimulación transcraneal.

## Referencias metodológicas clave

1. Wagner HH, Dray S. Generating spatially constrained null models for irregularly spaced data using Moran spectral randomization methods. *Methods in Ecology and Evolution*. 2015;6:1169–1178. https://doi.org/10.1111/2041-210X.12407
2. Burt JB, Helmer M, Shinn M, Anticevic A, Murray JD. Generative modeling of brain maps with spatial autocorrelation. *NeuroImage*. 2020;220:117038. https://doi.org/10.1016/j.neuroimage.2020.117038
3. Markello RD, Misic B. Comparing spatial null models for brain maps. *NeuroImage*. 2021;236:118052. https://doi.org/10.1016/j.neuroimage.2021.118052
4. Markello RD, Arnatkeviciute A, Poline J-B, Fulcher BD, Fornito A, Misic B. Standardizing workflows in imaging transcriptomics with the abagen toolbox. *eLife*. 2021;10:e72129. https://doi.org/10.7554/eLife.72129
5. Siletti K, Hodge R, Mossi Albiach A, et al. Transcriptomic diversity of cell types across the adult human brain. *Science*. 2023;382(6667):eadd7046. https://doi.org/10.1126/science.add7046
6. Allen Institute. Allen Human Brain Atlas API documentation. https://brain-map.org/support/documentation/human-brain-atlas-api
7. Allen Institute. Terms of Use. https://alleninstitute.org/legal/terms-of-use
8. Allen Institute. Citation Policy. https://alleninstitute.org/legal/citation-policy
9. Lee J, Hever A, Willhite D, Zlotnik A, Hevezi P. Effects of RNA degradation on gene expression analysis of human postmortem tissues. *FASEB Journal*. 2005;19:1356–1358. https://doi.org/10.1096/fj.04-3552fje
10. Roth RB, Hevezi P, Lee J, et al. Gene expression analyses reveal molecular relationships among 20 regions of the human CNS. *Neurogenetics*. 2006;7:67–80. https://doi.org/10.1007/s10048-006-0032-6. GEO: GSE3526.
11. Ushach I, Burkhardt AM, Martinez C, et al. METEORIN-LIKE is a cytokine associated with barrier tissues and alternatively activated macrophages. *Clinical Immunology*. 2015;156:119–127. https://doi.org/10.1016/j.clim.2014.11.006
12. Hawrylycz MJ, Lein ES, Guillozet-Bongaarts AL, et al. An anatomically comprehensive atlas of the adult human brain transcriptome. *Nature*. 2012;489:391–399. https://doi.org/10.1038/nature11405
13. Human Protein Atlas. Brain data, version 25.1. Accessed 9 September 2026. https://www.proteinatlas.org/humanproteome/brain/data
14. National Center for Biotechnology Information. Gene Expression Omnibus platform GPL570: Affymetrix Human Genome U133 Plus 2.0 Array. Accessed 9 September 2026. https://www.ncbi.nlm.nih.gov/geo/query/acc.cgi?acc=GPL570
