## Supplementary Archive S1 for "An anatomy-constrained transcriptomic screen identifies a 19-gene panel with an ordered regional similarity pattern in the human somatosensory system": figure_captions_spatial_v1_1.md

# Draft spatial-robustness figure captions — route v1.1-A04

These figures report post-discovery robustness analyses in the same Allen Human Brain Atlas (AHBA) dataset in which the topology was identified. They are not independent confirmation. The analyses do not establish anatomical connectivity, direction, molecular flow, causality, route exclusivity, a pain or fibromyalgia mechanism, or a stimulation target.

## Figure 1. Frozen spatial-null double gate and sensitivity models

The frozen endpoint was the six-donor mean adjacency contrast, \(T=[(\rho_{Cu/Gr,DTLv}+\rho_{DTLv,postcentral})/2]-\rho_{Cu/Gr,postcentral}\), evaluated across the 19-gene panel (observed \(T=0.345175\)). (A) Black diamonds mark the observed statistic and colored ticks mark each spatial-null 95th percentile; connecting segments show the signed separation between them. Orange denotes blockwise models and blue denotes whole-donor models. The shaded rows are the two co-primary components of the prespecified intersection–union double gate. P1A placed the observed statistic just below its 95th percentile (\(q_{.95}=0.345906\), difference \(=-0.000731\)); P1B placed it well above its 95th percentile (\(q_{.95}=0.221499\), difference \(=0.123677\)). The four sensitivity thresholds were 0.336849 (S1), 0.221345 (S2), 0.337135 (S3A), and 0.221791 (S3B). (B) Points show plus-one empirical upper-tail \(p\) values and horizontal intervals show 95% Wilson intervals for the corresponding raw exceedance fractions, \(b/N\). The dashed line marks 0.05. P1A was Monte Carlo-borderline and failed the frozen numerical rule (2,558/50,000 exceedances; \(p=0.051179\); Wilson interval, 0.049263–0.053126), whereas P1B passed (192/50,000; \(p=0.003860\); 0.003335–0.004422). S1, S2, S3A, and S3B yielded \(p=0.043196\), 0.005099, 0.043996, and 0.004500, respectively. Because both co-primary components had to pass, \(p_{IUT}=\max(p_{P1A},p_{P1B})=0.051179\), the double gate did not pass, and the frozen classification was `WHOLE_ONLY_SPATIAL_ROBUSTNESS`. The fact that five of six displayed models passed their individual numerical criteria is not a voting rule and does not override the failed co-primary gate.

## Figure 2. Leave-one-donor-out sensitivity under the two primary spatial models

The endpoint was recomputed after omitting each AHBA donor. Diamonds mark observed leave-one-donor-out (LODO) values; colored ticks mark the corresponding spatial-null 95th percentiles. Filled diamonds indicate values above the threshold and open diamonds indicate values at or below it. All 12 observed LODO values were positive. Under P1A, 3 of 6 omissions exceeded the threshold (plus-one empirical upper-tail \(p\) range, 0.023060–0.186616); under P1B, 6 of 6 did so (\(p\) range, 0.001000–0.029539). The six five-donor subsets overlap extensively, so these counts are sensitivity diagnostics—not votes, independent experiments, or independent replications—and they do not alter the failed primary double gate.

## Abbreviations and visual encoding

- **AHBA:** Allen Human Brain Atlas.
- **Cu/Gr:** cuneate and gracile nuclei aggregate.
- **DTLv:** ventral division of the dorsal thalamus aggregate.
- **P1A:** blockwise union-5-nearest-neighbor spatial model with preserved linear trends.
- **P1B:** whole-donor union-6-nearest-neighbor spatial model with preserved linear trends and top-level-structure fixed effects.
- **S1/S2:** blockwise/whole-donor dense-distance sensitivity models.
- **S3A/S3B:** blockwise/whole-donor nearest-neighbor sensitivities after within-donor coordinate-axis standardization.
- **LODO:** leave one donor out.
- Orange identifies blockwise spatial models; blue identifies whole-donor spatial models. The palette is color-vision accessible.
