## Supplementary Archive S1 for "An anatomy-constrained transcriptomic screen identifies a 19-gene panel with an ordered regional similarity pattern in the human somatosensory system": plot_spatial_v1_1.py

```
#!/usr/bin/env python3
"""Create deterministic publication figures from the validated v1.1-A04 analysis.

The endpoint analysis is read-only.  Figure values are independently replayed
from the sealed spatial-null tables before plotting.  Outputs are written only
to the requested figure directory.
"""

from __future__ import annotations

import argparse
import hashlib
import json
import os
import platform
from pathlib import Path
from statistics import NormalDist
import tempfile
import xml.etree.ElementTree as ET

import matplotlib as mpl

mpl.use("Agg")
import matplotlib.pyplot as plt
from matplotlib import font_manager
from matplotlib.lines import Line2D
import numpy as np
import pandas as pd
from PIL import Image, __version__ as pillow_version


BLOCK_COLOR = "#D55E00"
WHOLE_COLOR = "#0072B2"
INK = "#161616"
MID_GRAY = "#6B6B6B"
LIGHT_GRAY = "#D5D5D5"
FAIL_COLOR = "#9B1B30"
TAIL_TOLERANCE = 1e-15

MODEL_CONFIG = (
    {
        "id": "P1A_BLOCKWISE_KNN5_TREND_PRESERVED",
        "label": "P1A  Blockwise kNN5",
        "scope": "block",
        "primary": True,
    },
    {
        "id": "P1B_WHOLE_DONOR_KNN6_FIXED_EFFECTS",
        "label": "P1B  Whole-donor kNN6",
        "scope": "whole",
        "primary": True,
    },
    {
        "id": "S1_BLOCKWISE_DENSE_NATIVE",
        "label": "S1  Blockwise dense",
        "scope": "block",
        "primary": False,
    },
    {
        "id": "S2_WHOLE_DONOR_DENSE_NATIVE",
        "label": "S2  Whole-donor dense",
        "scope": "whole",
        "primary": False,
    },
    {
        "id": "S3A_BLOCKWISE_KNN5_AXIS_Z",
        "label": "S3A  Blockwise kNN5 (scaled)",
        "scope": "block",
        "primary": False,
    },
    {
        "id": "S3B_WHOLE_DONOR_KNN8_AXIS_Z",
        "label": "S3B  Whole-donor kNN8 (scaled)",
        "scope": "whole",
        "primary": False,
    },
)

PRIMARY_MODELS = tuple(item["id"] for item in MODEL_CONFIG if item["primary"])
MODEL_IDS = tuple(item["id"] for item in MODEL_CONFIG)
Y_POSITIONS = np.array([6.25, 5.25, 3.65, 2.65, 1.65, 0.65], dtype=float)

REQUIRED_ANALYSIS_FILES = (
    "summary_v1_1.json",
    "spatial_model_specifications_v1_1.csv",
    "spatial_lodo_v1_1.csv",
    "observed_topology_donor_v1_1.csv",
    "spatial_analysis_hashes_v1_1.txt",
    *(f"spatial_null_{model_id}_v1_1.csv.gz" for model_id in MODEL_IDS),
)


def sha256(path: Path) -> str:
    digest = hashlib.sha256()
    with path.open("rb") as handle:
        for block in iter(lambda: handle.read(1024 * 1024), b""):
            digest.update(block)
    return digest.hexdigest()


def atomic_write_bytes(destination: Path, payload: bytes) -> None:
    descriptor, temporary_name = tempfile.mkstemp(
        prefix=f".{destination.stem}.", suffix=destination.suffix, dir=destination.parent
    )
    temporary_path = Path(temporary_name)
    try:
        with os.fdopen(descriptor, "wb") as handle:
            handle.write(payload)
            handle.flush()
            os.fsync(handle.fileno())
        os.replace(temporary_path, destination)
    finally:
        temporary_path.unlink(missing_ok=True)


def parse_analysis_hashes(manifest_path: Path) -> dict[str, str]:
    records: dict[str, str] = {}
    for line_number, raw_line in enumerate(
        manifest_path.read_text(encoding="utf-8").splitlines(), start=1
    ):
        if not raw_line.strip():
            continue
        parts = raw_line.split(maxsplit=1)
        if len(parts) != 2 or len(parts[0]) != 64:
            raise ValueError(f"Malformed analysis hash line {line_number}")
        digest, name = parts[0].lower(), parts[1].strip()
        if any(character not in "0123456789abcdef" for character in digest):
            raise ValueError(f"Invalid SHA-256 on analysis hash line {line_number}")
        if Path(name).name != name or name in records:
            raise ValueError(f"Unsafe or duplicate analysis filename: {name!r}")
        records[name] = digest
    if not records:
        raise ValueError("Empty analysis hash manifest")
    return records


def verify_validated_analysis(
    analysis_dir: Path, validation_path: Path
) -> tuple[dict[str, Path], dict, dict]:
    missing = [
        str(analysis_dir / name)
        for name in REQUIRED_ANALYSIS_FILES
        if not (analysis_dir / name).is_file()
    ]
    if missing:
        raise FileNotFoundError("Missing required analysis files: " + ", ".join(missing))

    hash_manifest = analysis_dir / "spatial_analysis_hashes_v1_1.txt"
    declared = parse_analysis_hashes(hash_manifest)
    for name, expected in declared.items():
        path = analysis_dir / name
        if not path.is_file():
            raise FileNotFoundError(f"Analysis manifest entry is missing: {path}")
        observed = sha256(path)
        if observed != expected:
            raise RuntimeError(f"Analysis hash mismatch for {name}: {observed} != {expected}")

    if not validation_path.is_file():
        raise FileNotFoundError(f"Missing spatial validation record: {validation_path}")
    validation = json.loads(validation_path.read_text(encoding="utf-8"))
    if validation.get("status") != "PASS":
        raise RuntimeError("The spatial analysis does not have PASS validation status")
    if validation.get("analysis_manifest_sha256") != sha256(hash_manifest):
        raise RuntimeError("Validation record does not match the analysis hash manifest")
    if not validation.get("independent_execution", {}).get("all_files_byte_identical"):
        raise RuntimeError("Independent analysis execution was not byte-identical")

    summary_path = analysis_dir / "summary_v1_1.json"
    if validation.get("summary_sha256") != sha256(summary_path):
        raise RuntimeError("Validation record does not match the v1.1 summary")
    summary = json.loads(summary_path.read_text(encoding="utf-8"))
    if summary.get("execution_revision") != "v1.1-A04_DEPENDENCY_AND_AUDIT_SEAL":
        raise ValueError("Unexpected spatial endpoint revision")
    if summary.get("interpretation_status") != (
        "POST_DISCOVERY_SPATIAL_ROBUSTNESS_NOT_INDEPENDENT_CONFIRMATION"
    ):
        raise ValueError("Unexpected interpretation status")
    if summary.get("classification") != "WHOLE_ONLY_SPATIAL_ROBUSTNESS":
        raise ValueError("Unexpected frozen v1.1 classification")
    if validation.get("classification") != summary["classification"]:
        raise RuntimeError("Validation classification does not match the summary")

    paths = {name: analysis_dir / name for name in declared}
    return paths, summary, validation


def wilson_interval(successes: int, total: int, alpha: float = 0.05) -> tuple[float, float]:
    z = NormalDist().inv_cdf(1.0 - alpha / 2.0)
    fraction = successes / total
    denominator = 1.0 + z * z / total
    center = (fraction + z * z / (2.0 * total)) / denominator
    half_width = (
        z
        * np.sqrt(fraction * (1.0 - fraction) / total + z * z / (4.0 * total * total))
        / denominator
    )
    return float(center - half_width), float(center + half_width)


def load_and_replay_results(
    analysis_dir: Path, summary: dict
) -> tuple[pd.DataFrame, dict[str, pd.DataFrame], pd.DataFrame]:
    specs = pd.read_csv(analysis_dir / "spatial_model_specifications_v1_1.csv")
    if tuple(specs["model_id"].astype(str)) != MODEL_IDS:
        raise ValueError("Unexpected spatial model order or membership")

    summary_specs = tuple(item["model_id"] for item in summary["model_specifications"])
    if summary_specs != MODEL_IDS:
        raise ValueError("Summary model order does not match the frozen figure order")

    observed = float(summary["observed"]["mean_adjacency_contrast"])
    model_rows: list[dict] = []
    nulls: dict[str, pd.DataFrame] = {}
    for config in MODEL_CONFIG:
        model_id = config["id"]
        model_summary = summary["models"][model_id]
        null_path = analysis_dir / f"spatial_null_{model_id}_v1_1.csv.gz"
        frame = pd.read_csv(null_path)
        draws = int(model_summary["null_n"])
        if len(frame) != draws or not np.array_equal(
            frame["draw_id"].to_numpy(int), np.arange(1, draws + 1)
        ):
            raise ValueError(f"{model_id}: invalid draw count or identifiers")
        values = frame["mean_adjacency_contrast"].to_numpy(float)
        if not np.isfinite(values).all():
            raise ValueError(f"{model_id}: nonfinite null statistic")
        q95 = float(np.quantile(values, 0.95, method="linear"))
        exceedances = int(np.count_nonzero(values >= observed - TAIL_TOLERANCE))
        p_value = (exceedances + 1) / (draws + 1)
        wilson_low, wilson_high = wilson_interval(exceedances, draws)
        numeric_pass = bool(p_value <= 0.05 and observed > q95)

        checks = (
            np.isclose(q95, float(model_summary["null_q95_linear"]), atol=1e-15, rtol=0),
            exceedances == int(model_summary["exceedance_count"]),
            np.isclose(
                p_value,
                float(model_summary["empirical_p_plus_one"]),
                atol=1e-15,
                rtol=0,
            ),
            np.allclose(
                [wilson_low, wilson_high],
                model_summary["exceedance_fraction_wilson95"],
                atol=1e-12,
                rtol=0,
            ),
            numeric_pass is bool(model_summary["numeric_pass"]),
            (observed > q95) is bool(model_summary["observed_beyond_q95"]),
        )
        if not all(checks):
            raise RuntimeError(f"{model_id}: replay does not match sealed summary")

        model_rows.append(
            {
                **config,
                "observed": observed,
                "q95": q95,
                "difference": observed - q95,
                "p": p_value,
                "wilson_low": wilson_low,
                "wilson_high": wilson_high,
                "numeric_pass": numeric_pass,
            }
        )
        nulls[model_id] = frame

    results = pd.DataFrame(model_rows)
    primary_p = float(results.loc[results["primary"], "p"].max())
    if not np.isclose(
        primary_p,
        float(summary["primary_double_gate"]["intersection_union_p_max"]),
        atol=1e-15,
        rtol=0,
    ):
        raise RuntimeError("Intersection–union p-value does not replay")
    if bool(results.loc[results["primary"], "numeric_pass"].all()):
        raise RuntimeError("Unexpected double-gate pass in sealed v1.1 results")

    lodo = pd.read_csv(analysis_dir / "spatial_lodo_v1_1.csv")
    expected_lodo_models = set(PRIMARY_MODELS)
    if len(lodo) != 12 or set(lodo["model_id"].astype(str)) != expected_lodo_models:
        raise ValueError("Unexpected LODO row count or model membership")
    donors = pd.read_csv(analysis_dir / "observed_topology_donor_v1_1.csv")
    if len(donors) != 6 or donors["donor_id"].duplicated().any():
        raise ValueError("Invalid observed donor table")
    donor_order = donors["donor_id"].astype(str).tolist()
    observed_by_donor = dict(
        zip(donor_order, donors["adjacency_contrast"].to_numpy(float), strict=True)
    )

    for model_id in PRIMARY_MODELS:
        frame = nulls[model_id]
        subset = lodo.loc[lodo["model_id"] == model_id].copy()
        subset["omitted_donor"] = subset["omitted_donor"].astype(str)
        subset = subset.set_index("omitted_donor").loc[donor_order]
        for donor_id, row in subset.iterrows():
            retained = [candidate for candidate in donor_order if candidate != donor_id]
            null_values = frame[
                [f"donor_{candidate}_adjacency_contrast" for candidate in retained]
            ].mean(axis=1).to_numpy(float)
            observed_lodo = float(np.mean([observed_by_donor[candidate] for candidate in retained]))
            q95_lodo = float(np.quantile(null_values, 0.95, method="linear"))
            exceedances = int(
                np.count_nonzero(null_values >= observed_lodo - TAIL_TOLERANCE)
            )
            p_lodo = (exceedances + 1) / (len(null_values) + 1)
            recorded_pass = bool(row["observed_beyond_q95"])
            if not (
                np.isclose(observed_lodo, float(row["observed"]), atol=1e-15, rtol=0)
                and np.isclose(q95_lodo, float(row["null_q95_linear"]), atol=1e-15, rtol=0)
                and exceedances == int(row["exceedance_count"])
                and np.isclose(
                    p_lodo, float(row["empirical_p_plus_one"]), atol=1e-15, rtol=0
                )
                and recorded_pass == bool(observed_lodo > q95_lodo)
            ):
                raise RuntimeError(f"{model_id}/{donor_id}: LODO replay mismatch")

    return results, nulls, lodo


def configure_matplotlib() -> None:
    mpl.rcParams.update(
        {
            "font.family": "DejaVu Sans",
            "font.size": 8.3,
            "axes.titlesize": 9.3,
            "axes.labelsize": 8.8,
            "axes.linewidth": 0.75,
            "axes.spines.top": False,
            "axes.spines.right": False,
            "axes.unicode_minus": True,
            "xtick.labelsize": 7.8,
            "ytick.labelsize": 7.8,
            "xtick.major.width": 0.7,
            "ytick.major.width": 0.7,
            "legend.fontsize": 7.5,
            "figure.dpi": 120,
            "savefig.dpi": 600,
            "savefig.facecolor": "white",
            "svg.hashsalt": "route-v1.1-A04-spatial-figures",
            "pdf.fonttype": 42,
            "ps.fonttype": 42,
        }
    )


def model_color(scope: str) -> str:
    return BLOCK_COLOR if scope == "block" else WHOLE_COLOR


def add_status_note(fig: mpl.figure.Figure) -> None:
    fig.text(
        0.99,
        0.943,
        "Post-discovery AHBA robustness analysis; not independent confirmation",
        ha="right",
        va="top",
        fontsize=6.4,
        color="#555555",
    )


def save_figure(fig: mpl.figure.Figure, base: Path) -> list[Path]:
    outputs: list[Path] = []
    metadata_by_suffix = {
        ".png": {"Software": "matplotlib; route v1.1-A04 spatial figures"},
        ".svg": {"Creator": "matplotlib; route v1.1-A04 spatial figures", "Date": None},
        ".pdf": {
            "Creator": "matplotlib; route v1.1-A04 spatial figures",
            "CreationDate": None,
            "ModDate": None,
        },
    }
    for suffix in (".png", ".svg", ".pdf"):
        destination = base.with_suffix(suffix)
        descriptor, temporary_name = tempfile.mkstemp(
            prefix=f".{base.name}.", suffix=suffix, dir=base.parent
        )
        os.close(descriptor)
        temporary_path = Path(temporary_name)
        try:
            fig.savefig(
                temporary_path,
                format=suffix.removeprefix("."),
                bbox_inches="tight",
                pad_inches=0.04,
                metadata=metadata_by_suffix[suffix],
            )
            if temporary_path.stat().st_size == 0:
                raise RuntimeError(f"Empty temporary figure output: {temporary_path}")
            os.replace(temporary_path, destination)
        finally:
            temporary_path.unlink(missing_ok=True)
        outputs.append(destination)
    plt.close(fig)
    return outputs


def verify_figure_file(path: Path) -> None:
    if not path.is_file() or path.stat().st_size == 0:
        raise RuntimeError(f"Missing or empty figure output: {path}")
    if path.suffix == ".png":
        with Image.open(path) as image:
            image.verify()
    elif path.suffix == ".svg":
        ET.parse(path)
    elif path.suffix == ".pdf":
        payload = path.read_bytes()
        if not payload.startswith(b"%PDF-") or b"%%EOF" not in payload[-1024:]:
            raise RuntimeError(f"Invalid PDF envelope: {path}")
    else:
        raise ValueError(f"Unsupported figure extension: {path.suffix}")


def decorate_model_axis(axis: mpl.axes.Axes, show_labels: bool) -> None:
    axis.axhspan(4.75, 6.65, color="#F2F2F2", zorder=0)
    axis.axhline(4.48, color="#B8B8B8", linewidth=0.65, zorder=0)
    axis.set_ylim(0.15, 7.08)
    axis.set_yticks(Y_POSITIONS)
    if show_labels:
        axis.set_yticklabels([item["label"] for item in MODEL_CONFIG])
        for index, label in enumerate(axis.get_yticklabels()):
            if index < 2:
                label.set_fontweight("bold")
        axis.tick_params(axis="y", length=0, pad=4)
    else:
        axis.tick_params(axis="y", labelleft=False, length=0)
    axis.text(
        0.01,
        0.985,
        "CO-PRIMARY DOUBLE GATE",
        transform=axis.transAxes,
        ha="left",
        va="top",
        fontsize=6.5,
        fontweight="bold",
        color="#555555",
    )
    axis.text(
        0.01,
        0.611,
        "SPATIAL SENSITIVITIES",
        transform=axis.transAxes,
        ha="left",
        va="top",
        fontsize=6.5,
        fontweight="bold",
        color="#555555",
    )
    axis.grid(axis="x", color=LIGHT_GRAY, linewidth=0.55, alpha=0.72)
    axis.set_axisbelow(True)


def figure_1_double_gate(output_dir: Path, results: pd.DataFrame) -> list[Path]:
    fig, axes = plt.subplots(
        1,
        2,
        figsize=(7.25, 4.55),
        sharey=True,
        gridspec_kw={"width_ratios": [1.05, 1.0]},
    )
    decorate_model_axis(axes[0], show_labels=True)
    decorate_model_axis(axes[1], show_labels=False)

    observed = float(results["observed"].iloc[0])
    for y, (_, row) in zip(Y_POSITIONS, results.iterrows(), strict=True):
        color = model_color(str(row["scope"]))
        q95 = float(row["q95"])
        axes[0].plot(
            [q95, observed],
            [y, y],
            color=color,
            linewidth=1.7,
            solid_capstyle="round",
            zorder=2,
        )
        axes[0].scatter(
            [q95],
            [y],
            marker="|",
            s=165,
            linewidth=2.2,
            color=color,
            zorder=3,
        )
        axes[0].scatter(
            [observed],
            [y],
            marker="D",
            s=34,
            facecolor=INK,
            edgecolor="white",
            linewidth=0.45,
            zorder=4,
        )
        status = "PASS" if bool(row["numeric_pass"]) else "FAIL"
        axes[0].text(
            0.985,
            y,
            status,
            transform=axes[0].get_yaxis_transform(),
            ha="right",
            va="center",
            fontsize=6.6,
            fontweight="bold" if status == "FAIL" else "normal",
            color=FAIL_COLOR if status == "FAIL" else "#444444",
        )

        p_value = float(row["p"])
        low = float(row["wilson_low"])
        high = float(row["wilson_high"])
        marker = "D" if bool(row["primary"]) else "o"
        passed = bool(row["numeric_pass"])
        axes[1].errorbar(
            [p_value],
            [y],
            xerr=np.array([[p_value - low], [high - p_value]]),
            fmt=marker,
            markersize=5.4,
            markerfacecolor=color if passed else "white",
            markeredgecolor=color,
            markeredgewidth=1.25,
            ecolor=color,
            elinewidth=1.25,
            capsize=2.3,
            capthick=1.0,
            zorder=3,
        )
        axes[1].text(
            0.985,
            y,
            f"{p_value:.4f}",
            transform=axes[1].get_yaxis_transform(),
            ha="right",
            va="center",
            fontsize=6.7,
            color=FAIL_COLOR if not passed else "#444444",
            fontweight="bold" if not passed else "normal",
        )

    axes[0].set_xlim(0.205, 0.395)
    axes[0].set_xticks([0.22, 0.26, 0.30, 0.34, 0.38])
    axes[0].set_xlabel(r"Adjacency contrast, $T$")
    axes[0].set_title("A   Observed T vs null q95", loc="left", fontweight="bold")

    axes[1].axvline(0.05, color=MID_GRAY, linestyle=(0, (4, 2)), linewidth=1.15, zorder=1)
    axes[1].set_xlim(-0.0015, 0.066)
    axes[1].set_xticks([0.00, 0.01, 0.02, 0.03, 0.04, 0.05, 0.06])
    axes[1].set_xlabel(r"Plus-one empirical $p$")
    axes[1].set_title("B   Empirical upper-tail p", loc="left", fontweight="bold")

    fig.legend(
        handles=[
            Line2D(
                [0], [0], marker="D", color="none", markerfacecolor=INK,
                markeredgecolor="white", markersize=5.8, label="Observed T",
            ),
            Line2D(
                [0], [0], marker="|", color=MID_GRAY, linestyle="none",
                markersize=10, markeredgewidth=2.2, label="Null 95th percentile",
            ),
            Line2D([0], [0], color=BLOCK_COLOR, lw=2, label="Blockwise model"),
            Line2D([0], [0], color=WHOLE_COLOR, lw=2, label="Whole-donor model"),
        ],
        frameon=False,
        loc="lower center",
        bbox_to_anchor=(0.55, 0.112),
        ncol=4,
        columnspacing=1.15,
        handletextpad=0.45,
        handlelength=1.9,
    )
    fig.text(
        0.55,
        0.025,
        "Primary intersection–union decision: P1A FAIL + P1B PASS  →  DOUBLE GATE NOT PASSED\n"
        "Frozen classification: WHOLE-ONLY SPATIAL ROBUSTNESS",
        ha="center",
        va="bottom",
        fontsize=7.4,
        fontweight="bold",
        linespacing=1.35,
        color="#303030",
        bbox={"boxstyle": "round,pad=0.35", "fc": "#F3F3F3", "ec": "#A8A8A8", "lw": 0.7},
    )
    fig.suptitle(
        "Frozen spatial-null double gate and sensitivity models",
        x=0.02,
        y=0.992,
        ha="left",
        fontsize=11,
        fontweight="bold",
    )
    add_status_note(fig)
    fig.subplots_adjust(left=0.305, right=0.995, top=0.855, bottom=0.245, wspace=0.18)
    return save_figure(fig, output_dir / "figure_1_spatial_double_gate_v1_1")


def figure_2_lodo(output_dir: Path, lodo: pd.DataFrame) -> list[Path]:
    donor_order = (
        lodo.loc[lodo["model_id"] == PRIMARY_MODELS[0], "omitted_donor"]
        .astype(str)
        .tolist()
    )
    if len(donor_order) != 6 or set(lodo["omitted_donor"].astype(str)) != set(donor_order):
        raise ValueError("Unexpected donor membership in LODO results")

    titles = {
        PRIMARY_MODELS[0]: "A   P1A blockwise kNN5",
        PRIMARY_MODELS[1]: "B   P1B whole-donor kNN6",
    }
    fig, axes = plt.subplots(1, 2, figsize=(7.25, 3.55), sharex=True, sharey=True)
    for axis, model_id in zip(axes, PRIMARY_MODELS, strict=True):
        subset = lodo.loc[lodo["model_id"] == model_id].copy()
        subset["omitted_donor"] = subset["omitted_donor"].astype(str)
        subset = subset.set_index("omitted_donor").loc[donor_order]
        y = np.arange(len(donor_order), dtype=float)[::-1]
        observed = subset["observed"].to_numpy(float)
        q95 = subset["null_q95_linear"].to_numpy(float)
        beyond = observed > q95
        recorded = subset["observed_beyond_q95"].astype(bool).to_numpy()
        if not np.array_equal(beyond, recorded):
            raise RuntimeError(f"{model_id}: LODO threshold indicators do not replay")
        color = BLOCK_COLOR if model_id == PRIMARY_MODELS[0] else WHOLE_COLOR

        for yi, observed_value, threshold in zip(y, observed, q95, strict=True):
            axis.plot(
                [min(observed_value, threshold), max(observed_value, threshold)],
                [yi, yi],
                color="#B5B5B5",
                linewidth=1.1,
                zorder=1,
            )
        axis.scatter(
            q95,
            y,
            marker="|",
            s=170,
            linewidth=2.2,
            color=color,
            zorder=2,
        )
        axis.scatter(
            observed[beyond],
            y[beyond],
            marker="D",
            s=36,
            facecolor=INK,
            edgecolor=INK,
            linewidth=0.8,
            zorder=3,
        )
        axis.scatter(
            observed[~beyond],
            y[~beyond],
            marker="D",
            s=36,
            facecolor="white",
            edgecolor=INK,
            linewidth=1.0,
            zorder=3,
        )
        axis.set_title(titles[model_id], loc="left", fontweight="bold")
        axis.set_xlabel(r"LODO adjacency contrast, $T$")
        axis.set_xlim(0.215, 0.445)
        axis.set_xticks([0.22, 0.26, 0.30, 0.34, 0.38, 0.42])
        axis.grid(axis="x", color=LIGHT_GRAY, linewidth=0.55, alpha=0.72)
        axis.set_axisbelow(True)
        axis.text(
            0.97,
            0.06,
            f"{int(beyond.sum())}/6 above null q95",
            transform=axis.transAxes,
            ha="right",
            va="bottom",
            fontsize=7.5,
            fontweight="bold",
            color=color,
        )

    axes[0].set_yticks(np.arange(len(donor_order), dtype=float)[::-1], donor_order)
    axes[0].set_ylabel("Omitted AHBA donor")
    axes[0].tick_params(axis="y", length=0)
    axes[1].tick_params(axis="y", labelleft=False, length=0)
    axes[0].legend(
        handles=[
            Line2D(
                [0], [0], marker="D", color="none", markerfacecolor=INK,
                markeredgecolor=INK, markersize=5.5, label="Observed above q95",
            ),
            Line2D(
                [0], [0], marker="D", color="none", markerfacecolor="white",
                markeredgecolor=INK, markersize=5.5, label="Observed at/below q95",
            ),
            Line2D(
                [0], [0], marker="|", color=MID_GRAY, linestyle="none",
                markersize=10, markeredgewidth=2.2, label="Spatial-null q95",
            ),
        ],
        frameon=False,
        loc="upper left",
        bbox_to_anchor=(0.0, -0.235),
        ncol=3,
        columnspacing=1.2,
        handletextpad=0.45,
    )
    fig.text(
        0.99,
        0.018,
        "Diagnostic only: the six five-donor subsets overlap; counts are not votes or independent replications.",
        ha="right",
        va="bottom",
        fontsize=6.6,
        color="#555555",
    )
    fig.suptitle(
        "Leave-one-donor-out sensitivity under the primary spatial models",
        x=0.02,
        y=0.99,
        ha="left",
        fontsize=11,
        fontweight="bold",
    )
    add_status_note(fig)
    fig.subplots_adjust(left=0.115, right=0.995, top=0.84, bottom=0.285, wspace=0.14)
    return save_figure(fig, output_dir / "figure_2_spatial_lodo_v1_1")


def main() -> None:
    parser = argparse.ArgumentParser(description=__doc__)
    parser.add_argument(
        "--analysis-dir",
        type=Path,
        default=Path(__file__).resolve().parents[1] / "analysis",
    )
    parser.add_argument(
        "--validation-file",
        type=Path,
        default=Path(__file__).resolve().parents[1]
        / "verification"
        / "spatial_validation_v1_1.json",
    )
    parser.add_argument(
        "--output-dir",
        type=Path,
        default=Path(__file__).resolve().parents[1] / "figures",
    )
    parser.add_argument(
        "--caption-source",
        type=Path,
        default=Path(__file__).resolve().with_name("figure_captions_spatial_v1_1.md"),
    )
    args = parser.parse_args()
    analysis_dir = args.analysis_dir.resolve()
    validation_path = args.validation_file.resolve()
    output_dir = args.output_dir.resolve()
    caption_source = args.caption_source.resolve()
    if not caption_source.is_file():
        raise FileNotFoundError(f"Missing caption source: {caption_source}")
    output_dir.mkdir(parents=True, exist_ok=True)

    analysis_paths, summary, validation = verify_validated_analysis(
        analysis_dir, validation_path
    )
    results, _nulls, lodo = load_and_replay_results(analysis_dir, summary)
    configure_matplotlib()

    outputs: list[Path] = []
    outputs += figure_1_double_gate(output_dir, results)
    outputs += figure_2_lodo(output_dir, lodo)
    for output in outputs:
        verify_figure_file(output)

    caption_path = output_dir / "figure_captions_spatial_v1_1.md"
    if caption_source != caption_path.resolve():
        atomic_write_bytes(caption_path, caption_source.read_bytes())

    manifest = {
        "version": "v1.1",
        "execution_revision": summary["execution_revision"],
        "analysis_validation": {
            "status": validation["status"],
            "analysis_manifest_sha256": validation["analysis_manifest_sha256"],
            "summary_sha256": validation["summary_sha256"],
            "independent_execution_byte_identical": validation["independent_execution"][
                "all_files_byte_identical"
            ],
            "validation_record_path": "verification/spatial_validation_v1_1.json",
            "validation_record_sha256": sha256(validation_path),
        },
        "classification": summary["classification"],
        "interpretation_status": summary["interpretation_status"],
        "claim_guardrail": summary["claim_guardrail"],
        "primary_double_gate": summary["primary_double_gate"],
        "generator_file": {
            "path": "scripts/plot_spatial_v1_1.py",
            "bytes": Path(__file__).resolve().stat().st_size,
            "sha256": sha256(Path(__file__).resolve()),
        },
        "caption_source": {
            "path": "scripts/figure_captions_spatial_v1_1.md",
            "bytes": caption_source.stat().st_size,
            "sha256": sha256(caption_source),
        },
        "runtime": {
            "python": platform.python_version(),
            "matplotlib": mpl.__version__,
            "numpy": np.__version__,
            "pandas": pd.__version__,
            "pillow": pillow_version,
            "backend": str(mpl.get_backend()),
            "font_family": "DejaVu Sans",
            "font_file_sha256": sha256(Path(font_manager.findfont("DejaVu Sans"))),
        },
        "validated_analysis_files": {
            name: {"bytes": path.stat().st_size, "sha256": sha256(path)}
            for name, path in sorted(analysis_paths.items())
        },
        "output_files": {
            output.name: {"bytes": output.stat().st_size, "sha256": sha256(output)}
            for output in sorted(outputs)
        },
        "caption_file": {
            "path": caption_path.name,
            "bytes": caption_path.stat().st_size,
            "sha256": sha256(caption_path),
        },
    }
    manifest_path = output_dir / "figure_manifest_spatial_v1_1.json"
    atomic_write_bytes(
        manifest_path,
        (json.dumps(manifest, indent=2, sort_keys=True) + "\n").encode("utf-8"),
    )
    print(f"Generated and verified {len(outputs)} figure files in {output_dir}")
    print(f"Manifest: {manifest_path} ({sha256(manifest_path)})")


if __name__ == "__main__":
    main()
```
