## Supplementary Archive S1 for "An anatomy-constrained transcriptomic screen identifies a 19-gene panel with an ordered regional similarity pattern in the human somatosensory system": verify_spatial_outputs_v1_1.py

```
#!/usr/bin/env python3
"""Independently validate the promoted v1.1 spatial-analysis package."""

from __future__ import annotations

import hashlib
import json
import math
from pathlib import Path
from typing import Any

import numpy as np
import pandas as pd


HERE = Path(__file__).resolve().parents[1]
PROJECT = HERE.parents[1]
ANALYSIS = HERE / "analysis"
VERIFICATION = HERE / "verification"
REFERENCE_RUN = HERE / "verification_runs" / "spatial_A04_run1"
SUMMARY_PATH = ANALYSIS / "summary_v1_1.json"
MANIFEST_PATH = ANALYSIS / "spatial_analysis_hashes_v1_1.txt"
DONORS = [9861, 10021, 12876, 14380, 15496, 15697]
FLOAT_TOLERANCE = 1e-12
TAIL_TOLERANCE = 1e-15


def sha256(path: Path) -> str:
    digest = hashlib.sha256()
    with path.open("rb") as handle:
        for block in iter(lambda: handle.read(1024 * 1024), b""):
            digest.update(block)
    return digest.hexdigest()


def assert_close(observed: float, expected: float, label: str) -> None:
    if not math.isclose(float(observed), float(expected), rel_tol=0.0, abs_tol=FLOAT_TOLERANCE):
        raise RuntimeError(f"{label}: {observed!r} != {expected!r}")


def wilson_interval(successes: int, total: int) -> tuple[float, float]:
    z = 1.959963984540054
    proportion = successes / total
    denominator = 1 + z * z / total
    centre = (proportion + z * z / (2 * total)) / denominator
    half = z * math.sqrt(
        proportion * (1 - proportion) / total + z * z / (4 * total * total)
    ) / denominator
    return centre - half, centre + half


def parse_manifest() -> dict[str, str]:
    entries: dict[str, str] = {}
    for line in MANIFEST_PATH.read_text(encoding="utf-8").splitlines():
        digest, filename = line.split("  ", 1)
        if filename in entries or len(digest) != 64:
            raise RuntimeError("Malformed or duplicate output-manifest entry")
        entries[filename] = digest
    actual = {path.name for path in ANALYSIS.iterdir() if path.is_file()}
    expected_actual = set(entries) | {MANIFEST_PATH.name}
    if actual != expected_actual:
        raise RuntimeError(f"Analysis file set differs from manifest: {actual ^ expected_actual}")
    for filename, digest in entries.items():
        if sha256(ANALYSIS / filename) != digest:
            raise RuntimeError(f"Output hash mismatch: {filename}")
    return entries


def validate_model(
    model: dict[str, Any], summary: dict[str, Any], observed: float
) -> tuple[pd.DataFrame, dict[str, Any]]:
    model_id = model["model_id"]
    path = ANALYSIS / f"spatial_null_{model_id}_v1_1.csv.gz"
    frame = pd.read_csv(path, float_precision="round_trip")
    draws = int(model["draws"])
    expected_columns = ["draw_id", "mean_adjacency_contrast"] + [
        f"donor_{donor}_adjacency_contrast" for donor in DONORS
    ]
    if frame.columns.tolist() != expected_columns:
        raise RuntimeError(f"Unexpected null columns for {model_id}")
    if len(frame) != draws or frame["draw_id"].tolist() != list(range(1, draws + 1)):
        raise RuntimeError(f"Unexpected draw sequence for {model_id}")
    values = frame["mean_adjacency_contrast"].to_numpy(float)
    donor_values = frame[expected_columns[2:]].to_numpy(float)
    if not np.isfinite(values).all() or not np.isfinite(donor_values).all():
        raise RuntimeError(f"Nonfinite null value for {model_id}")
    if not np.allclose(values, donor_values.mean(axis=1), atol=3e-16, rtol=0.0):
        raise RuntimeError(f"Aggregate is not donor mean for {model_id}")
    exceedances = int(np.sum(values >= observed - TAIL_TOLERANCE))
    empirical_p = (1 + exceedances) / (1 + draws)
    q95 = float(np.quantile(values, 0.95, method="linear"))
    low, high = wilson_interval(exceedances, draws)
    expected = summary["models"][model_id]
    exact_integer_checks = {
        "null_n": draws,
        "exceedance_count": exceedances,
        "unique_statistic_count": int(np.unique(values).size),
    }
    for key, value in exact_integer_checks.items():
        if int(expected[key]) != value:
            raise RuntimeError(f"{model_id}/{key} mismatch")
    float_checks = {
        "observed": observed,
        "null_mean": float(values.mean()),
        "null_sd_ddof1": float(values.std(ddof=1)),
        "null_variance_ddof1": float(values.var(ddof=1)),
        "null_q95_linear": q95,
        "empirical_p_plus_one": empirical_p,
        "exceedance_fraction": exceedances / draws,
    }
    for key, value in float_checks.items():
        assert_close(expected[key], value, f"{model_id}/{key}")
    assert_close(expected["exceedance_fraction_wilson95"][0], low, f"{model_id}/Wilson low")
    assert_close(expected["exceedance_fraction_wilson95"][1], high, f"{model_id}/Wilson high")
    donor_sd = donor_values.std(axis=0, ddof=1)
    for index, donor in enumerate(DONORS):
        assert_close(
            expected["donor_contrast_sd_ddof1"][str(donor)],
            donor_sd[index],
            f"{model_id}/donor SD/{donor}",
        )
    pass_rule = bool(empirical_p <= 0.05 and observed > q95)
    borderline = bool(low <= 0.05 <= high)
    if bool(expected["numeric_pass"]) != pass_rule:
        raise RuntimeError(f"Pass classification mismatch for {model_id}")
    if bool(expected["monte_carlo_borderline_at_0_05"]) != borderline:
        raise RuntimeError(f"Wilson classification mismatch for {model_id}")
    if values.var(ddof=1) <= 0 or np.unique(values).size <= 1 or np.any(donor_sd <= 0):
        raise RuntimeError(f"Endpoint-diversity gate failed for {model_id}")
    return frame, {
        "model_id": model_id,
        "draws": draws,
        "exceedances": exceedances,
        "empirical_p_plus_one": empirical_p,
        "q95": q95,
        "numeric_pass": pass_rule,
        "wilson95": [low, high],
    }


def validate_lodo(
    lodo: pd.DataFrame,
    model_frames: dict[str, pd.DataFrame],
    observed_donor: pd.DataFrame,
) -> None:
    primary_ids = [
        "P1A_BLOCKWISE_KNN5_TREND_PRESERVED",
        "P1B_WHOLE_DONOR_KNN6_FIXED_EFFECTS",
    ]
    if len(lodo) != 12 or set(lodo["model_id"]) != set(primary_ids):
        raise RuntimeError("Unexpected LODO rows")
    observed_by_donor = observed_donor.set_index("donor_id")["adjacency_contrast"]
    for model_id in primary_ids:
        frame = model_frames[model_id]
        for omitted in DONORS:
            row = lodo[(lodo["model_id"] == model_id) & (lodo["omitted_donor"] == omitted)]
            if len(row) != 1:
                raise RuntimeError("Missing or duplicate LODO row")
            row = row.iloc[0]
            retained = [donor for donor in DONORS if donor != omitted]
            null = frame[[f"donor_{donor}_adjacency_contrast" for donor in retained]].mean(axis=1).to_numpy(float)
            observed = float(observed_by_donor.loc[retained].mean())
            exceedances = int(np.sum(null >= observed - TAIL_TOLERANCE))
            p_value = (1 + exceedances) / (1 + len(null))
            q95 = float(np.quantile(null, 0.95, method="linear"))
            assert_close(row["observed"], observed, "LODO observed")
            assert_close(row["empirical_p_plus_one"], p_value, "LODO p")
            assert_close(row["null_q95_linear"], q95, "LODO q95")
            if int(row["exceedance_count"]) != exceedances:
                raise RuntimeError("LODO exceedance mismatch")


def validate_qc() -> dict[str, float]:
    qc = pd.read_csv(ANALYSIS / "spatial_block_qc_v1_1.csv")
    invariant = pd.read_csv(ANALYSIS / "spatial_invariant_audit_v1_1.csv")
    if len(qc) != 72 or len(invariant) != 72:
        raise RuntimeError("Unexpected spatial-fit QC row count")
    if not qc["connected"].all() or not invariant["connected"].all():
        raise RuntimeError("Disconnected fitted spatial graph")
    gates = {
        "max_basis_orthogonality_error": (qc, 1e-9),
        "max_design_orthogonality_error": (qc, 1e-9),
        "max_all_plus_reconstruction_error": (qc, 1e-9),
        "max_fitted_nuisance_difference": (invariant, 1e-9),
        "max_residual_sd_difference_ddof1": (invariant, 1e-8),
        "max_residual_gene_covariance_difference_ddof1": (invariant, 1e-8),
        "max_residual_gene_correlation_difference": (invariant, 1e-8),
        "max_residual_moran_i_difference": (invariant, 1e-8),
    }
    maxima: dict[str, float] = {}
    for column, (frame, limit) in gates.items():
        maximum = float(frame[column].max())
        if not np.isfinite(maximum) or maximum > limit:
            raise RuntimeError(f"QC gate failed: {column}")
        maxima[column] = maximum
    whole_projection = invariant[invariant["scope"] == "whole"][
        "targeted_projection_max_difference"
    ]
    if whole_projection.isna().any() or float(whole_projection.max()) > 1e-9:
        raise RuntimeError("Whole-donor targeted-projection gate failed")
    if (qc["minimum_residual_gene_variance_ddof1"] <= 0).any():
        raise RuntimeError("Nonpositive residual variance in QC")
    return maxima


def validate_ledger() -> int:
    ledger = pd.read_csv(
        ANALYSIS / "verified_spatial_inputs_v1_1.csv", float_precision="round_trip"
    )
    if (
        len(ledger) != 59
        or ledger["source_id"].nunique() != 59
        or int(ledger["source_id"].str.startswith("raw_probe::").sum()) != 44
    ):
        raise RuntimeError("Unexpected verified-input ledger")
    if not ledger["verified_before_endpoint"].all() or not ledger["verified_after_endpoint"].all():
        raise RuntimeError("Input ledger contains an unverified item")
    for row in ledger.itertuples(index=False):
        path = PROJECT / row.path
        if not path.is_file() or path.stat().st_size != int(row.bytes) or sha256(path) != row.sha256:
            raise RuntimeError(f"Current input differs from ledger: {row.source_id}")
    return len(ledger)


def validate_independent_execution() -> dict[str, Any]:
    if not REFERENCE_RUN.is_dir():
        raise RuntimeError("Preserved first A04 execution is missing")
    reference_files = sorted(path.name for path in REFERENCE_RUN.iterdir() if path.is_file())
    final_files = sorted(path.name for path in ANALYSIS.iterdir() if path.is_file())
    if reference_files != final_files:
        raise RuntimeError("Independent execution file sets differ")
    rows = []
    for filename in final_files:
        reference_hash = sha256(REFERENCE_RUN / filename)
        final_hash = sha256(ANALYSIS / filename)
        if reference_hash != final_hash:
            raise RuntimeError(f"Independent execution differs: {filename}")
        rows.append({"file": filename, "sha256": final_hash})
    return {
        "preserved_first_run": str(REFERENCE_RUN.relative_to(PROJECT)),
        "final_second_run": str(ANALYSIS.relative_to(PROJECT)),
        "file_count": len(rows),
        "all_files_byte_identical": True,
        "files": rows,
    }


def main() -> int:
    manifest = parse_manifest()
    summary = json.loads(SUMMARY_PATH.read_text(encoding="utf-8"))
    observed = float(summary["observed"]["mean_adjacency_contrast"])
    observed_donor = pd.read_csv(
        ANALYSIS / "observed_topology_donor_v1_1.csv", float_precision="round_trip"
    )
    if len(observed_donor) != 6 or observed_donor["donor_id"].astype(int).tolist() != DONORS:
        raise RuntimeError("Unexpected observed donor table")
    assert_close(observed_donor["adjacency_contrast"].mean(), observed, "Observed aggregate")
    model_frames: dict[str, pd.DataFrame] = {}
    model_checks = []
    for model in summary["model_specifications"]:
        frame, check = validate_model(model, summary, observed)
        model_frames[model["model_id"]] = frame
        model_checks.append(check)
    p1a = summary["models"]["P1A_BLOCKWISE_KNN5_TREND_PRESERVED"]
    p1b = summary["models"]["P1B_WHOLE_DONOR_KNN6_FIXED_EFFECTS"]
    expected_classification = (
        "SPATIAL_ROBUST_DOUBLE_GATE"
        if p1a["numeric_pass"] and p1b["numeric_pass"]
        else "BLOCK_ONLY_SPATIAL_ROBUSTNESS"
        if p1a["numeric_pass"]
        else "WHOLE_ONLY_SPATIAL_ROBUSTNESS"
        if p1b["numeric_pass"]
        else "SPATIAL_NOT_ROBUST"
    )
    if summary["classification"] != expected_classification:
        raise RuntimeError("Primary spatial classification mismatch")
    assert_close(
        summary["primary_double_gate"]["intersection_union_p_max"],
        max(p1a["empirical_p_plus_one"], p1b["empirical_p_plus_one"]),
        "Intersection-union p",
    )
    validate_lodo(
        pd.read_csv(ANALYSIS / "spatial_lodo_v1_1.csv"),
        model_frames,
        observed_donor,
    )
    counts = pd.read_csv(ANALYSIS / "spatial_target_sample_counts_v1_1.csv")
    if len(counts) != 18 or (counts[["target_sample_count", "sibling_sample_count"]] <= 0).any().any():
        raise RuntimeError("Invalid target/sibling count table")
    qc_maxima = validate_qc()
    ledger_rows = validate_ledger()
    report = {
        "status": "PASS",
        "analysis_manifest_entry_count": len(manifest),
        "verified_input_ledger_rows": ledger_rows,
        "observed": observed,
        "classification": summary["classification"],
        "intersection_union_p_max": summary["primary_double_gate"][
            "intersection_union_p_max"
        ],
        "model_checks": model_checks,
        "qc_maxima": qc_maxima,
        "summary_sha256": sha256(SUMMARY_PATH),
        "analysis_manifest_sha256": sha256(MANIFEST_PATH),
        "independent_execution": validate_independent_execution(),
        "validator_sha256": sha256(Path(__file__).resolve()),
    }
    VERIFICATION.mkdir(parents=True, exist_ok=True)
    output = VERIFICATION / "spatial_validation_v1_1.json"
    output.write_text(json.dumps(report, indent=2, sort_keys=True) + "\n", encoding="utf-8")
    print(json.dumps(report, indent=2, sort_keys=True))
    return 0


if __name__ == "__main__":
    raise SystemExit(main())
```
