## Supplementary Archive S1 for "An anatomy-constrained transcriptomic screen identifies a 19-gene panel with an ordered regional similarity pattern in the human somatosensory system": independent_validation_feasibility_v1_2.md

# Independent validation feasibility assessment v1.2

**Project:** anatomy-constrained transcriptomic adjacency across human somatosensory relay-region proxies

**Panel:** 19 genes associated with the DCML candidate set

**Assessment date:** 2026-09-09

**Document status:** feasibility and protocol-design record; no new endpoint in a proposed independent resource was executed or inspected for this assessment. Previously inspected GSE3526 source-lineage results are recorded below as contamination, not validation.

## Executive conclusion

No single public adult-human transcriptomic resource was identified that simultaneously provides:

1. isolated cuneate and/or gracile nuclei;
2. exact ventral posterolateral thalamic nucleus (VPL);
3. postcentral gyrus or primary somatosensory cortex (S1);
4. all three regions in multiple matched donors;
5. a dataset that is both independent of AHBA and completely uninspected for this 19-gene panel.

The defensible v1.2 strategy is therefore **prospectively frozen multi-dataset external evaluation**, not a claim of definitive replication in one atlas. The term triangulation is reserved for the eventual case in which multiple prespecified, non-equivalent analyses are actually executed and support the resource-specific directions reported.

The strongest immediate resource is the Siletti/Human Brain Cell Atlas because it contains the highest-specificity available three-compartment proxy—MoSR, a VPL-labelled/targeted dissection, and S1C—and its specimens and assay are independent of AHBA. The highest-specificity matched endpoint nevertheless contains only one Siletti donor. MoSR is not isolated Cu/Gr, and the VPL-labelled dissection has an ontology-label discrepancy and possible adjacent-tissue overlap. Moreover, Siletti was already used in v0.11 for broad regional/cellular attribution of this panel. It is therefore **not a fully held-out validation dataset**. A newly frozen dissection-level analysis can still provide valuable external-assay evidence, but it must be described accurately.

FANTOM5 and BrainSpan are independent of AHBA, and no prior inspection of their proposed endpoints was found in the recovered project provenance; this is an operational record rather than proof that nobody has inspected them. Each has materially incomplete anatomy. Their results must remain separate, non-equivalent tests. **No meta-p value may combine non-equivalent anatomical proxies or endpoints.**

GSE3526 is explicitly excluded from that external strategy. It is a public cohort in the documented historical source lineage of BIGE and a donor-level reconstruction target; formal sample- and processing-level reconciliation has not established numerical equivalence to the internally analyzed aggregate. Its broad-node endpoints, background sensitivities and donor heterogeneity were already inspected before v1.2. It can improve auditability and expose source-lineage heterogeneity, but it cannot supply independent, external or held-out validation.

## Independence categories used here

| Category | Definition | Interpretation |
|---|---|---|
| **U-I: no prior inspection documented, independent** | Different donors/cohort and assay from AHBA, with no inspection of the proposed 19-gene endpoint found in the recovered provenance | Eligible for a prospectively specified external test, subject to anatomical adequacy; this label is not a universal claim of virginity |
| **I-P: independent donors/platform, previously inspected** | Different specimens and assay from AHBA, but this panel or closely related regional contrasts were already examined | External evidence, but not a fully held-out validation dataset |
| **P-A: partial anatomy** | One or more target relay nodes are absent or represented only by a broad/non-equivalent proxy | Can test only a modular or descriptive secondary endpoint |
| **N-I: not independent** | Same source cohort as discovery or another direct dependency | Discovery reconstruction, audit, or internal sensitivity only |

These categories are orthogonal where needed. For example, BrainSpan is U-I but also P-A.

## Ranked feasibility table

| Rank | Resource | Anatomical coverage | Donors / samples and modality | Panel testability | Access and licensing | Independence classification | Chief caveat and appropriate role |
|---:|---|---|---|---|---|---|---|
| **1** | **Siletti et al. / Human Brain Cell Atlas v1.0** | S1C: postcentral gyrus/primary somatosensory cortex; MoSR: a mixed sensory-relay medullary proxy rather than isolated Cu/Gr; a VPL-labelled/targeted dissection with ontology and overlap caveats; also LP+VPL and ventral lateral nuclear proxies | Adult human snRNA-seq; 3 postmortem donors; 3,369,219 cells/nuclei from approximately 100 dissections. HCA reports S1C 38.2k cells, MoSR 13.6k, the VPL-labelled entry 6.9k, and LP+VPL 17.7k. Official sample metadata supports only one donor-matched highest-specificity MoSR–VPL-labelled–S1C triad | Yes. Whole-transcriptome counts permit the complete 19-gene panel, preferably as prespecified neuronal pseudobulk | HCA exports are CC BY 4.0. The Allen AWS mirror states CC BY-NC 4.0; the distribution route and its terms must be recorded. Academic publication is compatible, but any commercial reuse should be separately reviewed | **I-P**, not U-I. Donors and platform are independent of AHBA, but v0.11 already inspected this panel in Siletti at broad regional/cellular resolution | Best immediate external-assay analysis, but not a held-out replication. Both the medullary and thalamic nodes are dissection-level proxies; only one donor has the highest-specificity matched triad; cortical and cell-composition effects remain possible |
| **2** | **FANTOM5** | Two matched broad triads: medulla oblongata, whole thalamus, and parietal lobe. An exact postcentral-gyrus sample exists separately as an adult pool, not matched to either triad | CAGE on total RNA. Two adult female donors have complete broad triads: donor10196, age 60; donor10252, age 76 | Probably yes after a frozen promoter-to-gene aggregation and symbol/alias audit | Raw reads, alignments, CTSS and portal data are public; FANTOM5 data are released under CC BY 4.0 | **U-I + P-A** | Strongest three-compartment resource with no prior endpoint inspection found in the recovered provenance, but anatomy is broad, n=2, and CAGE measures promoter activity rather than conventional gene-level abundance |
| **3** | **BrainSpan / Human Brain Transcriptome** | S1C is present; thalamus is mediodorsal nucleus (MD), not VPL; medulla is absent | Developmental transcriptome across 16 regions. The primary exon-array study reports 1,340 tissue samples from 57 brains. In the current RNA-seq metadata, the adult subset contains 8 donors; 7 have paired S1C–MD samples | Yes at gene level for an adult-only modular endpoint | Normalized expression and metadata are publicly downloadable. Some sequence/genotype material requires dbGaP authorized access. The download portal displays Allen terms/citation policy rather than a dataset-specific CC license | **U-I + P-A** | Eligible only for a prespecified, non-equivalent thalamocortical modular sensitivity. MD is not a somatosensory thalamic relay and the medullary node is missing; it cannot corroborate a DCML edge without additional anatomical comparators and specificity controls |
| **4** | **Bakshi et al. / Stanley thalamic laser-capture dataset** | Exact VPL and anterior thalamic nucleus from the same subjects; no medulla and no S1 | Laser-capture microdissection of approximately 3,000 large thalamic neurons per region, followed by Affymetrix U133 Plus 2.0 microarray; small normal-control and schizophrenia subsets after QC | Technically yes if the complete sample-level matrices remain obtainable | Article and supplements are public. The original Stanley Genomics data endpoint and reusable-data license require confirmation before relying on this resource | **U-I + P-A** | Potential exact-VPL specificity check only. It cannot test the three-node topology, has small samples and disease-related sampling, and current raw-data availability is uncertain |
| **5** | **Braineac / UK Brain Expression Consortium, GSE60862** | Inferior olivary nucleus, thalamus sampled at lateral geniculate level, and frontal/temporal/occipital cortices; no Cu/Gr, VPL, parietal cortex, or S1 | Affymetrix Human Exon 1.0 ST; 1,231 samples from 134 neurologically and neuropathologically normal adults across 10 regions | Yes at gene/exon level | Public GEO deposition and Braineac resource; the GEO accession does not present a CC license, so repository and publication terms must be followed | **U-I + P-A** | Large independent cohort useful as an anatomical negative control or detectability audit. Its three sampled regions are not equivalent to the DCML relays, so it cannot be called replication |
| **6** | **Human Protein Atlas Brain v25.1** | Exact nucleus cuneatus, nucleus gracilis, ventral posterolateral thalamic nucleus, and postcentral gyrus subdivisions | MGI DNBSEQ RNA-seq; 966 samples across 193 subregions. Inspection of the official sample header gives cuneatus n=9, gracilis n=3, VPL n=6, and postcentral general/dorsal/middle/ventral n=16 in total | Yes, complete panel at transcript/gene level | Public downloads; HPA copyrightable database content is CC BY 4.0, subject to third-party constraints | **I-P**, not U-I | Anatomically the best public bulk resource, but exact relay residuals for this panel were already inspected in v0.4/v0.6. It is internal/post hoc corroboration only, not a held-out v1.2 confirmation |
| **7** | **GSE3526 whole-body expression cohort** | Broad medulla, thalamus, and parietal lobe; no nucleus-isolated relays | Affymetrix U133 Plus 2.0; 353 samples across 65 tissues and 10 donors. Eight donors have complete broad three-/four-node data; 24/32 arrays enter the endpoints. The series has 193 neural arrays in all 10 donors, of which 180 belong to the 8 complete donors and can influence their frozen z-score background. The 19 genes map to 39 historical GPL570 probes | Yes, subject to multi-target probe annotation and background-universe sensitivities | Public GEO deposition; source file must pass the recorded byte/hash gate and repository citation terms apply | **N-I; known before v1.2** | Public cohort in the documented historical BIGE source lineage; numerical equivalence to the internally analyzed aggregate is not established. Donor reconstruction and post-selection source-lineage sensitivity only; not independent validation. Positive T is a relative serial-order contrast, not positive coexpression or continuity |
| **8** | **GTEx v8** | Brain cortex and other broad brain tissues are present, but no medulla oblongata, thalamus, postcentral gyrus, or S1 sample is available | Adult bulk RNA-seq; 17,382 RNA-seq samples across 54 tissue sites. NIH pages describe the final cohort as 838 donors with genomic data, while release notices also report 948 postmortem donors represented in RNA-seq | Yes for general expression/detectability, not for the route endpoint | Open-access aggregate expression through GTEx Portal; individual-level protected data through dbGaP/AnVIL under controlled terms | **U-I + P-A**, anatomically ineligible | Cannot test the route, any route edge, or an anatomically meaningful three-node proxy. Use only as a general technical or detectability reference |

## Resource-specific evidence and source links

### 1. Siletti / Human Brain Cell Atlas

- The official HCA overview states that high-throughput snRNA-seq surveyed more than three million nuclei from approximately 100 dissections across the adult brain in three postmortem donors: [Human Brain Cell Atlas v1.0](https://data.humancellatlas.org/hca-bio-networks/nervous-system/atlases/brain-v1-0).
- The same page lists:
  - postcentral gyrus/primary somatosensory cortex, S1C, 38.2k cells;
  - sensory relay nuclei in medulla oblongata, MoSR, 13.6k cells;
  - LP+VPL, 17.7k cells;
  - the VPL-labelled entry, 6.9k cells.
- The entry is named VPL but carries an incongruent ontology/tissue label of “lateral posterior nucleus of thalamus.” The dissection record indicates VPL targeting but does not eliminate possible adjacent-tissue overlap. Both library identifiers and the dissection mapping must be verified from metadata or authoritative clarification before the endpoint is eligible; the label must never be silently upgraded to an anatomically pure VPL sample.
- The authors’ official repository provides expression matrices, cell/dissection metadata, raw-read links, and the tables used for this feasibility audit: [adult-human-brain repository](https://github.com/linnarsson-lab/adult-human-brain), [samples.xlsx](https://github.com/linnarsson-lab/adult-human-brain/blob/main/tables/samples.xlsx), [dissections.xlsx](https://github.com/linnarsson-lab/adult-human-brain/blob/main/tables/dissections.xlsx), and [donors.xlsx](https://github.com/linnarsson-lab/adult-human-brain/blob/main/tables/donors.xlsx).
- Allen’s public mirror documents the matrices and metadata and states a CC BY-NC 4.0 license: [Allen Brain Cell Atlas data access](https://alleninstitute.github.io/abc_atlas_access/descriptions/WHB-10Xv3.html).
- HCA states that downloaded/exported atlas data are licensed CC BY 4.0. This distribution-level difference should be preserved in the provenance record.
- Primary publication: [Siletti et al., Science 2023](https://doi.org/10.1126/science.add7046).

The official sample metadata supports the following prospective anatomical definitions without opening expression values:

| Analysis tier | Donor | Medullary node | Thalamic node | Cortical node | Status |
|---|---|---|---|---|---|
| Highest-specificity available | H19.30.001 | MoSR proxy, 10X379_1/2 | VPL-labelled/targeted, 10X392_5/6 | S1C, 10X277_7/8 | Only donor-matched triad at this specificity; neither subcortical node is anatomically pure |
| Proxy sensitivity | H18.30.002 | MoSR | LP+VPL | S1C | Thalamic proxy |
| Proxy sensitivity | H19.30.001 | MoSR proxy | VPL-labelled/targeted | S1C | Highest-specificity available proxy |
| Proxy sensitivity | H19.30.002 | MoSR | VLN annotated as VPM and VPL, 10X378_1/2 | S1C | Mixed thalamic proxy |

Technical libraries must be combined before inference. Library wells must never be treated as independent biological replicates.

### 2. FANTOM5

Official sample pages establish two donor-matched broad adult-human triads:

| Donor | Medulla | Thalamus | Parietal |
|---|---|---|---|
| donor10196, female, 60 | [FF:10174-103C3](https://fantom.gsc.riken.jp/5/sstar/FF%3A10174-103C3) | [FF:10168-103B6](https://fantom.gsc.riken.jp/5/sstar/FF%3A10168-103B6) | [FF:10171-103B9](https://fantom.gsc.riken.jp/5/sstar/FF%3A10171-103B9) |
| donor10252, female, 76 | [FF:10155-103A2](https://fantom.gsc.riken.jp/5/sstar/FF%3A10155-103A2) | [FF:10154-103A1](https://fantom.gsc.riken.jp/5/sstar/FF%3A10154-103A1) | [FF:10157-103A4](https://fantom.gsc.riken.jp/5/sstar/FF%3A10157-103A4) |

An exact postcentral sample exists as a separate mixed adult pool: [FF:10032-101E5](https://fantom.gsc.riken.jp/5/sstar/FF%3A10032-101E5). It must not be substituted into either donor-matched triad.

Portal and data access: [FANTOM5](https://fantom.gsc.riken.jp/5/).

### 3. BrainSpan / Human Brain Transcriptome

- Official overview and downloads: [BrainSpan](https://www.brainspan.org/) and [Developmental Transcriptome downloads](https://www.brainspan.org/static/download.html).
- The Human Brain Transcriptome portal lists the 16 sampled regions, including S1C and mediodorsal thalamic nucleus but not medulla: [HBT atlas](https://hbatlas.org/).
- Primary publication describing 1,340 samples from 57 brains: [Kang et al., Nature 2011](https://pubmed.ncbi.nlm.nih.gov/22031440/).
- Controlled-access molecular data description: [dbGaP phs000731.v2.p1](https://www.ncbi.nlm.nih.gov/projects/gap/cgi-bin/study.cgi?study_id=phs000731.v2.p1).

Any v1.2 use must filter to the adult subset before endpoint computation and must label MD as a non-equivalent thalamic proxy.

### 4. Exact-VPL laser-capture study

- Primary article: [Bakshi et al., 2020](https://onlinelibrary.wiley.com/doi/10.1155/2020/5176834).
- PubMed record: [PMID 32566292](https://pubmed.ncbi.nlm.nih.gov/32566292/).

No claim should depend on this resource until sample-level matrix availability, provenance, and reuse terms are verified.

### 5. Braineac / GSE60862

- Official accession: [GSE60862](https://www.ncbi.nlm.nih.gov/geo/query/acc.cgi?acc=GSE60862).
- Primary publication: [Ramasamy et al., Nature Neuroscience 2014](https://www.nature.com/articles/nn.3801) and [PubMed](https://pubmed.ncbi.nlm.nih.gov/25174004/).

Its regions are suitable as deliberate non-equivalent anatomical controls, not route relays.

### 6. Human Protein Atlas Brain

- Official data page, including 966 samples, 193 subregions, and exact regional names: [HPA Brain data](https://www.proteinatlas.org/humanproteome/brain/data).
- Transcriptomic methods: [HPA Brain transcriptomics](https://www.proteinatlas.org/humanproteome/brain/method/transcriptomics).
- License: [HPA license and citation](https://www.proteinatlas.org/about/licence).

Because the exact panel/region analysis was already examined in earlier versions, HPA must be fenced off from the v1.2 confirmatory gate.

### 7. GSE3526

- Official accession: [GSE3526](https://www.ncbi.nlm.nih.gov/geo/query/acc.cgi?acc=GSE3526).
- Primary CNS source-lineage publication: [Roth et al., Neurogenetics 2006](https://doi.org/10.1007/s10048-006-0032-6).
- Related BIGE provenance: [Lee et al., FASEB J 2005](https://doi.org/10.1096/fj.04-3552fje) and [Ushach et al., Clin Immunol 2015](https://doi.org/10.1016/j.clim.2014.11.006). Ushach contains the published description of BIGE as providing expression information across more than 105 human cells and tissues and cites Lee and Roth. This source chain does not establish that all of BIGE is identical to GSE3526.

The public reconstruction contains 353 GPL570 arrays across 65 tissues and 10 donors. There are no donor-by-tissue duplicates. Eight donors have complete broad-node data; 24 arrays form the three-node endpoint and 32 form the four-node endpoint. The series contains 193 arrays from the frozen neural universe across all 10 donors; 180 belong to the 8 complete donors and can influence their within-donor standardization. Deposited values are linearized RMA (range 4.538–33730.54). The historical `log2(x+1)` transform is appropriate; `log2(x)` gives an almost identical sensitivity.

Known pre-v1.2 results are:

| Node order | Probe collapse | Mean T | Positive donors | Descriptive label tail | Post-selection sign-flip tail |
|---|---|---:|---:|---:|---:|
| medulla–thalamus–parietal | median | 0.213706 | 5/8 | 0.099680 | 0.097656 |
| medulla–thalamus–parietal | mean | 0.221053 | 6/8 | 0.110654 | 0.109375 |
| medulla–thalamus–parietal | delegate | 0.191338 | 5/8 | 0.144033 | 0.113281 |
| DRG–medulla–thalamus–parietal | median | 0.152193 | 6/8 | 0.057699 Monte Carlo; 0.0578575 exact | 0.097656 |
| DRG–medulla–thalamus–parietal | mean | 0.165205 | 6/8 | 0.050609 Monte Carlo; 0.0487755 exact | 0.085938 |
| DRG–medulla–thalamus–parietal | delegate | 0.109722 | 6/8 | 0.132369 Monte Carlo; 0.1311227 exact | 0.140625 |

The exact four-node finite-space convolution shows that the mean branch crosses 0.05 solely because of Monte Carlo error. This cannot support a threshold narrative: tissue labels are not anatomically exchangeable, the panel/source were selected previously, and all label/sign fractions are descriptive rather than confirmatory p values. Future reruns must compute and report the exact deterministic label-space fraction alongside the archived Monte Carlo result.

Positive T is also not positive coexpression. With median probe collapse, donor-mean correlations for medulla–thalamus, thalamus–parietal and medulla–parietal were −0.022149, −0.028509 and −0.239035. T was positive because the serially adjacent pairs were less negative than the selected non-adjacent pair. In the four-node result, only DRG–medulla was positive (~0.203); the remaining serial edges were small or negative. Terms such as shared-expression continuity, molecular flow or tract support are not permitted.

The endpoint is strongly conditioned on its background universe:

| Background | Collapse | Three-node T | Four-node T |
|---|---|---:|---:|
| 20 neural tissues common to all eight donors | median | 0.243092 | 0.150073 |
| 20 neural tissues common to all eight donors | mean | 0.241776 | 0.172588 |
| 20 neural tissues common to all eight donors | delegate | 0.199890 | 0.102924 |
| All 65 tissues available within donor | median | 0.099123 | 0.045249 |
| All 65 tissues available within donor | mean | 0.101316 | 0.054678 |
| All 65 tissues available within donor | delegate | 0.055702 | −0.005994 |
| No within-donor z standardization | median | 0.003399 | 0.010526 |
| No within-donor z standardization | mean | −0.001645 | 0.008114 |
| No within-donor z standardization | delegate | 0.003509 | 0.000512 |

The common-20 result addresses donor-to-donor differences in neural-tissue availability and remains directionally similar. The all-65 and no-z alternatives change the estimand and demonstrate that the historical result is specifically a contrast relative to a neural background, not general or absolute regional similarity. Every report must state this conditioning and show the full sensitivity table; the background cannot be selected retrospectively.

There is substantial post hoc donor heterogeneity. For the three-node endpoint, all four recorded females exceeded all four recorded males: female-versus-male mean T was 0.563 versus −0.136 (median), 0.569 versus −0.127 (mean), and 0.529 versus −0.146 (delegate); the descriptive exact two-sided partition fraction is 2/70=0.0286. The four-node endpoint did not show the split (~0.14 versus ~0.16). RAI2 is the only X-linked panel gene, but omitting it left female-minus-male differences of 0.598/0.590/0.560 and complete separation for median/mean; all leave-one-gene differences remained positive (0.549–0.768), and jointly excluding RAI2 and C7orf25 still left 0.422–0.490. These are post hoc robustness diagnostics, not evidence of sex biology. Sex is perfectly confounded with these eight donor identities, n=4 per stratum, and only age/sex are available; cause of death, PMI, RIN and batch are missing. Any further subgroup analysis requires a frozen metadata-aware rule and must preserve the full limitation.

The GPL570 annotation snapshot inspected on 9 September 2026 creates gene-attribution ambiguity: all three historical probes assigned to RAB3GAP2 also map to AURKAPS1, while the only historical C7orf25 probe also maps to PSMA2. Corrected package v2_A02 archived the inspected mappings and ran a labelled 17-gene post hoc sensitivity excluding both historical assignments; mean contrasts remained positive across the three collapse methods. This directional persistence does not validate the remaining probes, make either ambiguous signal gene-specific, or create confirmatory evidence. A future rerun must freeze the exact annotation release and reproduce or supersede this handling; CEL-level processing with a contemporary custom definition remains prospective.

For a citable reproducible rerun, acquire `GSE3526_series_matrix.txt.gz` from the frozen official GEO route, require exactly 85,860,948 bytes and SHA-256 `77cc18cb606adb8028e0ef8e84db288f26b160308a364089c3876c0335c4b7d0` before parsing, use relative-path manifests, pin Python/dependencies in a machine-readable lock, record BLAS/LAPACK/thread variables, and require two byte-identical scientific runs. Corrected package `gse3526_donor_reconstruction_v2_A02.zip` implements these gates, omits the large public matrix from the archive, and reproduced 22/22 outputs byte-for-byte in two clean executions. Its SHA-256 is `4edd3d126023e730ae8c93bcf6f74bb193237524429c20d5108e63487ffadfae`; it remains a source-lineage audit, not validation.

This cohort is a reconstruction target, provenance audit and source-lineage post-selection sensitivity only. It is excluded from every independent-validation gate and from any meta-analysis of external resources.

### 8. GTEx

- NIH program and final-release summary: [NIH Common Fund GTEx](https://commonfund.nih.gov/GTEx).
- Official open-access downloads: [GTEx Portal datasets](https://gtexportal.org/home/datasets).
- The HPA official data page provides a compact listing of the available GTEx brain-region groupings and confirms the absence of medulla and thalamus: [HPA Brain data, GTEx section](https://www.proteinatlas.org/humanproteome/brain/data).

## Spatial transcriptomics assessment

No independent public adult-human spatial transcriptomic atlas was identified that spans isolated Cu/Gr, VPL-specific thalamus, and S1/postcentral cortex.

The multiplexed RNA-FISH/EEL files linked from the Siletti repository belong to the same study/atlas; some material may involve donors outside the three whole-brain snRNA-seq donors, so donor independence must not be denied categorically. They remain ineligible as an independent route-level validation because they are study-linked, use a targeted panel, and do not provide the required three-node coverage or assured coverage of all 19 genes.

Whole-brain, high-resolution transcriptomic/spatial atlases are available in mouse. The principal example is [Yao et al., Nature 2023](https://www.nature.com/articles/s41586-023-06812-z). Cross-species evidence may be used only as secondary mechanistic or orthology support; it cannot rescue failure of the human validation gate.

## Recommended frozen v1.2 validation architecture

### A. Pre-analysis seal

Before accessing highest-specificity dissection-level proxy expression for the 19-gene panel in Siletti, freeze and hash:

1. source version and distribution route;
2. expression and metadata file hashes;
3. the 19 HGNC symbols and any accepted aliases;
4. exact donor, dissection, and technical-library identifiers;
5. neuronal/non-neuronal class inclusion rules;
6. pseudobulk aggregation and normalization;
7. detection thresholds and missing-gene rules;
8. control-gene eligibility and matching rules;
9. random seed, number of null draws, and empirical-p convention;
10. the single primary statistic and pass/fail gate;
11. all secondary and sensitivity analyses;
12. exact permitted claims for each outcome.

No choice may be revised on the basis of observed expression of the 19 target genes.

Before the highest-specificity Siletti endpoint is eligible to run, both technical-library identifiers for the VPL-labelled/targeted dissection must pass a metadata-only gate. The entry-name versus ontology/tissue-label discrepancy and the stated possibility of adjacent-tissue overlap must be resolved or bounded from the authors' dissection records, NeMO metadata, or authoritative clarification. If the identity cannot be supported without examining target-panel expression, the VPL-labelled endpoint is `ANATOMICALLY_INELIGIBLE`; LP+VPL or VLN may remain declared proxy sensitivities but may not replace the failed primary gate.

### B. Primary Siletti endpoint: highest-specificity available proxy

Use only the H19.30.001 matched triad:

- MoSR proxy: 10X379_1 and 10X379_2;
- VPL-labelled/targeted dissection: 10X392_5 and 10X392_6;
- S1C: 10X277_7 and 10X277_8.

Combine technical libraries within each dissection. The biological unit is the donor/dissection, not the library.

The recommended primary expression unit is a neuronal pseudobulk generated by summing raw UMI counts under a frozen neuronal definition, followed by one prespecified normalization. All-nuclei or composition-standardized results should be sensitivities, not interchangeable alternatives.

Use the already motivated topology statistic:

$$
T =
\frac{\rho(\mathrm{MoSR},\mathrm{VPL})+
\rho(\mathrm{VPL},\mathrm{S1C})}{2}
- \rho(\mathrm{MoSR},\mathrm{S1C}),
$$

where each correlation is computed over the fixed 19 genes after the exact frozen transformation.

Calibrate T against a frozen collection of eligible control-gene panels. Use the plus-one empirical convention:

$$
p = \frac{b+1}{B+1}.
$$

The recommended single gate is:

$$
T_{\mathrm{observed}} > q_{0.95,\mathrm{null}}
\quad\text{and}\quad
p \leq 0.05.
$$

Failure of either condition is failure of the primary Siletti gate. Alternative cell strata, normalizations, proxies, or null definitions cannot replace it.

Because the highest-specificity endpoint has one donor, even a positive result does not establish population-level replication. Its inferential unit is the prespecified gene panel relative to matched gene panels in one independently assayed brain, using declared anatomical proxies.

### C. Siletti generalizability sensitivity

Run the same frozen statistic on the three donor-specific proxy triads listed above, allowing only the prespecified thalamic definition for each donor. Report:

1. T for each donor;
2. sign concordance;
3. the donor-average T;
4. leave-one-donor-out values;
5. the corresponding frozen-panel null calibration.

This analysis is secondary because its thalamic node changes across donors. It cannot supersede a failed or ineligible highest-specificity VPL-labelled gate.

### D. Cell-composition sensitivities

Recommended order:

1. neuronal pseudobulk as primary;
2. prespecified broad neuronal-class pseudobulks as secondary;
3. all-nuclei pseudobulk with common frozen class weights as sensitivity;
4. unweighted all-nuclei pseudobulk as descriptive only.

The Siletti sampling/enrichment design makes naive all-cell comparisons vulnerable to composition differences.

### E. External resources with no prior endpoint inspection found

1. **FANTOM5 broad triads:** compute the exact frozen T separately in donor10196 and donor10252 after a prespecified promoter-to-gene aggregation. Require the same direction in both donors for descriptive concordance. Do not pool the exact postcentral sample into these donor-specific triads.
2. **BrainSpan adult S1C–MD module:** test only a prespecified, non-equivalent two-node profile/correlation sensitivity in the seven paired adult donors. Do not describe MD as VPL, as a DCML edge, or as corroboration without additional anatomical comparators and specificity controls.
3. **Optional exact-VPL laser-capture audit:** use only if the full sample-level data and terms are confirmed. Treat it as a one-node specificity analysis.
4. **Braineac:** use, if useful, as an anatomical non-equivalent control or panel detectability audit.

These resources do not test the same null hypothesis. **Do not combine their p values into a meta-p, Fisher statistic, Stouffer score, or a single confirmatory probability.** Report the results in parallel with explicit endpoint and anatomy labels.

## Permitted interpretation

Permitted wording must depend on the actual pattern of results:

- **If only the highest-specificity Siletti endpoint passes:**

  > The prespecified adjacency contrast was observed in one Siletti donor using a MoSR proxy, a VPL-labelled/targeted dissection, and S1C; this constitutes external-assay corroboration, not independent replication.

- **If Siletti passes and both prespecified FANTOM5 donor-specific broad triads are directionally concordant:** the preceding sentence may be followed by a separate statement that broad-anatomy CAGE results were directionally concordant in two donors. It must not be merged into a common probability or called exact anatomical replication.

- **If the BrainSpan modular sensitivity is favourable:** report it only as a result for the non-equivalent MD–S1C module. Do not use it to strengthen a three-node or VPL claim.

- **If any frozen external gate fails, is anatomically ineligible, or disagrees:** state that outcome beside favourable results. “Triangulation” may be used only when the named prespecified resources were actually executed and their separately defined results support the stated, resource-specific directions.

The manuscript must also state:

> The Siletti atlas was not fully held out, because earlier work had examined the same panel for broad regional and cellular attribution; the highest-specificity dissection-level proxy endpoint was specified prospectively.

Avoid:

- “independent atlas replication” without qualification;
- “validated in three independent atlases”;
- “confirmed across multiple donors” based on the one highest-specificity Siletti triad;
- claims that MoSR equals isolated cuneate/gracile nuclei;
- claims that BrainSpan MD or Braineac LGN equals VPL;
- any combined significance measure across non-equivalent proxies.

## Route to definitive confirmation

A truly independent, population-level human confirmation would require newly generated matched tissue from the same donors:

- isolated cuneate and gracile nuclei;
- exact VPL;
- postcentral/S1 cortex;
- a prespecified minimum number of neurologically normal adult donors;
- bulk RNA-seq or, preferably, snRNA-seq with a frozen analysis plan.

The NIH NeuroBioBank can support a prospective tissue request but is not a ready-made transcriptomic atlas. Requests require an institution with an active FWA, a PI CV/biosketch, a signed material transfer agreement, repository and NIH review, and specimen availability: [NIH NeuroBioBank researcher process](https://neurobiobank.nih.gov/researchers/).

## Audit statement

This document records dataset contents, anatomical eligibility, independence status, access/licensing constraints, known pre-v1.2 GSE3526 source-lineage dependence, and a proposed decision framework. It does not contain results from, or inspection of, the proposed Siletti dissection-level expression endpoint, FANTOM5 target-panel endpoint, BrainSpan target-panel endpoint, or any other new v1.2 validation endpoint. The GSE3526 results above were already inspected and are fenced as source-lineage post-selection sensitivity, not validation; numerical equivalence to the internally analyzed BIGE aggregate remains unestablished.
