## Supplementary Archive S1 for "An anatomy-constrained transcriptomic screen identifies a 19-gene panel with an ordered regional similarity pattern in the human somatosensory system": README_protocol_v1_2.md

# Protocolo v1.2 — borrador para auditoría

**Estado abreviado:** `DRAFT_NOT_FROZEN_NOT_EXECUTABLE`. Este documento no autoriza calcular endpoints. El estado completo y gobernante está en el JSON. La búsqueda externa está completa y no identificó un único conjunto mantenido fuera de todo análisis previo, exacto y multidonante. La auditoría raíz concluyó que el diseño es científicamente viable como borrador. La autoridad para analizar el agregado BIGE interno y publicar estadísticas, tablas y figuras agregadas no reconstructivas ya quedó registrada; todavía faltan la factibilidad sólo-covariables, la revisión estática/estadística del ejecutable, el allowlist/exclusion audit archivo por archivo para la liberación pública, la decisión y protocolo separado para recursos externos, y el sellado final por hashes.

La decisión detallada está en `root_audit_v1_2_draft.md`: `CONDITIONAL_GO_AS_DESIGN_DRAFT__NO_GO_TO_FREEZE_OR_EXECUTE`.

## Objetivo

v1.2 está diseñado para resolver las objeciones que todavía pueden modificar de manera material el manuscrito antes de enviarlo:

1. nombrar y estimar correctamente el resultado como **contraste de adyacencia entre tres regiones**;
2. separar si el contraste proviene de similitud elevada entre pares adyacentes, de baja similitud entre extremos o de ambas;
3. formalizar la influencia de genes individuales sin seleccionar o eliminar genes;
4. medir dependencia del colapso de sondas y del procesamiento;
5. evaluar una sensibilidad explícita a composición celular;
6. fortalecer el nulo de selección emparejando coexpresión basal, estabilidad entre donantes y autocorrelación espacial;
7. decidir con honestidad si un nulo combinado selección+espacio es ejecutable y qué puede —y qué no puede— demostrar.

El protocolo no intenta convertir evidencia posdescubrimiento en validación independiente. Los resultados y decisiones congelados de v1.0 y v1.1 permanecen intactos.

## Cambio obligatorio de lenguaje

El endpoint principal conserva exactamente la aritmética previa, pero su nombre defendible es **three-region adjacency contrast**:

\[
T = \operatorname{mean}_{d=1}^{6}\left[
\frac{\rho_{12,d}+\rho_{23,d}}{2}-\rho_{13,d}
\right].
\]

Aquí, \(\rho_{12}\) es Cu/Gr–DTLv, \(\rho_{23}\) es DTLv–postcentral y \(\rho_{13}\) es Cu/Gr–postcentral. Un \(T>0\) sólo demuestra una diferencia relativa. No demuestra conectividad, flujo molecular ni continuidad de un tracto.

La agregación siempre respeta al donante. En cada donante se calculan primero \(\rho_{12}\), \(\rho_{23}\), \(\rho_{13}\), A, B, E y T a través de los genes; después se obtiene por separado la media aritmética de cada estadístico entre los seis donantes. En particular, `B_group` es la media de los seis mínimos intradonante, no el mínimo de dos correlaciones ya promediadas. Nunca se agrupan muestras entre donantes ni se correlacionan perfiles regionales promediados entre donantes.

Para afirmar que existe **similitud adyacente elevada**, además deben sostenerse bajo los dos modelos espaciales primarios:

- \(A=(\rho_{12}+\rho_{23})/2\), la similitud adyacente media;
- \(B=\min(\rho_{12},\rho_{23})\), el cuello de botella que exige contribución de ambos pares.

La similitud entre extremos \(E=\rho_{13}\) se prueba y reporta por separado. Si sólo \(E\) es inusual, el lenguaje correcto es “separación de extremos”, no “similitud elevada entre relés adyacentes”.

## Qué permanece congelado

- Los 19 genes y su orden.
- Los seis donantes AHBA.
- Los tres nodos, sus reglas anatómicas y el uso de muestras izquierdas/no pareadas.
- El residual `mediana(target) - mediana(siblings)` dentro de macrocompartimento.
- El kernel de Spearman, las colas y las reglas empíricas `+1`.
- La decisión histórica de v1.0 y el doble gate espacial P1A/P1B de v1.1.

En particular, no se aumentará el número de sorteos del componente espacial fronterizo, no se cambiará su semilla y no se sustituirá por otro valor de `k`. La descomposición regenerará **las mismas corrientes selladas** de v1.1 y deberá reproducir el \(T\) archivado por sorteo con tolerancia `1e-12` antes de liberar A, B o E.

## Estado de contaminación conocido

El ledger completo no se limita a leave-one-gene. Antes de este borrador ya se habían inspeccionado:

- el endpoint P0 mediana y todos los resultados v0.3–v1.1;
- la rama de media multisonde (`T=0.3845029239766082`; calibración histórica `p=0.007079858402831944`);
- la sonda delegada (`T=0.19005847953216373`; `p=0.2910141797164057`);
- A, B y E observados y sus calibraciones condicionales v1.0;
- los valores T favorables de los cuatro modelos espaciales secundarios v1.1.

Por ello, la media multisonde y los componentes de los modelos secundarios no son endpoint-blind. Sus recalibraciones nuevas sólo pueden ser sensibilidades o descomposiciones posdescubrimiento. Ninguna rama AHBA v1.2 constituye confirmación independiente.

El revisor escéptico ya calculó y comunicó que los 19 contrastes observados al omitir un gen abarcan **0.28896–0.41091**. Por tanto, leave-one-gene ya no es endpoint-blind.

Su análisis se conserva como diagnóstico formal pos hoc:

- se calculará el mínimo de los 19 valores leave-one-gene;
- se calibrará descriptivamente contra el nulo de pseudorrutas y ambos nulos espaciales;
- se reportarán cambios, concentración de influencia y percentiles;
- no tendrá gate, etiqueta de éxito, papel decisorio ni capacidad de reforzar el titular;
- no podrá utilizarse para eliminar genes, elegir procesamiento o definir el conjunto externo.

También se conocía antes de v1.2 la reconstrucción pública por donante de **GSE3526**, una cohorte del linaje fuente histórico documentado de BIGE. No se ha establecido mediante reconciliación de muestras y procesamiento que su agregado sea numéricamente equivalente al extracto BIGE efectivamente analizado. Ocho donantes tenían datos completos. En la tríada amplia medulla–thalamus–parietal, el T medio fue 0.213706 (mediana; 5/8 donantes positivos), 0.221053 (media; 6/8) y 0.191338 (sonda delegada; 5/8). Las fracciones descriptivas de cola por etiquetas/signos fueron, respectivamente, 0.099680/0.097656, 0.110654/0.109375 y 0.144033/0.113281. En la cadena DRG–medulla–thalamus–parietal, T fue 0.152193, 0.165205 y 0.109722; 6/8 donantes fueron positivos en las tres ramas y las fracciones Monte Carlo de etiquetas/signos fueron 0.057699/0.097656, 0.050609/0.085938 y 0.132369/0.140625. Ninguna de esas fracciones Monte Carlo o sign-flip fue menor de 0.05.

La convolución exacta del espacio finito de etiquetas de cuatro nodos da fracciones descriptivas 0.0578575, 0.0487755 y 0.1311227 para mediana, media y delegada. La rama de media cruza 0.05 sólo por error Monte Carlo (0.0506095 frente a 0.0487755). Esto no crea un resultado “significativo”: la intercambiabilidad anatómica sigue siendo falsa, el análisis es posselección y el cruce mismo demuestra que no se admite ninguna narrativa basada en umbral. Un rerun debe calcular la fracción exacta determinista y mostrarla junto a la cifra Monte Carlo archivada.

El signo positivo de T **no significa coexpresión positiva ni continuidad**. Con colapso por mediana, las correlaciones medias medulla–thalamus, thalamus–parietal y medulla–parietal fueron −0.022149, −0.028509 y −0.239035: los pares serialmente adyacentes fueron menos negativos que el par de extremos seleccionado. En la cadena de cuatro nodos sólo DRG–medulla fue positivo (0.203289); las demás aristas seriales fueron pequeñas o negativas.

Se conoce además una heterogeneidad pos hoc perfectamente separada por sexo registrado en la tríada: las cuatro mujeres superaron a los cuatro hombres, con T medio mujer/hombre de 0.563/−0.136 (mediana), 0.569/−0.127 (media) y 0.529/−0.146 (delegada); la fracción exacta bilateral descriptiva es 2/70=0.0286. Con n=4+4 y sexo confundido con identidad de donante, esto no permite inferir un efecto sexual. No depende sólo de RAI2, el único gen del panel ligado al X: al omitirlo, la diferencia mujer−hombre fue 0.598/0.590/0.560; cada leave-one-gene conservó diferencia positiva (0.549–0.768), y excluir juntos RAI2 y C7orf25 aún dejó 0.422–0.490. La cadena de cuatro nodos no mostró el mismo patrón (aproximadamente 0.14 frente a 0.16). Sólo hay edad y sexo; faltan causa de muerte, PMI, RIN y lote. Cualquier análisis adicional debe congelar antes una sensibilidad consciente de metadatos y reportar esta heterogeneidad completa, sin convertir estos diagnósticos pos hoc en biología sexual.

GSE3526 queda así clasificado como **sensibilidad posselección del linaje fuente**, no como validación independiente. La equivalencia numérica con el agregado BIGE interno no está establecida. Las permutaciones de etiquetas carecen de intercambiabilidad anatómica y los sign-flips son posselección; sus fracciones son descriptivas, no valores p confirmatorios.

Antes de congelar se exigirá una declaración auditada que enumere este ledger y confirme que no se abrió ningún otro endpoint v1.2. Cualquier acceso adicional no registrado degrada todos los análisis v1.2 a exploratorios pos hoc.

## Factibilidad con los archivos existentes

Para AHBA v1.2 y los recursos externos propuestos, la inspección se limitó a identidades, esquemas, dimensiones, hashes y conectividad basada sólo en coordenadas; no se abrió ningún endpoint nuevo. La excepción explícita es GSE3526: su reconstrucción del linaje fuente y nuevos diagnósticos pos hoc fueron inspeccionados antes del sellado y quedaron completos en el ledger, sin papel de calificación ni validación.

| Recurso | Disponibilidad actual | Consecuencia |
|---|---:|---|
| Caché AHBA v0.10 | 324 lotes, 3,234 sondas, 3,702 muestras por sonda, 1,390 símbolos | El inventario v0.10 contiene 763 símbolos = 19 candidatos + 744 controles; el universo analítico común N1 contiene 761 = 19 + 742 tras excluir dos controles sin contrastes celulares completos |
| Metadatos AHBA | 3,702 muestras con donante, ontología y coordenadas MRI nativas | Permite excluir por completo los tres macrocompartimentos del endpoint al construir covariables |
| Agregado regional BIGE mantenido internamente | 10,262 genes colapsados × 23 regiones agregadas; plataforma Affymetrix Human Genome U133 Plus 2.0 | Permite coexpresión basal fuera de la pseudorruta, pero no inferencia por donante ni autocorrelación espacial. Su análisis y la publicación de resultados agregados no reconstructivos están autorizados; la matriz completa y exportaciones reconstructivas no se redistribuyen |
| Reconstrucción pública GSE3526 | 353 arrays GPL570, 65 tejidos, 10 donantes; 8 completos para las comparaciones amplias; 19 genes/39 sondas históricas | Sensibilidad del linaje fuente ya inspeccionada, no validación; equivalencia numérica con el agregado BIGE interno no establecida. Un rerun exige descarga oficial con 85,860,948 bytes y SHA-256 `77cc18cb606adb8028e0ef8e84db288f26b160308a364089c3876c0335c4b7d0`, entorno bloqueado y auditoría de anotación de sondas |
| HPA/Siletti | expresión por clúster y metadatos de seis clases celulares amplias | Permite construir scores celulares de referencia; no constituye deconvolución de fracciones |
| Insumos completos de abagen | No están presentes: faltan los archivos PACall/background por donante | No se puede llamar “abagen” al procesamiento disponible; cualquier rama canónica requiere adquisición y enmienda previa al endpoint |

El caché canónico debe verificarse contra **su propio manifiesto de rerun**, porque otra copia histórica fue recuperada con una codificación gzip distinta. Mezclar manifiestos sería un hard stop aunque el contenido descomprimido pareciera equivalente.

### Procedencia y gobernanza de BIGE/GSE3526

La descripción autorizada para el manuscrito es: **agregado regional derivado de Body Index of Gene Expression (BIGE), mantenido internamente y no redistribuible**, procedente de perfiles Affymetrix U133 Plus 2.0. Ushach et al. describen BIGE como una base amplia con información de más de 105 células y tejidos humanos y citan a Lee et al. (2005) y Roth et al. (2006). Roth documenta 20 regiones del SNC y la cohorte pública GSE3526. Esto no autoriza a afirmar que GSE3526 sea la totalidad de BIGE ni que toda la matriz interna sea idéntica a una única serie pública.

La autorización registrada permite análisis y publicación de estadísticas, tablas, figuras y resúmenes no reconstructivos. Prohíbe redistribuir la matriz interna completa o exportaciones completas por fila/columna o de otra forma reconstructivas. Antes de promover un paquete público, cada archivo derivado de BIGE debe entrar en un allowlist o exclusion list explícito.

El paquete GSE3526 original pasó su integridad y una reejecución limpia desde la matriz oficial reprodujo los ocho CSV científicos byte a byte, pero no era autónomo. El paquete corregido `gse3526_donor_reconstruction_v2_A02.zip` añade descarga oficial con fallo cerrado por tamaño/hash, manifiestos relativos, locks de entorno, convolución exacta y sensibilidades de fondo, anotación y metadatos. Dos ejecuciones limpias reprodujeron 22/22 salidas byte a byte. El ZIP contiene 46 archivos y excluye deliberadamente la matriz GEO de 85,860,948 bytes, que se descarga por separado. Los valores depositados son RMA linearizado (rango 4.538–33730.54); `log2(x+1)` es apropiado y `log2(x)` produjo una sensibilidad casi idéntica. No hay duplicados donante×tejido: 24/32 arrays entran en los endpoints de tres/cuatro nodos. La serie contiene 193 arrays neurales en los diez donantes; 180 pertenecen a los ocho donantes completos y pueden influir en sus z intradonante.

Ese fondo es una dependencia mayor. Para mediana/media/delegada, el T de tres/cuatro nodos fue 0.243092/0.150073, 0.241776/0.172588 y 0.199890/0.102924 con los 20 tejidos neurales comunes; 0.099123/0.045249, 0.101316/0.054678 y 0.055702/−0.005994 con los 65 tejidos; y 0.003399/0.010526, −0.001645/0.008114 y 0.003509/0.000512 sin z. El resultado debe llamarse contraste condicionado al universo neural congelado, no similitud regional general, y la sensibilidad del universo de fondo debe acompañar todo reporte.

Además, la anotación GPL570 vigente asigna las tres sondas históricas de RAB3GAP2 también a AURKAPS1 y la única sonda histórica de C7orf25 también a PSMA2. El paquete corregido archiva el mapa inspeccionado y ejecuta una sensibilidad pos hoc de 17 genes que excluye ambas asignaciones históricas. La dirección se conservó, pero esto no valida las sondas restantes ni permite atribución específica a esos dos genes. Un rerun futuro debe congelar la versión exacta de anotación y reproducir o superar este manejo; el reprocesamiento CEL con una definición contemporánea continúa pendiente.

## Calificadores posdescubrimiento propuestos

La especificación, semillas, multiplicidad y gates que siguen **se proponen para quedar fijados únicamente tras auditoría y sellado**. Cuando un resultado nuevo no haya sido inspeccionado antes del sellado, el término máximo será **prospectively specified post-discovery qualifier**. Todos reutilizan AHBA, varios componentes observados ya eran conocidos y ninguno constituye inferencia confirmatoria ni confirmación independiente.

### C1. Nulo de pseudorrutas con matching basal reforzado

Se mantiene la mecánica de pseudorrutas de v1.0: misma elegibilidad, exclusión de los 23 candidatos históricos, mismos seis estratos tier×número de sondas y muestreo uniforme.

La línea histórica queda identificada por los hashes del protocolo v1.0, sus cuatro enmiendas y el generador registrados en el JSON. Deben reproducirse cinco pseudorrutas con totales exactos A=0/B=3/C=16, cero que además reproduzcan toda la tabla tier×sonda y 134 rutas elegibles. El soporte cero del replay estricto implica que N1 sólo calibra dentro del universo reconstruido; no identifica una probabilidad incondicional de toda la historia adaptativa.

La diferencia es que cada panel deberá emparejar además:

- Moran I residual medio y su variabilidad entre donantes;
- estabilidad de expresión entre donantes;
- mediana de coexpresión y mediana de coexpresión absoluta en muestras AHBA fuera de los tres macrocompartimentos del endpoint;
- las dos métricas equivalentes en las 19 regiones privadas externas a la ruta de selección de cada panel.

El grafo basal será 6-NN simétrico con peso inverso a distancia MRI nativa. El `k=6` surgió de una prueba basada únicamente en coordenadas: fue el mínimo conectado común después de excluir los tres macrocompartimentos del endpoint.

Por rama se ejecutará un piloto de 100,000 propuestas sólo de covariables y después se exigirán 50,000 paneles aceptados dentro de un máximo de 50,000,000 propuestas.

| Rama | Semilla piloto | Semilla de paneles |
|---|---:|---:|
| Mediana congelada | 20260920 | 20260921 |
| Media de todas las sondas | 20260922 | 20260923 |
| Sonda por estabilidad diferencial sin endpoint | 20260924 | 20260925 |

El análisis principal C1 usa la rama de mediana. Las otras dos ramas pertenecen a sensibilidad de procesamiento.

Cada corriente usa PCG64. Se elige uniformemente una ruta elegible, se recorren los seis estratos en orden léxico y se muestrea sin reemplazo dentro de cada estrato. Se conservan las primeras 50,000 propuestas aceptadas en orden cronológico; repeticiones exactas entre paneles están permitidas y se registran, sin deduplicación ni reponderación.

Para métricas por gen, el denominador SMD queda fijado como

\[
s_{ref}=\sqrt{\frac{\operatorname{var}_{ddof=1}(candidatos)+\operatorname{var}_{ddof=1}(742\ controles)}{2}}.
\]

Las cuatro métricas a nivel de panel usan la desviación `ddof=1` de las 100,000 propuestas piloto como denominador. Moran I se calcula sobre residuales centrados con el grafo simétrico 6-NN sin normalizar, usando la fórmula y convenciones completas del JSON. Cada propuesta debe cumplir calipers estandarizados de magnitud `≤0.25`: SMD para métricas por gen y diferencia dividida por la SD piloto para métricas de panel. El ensemble final exige `|SMD|≤0.15`, razón de varianzas `[0.50,2.00]`, KS ponderado `≤0.20`, al menos 20 rutas, ninguna con más de 20% y número efectivo inverso de Simpson `≥15`. Los pesos de símbolo son sus apariciones divididas entre `50,000×19`; ningún umbral puede relajarse después del endpoint.

C1 empareja resúmenes seleccionados de Moran I, estabilidad y coexpresión fuera del endpoint; no preserva la covarianza espacial o variogramas completos ni reproduce toda la investigación adaptativa. La autoridad para analizar el agregado BIGE y publicar resultados agregados no reconstructivos ya está registrada, por lo que ese bloqueo analítico se considera resuelto. La matriz completa y los derivados reconstructivos siguen excluidos. Hasta completar el allowlist/exclusion audit archivo por archivo, la reproducibilidad pública se describirá como restringida y no se promoverá ningún paquete como `public-data-only`.

### C2. Descomposición del nulo espacial

Se reconstruirán A, B y E para:

- P1A blockwise, 50,000 sorteos, semilla raíz histórica `20260912`;
- P1B whole-donor, 50,000 sorteos, semilla raíz histórica `20260913`.

No habrá sorteos adicionales. Antes de inspeccionar los componentes, cada vector \(A-E\) deberá coincidir con el archivo v1.1 correspondiente a `1e-12`.

Las seis pruebas A/B/E se corregirán juntas con Holm. Para considerar sustentada la frase “similitud adyacente elevada”, A y B deben superar p ajustada, q95 y el gate de Wilson en **ambos** modelos. Para llamar a E condicionalmente bajo, sus dos pruebas inferiores deben superar Holm, quedar por debajo de q05 y tener límite superior Wilson `<0.05`; si la conjunción falla se reportan los modelos por separado, sin rótulo binario de separación. C2 nunca cambia la clasificación histórica de v1.1.

## Sensibilidades prespecificadas

### S1. Procesamiento y sondas

Se compararán dos alternativas ejecutables con los mapas existentes:

1. **Media de todas las sondas**, siguiendo la línea float32 ya implementada en v0.10.
2. **Sonda de máxima estabilidad diferencial sin endpoint**: se agregan sondas por `structure_id`, se correlacionan regiones compartidas entre los 15 pares de donantes y se selecciona la mayor estabilidad; los tres macrocompartimentos del endpoint quedan excluidos de esta selección.

Cada rama se evaluará contra su nulo C1 y contra P1A/P1B, con 50,000 valores por referencia. Las seis pruebas se corrigen con Holm. Es una sensibilidad; no una repetición independiente.

La media multisonde ya tenía un T y una calibración histórica inspeccionados en v0.10; su nueva calibración es, por tanto, una sensibilidad pos hoc recalibrada. La rama de estabilidad diferencial no se ha inspeccionado, pero sigue reutilizando AHBA posdescubrimiento y tampoco puede calificar el titular. La sonda delegada de Allen queda como diagnóstico porque esa rama ya se inspeccionó en v0.10.

### S2. Proxy de composición celular preservado

Desde HPA/Siletti se elegirán, sin usar AHBA target, 25 marcadores por cada una de seis clases amplias: neuronal, glial, endotelial/mural, ciliada, sangre/inmune y mesenquimal. Se incluyen sólo clústeres autorizados para agregación; se excluyen `Splatter`, `Miscellaneous` y los 23 candidatos históricos. El score de marcador es el contraste ponderado por número de células entre una clase y las demás, resumido por la mediana de al menos tres regiones; se eligen los 25 genes mejor puntuados dentro de los 1,390 símbolos AHBA, con empates léxicos y solapamiento entre clases permitido y registrado.

En AHBA se construirán seis scores por muestra, se estandarizarán dentro de donante y, **para cada modelo y cada bloque ajustado por separado**, se residualizarán contra su diseño espacial base. Una SVD determinista retendrá los tres primeros componentes con convención de signo fijada. Para que el subespacio tridimensional sea identificable, la brecha entre los valores singulares tercero y cuarto debe superar `max(1e-10, 1e-8×s1)`; si no, la rama queda inconclusa. Nunca se reutiliza la SVD de donante completo dentro de P1A ni se concatenan bloques. Los tres componentes se incorporan al nuisance de MSR, de modo que la parte de expresión asociada al proxy utilizado permanezca fija en cada sustituto.

| Modelo | Sorteos | Semilla raíz |
|---|---:|---:|
| P1A + tres componentes celulares | 50,000 | 20260926 |
| P1B + tres componentes celulares | 50,000 | 20260927 |

Las dos pruebas se corrigen con Holm. El resultado sólo permite decir “robusto al score celular de referencia utilizado”; nunca “libre de composición celular”.

### S3. Componentes bajo modelos espaciales secundarios

Los cuatro modelos de sensibilidad de v1.1 se regeneran con sus 10,000 sorteos históricos y se descomponen en A/B/E. Sus T y resultados nominales favorables ya habían sido inspeccionados; sólo los componentes son nuevos. Primero deben reproducir el \(T\) archivado. Los 12 valores se corrigen con Holm y permanecen como descomposición diagnóstica de modelos ya conocidos, sin papel decisorio.

### S4. Marginales, distancias y variogramas: sensibilidad de compromiso

El muestreo AHBA es irregular y el mayor borde retenido por los grafos P1A congelados mide **49.1325 mm**. Antes de cualquier endpoint v1.2 se exigirá reproducir ese máximo dentro de `0.0001 mm` y exportar, por bloque, el resumen prespecificado de longitudes: recuento, mínimo, mediana, q90, q95, q99 y máximo. Los bins exactos de distancia quedan enumerados en el JSON y usan únicamente coordenadas. En ellos se auditarán semivariogramas de los residuales originales y sustitutos; los bins con menos de 20 pares quedarán como `NA` y nunca se fusionarán después de abrir resultados.

Como diagnóstico, los primeros 10,000 sorteos ya sellados de P1A y P1B se remapearán por rangos a la distribución residual empírica de cada gen. Esto conserva exactamente el marginal univariado **del residual de cada gen dentro de cada bloque ajustado**; no implica conservar el marginal del mapa reconstruido completo y **no** conserva simultáneamente la ortogonalidad al nuisance, la covarianza entre genes, el espectro/Moran I ni el variograma. Se cuantificará la distorsión de cada propiedad, incluidos semivariogramas en los primeros 100 sorteos auditados, y se reportarán T/A/B/E. Los dos T recibirán Holm sólo por transparencia, sin gate, función decisoria ni capacidad de rescatar el doble gate v1.1.

### Diagnósticos D1–D3

- **D1:** el mínimo leave-one-gene y la concentración de influencia se calibrarán descriptivamente contra N1-P0, P1A y P1B con 50,000 sorteos cada uno y Holm entre las tres referencias; la contaminación ya registrada impide todo gate o selección génica.
- **D2:** se reutilizarán exactamente los 50,000 sorteos del ensemble de elección de sonda v1.0-S5, sin valor p ni papel decisorio.
- **D3:** se construirán por separado estimandos residualizados por score celular para los 18 bloques P1A y los seis bloques P1B. Cambian el estimando, se reportan lado a lado y nunca reemplazan el endpoint original.

No se conoce en este entorno una implementación multivariada validada que preserve a la vez nuisance, marginales, estructura espacial y covarianza entre genes. Un sustituto tipo BrainSMASH aplicado independientemente a cada gen tampoco es equivalente para este endpoint, porque destruye la covarianza transversal que participa en las correlaciones de perfiles.

## Multiplicidad y gates

- Todas las colas son unilaterales y se fijan antes del endpoint.
- Los valores empíricos usan `(1+b)/(N+1)` y comparación inclusiva; no se eliminan valores ni se reduce el denominador.
- Se reportan `b`, p, q05/q95 y el intervalo Wilson bilateral de 95% para `b/N`. Wilson es sólo un gate auxiliar conservador de precisión Monte Carlo: no es un intervalo del efecto biológico, no es simultáneo y no garantiza FWER; la multiplicidad se gobierna por Holm donde corresponda.
- Un gate de calificación prospectiva exige p/quantil, p de Holm cuando corresponda y límite superior de Wilson `<0.05`.
- C1 precede a C2 en una secuencia fija. Si C1 falla, C2 todavía se reporta, pero sólo como análisis exploratorio de componentes.
- Los leave-one-gene se corrigen descriptivamente con Holm, pero nunca entran a la secuencia por la contaminación ya registrada.
- Las familias de procesamiento y célula tienen corrección Holm propia y no se usan para fabricar una p “omnibus”.

El doble gate histórico de v1.1 **ya falló**. Por ello v1.2 nunca podrá asignar una nueva etiqueta de robustez espacial global, rescatar la clasificación histórica ni sostener robustez espacial general, sin importar cómo salgan C1, C2 o las sensibilidades. C1 sólo puede calificar el lenguaje sobre selección; C2 sólo puede distinguir similitud adyacente elevada de separación entre extremos.

## Nulo combinado selección + espacio

Es **condicionalmente factible**, no automáticamente factible. Los mapas completos parecen disponibles para el universo requerido, pero antes se exige:

- verificación de los 324 lotes y todas las identidades;
- nulo C1 técnicamente válido;
- precomputación y auditoría de proyecciones espectrales para todos los genes;
- benchmark de memoria/tiempo sin endpoints;
- revisión estática independiente del código y de la independencia de corrientes.

Si todo pasa, se proponen sólo como diagnóstico:

| Referencia | Sorteos | Semilla raíz |
|---|---:|---:|
| Panel C1 + espacial P1A | 10,000 | 20260928 |
| Panel C1 + espacial P1B | 10,000 | 20260929 |

Se usa únicamente P0. En cada sorteo se selecciona uniformemente con reemplazo un panel aceptado N1 y se genera una corriente espacial independiente por bloque; el mismo vector de signos se aplica conjuntamente a sus 19 genes. La proyección optimizada debe coincidir con reconstrucción directa dentro de `1e-9` en 100 sorteos de auditoría. El endpoint es T agregado por donante, cola superior y p `+1`; los dos valores se ajustan con Holm sólo descriptivamente, sin umbral ni etiqueta de aprobación.

Este nulo pregunta qué ocurre con un **panel elegible aleatorio colocado espacialmente al azar**. No prueba la conjunción de “selección irrelevante” y “espacio irrelevante”. Un resultado favorable puede coexistir con el fracaso de uno de los dos nulos separados. Por ello jamás reemplaza ni promedia v1.0/v1.1 y no puede rescatar un componente espacial.

## Lo que no puede rescatarse pos hoc

- Repetir P1A con más sorteos, nueva semilla, otro `k`, otra distancia o distinto nuisance.
- Promediar P1A y P1B o escoger el modelo espacial favorable.
- Tratar el nulo combinado como sustituto de los dos nulos separados.
- Llamar validación a cualquier reanálisis, split o leave-one-donor dentro de AHBA.
- Elegir sonda o pipeline después de inspeccionar sus valores y luego llamarlo confirmatorio.
- Llamar deconvolución al proxy HPA/Siletti.
- Inferir exclusividad de DCML, conectividad, dirección, causalidad, fibromialgia, dolor o un blanco de estimulación.

## Hard stops

El alcance del fallo se decide antes de ejecutar. Una falla de hashes/linaje compartido, acceso no autorizado a endpoints o uso del agregado BIGE fuera del alcance autorizado detiene todo v1.2. La autoridad analítica y para publicar resultados agregados no reconstructivos ya está registrada; la falta del allowlist público bloquea la liberación, no el análisis interno autorizado. Una falla de N1-P0 bloquea la secuencia C1→C2; una falla de replay P1A/P1B vuelve C2 inconcluso. Una falla confinada a P1/P2, célula, S3/S4, D1–D4, la reconstrucción GSE3526 o un recurso externo detiene sólo esa rama, que debe reportarse como `TECHNICALLY_INCONCLUSIVE` y no puede omitirse ni reemplazarse.

Los eventos de detención aplicables son:

- se accede a un endpoint nuevo antes de sellar protocolo y ejecutable;
- falla un hash, identidad de sonda/muestra, alineación, recuento anatómico o replay histórico;
- un nulo no alcanza el número exacto de sorteos/paneles o falla balance/diversidad;
- un grafo no es conectado, el diseño pierde rango o falla un invariante MSR;
- una rama de procesamiento carece de sondas completas —fallo local de esa sensibilidad—;
- no existen 25 marcadores por clase, algún score residualizado tiene rango menor de tres o quedan menos de diez modos residuales —fallo local de S2/D3—;
- se intenta abrir un conjunto externo antes de congelar su mapeo anatómico y análisis;
- un defecto produce outputs parciales: se preserva el incidente, no se interpretan los endpoints y se requiere una enmienda hash-linked antes de reintentar.
- para GSE3526, no coinciden tamaño/hash de la matriz oficial, falta el lock de entorno, se omite la convolución exacta del espacio de etiquetas, no se resuelve explícitamente la anotación multiblanco RAB3GAP2/AURKAPS1 y C7orf25/PSMA2, o falla el replay científico —fallo local de esa sensibilidad—.

## Antes de congelar

El estado sólo puede cambiar mediante un **nuevo** `protocol_v1_2.json`; este borrador se conserva intacto como procedencia. Antes se requieren:

1. ledger firmado de contaminación y auditoría raíz;
2. revisión estadística/metodológica independiente;
3. decisión documentada sobre el protocolo externo separado;
4. auditoría hash de los 324 lotes canónicos y de todos los insumos;
5. factibilidad exclusivamente de covariables para N1, marcadores y rangos por bloque;
6. revisión estática y pruebas sintéticas del código, sin endpoints;
7. ejecutable, entorno, ledger y semillas sellados por hash.

Antes de **liberar públicamente** —no antes del análisis interno autorizado— se requiere además el allowlist/exclusion audit archivo por archivo para todo derivado BIGE y la verificación explícita de que no se incluyó la matriz completa ni una exportación reconstructiva.

Si se ejecuta, serán obligatorios los balances N1, auditorías de sonda/célula/distancia/variograma, replay T→A−E, tablas crudas y Holm, manifiesto del entorno, doble ejecución byte-idéntica y una matriz que vincule cada patrón de gates con el lenguaje permitido.

El claim máximo dentro del mismo AHBA es: “En datos AHBA previamente inspeccionados, el panel congelado de 19 genes muestra un contraste relativo de adyacencia entre tres regiones, acompañado de sensibilidades explícitas a selección, modelo espacial, procesamiento, influencia génica y proxy celular”. Todo resultado fallido, limítrofe o inconcluso debe aparecer junto a los favorables.

## Validación externa

La búsqueda documentada **no identificó un único conjunto mantenido fuera de todo análisis previo, anatómicamente exacto y multidonante** que cubra Cu/Gr, VPL y S1/postcentral. La opción defendible es evaluar recursos externos con protocolos y límites separados; sólo podrá llamarse triangulación si los análisis prespecificados se ejecutan y sus resultados apoyan las direcciones específicas que se nombren:

| Recurso | Independencia | Cobertura | Papel máximo |
|---|---|---|---|
| Siletti/Human Brain Cell Atlas | Donantes/plataforma independientes, pero el panel ya fue inspeccionado ampliamente en v0.11 | MoSR proxy–VPL-labelled/targeted–S1C; sólo un donante forma la tríada de mayor especificidad disponible | Evidencia externa dissection-level, no réplica con datos mantenidos fuera de análisis previos ni poblacional |
| FANTOM5 | Independiente de AHBA; no se encontró inspección previa de este endpoint en la procedencia recuperada | Dos tríadas amplias medulla–thalamus–parietal, CAGE; la muestra postcentral exacta es un pool separado y no se mezclará con ellas | Corroboración direccional de anatomía amplia en dos donantes, si ambos resultados prespecificados concuerdan |
| BrainSpan | Independiente de AHBA; no se encontró inspección previa del módulo propuesto en la procedencia recuperada | S1C–tálamo mediodorsal; falta médula y MD no es VPL | Sensibilidad modular talamocortical no equivalente; no es una arista DCML sin comparadores y controles de especificidad |
| GSE3526 | **No independiente**: cohorte pública del linaje fuente histórico y resultados ya inspeccionados; equivalencia numérica con el agregado BIGE interno no establecida | Medulla–thalamus–parietal amplia; ocho donantes completos; sin núcleos aislados | Reconstrucción, auditoría y sensibilidad posselección del linaje fuente solamente; queda excluido de cualquier gate externo |

Siletti tiene además un gate anatómico previo: la etiqueta “VPL” y su ontology/tissue label son incongruentes, y la disección puede incluir tejido adyacente. Deben verificarse ambas bibliotecas VPL-labelled/targeted únicamente con metadatos, registros de disección o aclaración autoritativa antes de abrir expresión. MoSR seguirá declarado como proxy medular, no Cu/Gr aislado. Si la identidad talámica no puede sostenerse, ese endpoint queda `ANATOMICALLY_INELIGIBLE`; LP+VPL o VLN sólo pueden permanecer como sensibilidades proxy.

Antes de cualquiera de estos endpoints se requiere un **protocolo de evaluación externa separado, hash-sealed**, que fije versión, donantes/dissecciones, bibliotecas técnicas, ontología, cobertura génica, pseudobulk o agregación promotor-gen, normalización, paneles control, endpoint, semillas, sorteos, multiplicidad y lenguaje permitido. El término triangulación queda reservado para análisis efectivamente ejecutados y concordantes en los sentidos específicos que se reporten.

Los tres recursos externos propuestos tienen anatomía, unidades y endpoints no equivalentes. Se reportarán en paralelo: queda prohibido combinarlos mediante meta-p, Fisher, Stouffer o una probabilidad omnibus, y un proxy favorable no puede sustituir un endpoint primario inelegible. GSE3526 no entra en ese conjunto externo ni en ninguna combinación. Si sólo el análisis Siletti de mayor especificidad pasa, el lenguaje máximo será corroboración en un único espécimen ensayado independientemente, no replicación independiente; cualquier mención de corroboración FANTOM5 o sensibilidad BrainSpan quedará condicionada a sus resultados reales y se formulará por separado.

Si no se ejecuta esa triangulación, el paper puede avanzar sólo como estudio AHBA posdescubrimiento con soporte espacial convergente pero no uniforme; v1.2 no cambia ese estatus.

## Referencias metodológicas

- [Lee et al., 2005 — expresión en tejidos humanos post mortem](https://doi.org/10.1096/fj.04-3552fje)
- [Roth et al., 2006 — relaciones moleculares entre 20 regiones del SNC](https://doi.org/10.1007/s10048-006-0032-6)
- [GEO GSE3526 — cohorte pública fuente](https://www.ncbi.nlm.nih.gov/geo/query/acc.cgi?acc=GSE3526)
- [Ushach et al., 2015 — descripción publicada de BIGE](https://doi.org/10.1016/j.clim.2014.11.006)
- [Markello et al., 2021 — abagen y variabilidad de pipelines](https://elifesciences.org/articles/72129)
- [Documentación estable de `abagen.get_expression_data`](https://abagen.readthedocs.io/en/stable/generated/abagen.get_expression_data.html)
- [Wagner y Dray, 2015 — Moran spectral randomization](https://doi.org/10.1111/2041-210X.12407)

Este README es un resumen humano no ejecutable. El detalle propuesto de calipers, QC, multiplicidad, semillas y límites está en `protocol_v1_2_draft.json`; ante una diferencia gobierna únicamente el **futuro JSON congelado**, que todavía no existe.
