## Supplementary Archive S1 for "An anatomy-constrained transcriptomic screen identifies a 19-gene panel with an ordered regional similarity pattern in the human somatosensory system": root_audit_v1_2_draft.md

# Auditoría raíz del borrador v1.2

**Fecha:** 2026-09-09

**Veredicto:** `CONDITIONAL_GO_AS_DESIGN_DRAFT__NO_GO_TO_FREEZE_OR_EXECUTE`

**Acceso a endpoints:** no se generó ni inspeccionó ningún endpoint AHBA v1.2 ni de los recursos externos propuestos. La reconstrucción GSE3526 del linaje fuente y sus diagnósticos pos hoc sí se inspeccionaron antes del sellado y están registrados como contaminación sin papel de calificación o validación; la equivalencia numérica con el agregado BIGE interno no está establecida.

## Decisión

El diseño v1.2 puede continuar como protocolo en construcción. No contiene un error estadístico fatal y aborda amenazas materiales del manuscrito: coexpresión basal, autocorrelación espacial, dependencia de sonda/procesamiento, composición celular, descomposición A/B/E y ausencia de confirmación externa exacta.

No está listo para congelarse ni ejecutarse. El JSON sigue marcado `DRAFT_NOT_FROZEN_NOT_EXECUTABLE`. Cualquier cálculo inferencial antes de completar los bloqueantes siguientes invalidaría el rótulo de calificación prospectiva posdescubrimiento.

La autoridad para analizar el agregado regional BIGE mantenido internamente y publicar estadísticas, tablas y figuras agregadas no reconstructivas quedó registrada por atestación del investigador. No se autoriza redistribuir la matriz completa ni exportaciones reconstructivas. Por tanto, se resolvió el bloqueo de autoridad analítica/publicación, pero permanece pendiente un allowlist/exclusion audit archivo por archivo antes de cualquier liberación pública.

## Base factual que no se reabre

- El endpoint AHBA conocido es `T=0.3451754385964913`.
- P1A blockwise falló: 2,558/50,000, `p=0.05117897642047159`, observado por debajo de `q95=0.345906432748538`.
- P1B whole-donor pasó: 192/50,000, `p=0.0038599228015439692`, `q95=0.2214985380116953`.
- La conjunción espacial primaria falló; el soporte es convergente pero no uniforme y dependiente del modelo.
- Los análisis de selección v1.0 y espacio v1.1 prueban nulos distintos y sus probabilidades no pueden combinarse.
- Ningún análisis dentro de AHBA constituye confirmación independiente.
- GSE3526 es una cohorte pública del linaje fuente histórico documentado de BIGE y permite una reconstrucción por donante; no se ha establecido equivalencia numérica con el agregado interno. Sus endpoints amplios ya fueron inspeccionados, por lo que sólo constituye una sensibilidad posselección del mismo linaje; no validación independiente.

## Correcciones incorporadas durante la auditoría

1. Se amplió el ledger de contaminación para incluir P0, media multisonde, sonda delegada, A/B/E, sensibilidades espaciales conocidas y leave-one-gene.
2. Se corrigió el universo: 763 símbolos almacenados corresponden a 19 candidatos + 744 controles v0.10; N1 usa 19 + 742 controles completos = 761.
3. N1 ahora identifica por hash la línea v1.0 y fija propuesta, orden, duplicados, denominadores SMD, Moran I, pesos, KS, varianzas y diversidad de rutas.
4. Se declaró que N1 sólo calibra dentro del universo reconstruido y empareja resúmenes espaciales seleccionados; no reproduce toda la historia adaptativa ni preserva la estructura espacial completa.
5. C2 recibió gates separados y completos para similitud adyacente y similitud baja entre extremos.
6. Se fijó el dominio por modelo y bloque para SVD/regresión del proxy celular.
7. El nulo combinado D4 recibió endpoint, cola, regla `+1`, familia Holm y papel exclusivamente diagnóstico.
8. Se distinguieron fallos globales, fallos de la secuencia primaria y fallos locales de sensibilidad.
9. Wilson quedó definido como control auxiliar de precisión Monte Carlo, no como intervalo del efecto ni garantía simultánea.
10. Se registró la autorización para analizar el agregado BIGE y publicar resultados agregados no reconstructivos; se mantuvo la prohibición de redistribuir la matriz/exportaciones reconstructivas y se añadió un allowlist público archivo por archivo.
11. Se corrigió la sobreespecificidad anatómica externa: Siletti usa MoSR proxy, una disección VPL-labelled/targeted y S1C; no “Cu/Gr y VPL exactos”.
12. Se condicionó cualquier lenguaje de triangulación a los resultados efectivamente observados y se limitó BrainSpan a una sensibilidad modular no equivalente.
13. Se incorporó GSE3526 al ledger de contaminación con sus seis ramas conocidas, su interpretación como contraste relativo y su exclusión explícita de toda validación externa.
14. Se exigieron descarga y hash de la matriz oficial, entorno reproducible bloqueado, manifiestos relativos y auditoría GPL570 multiblanco para RAB3GAP2/AURKAPS1 y C7orf25/PSMA2.
15. Se registró la heterogeneidad pos hoc por sexo de la tríada GSE3526 como señal confundida con donante que exige sensibilidad consciente de metadatos y no admite una afirmación de efecto sexual.

## Bloqueantes antes de congelar y gate separado de liberación

| Bloqueante | Criterio de salida |
|---|---|
| Gobernanza de liberación BIGE — sólo liberación pública | Allowlist/exclusion audit archivo por archivo; exclusión demostrada de la matriz completa y de exportaciones reconstructivas. La autoridad analítica y para publicar resultados agregados ya está documentada; este punto no bloquea el análisis interno autorizado |
| Caché canónico | 324 lotes y todos los insumos reproducen hashes, identidades y cobertura esperada |
| Factibilidad sólo-covariables | N1 puede alcanzar aceptación/QC sin endpoint; los marcadores y matrices por bloque cumplen conteo/rango |
| Código | Ejecutable completo, revisión estática independiente y tests sintéticos de error tipo I/invariantes |
| Revisión final | Revisión estadística/metodológica independiente del JSON congelado y del ejecutable completo |
| Protocolo externo | Documento separado sellado; gate anatómico Siletti resuelto por metadatos antes de expresión |
| Sellado | Nuevo `protocol_v1_2.json`, ejecutable, entorno, inputs y declaración de ceguera con hashes |

## Auditoría de procedencia BIGE/GSE3526

La matriz puede describirse como un **agregado regional derivado de Body Index of Gene Expression (BIGE), mantenido internamente y no redistribuible**, sobre la plataforma Affymetrix Human Genome U133 Plus 2.0. La cadena documental es Lee et al. (FASEB J, 2005; doi:10.1096/fj.04-3552fje), Roth et al. (Neurogenetics, 2006; doi:10.1007/s10048-006-0032-6; GSE3526) y Ushach et al. (Clin Immunol, 2015; doi:10.1016/j.clim.2014.11.006), fuente de la descripción de BIGE como base con más de 105 células y tejidos humanos. GSE3526 es una cohorte/componente público de esa procedencia, no la demostración de que toda BIGE o toda la matriz interna equivalgan a una única serie GEO.

La reconstrucción GSE3526 contiene 353 arrays, 65 tejidos y 10 donantes; ocho donantes completan las comparaciones amplias, sin duplicados donante×tejido. Los endpoints usan 24/32 arrays. La serie contiene 193 arrays del universo neural en los diez donantes; 180 pertenecen a los ocho donantes completos y pueden influir en su z intradonante. Los valores depositados son RMA linearizado (4.538–33730.54); `log2(x+1)` es adecuado y `log2(x)` resultó casi idéntico. En medulla–thalamus–parietal, T fue 0.213706/0.221053/0.191338 para mediana/media/delegada; en DRG–medulla–thalamus–parietal fue 0.152193/0.165205/0.109722. Ninguna fracción Monte Carlo archivada o sign-flip fue `<0.05`. La convolución exacta del espacio de etiquetas de cuatro nodos produce 0.0578575/0.0487755/0.1311227; la rama media cruza 0.05 sólo por error Monte Carlo (0.0506095 frente a 0.0487755), por lo que queda prohibida toda narrativa de umbral. Para la tríada con mediana, las correlaciones medias fueron −0.022149, −0.028509 y −0.239035: T positivo refleja que los pares adyacentes son **menos negativos** que los extremos, no coexpresión positiva, continuidad o flujo. En cuatro nodos sólo DRG–medulla fue positiva (~0.203); las demás aristas seriales fueron pequeñas o negativas.

El resultado depende materialmente del universo usado para estandarizar. Para mediana/media/delegada, T de tres/cuatro nodos fue 0.243092/0.150073, 0.241776/0.172588 y 0.199890/0.102924 con 20 tejidos neurales comunes; 0.099123/0.045249, 0.101316/0.054678 y 0.055702/−0.005994 con los 65 tejidos; y 0.003399/0.010526, −0.001645/0.008114 y 0.003509/0.000512 sin z. Debe presentarse como contraste condicionado al fondo neural congelado, no como similitud regional general. La sensibilidad de universo de fondo es obligatoria y no puede usarse para escoger retrospectivamente la normalización favorable.

La tríada también mostró separación completa pos hoc por sexo registrado (n=4 mujeres, n=4 hombres; T mujer/hombre 0.563/−0.136, 0.569/−0.127 y 0.529/−0.146; partición exacta bilateral descriptiva 2/70=0.0286), ausente en cuatro nodos (~0.14 frente a ~0.16). Esto está confundido con donante y no sustenta biología sexual. No depende sólo de RAI2, el único gen ligado al X del panel: al omitir RAI2, las diferencias mujer−hombre fueron 0.598/0.590/0.560, y todo leave-one-gene conservó diferencia positiva (0.549–0.768); C7orf25 y RAI2 fueron los reductores más fuertes, pero excluir ambos aún dejó 0.422–0.490. Sólo están disponibles edad y sexo; faltan causa de muerte, PMI, RIN y lote. Todo ello es diagnóstico pos hoc y sólo obliga a reportar heterogeneidad y congelar una regla consciente de metadatos antes de análisis adicionales.

El paquete corregido `gse3526_donor_reconstruction_v2_A02.zip` implementa descarga con verificación de 85,860,948 bytes y SHA-256 `77cc18cb606adb8028e0ef8e84db288f26b160308a364089c3876c0335c4b7d0`, dependencias y entorno BLAS/LAPACK/hilos registrados en archivos legibles por máquina, rutas relativas, dos reejecuciones con 22/22 outputs científicos idénticos byte por byte y convolución exacta del espacio finito de etiquetas. También archiva el mapa GPL570 inspeccionado y ejecuta una sensibilidad post hoc de 17 genes que excluye RAB3GAP2 y C7orf25; la dirección persiste sin resolver la atribución génica ni la circularidad. El ZIP tiene 46 archivos, 1,199,750 bytes y SHA-256 `4edd3d126023e730ae8c93bcf6f74bb193237524429c20d5108e63487ffadfae`, y omite la matriz GEO. Un fallo futuro de estos gates es local a esta sensibilidad y no puede ocultarse.

## Decisión externa

La búsqueda documentada no identificó un atlas adulto humano que se haya mantenido fuera de todo análisis previo y sea, a la vez, anatómicamente exacto, multidonante y completo para Cu/Gr–VPL–S1/postcentral.

- **Siletti/Human Brain Cell Atlas:** mejor proxy de tres compartimentos y ensayo independiente de AHBA, pero ya inspeccionado a resolución amplia y con una sola tríada de mayor especificidad; no es réplica con datos mantenidos fuera de análisis previos ni poblacional.
- **FANTOM5:** dos tríadas amplias medulla–thalamus–parietal en CAGE; útil sólo con agregación promotor-gen congelada y concordancia prespecificada.
- **BrainSpan:** carece de médula y usa MD, no VPL; máximo papel de sensibilidad modular talamocortical no equivalente.
- **GSE3526:** excluido del conjunto externo por pertenecer al linaje fuente histórico documentado y por tener endpoints ya inspeccionados; sólo auditoría/reconstrucción y sensibilidad postselección del mismo linaje. La equivalencia numérica con el agregado BIGE interno no está establecida.

Los resultados de estos recursos deben permanecer separados. Se prohíbe un meta-p entre anatomías, plataformas o endpoints no equivalentes.

## Límite editorial

El artículo tiene `GO` condicionado como estudio posdescubrimiento transparente centrado en la dependencia del modelo y el fallo del doble gate. Tiene `NO-GO` para presentarse como validación biológica fuerte, especificidad de DCML, conectividad, dirección, mecanismo de fibromialgia/dolor o selección de un blanco de estimulación.

## Gobernanza del paquete v1.1

El paquete canónico `DCML19_transcriptomic_topology_v1_1_A04.zip` (25,644,708 bytes; SHA-256 `bedffc8837d0880dfae263e38df97e03370fc63243aa9d4abca1c6c26c05b357`) pasó auditoría interna de integridad: ZIP válido, 104/104 entradas del manifiesto verificadas y manuscrito/informe idénticos a sus copias empaquetadas. Su manifiesto interno `MANIFEST_SHA256.txt` tiene SHA-256 `1f883ae6ed841f3a88ce0f9b1a6f27ab3c653e4d276a3915d2b5237963a2d1cd`. No contiene los cuatro insumos privados conocidos. Sí contiene resúmenes derivados y ledgers con información de procedencia privada; por ello permanece clasificado como paquete científico/auditor interno hasta pasar el allowlist/exclusion audit archivo por archivo. Esto no contradice la autoridad ya registrada para publicar resultados agregados no reconstructivos.
