## Supplementary figures and images for "An anatomy-constrained transcriptomic screen identifies a 19-gene panel with an ordered regional similarity pattern in the human somatosensory system"

### figure_1_observed_donor_structure_v1_0.png

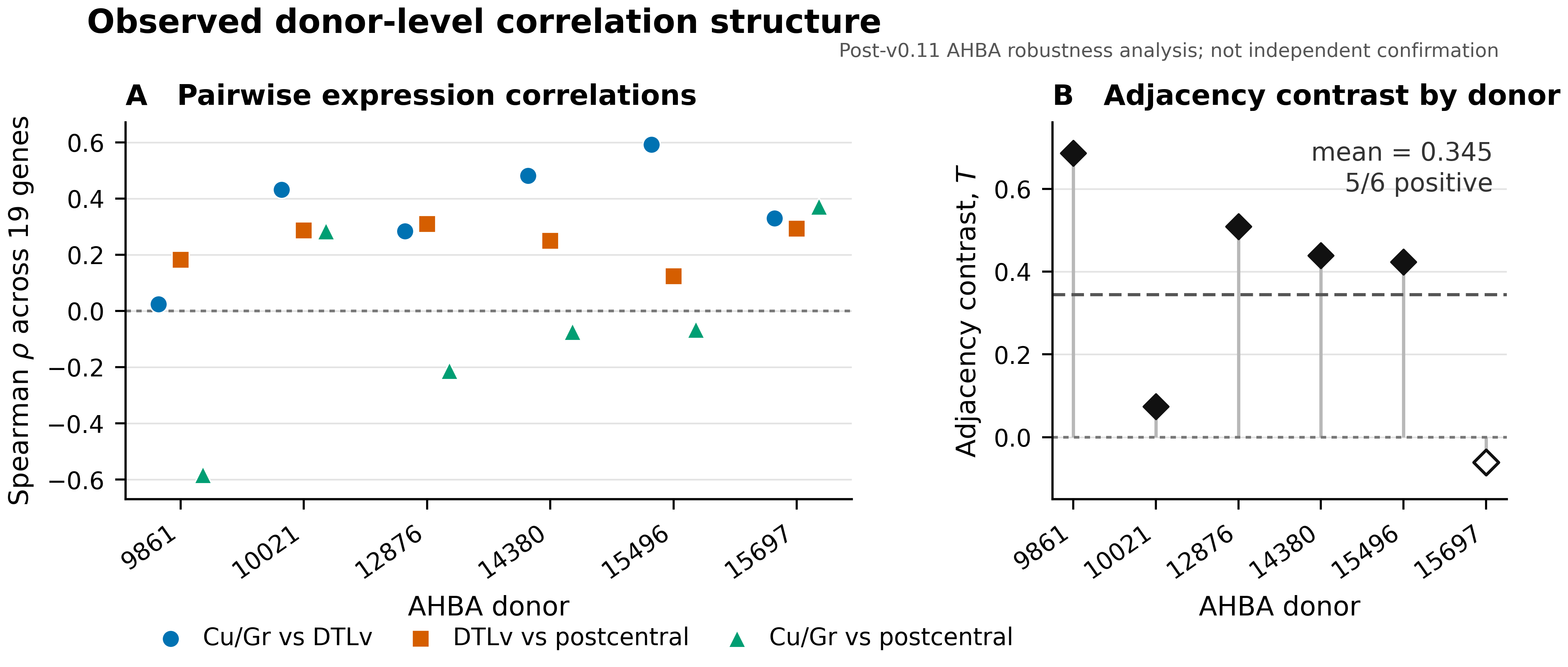

### figure_1_spatial_double_gate_v1_1.pdf

#### Frozen spatial-null double gate and sensitivity models

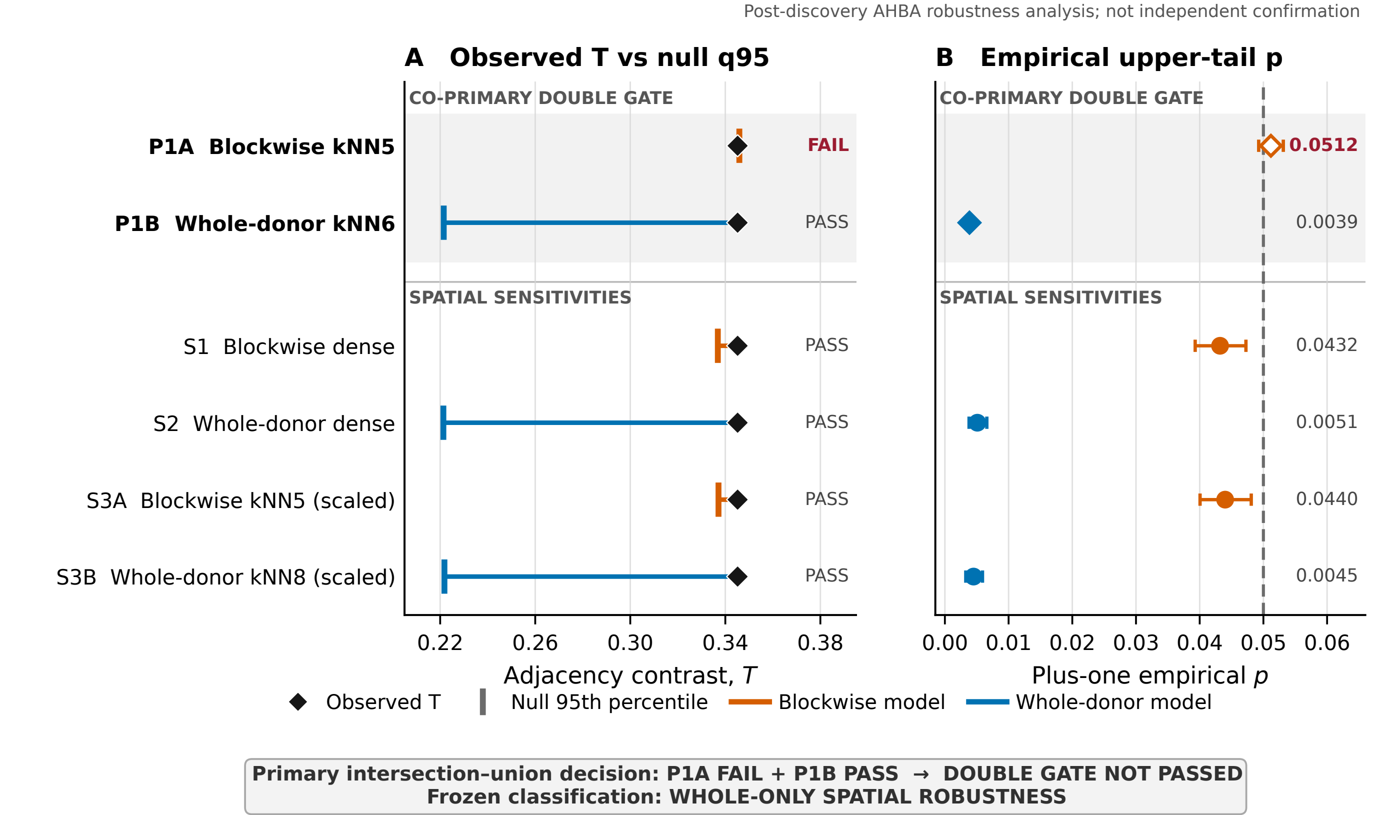

### figure_1_spatial_double_gate_v1_1.png

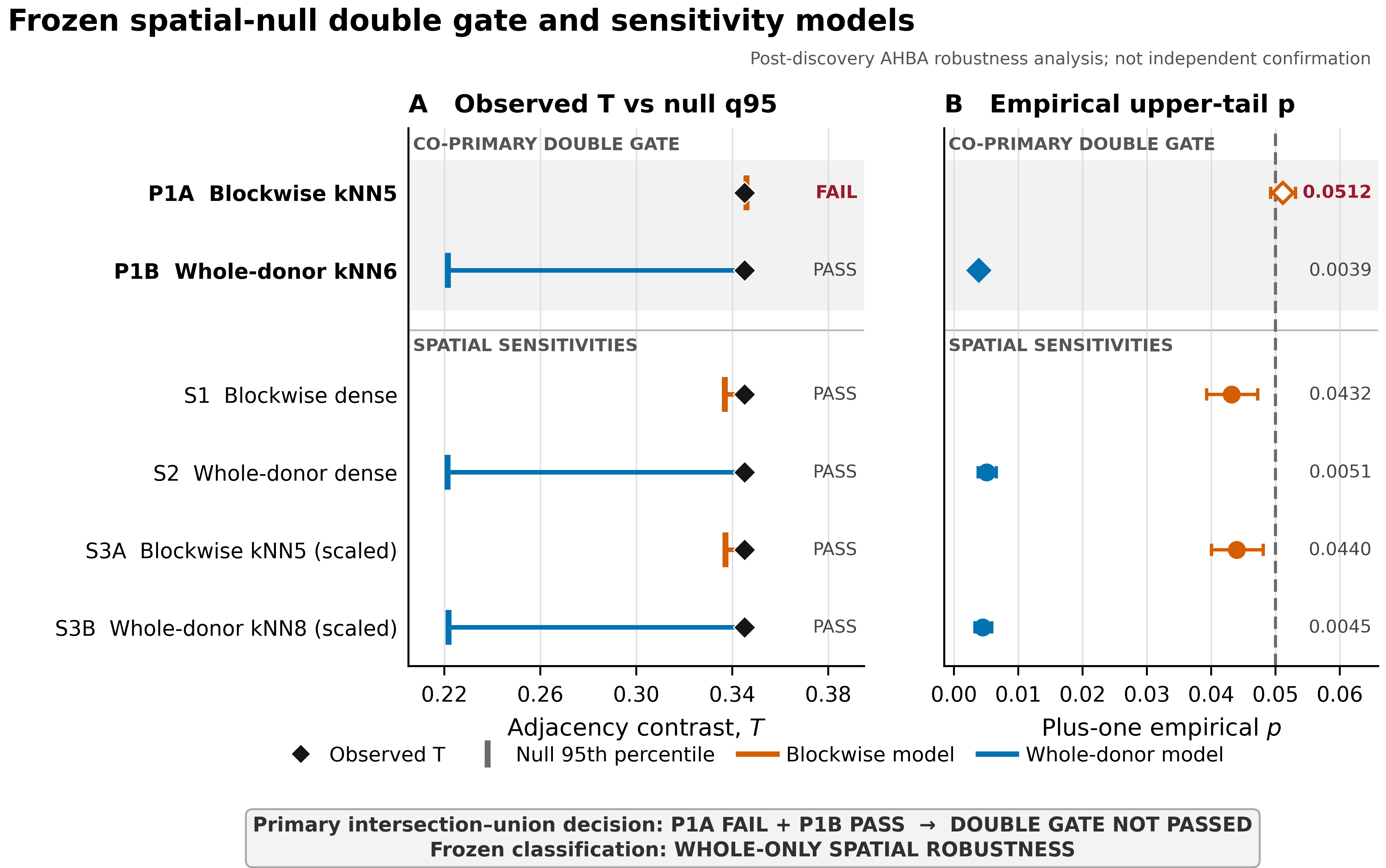

### figure_2_conditional_nulls_v1_0.png

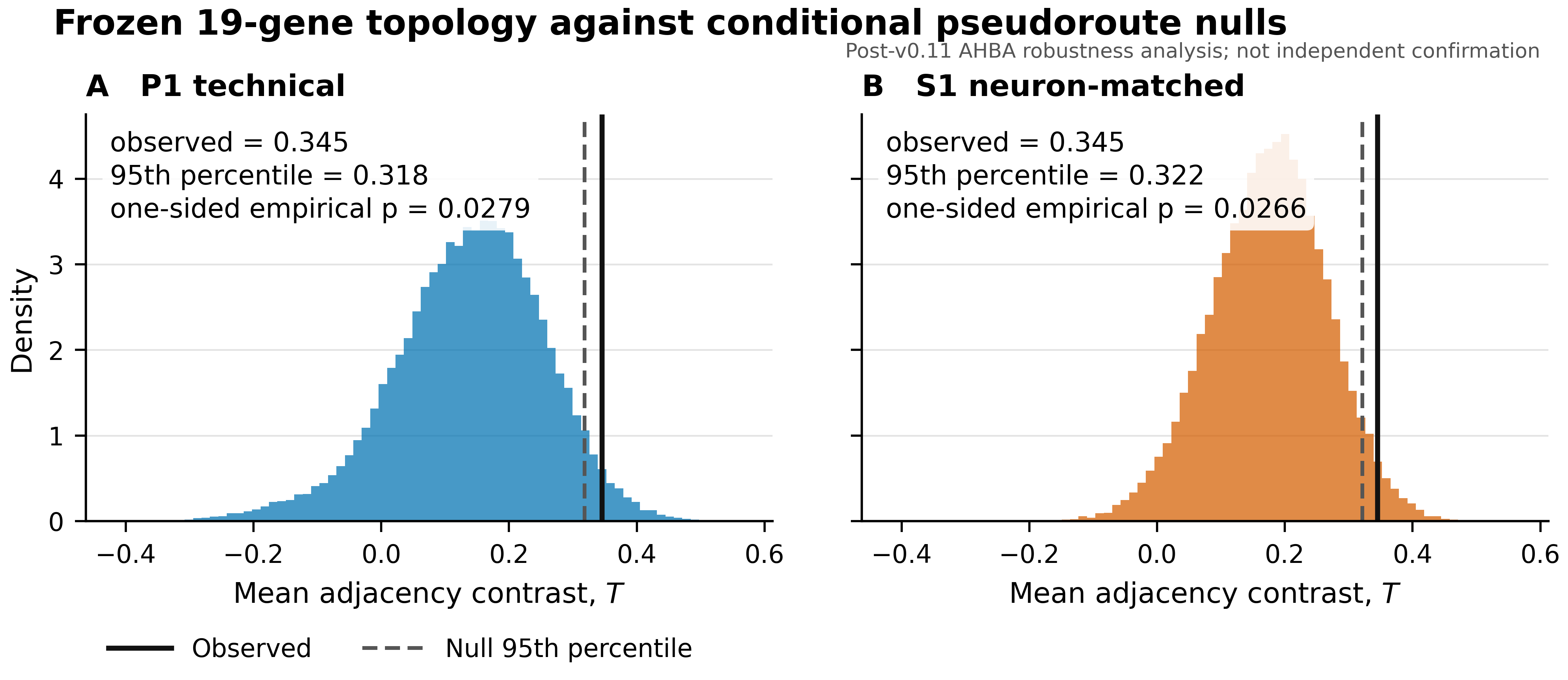

### figure_2_spatial_lodo_v1_1.pdf

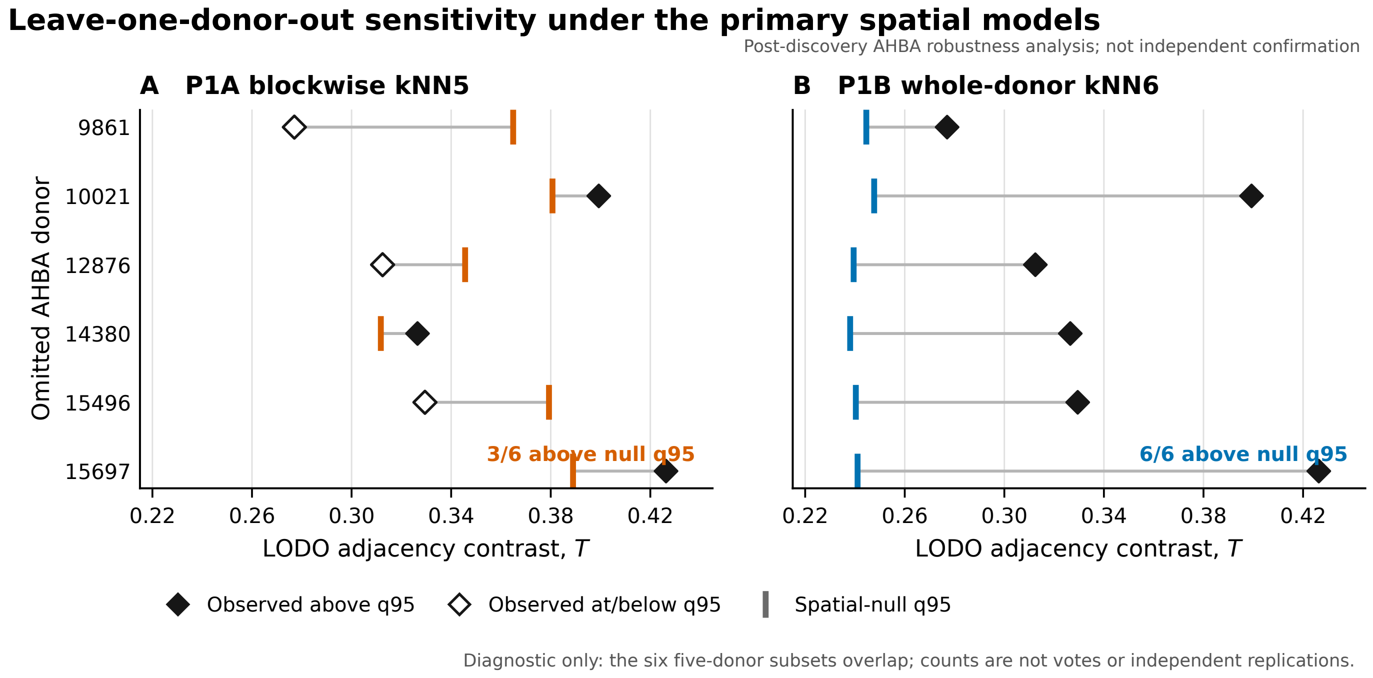

### figure_2_spatial_lodo_v1_1.png

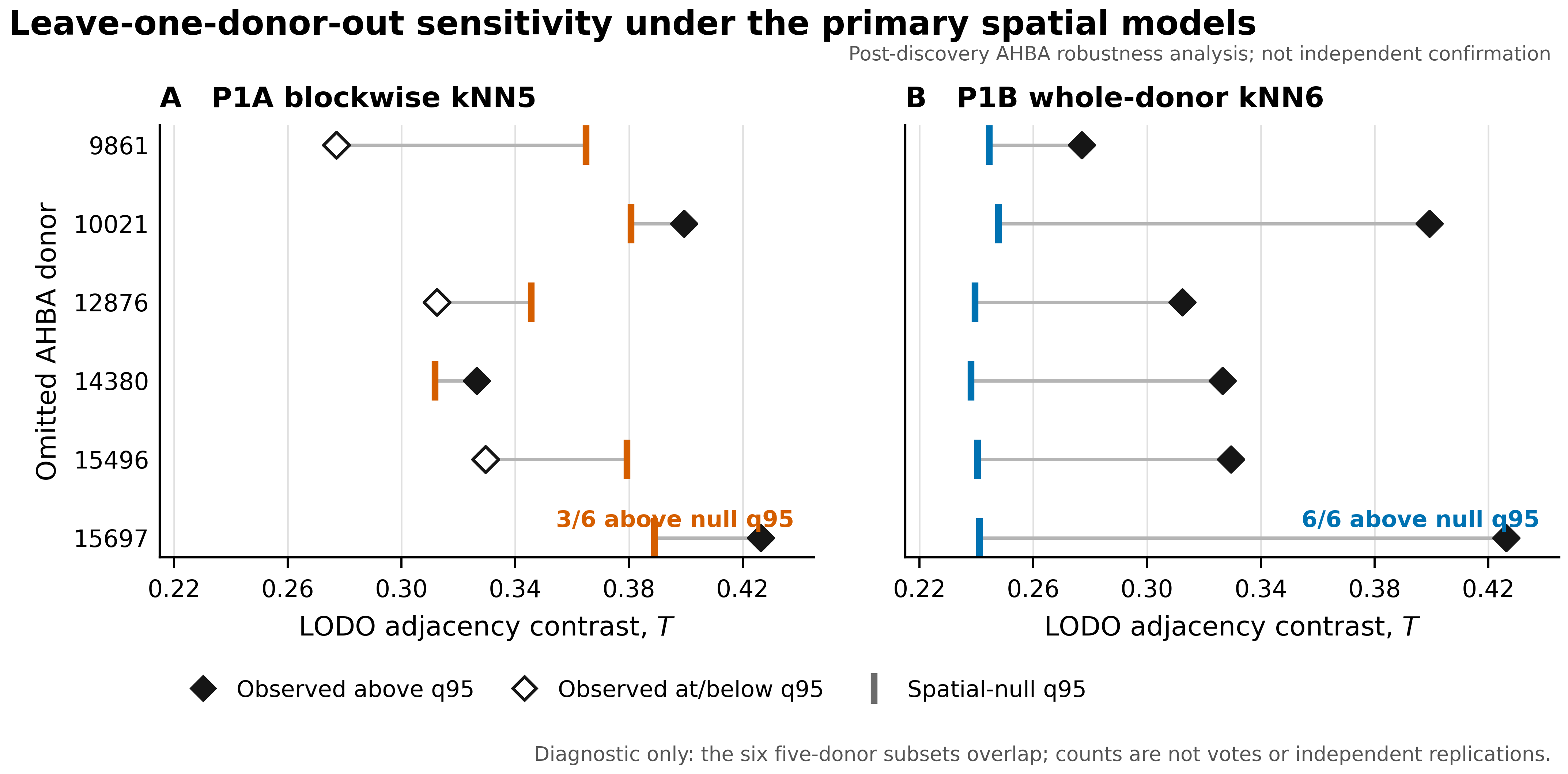

### figure_3_component_decomposition_v1_0.png

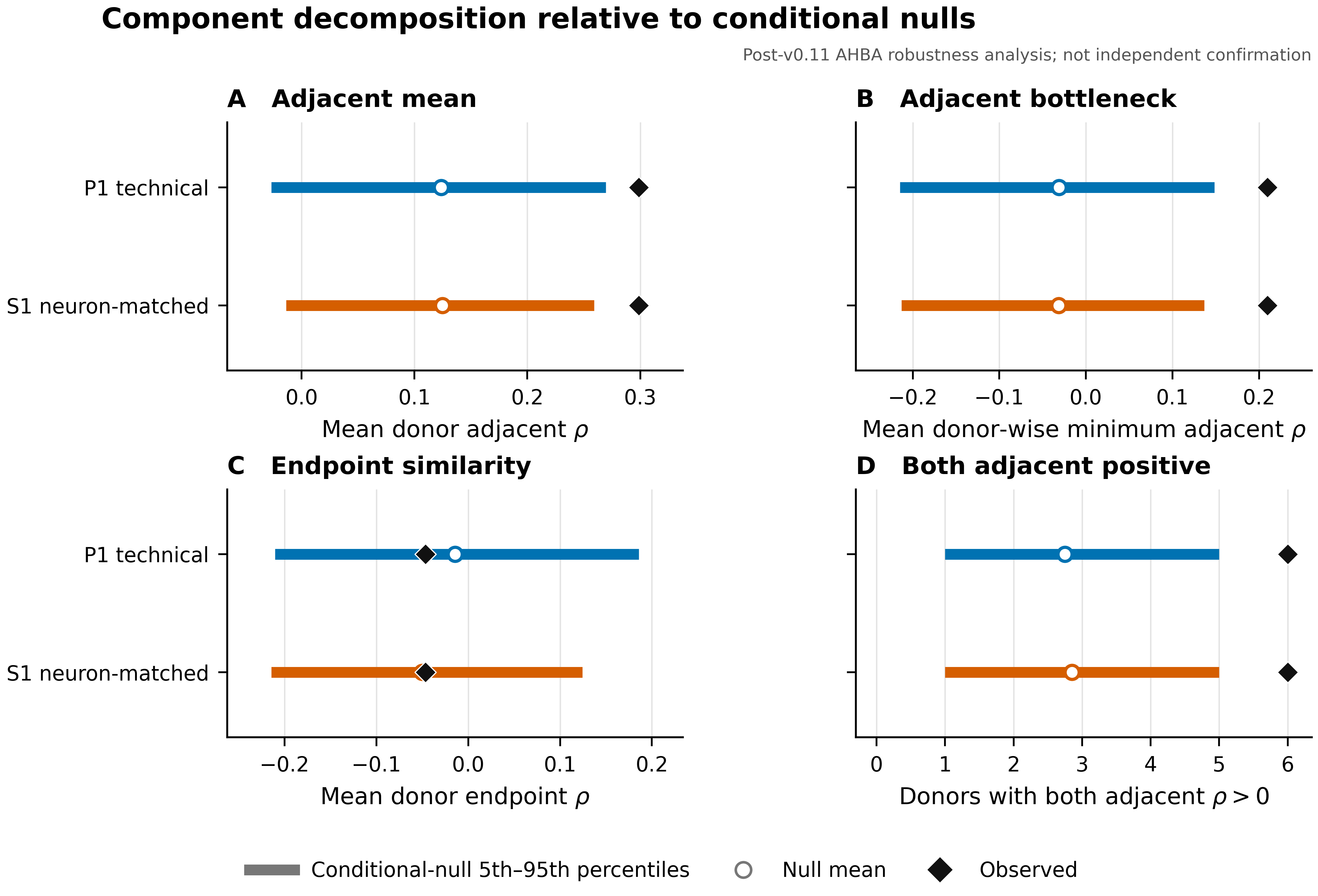

### figure_4_lodo_sensitivity_v1_0.pdf

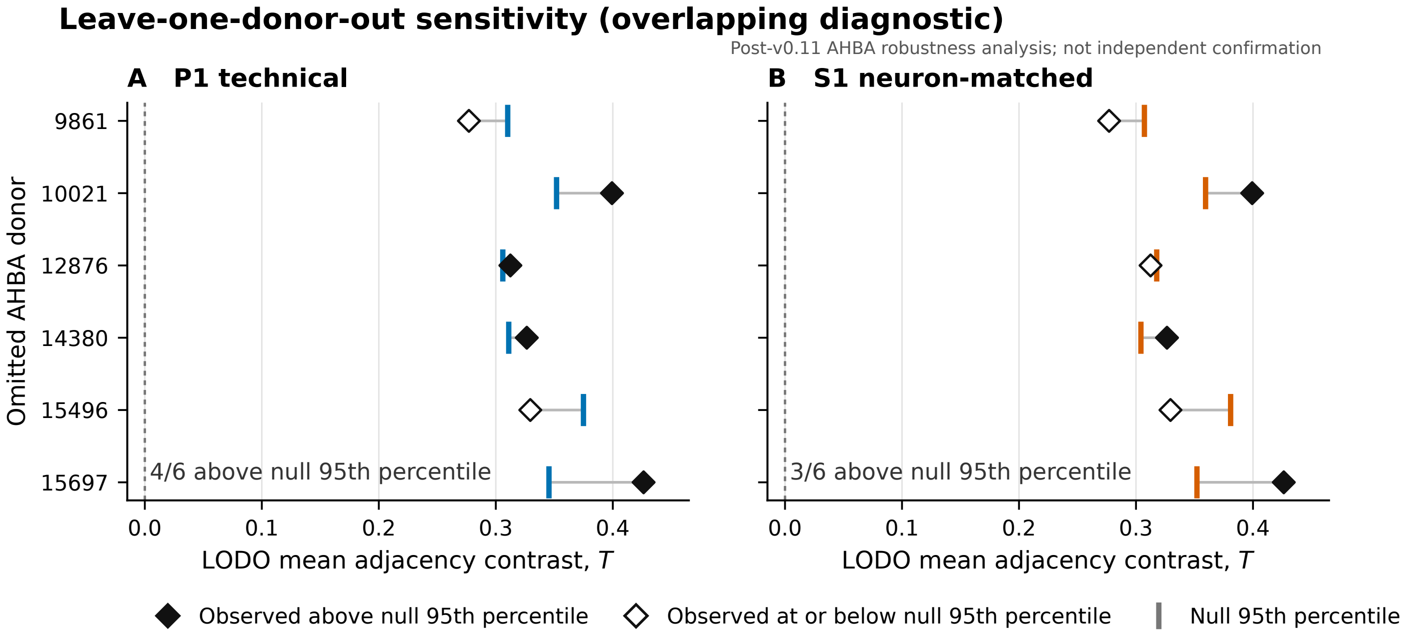

### figure_4_lodo_sensitivity_v1_0.png

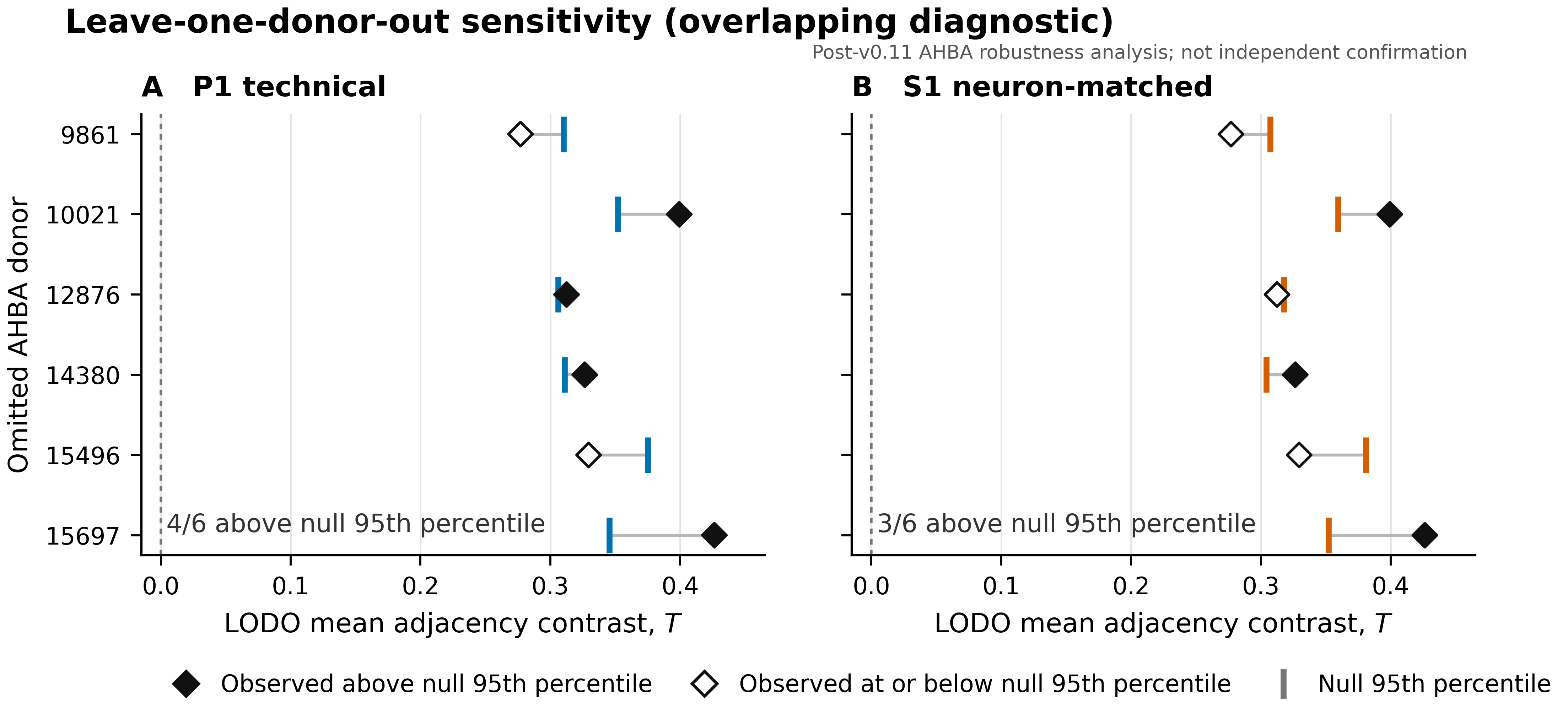

### figure_S_gse3526_donor_heterogeneity_v1.png

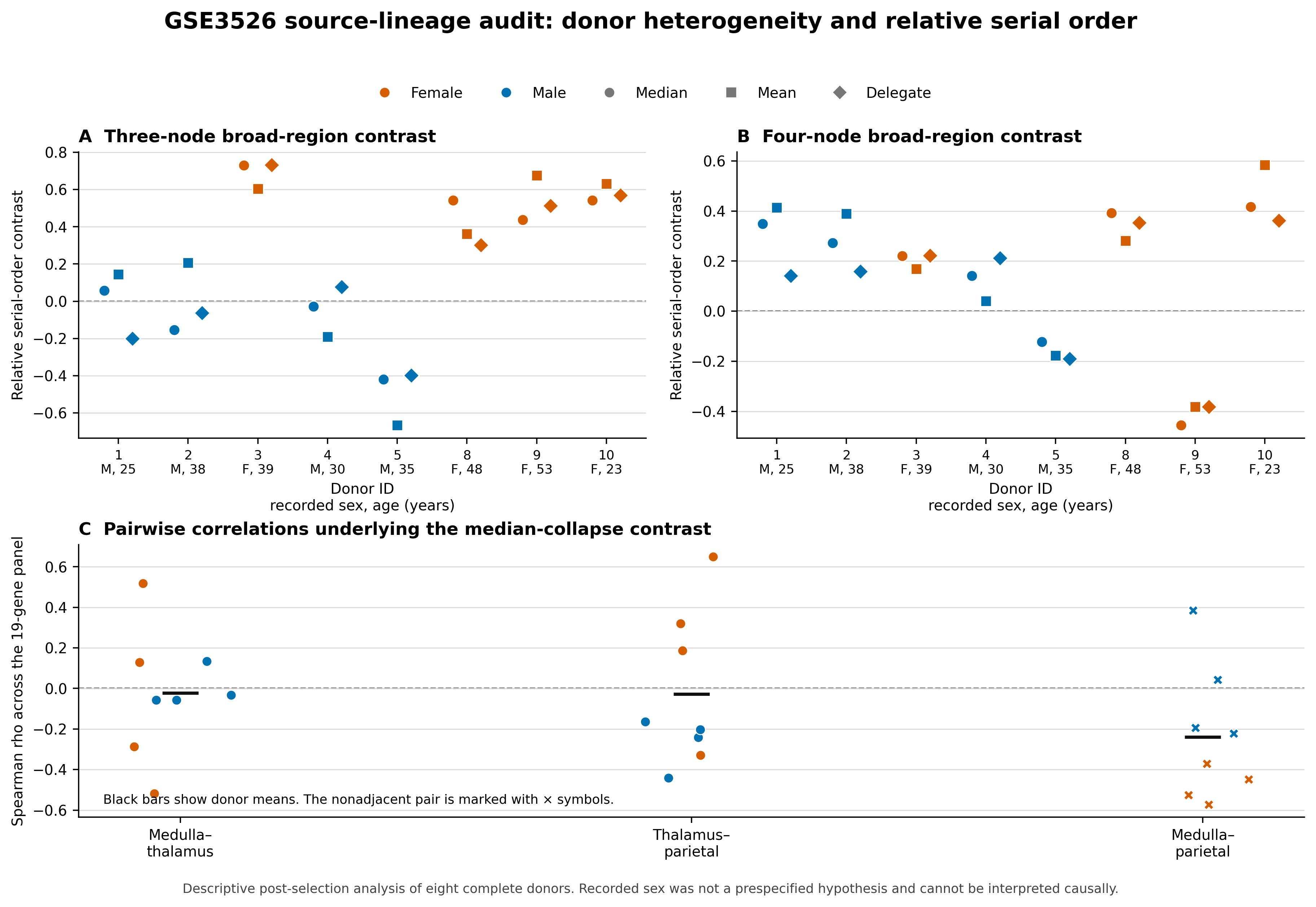
